# Convergent evolution of resistance pathways during early stage breast cancer treatment with combination cell cycle (CDK) and endocrine inhibitors

**DOI:** 10.1101/2021.01.19.427299

**Authors:** Jason I. Griffiths, Jinfeng Chen, Patrick A. Cosgrove, Anne O’Dea, Priyanka Sharma, Cynthia Ma, Meghna Trivedi, Kevin Kalinsky, Kari B. Wisinski, Ruth O’Regan, Issam Makhoul, Laura M. Spring, Aditya Bardia, Frederick R. Adler, Adam L. Cohen, Jeffrey T. Chang, Qamar J. Khan, Andrea H. Bild

**Affiliations:** Department of Medical Oncology & Therapeutics Research, City of Hope National Medical Center, 1500 East Duarte Road, Duarte, CA, 91010, USA; Department of Mathematics, University of Utah 155 South 1400 East, Salt Lake City, UT, 84112, USA; Division of Medical Oncology, University of Kansas Medical Center, Westwood, KS, 66160, USA; Division of Oncology, Washington University School of Medicine, St. Louis, MO, 63130, USA; Department of Medicine, Columbia University Irving Medical Center, NY, 10032, USA; Department of Medicine, University of Wisconsin School of Medicine and Public Health, Carbone Cancer Center, WI, 53726, USA; Division of Internal Medical Oncology, University of Arkansas for Medical Sciences, AR, 72205, USA; Department of Medicine, Massachusetts General Hospital Cancer Center and Harvard Medical School, MA, 02114, USA; School of Biological Sciences, University of Utah 257 South 1400 East, Salt Lake City, UT, 84112, USA; Department of Internal Medicine, Huntsman Cancer Institute, University of Utah, Salt Lake City, UT 84112, USA; Department of Integrative Biology and Pharmacology, School of Medicine, School of Biomedical Informatics, UT Health Sciences Center at Houston, Houston, TX, 77030, USA

## Abstract

Combining cyclin-dependent kinase (CDK) inhibitors with endocrine therapy improves outcomes for metastatic estrogen receptor positive (ER+), HER2 negative, breast cancer patients. However, the value of this combination in potentially curable earlier stage patients is not clear. Using single cell transcriptomic profiling, we examined the evolutionary trajectories of early stage breast cancer tumors using serial tumor biopsies from a clinical trial of preoperative endocrine therapy (letrozole) alone or in combination with the cell cycle inhibitor ribociclib. Applying hierarchical regression and Gaussian process mathematical modelling, we classified each tumor by whether it shrinks or persists with therapy and determined cancer phenotypes related to evolution of resistance and cell cycle transcriptional rewiring. We found that all patients’ tumors undergo subclonal evolution during therapy, irrespective of the clinical response. However, tumors subjected to endocrine therapy alone showed reduced diversity over time, while those facing combination therapy exhibited increased diversity. Despite different subclonal diversity, single nuclei RNA sequencing uncovered common phenotypic changes in tumor cells that persist following treatment. In these tumors, cancer cells with accelerated loss of estrogen signaling have convergent up-regulation of the JNK pathway, while cells that maintain estrogen signaling during therapy show potentiation of CDK4/6 activation consistent with ERBB4 and ERK signaling up-regulation. These convergent phenotypes were associated with growing tumors resistant to combination therapy. Cell cycle reconstruction identified that these tumors can rebound during combination therapy treatment, indicating stronger selection and promotion of a proliferative state. These results indicate that combination therapy in early stage ER+ breast cancers with ER and CDK inhibition drives rapid evolution of resistance via a shift from estrogen signaling to alternative growth factor receptor mediated proliferation and JNK signaling activation, concordant with a bypass in the G1 checkpoint.

## Introduction

Hormone receptor positive (estrogen receptor positive (ER+) and/or progesterone receptor positive (PR+)) breast cancer comprises 70-80% of all breast cancers (1). In ER+ breast cancer, estrogen receptors are activated by estrogen and transduction of this signal to the nucleus promotes cell proliferation and tumor growth. The primary treatment for ER+ breast cancer is endocrine therapy, which either depletes endogenous estrogen along with estrogen made by breast cancer cells using aromatase-inhibition (AI) or blocks ER activity through direct modulation or degradation (2-4). Approximately 90% of all patients with metastatic breast cancer eventually develop resistance to endocrine therapy and at least 33% of patients with early-stage disease will develop endocrine resistance (5, 6). Combination of endocrine therapy with cyclin-dependent kinase (CDK) 4/6 inhibitors has improved disease control in metastatic ER+ breast cancer and adjuvant trials to study the efficacy of this combination in early stage, non-metastatic breast cancer are ongoing (NATALEE) or have completed accrual and are in the follow-up phase (monarchE, PENELOPE-B and PALLAS) (7-10). CDK4/6 inhibitors combined with endocrine therapy are associated with a significant decrease in expression of the proliferation marker Ki-67 compared to endocrine therapy alone (11), resulting in a higher rate of complete cell cycle arrest in the neoadjuvant treatment of ER+ breast cancer, but the clinical significance of the reduction in proliferation from these combinations is not known (7-9). Preliminary results after short term follow-up from adjuvant trials have shown contradictory results, with the MonachE trial showing an improvement in invasive disease free survival with two years of adjuvant abemaciclib added to endocrine therapy, particularly in people whose tumors have high Ki-67 (12), while the PALLAS trial failed to show an improvement in the same endpoint with two years of adjuvant palbocilib (13). It is not known to what extent such differences are due to biologic effects of the drugs versus biologic differences in different populations of early breast cancer patients; therefore, additional research is needed to characterize the effects of CDK4/6 inhibitor and mechanisms of resistance in surviving cells.

CDK4/6 proteins form a complex with cyclin D that phosphorylates and deactivates the key cell cycle checkpoint regulator retinoblastoma protein (RB1), leading to E2F transcription factor activation and production of cell cycle promoting genes and progression from G1 to S phase of the cell cycle (14). In the context of breast cancer, binding of estrogen to ER and of growth factors binding to growth factor receptors (GFR) drive proliferation through cyclin D/CDK4/6 activation (15, 16). ER can also activate extracellular signal-regulated kinase (ERK) mitogen-activated protein kinase (MAPK) signaling and drive transcription of cyclin D genes and cell cycle progression (17). Some known mechanisms of endocrine resistance include loss or modification of the key estrogen receptor ESR1 and activation of the phosphoinositide 3-kinases (PI3K) or epidermal growth factor receptor (EGFR) pathways (18). Prior studies have also revealed mechanisms underlying resistance to CDK4/6 inhibitor treatment in the metastatic setting (19). These include disruption or loss of CDK6 and cyclin E2 (CCNE2), as well as activation of AKT1, RAS, ERBB2, and FGFR genes. The impact of treatment with these agents on the phenotypes and emergence of resistance in early, non-metastatic breast tumors remains unknown.

The phosphorylation of the human ER serine residue at position 118 is required for its full activity. Ser118 is phosphorylated by MAPK, specifically MAPK1/3 (ERK1/2), initiated by GFR activation (20). Likewise, estrogens also activate ERK1/2 through multiple signaling pathways, further highlighting the crosstalk among these pathways (21, 22). Additional MAPK pathways, including MAPK8-10 (Jun amino-terminal kinases; JNK1-3) and MAPK11-14 (p38α−γ), have also been shown to interact with ER signaling, but their role in response to therapy is unknown. Further, the role for all MAPK pathways in evolution of tumor cells to a resistant state has not been defined in tumors during therapy with endocrine and/or cell cycle inhibition.

To address the questions detailed above including the impact of therapy on signaling and response in early stage ER+ breast cancer, we addressed how resistance evolves in response to endocrine and cell cycle inhibitor therapies in early stage ER+ breast cancers. These analyses demonstrate multiple convergent phenotypes conveying resistance to combination endocrine and cell cycle inhibition therapy in early stage ER+ breast cancer.

## Results

### Clinical trial: Patient treatment and sample collection

We studied the evolution of endocrine and CDK inhibitor resistant cancer cell genotypes and phenotypes in post-menopausal women with node positive or >2 cm ER and or PR+, HER2 negative breast cancer enrolled in the FELINE trial (clinicaltrials.gov # NCT02712723) (23). This trial evaluated whether the addition of CDK inhibition to endocrine therapy in the neo-adjuvant setting improved the preoperative endocrine prognostic index (PEPI) as well as promotes sustained cell cycle arrest. Patients (n=120) were randomized equally into three arms: A) endocrine therapy alone (letrozole plus placebo), B) intermittent high dose combination therapy (letrozole plus ribociclib (600 mg/d, three on/one week off)) or C) continuous lower dose combination therapy (letrozole plus ribociclib (400 mg/d)) (Figure 1a). Patients were treated for six cycles (180 days) and biopsies were collected at baseline (day 0), following treatment initiation (day 14), and end of treatment (surgery around day 180).

**Figure 1.**
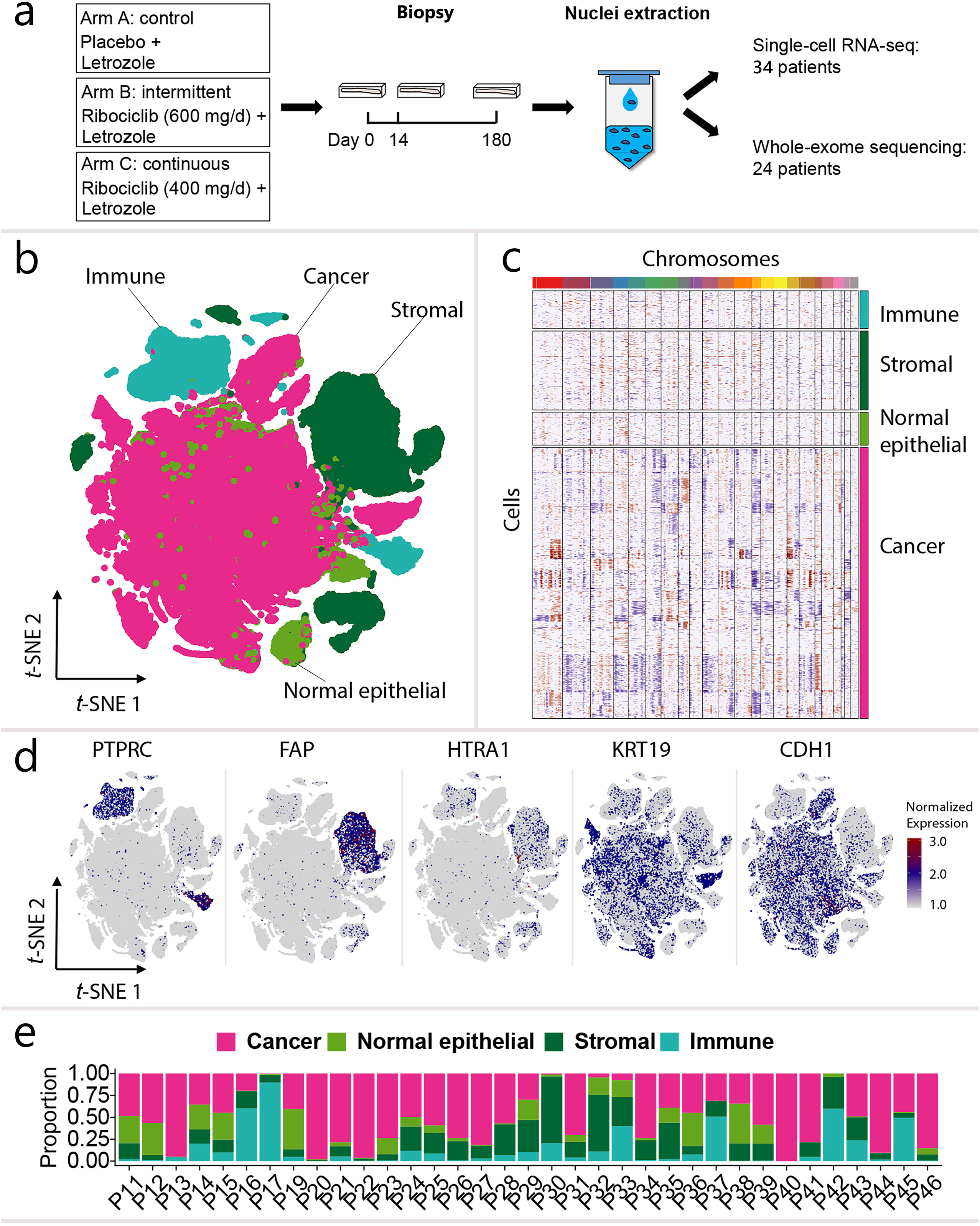
Landscape of tumor and macroenvironment of early stage ER+ breast cancer patients in FELINE trial. **a**, Schematic diagram of single-cell RNA-seq workflow. The data of 35 patients generated using the 10x genomics platform are shown in Fig. I b-d. The data of 10 patients generated using ICELLS platform are shown in Supplementary Fig. I. **b**, Distinction of different cell types is shown by a *t*-SNE plot of single cells of 35 ER+ breast cancer patient tumors (color=cell type). **c**, Gene copy number profile in cancer cells and neighboring normal cells. Blue color indicates copy number loss and red color indicates copy number gain. **d**, Expression of marker genes of cancer cells and normal epithelial cells (KRT19, CDHI), stromal cells (FAP, HTRAI), and immune cells (PTPRC). **e**, Proportion of cancer cells and neighboring normal cells in each patient across all timepoints.

We used tumor growth measurements over time to define resistance and sensitivity during the six months of therapy. Specifically, we mathematically reconstructed tumor size continuously over time during treatment using data from magnetic resonance imaging (MRI), ultrasound (US), mammogram (MG), clinical physical assessment (CA) taken throughout therapy, and the surgical pathology (SP) (24) observation. This estimation of tumor size over time was determine using a Gaussian Process Latent Variable Model to account for known biases and differences in accuracy between measurement modalities (25, 26). Groups of similar tumor trajectories were then identified using a Gaussian mixture model. Patients exhibiting either a sustained shrinkage in tumor size during treatment or an initial shrinkage followed by a plateau during treatment were classified as sensitive to therapy. Alternatively, tumors classified as resistant show either 1) no change in size during treatment, 2) a rebound indicated by an initial tumor shrinkage followed by growth during treatment, or 3) continual increased growth (Supplementary Figure 1a-b). Tumors defined as resistant had a significantly higher proportion of tumor remaining after therapy (>2/3 initial size) compared to sensitive tumors (t=4.45, p<0.001), and have a significant correlation between clinical and modeling classifications (Supplementary Figures 1a-c). All patient modeling classifications match RECIST 1.1 classification except for two patients. One tumor classified as resistant by the model but PR by RECIST 1.1 shrunk on imaging and clinical exam at day 90 but then both pathology and ultrasound show a rebound by day 180 following the emergence of resistance. The second tumor classified as sensitive by the model but marginally SD by RECIST 1.1 exhibited a steady decrease in tumor size when looking across all imaging data from day 0-180, resulting in 30% decrease in size at surgery.

Biopsies from the first 60 patients were processed and analyzed (20 patients from each treatment arm). The second set of 60 patients’ tumors is to be used as a validation cohort and was withheld. From 60 total patients, 45 had sufficient high-quality tissue in optimal cutting temperature compound (OCT) at day 0 and a follow-up time point. Serial single-cell RNA-sequencing profiles (scRNAseq) and whole exome sequencing (WES) pre- and post-treatment was performed as detailed in the methods (Figure 1a), with scRNAseq prioritized. From the 45 patients sampled, 34 had a sufficient number of cancer cells and high-quality sequencing data for scRNAseq analysis of the progression of tumor RNA phenotypes and 24 patients yielded WES data (Figure 1e shows patient samples processed on the 10x and Supplementary Table 1 shows patient samples processed using the ICELL8). For samples with sufficient DNA, WES (mean depth 234x) was performed on pre- and day 180 post-treatment tumor biopsies for 24 patients. Matched blood samples were sequenced in parallel (mean depth 230x) to identify somatic mutations (Supplementary Table 2).

### Genomic analysis of patient tumor samples

We obtained scRNAseq transcriptional profiles for 176,644 cells after filtering out low-quality cells and doublets (Supplementary Table 3). To correct single cell gene expression for differences in read depth between cells, counts were normalized using a zero-inflated negative binomial model (zinbwave) (27). Cells across patients were integrated using the Seurat normalization package and following the reciprocal PCA method (28). Cancer cells were first distinguished from normal cells by performing gene copy number analysis of the scRNAseq data using inferCNV for each single cell (29) (Supplementary Dataset 2). The inferCNV algorithm predicts the copy number profiles of each cell from its transcriptional profile. As shown in Figure 1c and Supplemental Figure 2b, some cells have frequent and pronounced changes in copy numbers, reflective of copy number alterations, and were therefore classified as cancer cells, while others show no copy number changes and are classified as normal cells. We confirmed this classification by projecting the transcriptomic profiles of cells into low dimensional space using t-SNE and found that cells with copy number alterations clustered together; and those that show no copy number alterations also cluster with themselves (30). Finally, marker gene expression shown in Figure 1d was then used to verify the cell type annotations. In sum, the cells were distinguished based on frequency of predicted copy number alterations, similarity in overall gene expression profiles, and expression of known cancer or normal cell markers. A total of 32,781 (18.56%) stromal cells and 16,672 (9.44%) immune cells were identified, using singleR cell type annotation and verified by cell type specific marker gene expression (Figures 1b and 1c, Supplementary Figure 2, and Supplementary Table 3) (31). Immune and stromal cells were clearly identifiable by expression of PTPRC and FAP/HTRA1, while cancer and normal epithelial cells expressed KRT19 and/or E-cadherin (CDH1) (Figure 1d).

### Tumors undergo subclonal evolution during treatment

To understand how selective pressures of endocrine and CDK4/6 inhibitors drove evolution of cancer cells, WES data was analyzed as detailed in the methods. On average, 99 non-synonymous mutations (range 7-916) and 89 indels (range 24-380) were detected in each sample (Supplementary Table 4). Two patients had substantially higher mutation burden compared to average (P21: 224-471 non-synonymous mutations vs. mean: 99 and P45: 912-916 vs. mean: 99, Supplementary Table 4) and shared a distinct mutational signature enriched in APOBEC signatures 2 and 13 (Supplementary Figure 3). Gene mutations known to be frequent in ER+ breast cancer were seen, including PIK3CA (46%), TP53 (29%), and MAP3K1 (21%) (Figure 2a). Gene copy number alterations (CNA) were also frequently identified in the WES data, including gains in AKT3, CCND1, CCNE2, CDK6, FGFR1 and losses in ESR1, RB1 and TP53 (Figure 2b). In general, copy number alterations were more frequent in resistant than sensitive tumors, with most present prior to therapy (Figure 2b, Supplementary Table 5). When summarized at the pathway level, cell cycle, TGF-beta, and TP53 pathway related genes were frequently mutated in this cohort, with no difference in occurrence rate between patients given endocrine alone versus combination therapy (Supplementary Figure 4 and Supplementary Table 6). Over time, allele frequency of variants in 19 genes (PTEN, GATA3, and others) increase in 10 patients, suggesting enrichment of clones carrying those mutations (Supplementary Table 7).

**Figure 2.**
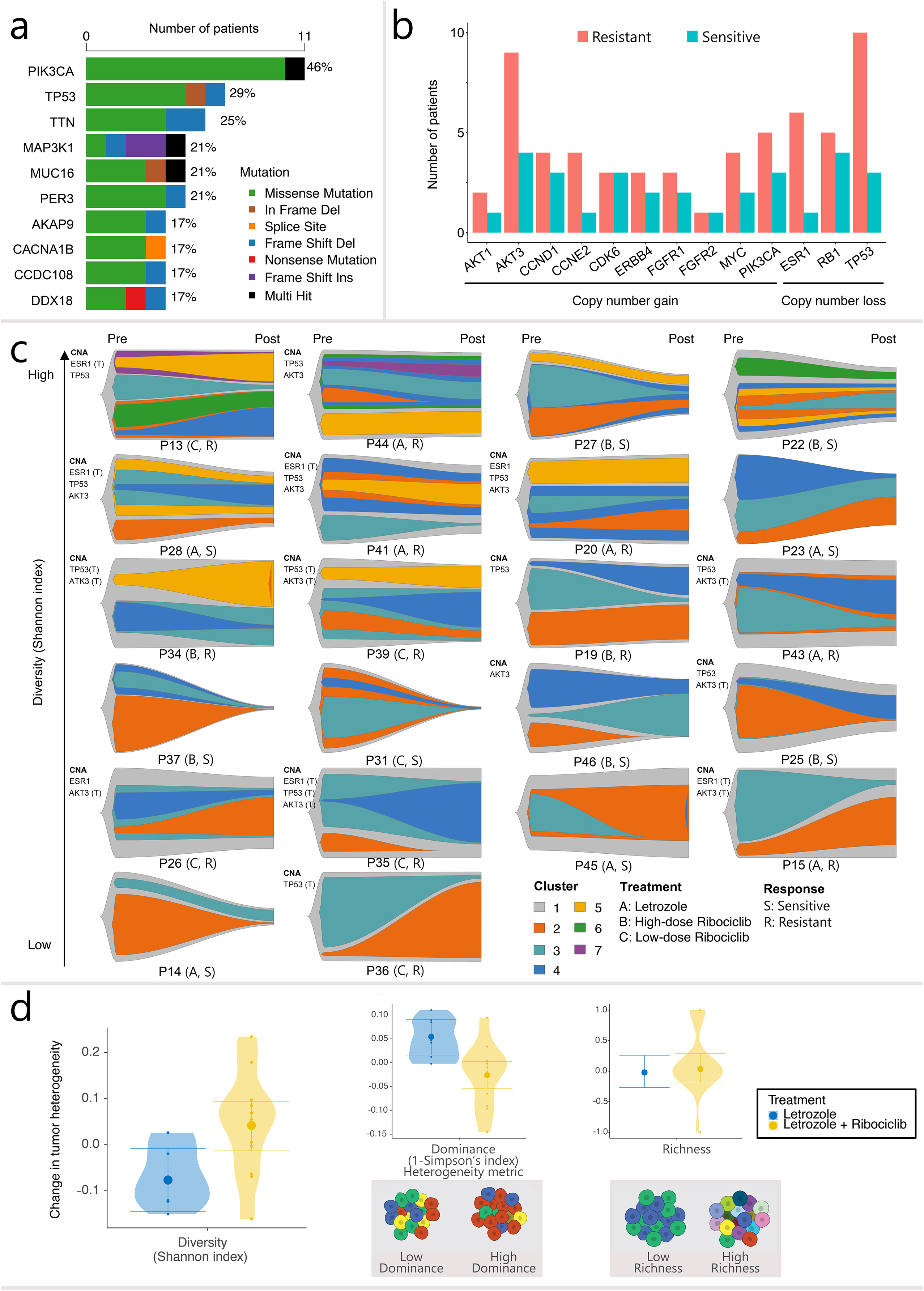
Evolution of genomic mutations in response to endocrine or combination therapy. **a**, Frequently mutated genes. Cancer driver genes were ranked based on number of tumors carrying large impact somatic mutations. Mutations were counted if they are present in one or both biopsies. The top 10 frequently mutated genes are shown. **b**, Copy number alteration of key resistant genes in sensitive and resistant tumors. Gene copy number alteration were counted for a patient if one or both biopsies carry the alteration. **c**, Clonal evolution in response to endocrine therapy or combination therapy. Patient tumors were ranked by initial diversity (Shannon index at Day 0). The height offishplot scaled to reflect changes of tumor size during treatment, normalized to 1 as maximal size as assessed by the Gaussian model. Two biopsies (pre- and post-treatment) were sequenced and used to perform analyses of clonal evolution for each tumor. Day 180 biopsies were used as post-treatment samples except P36, P44, P45, and P46. Only Day 14 biopsies were available for sequencing for these four patients. Copy number alteration (CNA) of ESRl loss, TP53 loss, and AKT3 gain are labelled for each patient when present at one or both biopsies. A “T” in parentheses indicates CNA is truncal in this tumor defined based on FACETS copy number analysis using WES and inferCNV copy number profiles using scRNAseq. **d**, Change of tumor heterogeneity in response to endocrine therapy or combination therapy, as measured by Shannon index of overall diversity, and broken down into the diversity components ofsubclonal dominance and richness. Violin curves show the between patient variability in the change of heterogeneity during the trial (small points show observed changes in diversity). Hierarchical regression models identified the average change in tumor heterogeneity (large points) during endocrine (blue) or combination therapy (yellow) (uncertainty quantified by 95% confidence interval error bars). Schematic diagrams show the distinction between differences in dominance and richness.

Subclonal cancer cell populations were identified using PyClone (32), after normalizing for cancer cell purity and copy-number changes. All patients’ tumors show polyclonal populations, with a range of 2-7 subclones present over the course of therapy (Figure 2c). Unlike later stage ER+ breast cancer (3) or triple negative breast cancer (TNBC) (33), few patients show bottleneck events, in which a single dominant subclone emerges during treatment, during the six month course of therapy, with the majority of patient tumors maintaining persistent polyclonal populations (Figure 2c) (27).

The evolution of subclonal tumor heterogeneity between treatment arms was further examined by assessing the frequency of mono- or poly-clonal populations over time (Figure 2d). For each biopsy sample, overall tumor diversity was measured using Shannon’s index (34). Diversity includes two components: richness and dominance. Richness was measured by the number of subclones in a tumor, whilst dominance reflects the abundance of each subclone and the uneven fraction of cells in each tumor subclone (Figure 2d). Changes in dominance were measured by the differences in Simpson’s dominance index over time (34). Tumor heterogeneity was found to decrease during endocrine treatment, due to an increase in the dominance of a resistant subclonal population (t=2.33, p<005). In contrast, overall tumor heterogeneity increased during combination treatment, as a result of a decreasing dominance of any one subclonal population (t=-5.06, p<0.001) (Supplementary Table 8). The increase in tumor genetic diversity under combined therapy suggests that multiple genetic mechanisms of resistance to ribociclib can lead to a resistance phenotype with similar fitness.

### Identifying resistant phenotypes during treatment

To determine how cancer cell phenotypes evolved during endocrine and combination treatment, we analyzed single cell pathway activity across all three time points (day 0, 14 and 180), using single sample Gene Set Enrichment Analysis (ssGSEA) scores (35). We identified pathways from the Molecular Signatures Database (MSigDB; c2 and hallmark pathways) that were significantly activated or deactivated during each treatment (Supplementary Table 9) (36). A hierarchical regression model determined overall and patient specific trajectories of pathway expression within an arm. The statistical significance (p-values) of pathway activation were corrected for by applying Holm’s conservative correction procedure (37). The most significantly altered ssGSEA pathways (n=87) were categorized into groups reflecting major biological processes, including estrogen receptor activity, signal transduction and proliferation (cell cycle activity) (Supplementary Table 10). The phenotypic heterogeneity in these three most prevalently dysregulated biological processes were assessed in detail through the extraction, dimension reduction, and model-based analysis of the individual genes constituting the detected pathways. Of note, at the end of 6 months of treatment, remaining tumor cells may include those with a resistance phenotype following an initial clinical response to therapy, as tumors are heterogeneous and often include resistant cancer cells (3, 33).

### Accelerated evolution of diminished estrogen signaling with combination treatment

Analysis of the pathway trajectories over time shows that persistent tumors treated with endocrine therapy alone maintained estrogen signaling following treatment (measured by Hallmark estrogen response early signature), indicating little or no sensitivity to therapy (t=0.77, p=0.45), while shrinking tumors showed a significant but modest decrease in estrogen signaling during endocrine therapy (Figure 3a, top left panel) (t=-2.138, p<0.05). This is in line with the expectation that effective response occurs when the aromatase inhibitor blocks estrogen production and reduces receptor activity. In contrast, combination therapy caused a stark decrease in estrogen signaling during treatment in persistent tumors (t=-2.721, p=0.05), indicating acquisition of a low estrogen dependent state in which estrogen receptor has diminished signaling activity (Figure 3a, top middle and right panels). Even though resistance patterns differ across treatment arms for tumors that grow on therapy, patients with tumors shrinking after combination therapy show similar diminished levels of endocrine signaling in cancer cells across all treatment arms.

**Figure 3.**
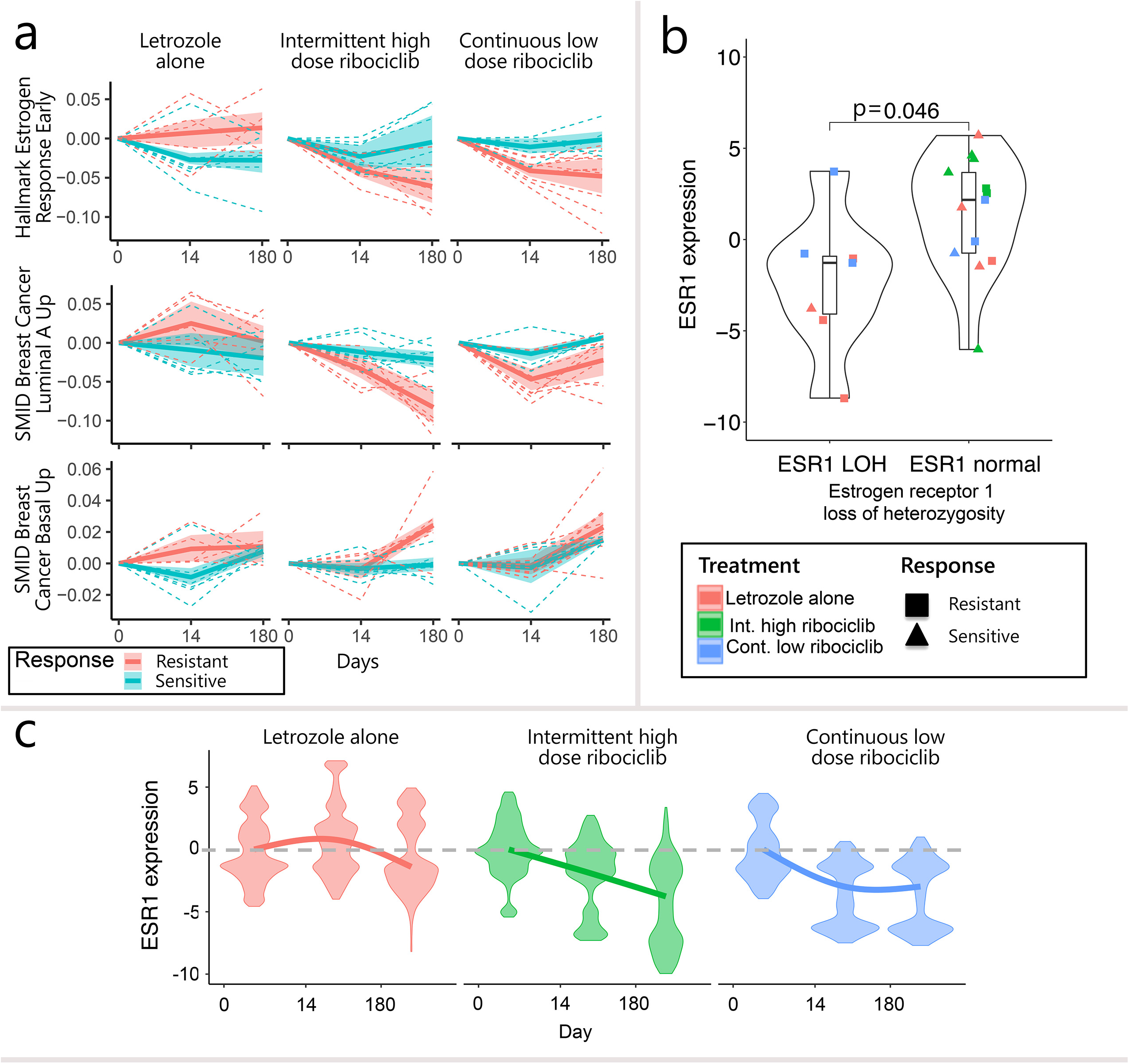
Accelerated evolution of estrogen independence during combination therapy. **a**, Changes in single cell estrogen receptor activity and signatures of estrogen dependent (luminal) and estrogen independent (basal) phenotypes during treatment (columns), in tumors resistant (persistent) or sensitive (shrinking) after therapy (red and blue, respectively). Pathway trends across tumors were determined using hierarchical regression (solid lines). Tumors’ pathway trajectories are shown by dashed lines and confidence intervals of model estimates shown by shaded regions. Pathway activity measured using the ssGSEA signatures (Hallmark estrogen response early, Smid breast cancer luminal A up and Smid breast cancer basal up respectively). **b**, Loss ofheterozygosity (LOH) of the estrogen receptor gene (ESRl) associated with reduced average expression in pre-treatment biopsies. Violin curves show the distribution of average ESRl expression in tumors with or without LOH. Points indicate patient specific averages (color indicates treatment and shape signifies tumor response). Box and whisker plots indicate the median and upper/lower quantiles of patient tumor expression. **c**, Reduction in ESRl expression accelerated under combination treatments, compared to endocrine therapy (columns: letrozole vs ribociclib treatments). Violin curves show the distribution of single cell expression during each treatment (color). Expression normalized relative to the baseline average (grey dashed line). Hierarchical generalized additive models predict the changes in expression during treatment, accounting initial patient specific difference (colored curves).

During combination therapy, diminished ER expression and pathway activity was accompanied by a transition from a luminal-like to basal-like characteristics (Figure 3a, middle and bottom panels) (transition under high dose combination therapy: t=2.85, p<0.05; low dose: t=2.97, p<0.05). The transition occurred repeatedly in different subclones from patient’s tumors (Supplementary Figure 5 and Supplementary Table 11). Cells with increased basal-like phenotype score had lower estrogen receptor (ESR1) expression (t=-5.77, p<0.001), and this negative association was strengthened under both ribociclib treatment arms (t=-3.15, p<0.005) (Supplementary Figure 6 and Supplementary Table 12). Estrogen signaling loss was also correlated with an increased signature of endocrine therapy resistance (Supplementary Table 12). The acute switch to an estrogen independent state of proliferation was only weakly observable during endocrine therapy alone, indicating that the selective pressure of combination therapy, especially at high doses, promoted accelerated evolution of the phenotypic switch away from estrogen signaling activity in resistant tumors. Of note, in sensitive tumors that shrink during treatment, combination therapy did not further decrease estrogen signaling as measured through transcriptional changes. Further, copy number increases in AKT3 or losses in ESR1 or TP53 at day 0 had only modest effect on ER expression during therapy (Supplementary Figure 7); however, larger samples sizes are needed to fully assess these relationships.

In addition to pathway analysis, we analyzed the change in the gene expression of ESR1 during treatment. We quantified the trajectories of ESR1 expression using a hierarchical generalized additive model to estimate decline of expression during each treatment, while accounting for differences in initial tumor expression. In line with other studies, we found from the WES data that loss of ESR1 heterozygosity (LOH) in tumor cells also significantly diminished expression of ESR1 mRNA (Figure 3b, W=20, p<0.05). In some patients this was found in the pre-treatment sample, indicating innate resistance to endocrine therapy (38, 39). Similar to estrogen pathway levels, combination therapy (especially intermittent high dose ribociclib treatment) accelerated the reduction of ESR1 mRNA levels over time (ESR1 reduction: t=-31.54, p<0.001) (Figure 3c), with persistent tumors in particular showing the most dramatic reductions in expression (t=-80.28, p<0.001) (Supplementary Figure 8). Shrinking tumors also show reductions in receptor mRNA levels, but more comparable to what occurred with letrozole alone (t=-79.50, p<0.001).

### Single cell RNA sequencing uncovers common transcriptional changes in the JNK MAPK pathway during treatment

In order to identify the proliferative stimuli in cancer cells with diminished estrogen signaling, we analyzed alternative growth pathways that were dysregulated during combination therapy (Supplementary Table 10). In addition to a loss of ER signaling, the pathway analysis detected several signatures of MAPK signaling network alterations. The JNK and ERK pathways are major component of the MAPK network, along with the p38 MAPK pathway; both showed divergent patterns of activity following combination therapy but not endocrine therapy alone (Figure 4a; Supplementary Figure 9; Supplementary Table 10).

**Figure 4.**
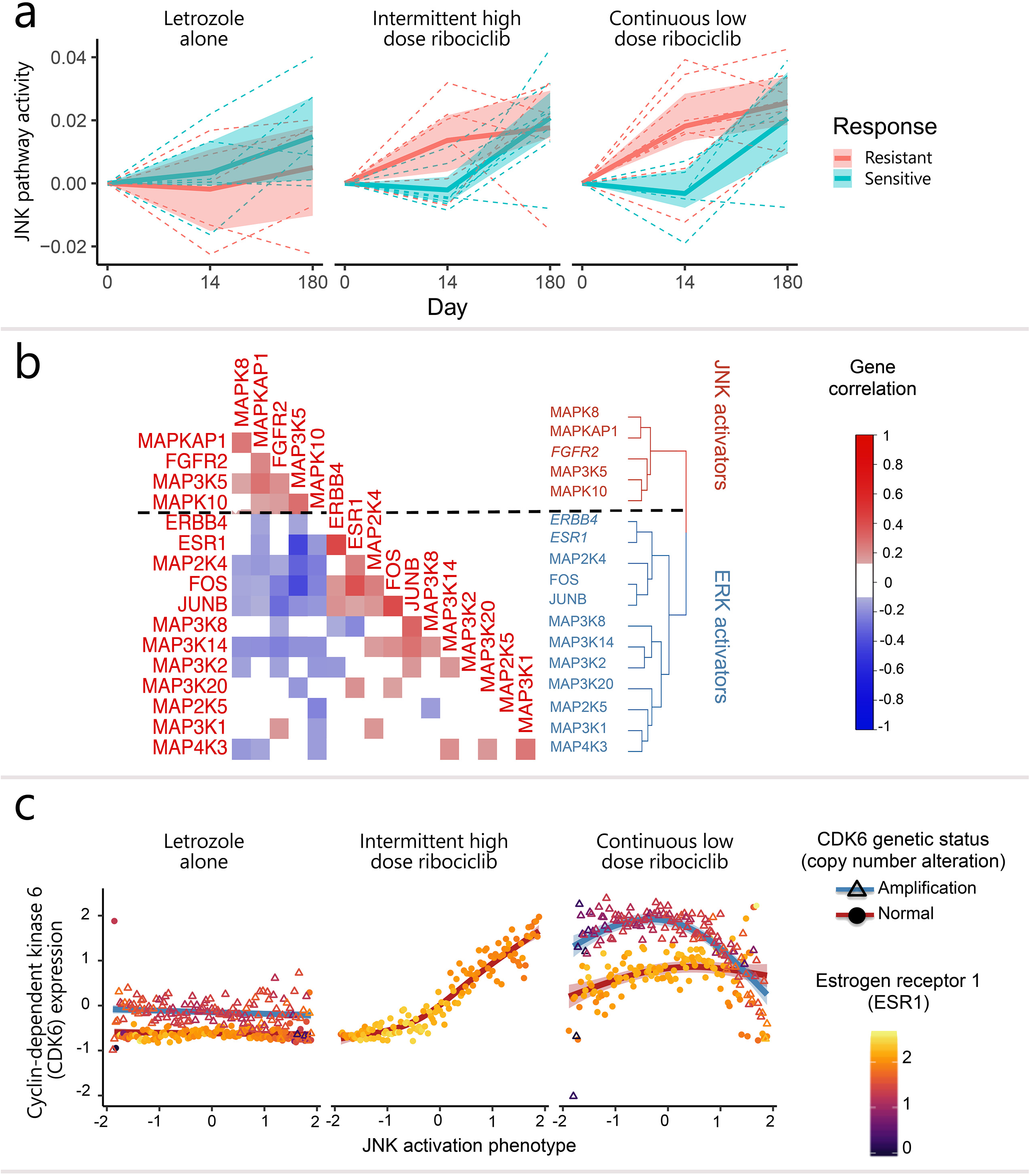
JNK pathway activation occurs during the emergence of combination therapy resistance and is associated with estrogen independence and increased CDK6 expression. **a**, JNK pathway activity increases during treatments (columns) in resistant tumors (red) compared to sensitive tumors (blue) at day 14, and tumor cells in most tumors at end of treatment (day 180). Pathway trends determined using hierarchical regression (solid lines). Tumors’ pathway trajectories are shown (dashed lines) along with confidence intervals of model estimates (shaded regions) (JNK ssGSEA pathway=St JNK MAPK). **b**, Heatmap of correlation between MAPK gene expression, showing the dichotomy between JNK and ERK activating genes across treatments (ESRl, ERBB4 and FGFR2 receptors added to indicate their relationship to MAPK genes). **c**, End of trial expression ofCDK6 (ribociclib target gene) in subclonal tumor populations with differing levels of JNK activation (phenotype integrates across MAPK gene expression) and for tumors with CDK6 genetic amplification (triangles and blue curves) or normal copy number (circles and red curves). Generalized additive models describe the relationship between JNK signaling activity and CDK6 expression at end of therapy (curves), with shaded regions indicating model confidence bands. Average estrogen receptor (ESRl) expression is shown for each subclonal population with differing JNK activation (point color gradient).

The JNK MAPK signaling pathway was upregulated during combination therapy in resistant tumor cells at day 14, but not in sensitive cells (Figure 4a) (high dose: t=3.10, p<0.01, low dose: t=2.40, p<0.05, resistant versus sensitive t=2.79, p<0.05). Tumor cells from all patients remaining after six months of combination therapy showed upregulation of JNK expression. This result indicates the acquired resistance in persistent cells, even in tumors that are initially sensitive and highlights the heterogeneous nature of these breast tumors (as shown in Figure 2c). Significant activation of the JNK pathway score was not seen in resistant tumors treated with endocrine therapy alone (Figure 4a left panel) (t=0.81, p=0.44), indicating its specific role in CDK inhibitor resistance.

Growth factor signaling can mimic estrogen action and ERK can phosphorylate ESR1, leading to estrogen independent activation (20-22). During combination therapy, resistant tumors became less dependent on ERK signaling (Supplementary Figure 9) (t=-2.83, p<0.05). In contrast, sensitive tumors maintained higher ERK signaling (no significant ERK change: t=0.89, p=0.41) perhaps due to the significant crosstalk between ERK and ER, including ER activation of growth factor signaling (40-43). Taken together with the activation of JNK signaling under combination therapy, the lack of ERK signal utilization further reflects the transition to an endocrine independent resistance state, with reduced reliance on the positive estrogen/ERK crosstalk (21, 44).

An inverse transcriptional expression pattern was found between JNK and ERK pathway genes in the single cell expression data (MAPK gene set in Supplementary Table 13). Within the JNK and ERK pathways, gene expression was positively correlated, while being inversely related to the other gene set (Figure 4b). This dichotomy was consistent in cells across treatments and timepoints (Supplementary Figure 10). An overall JNK activation phenotype score was then constructed which integrates the gene expression across MAPK genes, providing a pseudotime metric and placing cells on a continuum from JNK to ERK dominated signal transduction. Utilizing the collinearity of MAPK gene expression, we performed Uniform Manifold Approximation and Projection (UMAP) dimension reduction of MAPK genes and used the major axis of phenotypic variation to provide the JNK activation phenotype (Supplementary Figure 11). High JNK activation phenotypes were verified to correlate strongly with single cell upregulation of ASK1 and JNK1/3 genes, while low scores reflected ERK activation, including MEK4, MEKK1-3, and ERBB4 upregulation (Supplementary Figure 11).

To understand how JNK signaling relates to resistance, we examined the link between JNK activation and the expression of the ribociclib target gene, cyclin dependent kinase 6 (CDK6; CDK4 is not expressed in the scRNAseq data). Across each subclone in all tumors, the average CDK6 expression was calculated along with JNK activation levels (Figure 4c, red line). In addition to expression levels in copy number neutral cells, we also separately analyzed JNK levels in those subclones with amplified CDK6 (Figure 4c, blue line; no patients in arm B had CDK6 CNA). Using generalized additive models, we analyzed the nonlinear CDK6:JNK relationship in each treatment and CDK6 amplification group (Figure 4c).

Tumors converge on signaling states providing resistance to endocrine and combination therapy. Following combination therapy, JNK activation was concurrent with the upregulation of CDK6 in patients lacking pre-existing CDK6 copy number alterations (Figure 4c) (F=130.10, p<0.001). Subclonal populations with low JNK activation had low CDK6 expression (t=12.39, p<0.001), indicating that these cells were not in a proliferative state. In contrast, tumors with CDK6 copy number alterations had high CDK6 expression independent of their JNK activation status (t=16.03; p< 0.001), indicating that genetic alteration removed the requirement for altered signal transduction. Overall, tumors with higher JNK activation decreased in size less during combination therapy (t=13.84, p<0.001) (Supplementary Figure 12).

Estrogen signaling was explored between cancer populations differing in JNK activation and CDK6 amplification status. Subclonal populations with low estrogen receptor expression exhibited CDK6 upregulation (Figure 4c) (t=-3.89, p<0.001). In patients lacking CDK6 amplification (Figure 4c, filled circles), estrogen loss was linked to JNK activation (t=-6.13, p<0.001). Meanwhile, patients with CDK6 amplified tumors lost estrogen receptor expression independent of JNK activity (Figure 4c, open triangles) (t=-0.20, p=0.84). In the absence of CDK6 amplification, JNK activation provides an alternative pathway to estrogen independent proliferation under combined therapy. Increased expression of the anti-apoptotic MCL1 gene was also observed in persistent tumor cells after treatment (t=2.68, p<0.05) (Supplementary Figure 13), in line with studies showing JNK stabilization of MCL1 through phosphorylation (45, 46).

### ERBB4 and FGFR2 receptor tyrosine kinase upregulation is common in cancer cells after treatment but varies across treatment arms

Given that both JNK activation and CDK6 amplification allowed estrogen independence and/or potentiation, we next examined growth factor receptors that showed compensatory increases in expression. Receptor genes comprising ssGSEA signatures detected in the pathway analysis were identified along with genes coding for cell surface proteins, including receptors, listed in the cell surface protein atlas (n=1406) (47). For each patient, we compared pre- and post-treatment expression using analysis of variance and identified receptors with a) increased expression during treatment or b) that had initially higher expression in resistant tumors. Of these genes, we identified receptors with expression inversely correlated to single cell estrogen receptor activity (Supplementary Figure 14; Supplementary Table 14).

Following this method, we found that Erb-B4 Receptor Tyrosine Kinase 4 (ERBB4) gene was significantly upregulated relative to baseline in 50% of tumors (n=13), including all but one patient with CDK6 copy number amplification (n=4/5). ERBB4 is upstream of MAPK signaling and an ER coregulator that can drive proliferation via ERK signaling (48). Tumors lacking CDK6 amplification showed less ERBB4 upregulation than tumors with CDK6 amplification (Figure 5a, top row) (t=-13.06, p<0.001), indicating that it is a mechanism of resistance to endocrine but not cell cycle inhibition. Consistent with this hypothesis, higher expression of ERBB4 was also seen in tumors after endocrine monotherapy (t=11.88, p<0.001).

**Figure 5.**
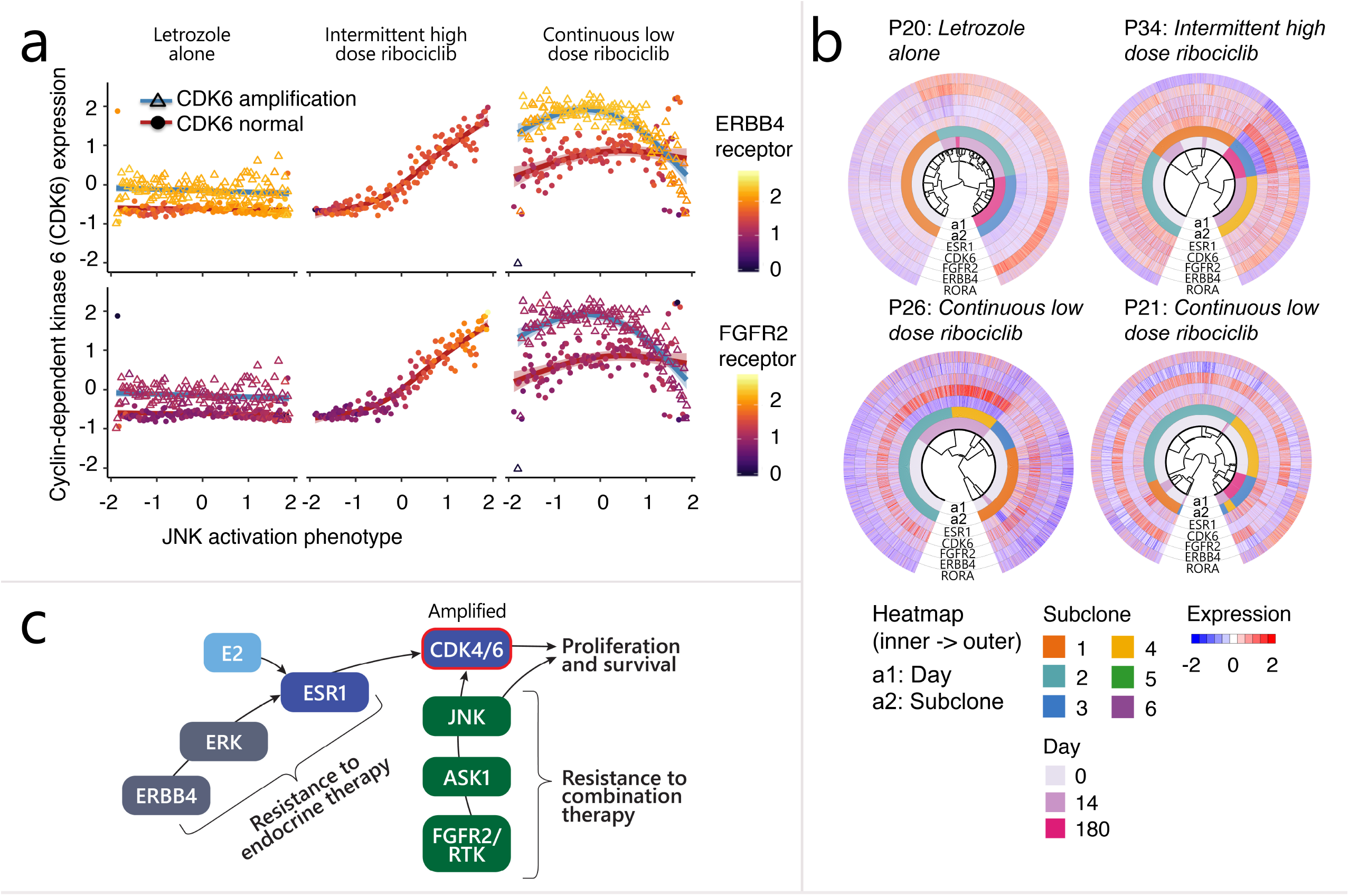
Activation of ERBB4 and FGFR2 as resistance mechanisms to endocrine and combination therapy. **a**, Average end of trial growth factor receptor expression (point color gradient) of ERBB4 (top) and FGFR2 (bottom) is shown for subclonal populations with differing JNK activation (x-axis) and for tumors with genetic amplification (triangles and blue curves) or normal copy number (circles and red curves). Cyclin-dependent kinase 6 (CDK6; ribociclib target gene) expression is shown for each population and generalized additive models (curves) describe the relationship between JNK signaling activity and CDK6 expression at end of therapy, (shaded regions indicate model confidence bands). **b**, Hierarchical clustering of tumors showing dysregulated ESRl as well as upregulated ERBB4 and/or FGFR2. For each patient’s tumor biopsies, a hierarchical clustering tree is shown at the center of circos plot. Cell annotation (timepoint and subclone) as well as expression of key resistance genes (ESRl, CDK6, FGFR2, ERBB4, RORA) were shown as heatmap. **c**, Schematic diagram showing resistance mechinisms driven by upregulation ofERBB4 and CDK6 amplification (red circle signifies amplification) or alternative signaling via FGFR2/RTK’s and JNK signal transduction.

Fibroblast Growth Factor Receptor 2 (FGFR2) was also found to be upregulated in JNK activated cells following high dose intermittent combination therapy when patients lacked CDK6 amplification (Figure 5a, bottom row). FGF receptors can also activate MAPK signaling (49, 50). This relationship was most evident in JNK activated cells following intermittent high dose combination therapy (t=18.38, p<0.001), with 64% (7/11) of patients showing high expression of FGFR2 (Supplementary Table 14). An additional receptor upregulated over time in resistant cells was RAR Related Orphan Receptor A (RORA), a nuclear receptor that potentially modulates expression of both aromatase enzyme (51) and the ribociclib target, CDK6, which controls cell cycle progression.

Single cell relationships were constructed, through a cluster tree, based on cellular copy number alterations and show that as tumor subclones evolve during treatment, the acquisition of alternative receptors is accompanied by a loss of ESR1 (Figure 5b; Supplementary Figure 15) (Supplementary Dataset 2 shows single-cell copy number alterations of each patients’ tumor). Clonal populations show consistent concordant loss of ESR1 as the growth factor/cell surface receptors are upregulated. As an example, subclones with reduced ESR1 and upregulated ERBB4 emerged 180 days after endocrine therapy only in tumors from P20 or combination therapy in tumors from P34 (Figure 5b). In addition, subclones with dysregulated ESR1 and upregulated FGFR2 emerged 180 days after endocrine therapy in P15 or disappeared 180 days after combination therapy in tumors from P21 (Figure 5b). As summarized in Figure 5c, in tumor cells with high estrogen signaling, potentiation of CDK4/6 activation can occur through ERBB4 and ERK upregulation and activation. Alternatively, cancer cells with diminished endocrine signaling can bypass CDK4/6 inhibition through upregulation of the JNK pathway. CDK6 amplification itself can potentiate its activity and correlates with cell proliferation. In sum, resistant cancer cell state reflects a phenotypic shift from ERK to JNK MAPK signaling and diminished estrogen signaling in tumors without CDK6 amplification. Tumors with CDK6 copy number amplification showed an alternative phenotype, with growth factor receptor upregulation.

### Functional consequences of therapy on the cell cycle: rewiring to bypass CDK inhibition

Cancer cell proliferation during endocrine and combination therapies was examined by measuring changes in cell cycle pathway activity. Cell cycle activity was initially inhibited by both endocrine therapy alone and combination therapy (Figure 6a) (Biocarta cell cycle pathway decline: t=-2.728, p<0.05). However, by the end of combination treatment, cell cycle activity had rebounded (t=2.678, p<0.05). Tumor cells that persisted following intermittent high dose combination therapy showed the largest initial reduction in cell cycle activity (t=-3.290, p<0.05), followed by the greatest proliferative rebound, suggesting stronger selection for the evolution of a cell cycle rewired to bypass the G1 checkpoint and allowing proliferation independent of estrogen deprivation. In contrast, tumors persistent to letrozole treatment alone exhibited the weaker initial and subsequent reductions in cell cycle activity compared to shrinking tumors, reflective of innate resistance.

**Figure 6.**
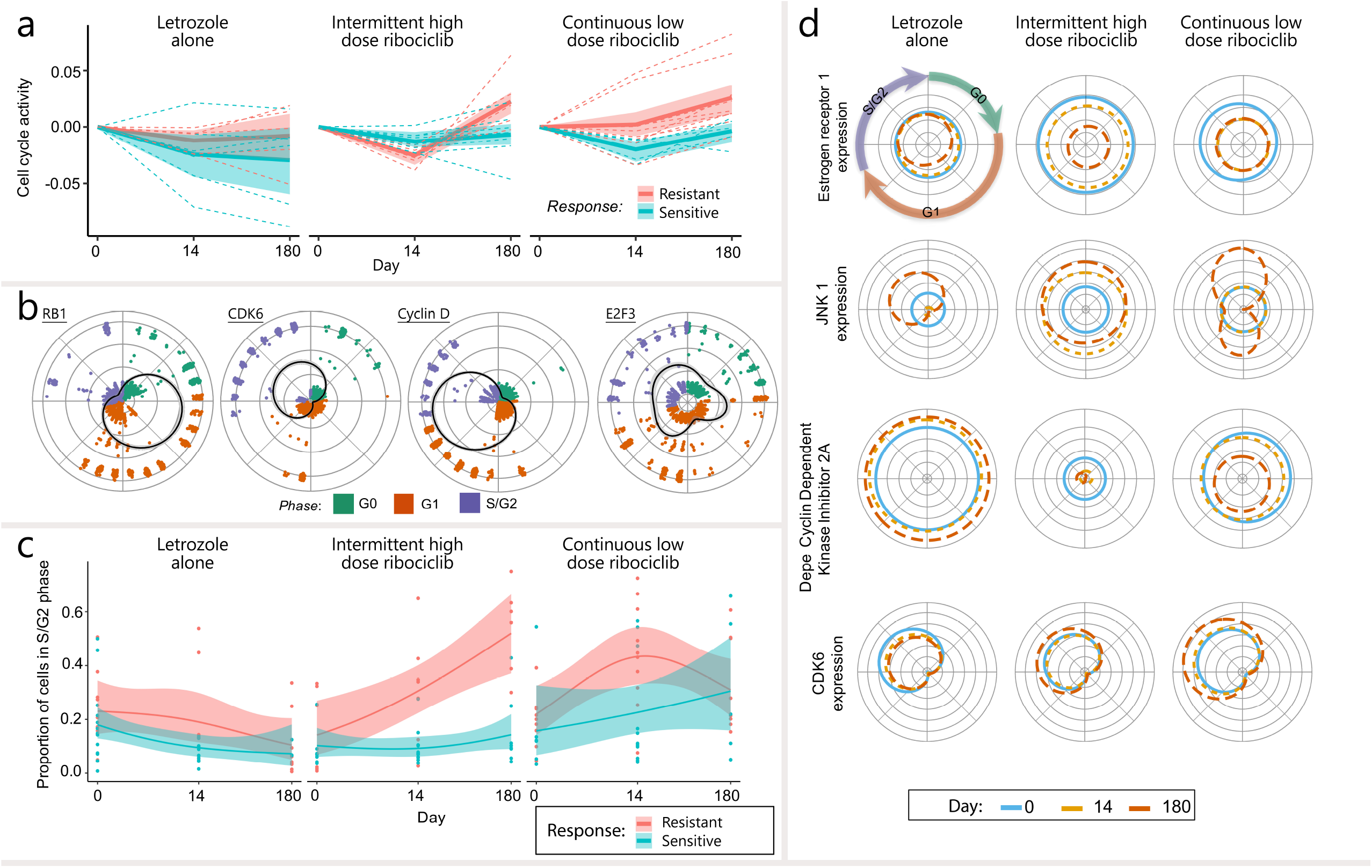
Cell cycle reactivates during combination therapy follows the loss of estrogen receptor expression, activation of JNKl, repression of the cell cycle inhibitor cyclin-dependent kinase 2A and upregulation of CDK6 during the G1 checkpoint phase. **a**, Cell cycle activity of resistant and sensitive tumors (red and blue, respectively) during treatment (columns= regimes). Trend in cell cycle activity, measured by the ssGSEA biocarta cell cycle pathway, are determined by hierarchical regression (solid lines). Tumor specific trajectories are shown (dashed lines) along with confidence intervals of model estimates (shaded regions). **b**, Visualization of the pseudotime cell cycle reconstruction obtained using the Markov model-based reCAT algorithm. The dynamics of key cell cycle gene expression across stages of the cell cycle (black lines) were recovered from cell specific gene expression (points), using cyclical generalized additive models. The recovered fluctuations of cell cycle gene expression were used to identify three distinct cell cycle phases (colors; GO, G 1 and S/G2), using a Gaussian mixture model. Cell cycle stages (clusters of cells) are colored by cell cycle phase and the distance of the point from the origin signifies the cells expression in that stage. Cell cycle orientation is consistent and comparable across circular plots. **c**, The proportion of S/G2 phase cells (passed the GI checkpoint) in samples from sensitive and resistant tumors (color) during each treatment (column). Logistic generalized additive models describe trends in S/G2 phase cell frequency over time (solid lines) and heterogeneity across tumors (shaded regions signify confidence bands). **d**, Changes in ESR I, JNK I, CDKN2A expression around the cell cycle and during treatment (columns). Colored lines show the expected gene expression of cells throughout the cell cycle prior (blue), during (orange) and after treatment (red). The distance from the center of the circle shows gene expression at point in the cell cycle.

A rewiring of the cell cycle regulatory pathways is required for resistant cells to bypass the G1 checkpoint (52-54). Cell cycle dysregulation is reflected in the changing distribution of cells across cell cycle phases and altered expression of cell cycle regulating genes. To reveal these alterations from single cell data, we extended the Markov model-based reCAT algorithm (55) to reconstruct the cancer cell cycle transitions (Supplementary Figure 16). Distinct cell cycle states, common across patients (Supplementary Figure 17) were identified by applying UMAP dimension reduction to the single cell expression of cell cycle related genes followed by clustering with a Gaussian mixture model. The states were connected by finding the shortest path that visited all states. Extending this algorithm, the dynamics of gene expression around distinct cell cycle phases was delineated using cyclical generalized additive models (Figure 6b). Through the reconstruction of the cell cycle from scRNAseq data (Figure 6b), we recovered expected events in the cell cycle, including a G1 checkpoint transition where cyclin D initially rises and is followed by CDK6 expression. Further, we observed that Retinoblastoma (RB1), the key G1 checkpoint protein, was downregulated in concert with increased expression of the E2F3 proliferation gene. This effect leads to a positive feedback in cyclin dependent kinase production and the commitment of cells to progress to S-phase.

The ability of cancer cells to bypass the G1 arrest checkpoint was assessed in detail by calculating the fraction of cancer cells in each patient biopsy that were in the division (S/G2) phase. During combination therapies, an increasing fraction of each patient’s cancer cells were found to be in the division and growth (S/G2) phase of the cell cycle (t=2.94, p<0.001). The frequency of proliferating cells increased in persistent tumors, especially those receiving high dose combination therapy (t=2.1, p<0.05). In contrast, those receiving endocrine therapy exhibited a decrease in the frequency of proliferating cells (t=-2.08, p<0.05), with no detectable difference between tumors that shrank or persisted while on therapy. This result indicates effective bypass of the ribociclib enhanced G1/S checkpoint in surviving subclonal populations.

Next, fluctuations in the expression of cell cycle, ER and JNK genes around the cell cycle were recovered by applying cyclical generalized additive models to the single cell gene expression of cells across cell cycle states. This approach predicted the smooth change in gene expression as cells transition around the cell cycle, including for genes not used to reconstruct the cell cycle. By applying this approach to cells sampled at different timepoints and from patients given different therapies, we distinguished whether treatment altered expression at specific cell cycle stages or if genes expression was dysregulated independent of the cell cycle.

ESR1 was expressed at consistent levels throughout the cell cycle (Figure 6d, top row). However, when looking across combination therapy arms, decreasing ESR1 expression was observed over time, accompanied by increased levels of JNK1 (Figure 6d, second row) (t=2.57, p<0.05) and its downstream target transcription factor JUNB (t= 10.10, p<0.001) (Supplementary Table 15). Further, during combination treatment, we observed a decrease in cyclin dependent kinase inhibitor 2A (CDKN2A (coding for p14 and p16)) (t=-3.07, p<0.005) and an increase in CDK6 expression in the G1 to S/G2 phases (t=4.81, p<0.001) (Figure 6d, bottom two rows). Taken together, these observations support the role of estrogen independent JNK signaling in decreasing cell cycle inhibition prior to the G1 checkpoint, thereby permitting cell cycle reactivation.

## Discussion

With their proven success in treatment of metastatic ER positive breast cancer (56-58), CDK 4/6 inhibitors are currently being tested in the treatment of early stage breast cancer (7-10). Neoadjuvant trials facilitate collection of tissue biopsies at various timepoints before, during and after therapy providing an opportunity to define the pathways that drive resistance in early stage breast cancer. To address this need, we studied the evolution to resistant states in patient tumor cells during treatment with endocrine therapy alone or in combination with the CDK4/6 inhibitor, ribociclib. Given the patient complexity, when monitoring tumors during neoadjuvant treatment, we used hierarchical statistical models to identify response related phenotypes. We then analyzed single cell heterogeneity in growth signal transduction and cell cycle pathways by integrating dimension reduction and pseudotime reconstruction approaches with generalized additive modelling techniques.

In patients treated with endocrine therapy alone, we see compensatory signaling between ESR1 and ERBB4 receptor tyrosine kinase, with activation of RTK and downstream ERK MAPK upregulation offsetting decreased ESR1 levels in resistant tumors. Compensatory expression of ESR1 signaling by EGFR and ERBB2 receptors has been detailed in cell lines, and EGFR, ERBB2 and IGF1 receptor (IGF1R) activity has been shown to cause tamoxifen resistance (2, 59). However, the role of these pathways has not previously been detailed in patient tumors during treatment. We uncovered convergence towards distinct pathways of resistance in patients treated with combined letrozole and ribociclib, including the evolution of estrogen independent proliferation through the upregulation of alternative growth factor receptors (e.g. ERBB4, FGFR2 and RORA) and JNK MAPK signaling including ASK1/MAP3K5 and JNK1-3 (MAPK8-10) (60, 61). 21% of tumors had genetic amplification of the CDK6 gene through copy number gain, directly enhancing transcription, while those lacking CDK6 amplification initially upregulated JNK signaling during treatment, providing an alternative route to upregulation of CDK6 levels. Lastly, upregulation of additional genes such as MAPKAP1 in JNK activated cells may also contribute to the promotion of survival and proliferation through modulation of MAPK signaling (62, 63). The relationship between resistance signaling pathways remains to be determined; however, this study shows that multiple bypass mechanisms exist that enable cells to survive therapy. Due to gene dropouts in scRNAseq data, transcription of all genes was not measured, and additional pathways and genes may also be dysregulated in this setting.

The JNK MAPK pathway is classically considered to be central to apoptotic signaling (64, 65). However, JNK has been found to drive aberrant tumor growth in a drosophila model system through modulating survival of nearby cells (66, 67), highlighting a potential role for JNK in tumorigenesis and cell-cell interactions within a tissue. Further, JNK has been shown to activate cell cycle regulated proteins such as CDK4 (68, 69). Regulation of these pathways can drive proliferation of ER+ tumors without a requirement for estrogen, which is essential as cells become ESR1-independent. Interestingly, a recent study showed p38 MAPK can independently drive cell cycle activation and proliferation (70, 71). The role of MAPK in proliferation independent of canonical cell cycle signaling and their role in drug resistance is an area of further investigation. Given the dual role of JNK as a proliferative and an apoptotic signal, it provides a promising target for evolution-based therapies. The CDK inhibitor abemaciclib, which recently demonstrated clinical benefit in early stage patients, is less specific for CDK4/6 than ribociclib, has a different CDK4/CDK6 inhibition ratio, and includes targets such as JNK (72). Despite similar clinical efficacy between these two agents in metastatic breast cancer, differences may be seen in the early setting based on target specificities of these different CDK inhibitors. Thus, our results suggest that JNK inhibition may explain the success of abemaciclib in the MonarchE adjuvant clinical trial (12), while the failure of palbociclib in the PALLAS adjuvant clinical trial (13) may be due to lack of JNK inhibition. They also lead to the hypothesis that adding JNK inhibitors to either palbocilib or ribociclib may improve their efficacy in early stage breast cancers.

ESR1 signaling loss occurred during treatments targeting this signaling pathway and was accelerated in patients receiving higher combination therapy doses. Further studies tracking the tumor mass during intermittent combination therapies will clarify the role of drug dose and timing on the rate of evolution of resistance phenotypes such as acquisition of endocrine independence. As breast cancer cells become independent of endocrine signaling, they require alternative pathways to drive proliferation. In contrast to earlier stage tumors, our prior studies with metastatic tumors show acquisition of cancer stem cell and EMT signaling during progression with chemotherapy and other targeted therapies (3). Therefore, it is possible that alternative pathways drive proliferation and response to CDK4/6 inhibitors in early versus late stage breast cancer. Studies such as this one highlight that efficacy in late stage cancer may not imply the same in early stage cancer, which is what was seen in the PALLAS trial of palbociclib (11, 73-75).

Changes in markers of proliferation after 2 weeks of endocrine therapy have been used as early indicators of efficacy (11, 76). Our results demonstrate that information beyond proliferation can be learned from early biopsies in neoadjuvant therapy trials. In addition to providing biological insight into mechanisms of resistance, scRNAseq data could provide clinically useful data to guide personalized next line therapy choices. As a prognostic marker, changes in JNK activation or cell cycle state could be tested at 2 weeks as a biomarker of the clinical efficacy of combination aromatase inhibitor-CDK4/6 inhibitor therapy in early breast cancer, to determine whether to continue therapy. Similarly, transcriptional changes reflecting shifts from luminal to basal-like characteristics or diminished expression of Rb concordant with increases in E2F proliferation genes may contribute to early biomarkers of resistance. Tumors that show an insufficient decrease in estrogen signaling and proliferation after two weeks of estrogen therapy alone may benefit from switching to another endocrine therapy or receiving growth factor receptor signaling inhibitors to achieve full inhibition of proliferation. While ERK signaling is correlated with estrogen receptor activity in tumors treated with endocrine therapy alone, based on our genomic analyses, a prior clinical trial with fulvestrant and selumetinib, a MEK1/2 inhibitor that blocks signaling directly upstream of ERK in breast cancer progressing after aromatase inhibitor therapy, failed to show efficacy (77), although that was in metastatic and not early breast cancers.

ER+ tumors treated with combination estrogen and CDK4/6 inhibition therapy that develop a loss of estrogen signaling and acquisition of ERBB4/FGFR2 and JNK signaling could be treated with targeted therapies that block pathways. Several options are approved for use or in clinical trials (78-80). Given that JNK activity has a dual role in cell proliferation and apoptosis, additional therapies targeting anti-apoptotic signaling should also be studied. The PALLET clinical trial showed that clinical response rates over 14 weeks under combination therapy and indicated that reduced tumor proliferation was balanced by diminished apoptosis, as measured by lower c-PARP expression (11). Anti-apoptotic treatments could be given either at time of progression, based on imaging, or following neoadjuvant therapy in patients with viable resistant cells at time of surgery. Given the potential for additional therapy to increase selective pressure, it will be imperative to rationally determine optimal dosing and timing of drug treatment regimens to reverse or prevent a resistant state while minimizing side effects.

In conclusion, our analysis identifies mechanisms of how tumors circumvent endocrine and CDK4/6 inhibition through phenotypic and subclonal evolution. These mechanisms include shifts to alternative proliferative signaling pathways, bypassing dependence on ER and CDK6 activation. This approach provides a method to detect resistance mechanisms early in cancer treatment to identify phenotypic targets directed at surviving tumor subclones in tumors. Absence

## Methods

### Patient cohort and sample collection

Patient tumor core biopsy samples were collected prospectively under Clinical Trial # NCT02712723, as a multicenter study led by Dr. Qamar Khan at the University of Kansas Medical Center (IND # 127673) entitled: Femara (letrozole) plus ribociclib (LEE011) or placebo as neo-adjuvant endocrine therapy for women with ER-positive, HER2-negative early breast cancer (FELINE Trial). FELINE is a randomized, placebo controlled, multicenter investigator-initiated trial. Patients in this trial were enrolled from 10 centers in the United States. Postmenopausal women with pathologically confirmed, non-metastatic, operable, invasive breast cancer and clinical tumor size of at least 2 cm were included. Invasive breast cancer had to be ER positive (≥66% of the cells positive or ER Allred score 6-8) and HER2 negative by ASCO-CAP guidelines. Patients were randomized 1:1:1 to one of three treatment arms. Arm A received letrozole plus placebo, Arm B letrozole plus ribociclib 600 mg daily for 21 out of 28 days of each cycle and Arm C received letrozole plus ribociclib 400 mg continuously. Protocol therapy was continued until the day before surgery. Mammogram, MRI and ultrasound of the affected breast were performed at baseline and a mammogram and ultrasound was performed at completion of neoadjuvant therapy. MRI of the breast was performed after completion of 2 cycles of treatment (Day 1 of cycle 3). Samples collection for tissue was mandatory. Three tumor core biopsies were collected temporally over the course of treatment: screening (Day 0), Cycle 1 Day 14 (Day 14), and end of trial (Day 180) using a 14-gauge needle. Immediately after collection research biopsy samples are snap frozen embedded in Optimal cutting temperature (OCT). Informed consent was obtained from all patients according to protocols approved by the institutional IRBs and in accordance with the Declaration of Helsinki. This study was approved by University of Kansas Institutional Review Board (protocol # CLEE011XUS10T).

### Nuclei isolation

On a dry-ice platform to keep tissue frozen, OCT embedded tumor core biopsies were placed in a 60mm petri dish and excess tumor margins were removed from core biopsies using a razor blade. Core biopsies were then placed in 1x PBS, pH 7.4 (Gibco, Cat# 10010) + Cytoprotective reagents (all components from Millipore Sigma, 0.54µM Necrostatin (Cat# N9037), 1.0µM HPN-07 (Cat# SML2163), 0.31µM sodium 3-hydroxybutyric Acid (Cat# 54965-10G-F), 78nM Q-VD-OPH (Cat# SML0063)) + 0.04% Bovine Serum Albumin (BSA, EMD Millipore, Cat # 12661525mL). Tissue was homogenized by mincing with a sterile razor blade in 2 mL of sterile 4:1 Lysis Buffer (10mM Tris-HCl, pH 7.8 (Teknova, Cat# T1078), 146mM NaCl (Alfa Aesar, Cat# J60434AK), 1mM CaCl2 (G-Biosciences, Cat# R040), 21mM MgCl2 (G-Biosciences, Cat# R004), 0.05% BSA (EMD Millipore, Cat# 12661525mL), 0.2% Igepal CA-630 (MP Biomedicals, Cat# 198596), DNase/RNase free water (Gibco, Cat# 10977)) : DAPI Buffer (106mM MgCl2, 50 µg/mL 4’, 6-diamidino-2-phenylindole (DAPI, Invitrogen, Cat# D1306), 5mM Ethylenediaminetetraacetic acid (EDTA, Quality Biological Inc., Cat# E522100ML), DNase/RNase free water) + 0.2 U/µL SUPERase-In RNase Inhibitor (Invitrogen, Cat# AM2694). Homogenized tissue was incubated for 15 min at 4°C to release nuclei. Lysate was then filtered through a 40 µm mesh filter (Falcon, Cat# 352340) collecting isolated nuclei in flow through. All downstream nuclei processing utilized Eppendorf DNA LoBind tubes to minimize nuclei loss. Nuclei were centrifuged at 4°C, 500 × g for 5 min and washed two times with 500-1,000 µL of Nuclei Resuspension Buffer (8mM Tris-HCl, pH7.8, 117mM NaCl, 0.8mM CaCl2, 38mM MgCl2, 0.04% BSA, 0.2U/µL SUPERase-In RNase Inhibitor, DNase/RNase free water) or 1x PBS + 1% BSA + 0.2U/µL SUPERase-In RNase Inhibitor. Nuclei were resuspended in 1x PBS + 1% BSA + 0.2U/µL SUPERase-In RNase Inhibitor at a target of 1,000 cell/µL, filtered using a 40 µm mesh filter, then counted on a hemocytometer by DAPI florescence on a Laxco LMI-6000 Inverted microscope with Florescence contrast or Invitrogen Countess equipped with DAPI filter and maintained on ice prior to scRNAseq.

### Exome data sequencing, variant calling, and copy number alteration

Whole-exome sequencing was performed for 24 patients with cancer cells that are present at both pre-(Day 0) and post-treatment (Day 14 or Day 180). Genomic DNA of tumor and matched germline samples were captured using the SureSelect Human All Exon v7 (Agilent) or xGen Exome Research Panel v2 (IDT). Enriched DNA samples were sequenced on an Illumina NovaSeq 6000 with 150 bp paired-end reads. Sequence analyses were performed with a Bioinformatics ExperT SYstem (BETSY) (81). Briefly, sequences were processed by trimmomatic v0.39-1 (82) (MINLEN:15 LEADING:3 TRAILING:3 SLIDINGWINDOW:4:15) to trim adaptors and low-quality sequences. The trimmed sequences were aligned to the hg19 human genome with BWA-MEM v0.7.17 (83-85) and bam files were sorted by sambamba v0.6.8 (https://lomereiter.github.io/sambamba/). PCR duplicates were marked by the Picard tool v2.18.4 (86). Local realignment around the 1000 Genomes Phase I indels were performed with the Genome Analysis Toolkit (GATK v3.8) (87).

Somatic single-nucleotide variations (SNVs) and small indels (≤ 50 bp) were identified with MuTect2 (implemented in GATK v3.8) (87), Strelka v2.9.2 (88), and Varscan2 v2.4.3-2 (89) using tumor-normal pairs. Variants with read depth ≥ 25, alternative allele read depth ≥ 5, variant allele frequency (VAF) ≥ 0.05 in tumor samples, and VAF ≤ 1% in normal samples were characterized as somatic variants for further analysis. Somatic variants were annotated with ANNOVAR v2018Apr16 (90). Somatic copy number, tumor purity, and ploidy were estimated from WES using FACETS v0.5.14 (91) and Sequenza v2.1.2 (92) (Supplementary Dataset 1). Multiple runs with different parameters were performed for each tool. A best model was chosen for each patient by manual inspection of log-ratio and copy number profile of each patient. Copy number alteration of cancer driver genes were determined based on the median log-ratio of the segment (cnlr.median) and copy number estimated by FACETS. A copy number gain was defined as genes located on segment with cnlr.median ≥ 0.2 and copy number > ploidy. A copy number loss was defined as genes located on segment with cnlr.median ≤ 0.2 and copy number < ploidy. A loss of heterozygosity (LOH) was defined as genes located on segment with minor allele completely lost (minor_cn = 0).

### Clonal structure and evolution

Somatic mutations obtained from the above analysis were clustered by PyClone v0.13.1 using the Beta Binomial model with 10,000 iterations. Tumor purity and copy number inferred by FACETS v0.5.14 or Sequenza v2.1.2 were used by PyClone to estimate cellular prevalence of somatic mutations and mutation clusters. The clonal evolution models were inferred by ClonEvol v0.99.11 (93) based on mutation clusters and cellular prevalence of somatic mutations predicted by PyClone. The truncal cluster was assigned to the cluster with cellular prevalence 80% in at least one sample. Mutations clusters with five or fewer mutations were discarded unless the cluster represented a truncal cluster. Mutation clusters showing similar changes across samples were merged when ClonEvol failed to predict clonal evolution models. Phylogenic trees of clonal structure were generated by ClonEvol. Fishplots were generated by fishplot v0.5 (94).

### Evolution of subclonal diversity

The evolution of tumor heterogeneity during treatment was assessed by quantifying the changes in subclonal diversity across patients in each arm. The relative frequency of cancer subclones (*p*(*i,T*))was calculated for biopsies taken from each patient (i) at the first and last treatment timepoints (T). The overall subclonal heterogeneity of each tumor sample was measured using the Shannon diversity index. To disentangle the two key components of diversity (richness and dominance), we calculated richness by the number of subclones identified per sample and measured dominance as 1 - Simpson’s index (Σ (*p*_*s*_(*i,T*)^2^)).

Changes in tumor diversity (D) over time (T) and between treatment arms (A), were assessed using a random effects model:

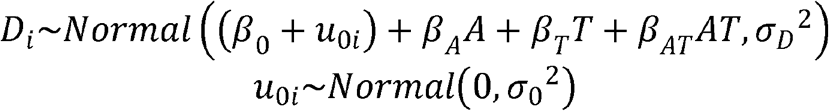

Pre-treatment differences in tumor diversity between patients in different arms were accounted for by the treatment-specific estimates of initial diversity (*β*_0_vs *β*_*A*_). Subsequent changes in diversity were described by treatment-specific temporal trend terms (*β*_*T*_and *β*_*AT*_). Individual variability in pre-treatment tumor heterogeneity was accounted for by allowing the model intercepts to vary among patients (*u*_0*i*_). Likelihood ratio tests were used to assess whether significant changes in tumor heterogeneity occurred during treatment. The likelihood of the above model was compared with that of nested null hypothesis models in which no change in tumor heterogeneity occurred (fixing *β*_*T*_ *& β*_*AT*_ *= 0*) or equal changes in tumor heterogeneity occurred across treatment groups (fixing *β*_*AT*_ *=* 0 but estimating *β*_*T*_). The likelihood function for these models was:

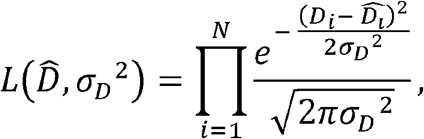

were 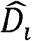is the expected diversity of the linear predictor and *N* is the total number of data points.

### Single nuclei RNA sequencing and data processing

Single cell RNA-Sequencing (scRNA-Seq) was performed on single nuclei suspensions using either the Takara Bio ICELL8 Single Cell System or the 10X Genomics Chromium to prepare cDNA sequencing libraries (Supplementary Table 3). 10x Genomics was used for 35 patients, ICELL8 platform was used for 10 patients. Samples processed on the ICELL8 Single Cell System (Takara Bio) were prepared using the SMARTer ICELL8 3 ‘DE Reagent Kit V2 (Takara Bio, Cat # 640167) from isolated nuclei. DAPI stained nuclei were diluted to a concentration of 60,000 cell/mL in 1x PBS + 1% BSA + 1x Second Diluent + 0.2U SUPERase·In RNase Inhibitor and dispensed onto the ICELL8 3 ‘DE Chip (Takara Bio, Cat# 640143) using the ICELL8 MultiSample NanoDispenser. Single nuclei candidates were selected using the ICELL8 imaging system with CellSelect Software (Takara Bio) selecting for DAPI positive nuclei and reverse transcription and sequencing library preparation was performed according to manufacturer instructions. ICELL8 cDNA sequencing libraries were sequenced at a depth of 200K reads/cell on Illumina HiSeq 2500 (read #1 = 26nt and read #2 = 100nt). ICELL8 scRNA-Seq was performed at the Integrative Genomics Core at City of Hope, Fulgent Genetics, and the High Throughput Genomics Core at Huntsman Cancer Institute (HCI) of University of Utah. Sequence reads were processed with BETSY, which aligned reads to reference genome (GRChg38) using STAR v2.6.0 (95). Counts on gene transcripts were calculated by featureCounts implemented in subread v1.5.2 (96). A gene-barcode matrix was generated for each sample containing counts for each gene in each barcode (cell).

Samples processed on the 10X Genomics Chromium were processed using the Chromium Single Cell 3 ‘V3 Kit (10X Genomics, Cat # 1000075) using isolated nuclei. Single nuclei were diluted to a target of 1,000 cells/µL in 1x PBS + 1.0% BSA + 0.2U/µL SUPERase·In RNase Inhibitor to generate GEM’s prepared at a target of 5,000 cells per sample. Barcoding, reverse transcription, and library preparation were performed according to manufacturer instructions. 10X Genomics generated cDNA libraries were sequenced on Illumina HiSeq 2500 or NovaSeq 6000 instruments using 150 bp paired-end sequencing at a median depth of 34K reads/cell. scRNA-Seq was performed at the Integrative Genomics Core at City of Hope, Fulgent Genetics, and the High Throughput Genomics Core at Huntsman Cancer Institute (HCI) of University of Utah. Sequence reads were processed with CellRanger v3.0.2 using reference genome (GRChg38). A gene-barcode matrix was generated for each sample containing counts of unique molecular identifiers (UMIs) for each gene in each barcode (cell).

### Copy number alteration and subclone analysis from scRNA

Copy number alterations of each cell were estimated based on the count matrix using R package ‘infercnv ‘v1.0.2 (cutoff=0, min_cells_per_gene = 100 or 500, cluster_by_groups=T, HMM=T, analysis_mode=“subclusters”), which uses gene expression intensity to predict copy number changes in tumor cells compared to normal reference cells. A subset of 500 immune cells or stromal cells from this study were used as reference for ‘infercnv ‘ analysis. Cancer cells from each patient were clustered based on the estimated gene copy numbers in each cell using ‘hclust ‘ from R package ‘fastcluster ‘v1.1.25 (97) (method=‘ward.D2’). Cell clusters were visualized in heatmap using R package ‘ComplexHeatmap ‘v2.0.0 (98). Clusters with distinct copy number profiles were defined as subclones for each patient. Single-cell phylogeny was performed based on hierarchical cluster analysis using R package ape v5.4-1 (99). Hierarchical clustering of tumor subclones were visualized using GraPhlAn v1.1.4 (100).

### Cell type classification

An integrative approach was used to classify cell types in samples from 45 patients. First, cell type of each cell was predicted using the R package ‘SingleR ‘v1.0.1 to generate preliminary cell type classifications for all patients. Second, we applied Seurat v3.1.1.9023 Reciprocal PCA integration workflow to 35 patients with 10x scRNA data to integrate cells from different samples (28). Patients with ICELL8 scRNA data were analyzed with standard Seurat workflow (101). Cell clusters were identified using ‘FindClusters ‘method (resolution=0.8) in R package Seurat v3.1.1.9023. Third, each cell cluster was then defined as epithelial cells, stromal cells (fibroblasts, endothelial cells), immune cells (macrophages, T cells, B cells) based on the maximum number of cells annotated as these cell types from ‘SingleR’. Expression profiles of marker genes were used to validate the cell type classifications: epithelial cells (KRT19, CDH1), immune cells (PTPRC), stromal cells (HTRA1, FAP). Finally, epithelial cells were classified into cancer cells and normal epithelial cells based on presence and absence of copy number alteration. Cell type annotation was summarized in Supplementary Table 3.

### Breast cancer intrinsic subtype prediction

Primary molecular subtypes of breast cancer (basal-like, HER2-enriched, luminal A, luminal B) for each cell was predicted based on the log(CPM+1) count matrix by R package ‘genefu ‘ v2.18.0 (102) with default parameters (sbt.model = scmod1). The predicted subtype that has the highest prediction probability was assigned as breast cancer intrinsic subtype of each cell. This analysis was performed only on patients with 10x scRNA data. Patients with ICELL8 scRNA have very few cancer cells, thereby were not included in this analysis.

### Gene Set enrichment analysis

The count matrix of each cell type was filtered to keep genes that expressed 1 or more counts in at least 10 cells. The filtered count matrix was normalized by R package ‘zinbwave ‘v 1.8.0 (27) with total number of counts and gene length and GC-content as covariates (K=2, X=“∼log (total number of counts)”, V=“∼ GC-content + log (gene length)”, epsilon=1000, normalizedValues=TRUE). Single sample Gene Set Enrichment Analysis (ssGSEA) scores of 50 hallmark signatures (MSigDB, hallmark) (36) and 4725 curated pathway signatures (MSigDB, c2) (36) were calculated for each cell based on the normalized count matrix using R package ‘GSVA ‘v1.30.0 (kcdf=“Gaussian”, method=‘ssgsea’) (103).

### Pathway analysis: identifying response related phenotypes

For each ssGSEA pathway in the C2-level and Hallmark pathway signatures, changes in cancer cell pathway activity (x) over time (T) was examined during each treatment arm of the trial (A). Phenotypic changes linked to treatment or differing between sensitive and resistant patients (R) were identified. A random effects model (104) with the following linear predictor and error structure was constructed for each pathway:

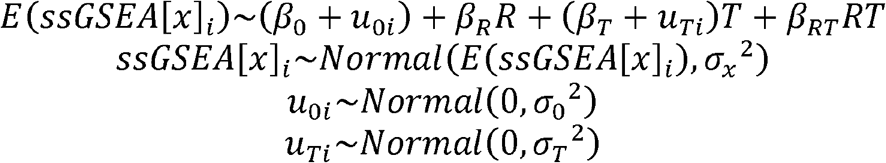

Initial differences in pathway activity between cancer cells from sensitive and resistant tumors, at the pre-treatment time point were captured by the group-specific intercepts (*β*_0_vs *β*_*R*_). Subsequent changes in pathway activity were described by temporal effect terms of sensitive and resistant tumors (*β*_*T*_and *β*_*RT*_). Preexisting individual variability in gene expression and patient specific susceptibility of pathway activation to therapy, were accounted for by allowing the model intercept and temporal effect to vary among patients (*i*) (*u*_0*i*_ and *u*_*Ti*_). Significant differences in pathway activity before or during treatment were identified between treatment arms and patient response groups, using likelihood ratio tests with multiple comparison corrections following Holm’s p-value correction. Compared to false discovery rate (FDR) correction, Holm’s adjustment was more conservative, avoiding the spurious detection of response related phenotypes. The likelihood of the full model was compared against that of nested null models in which: i) no change in pathway activity occurred in sensitive or resistant tumors (fixing *β*_*T*_ *& β*_*RT*_ *= 0*) or ii) equal changes in pathway activity occurred in sensitive and resistant tumors (fixing *β*_*RT*_*=* 0 but estimating *β*_*T*_). The likelihood function for each of these models was:

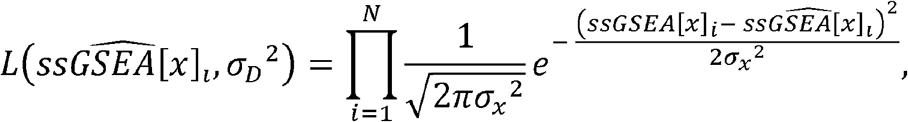

were 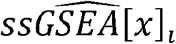 is the expected pathway activity (*E*(*ssGSEA*[*x*]_*i*_)) and *N* is the total number of datapoints.

The significance of pathway activation in sensitive and resistant tumors (non-zero *β*_*T*_and *β*_*RT*_ parameters) was then assessed using a two-tailed t-tests. The Satterthwaite method was applied to perform degree of freedom, t-statistic and p-value calculations for the hierarchical regression model coefficients, using the ‘lmerTest’ R package (105). The hierarchical regression model explicitly described the paired structure in our dataset, with earlier and later samples per patient, and this non-independence of cells within a sample determined the effective residual degrees of freedom. Consequently, the statistical test of whether pathway activity related to tumor growth was controlled for the pairing of earlier and later samples per patient tumor. Detected ssGSEA signatures were classified into functional categories (Supplementary Table 10). Genes contributing to each pathway category were identified from the MSigDB and used for downstream analyzes of each process.

### Assessing the loss of estrogen receptor expression

Changes in single cell ESR1 expression during treatment were assessed using a hierarchical generalized additive model (106). The nonlinear trajectory of ESR1 expression during the trial was described by a thin plate spline. Baseline differences in patients’ ESR1 expression was accounted for by a patient specific random intercept term. Single cell ESR1 expression over time was assessed separately for patients in each treatment arm.

### MAPK signaling network structure analysis: Determining co-regulation of gene sets in the MAPK pathways

Signal transduction, via MAPK pathways, forms a complex network with numerous kinases which crosstalk and regulate the activity of one another. To investigate the co-regulation of signal transduction states of cancer cells, we analyzed the pairwise correlation of MAPK gene expression. Hierarchical clustering of the correlation matrices showed the dichotomy between the expressions of kinases acting in the JNK versus the ERK pathways of signal transduction.

The primary phenotypic variation in MAPK signaling across cancer cells was determined by performing dimension reduction of MAPK gene expression, using UMAP (confirmed by Principle component analysis). To assess phenotypic variation across patients, without large patient samples biasing results, the dataset was initially down sampled to 100 cells per biopsy. The UMAP model was then trained and used to calculate the phenotypic scores of the full dataset. This analysis confirmed that JNK activation status was the primary component of heterogeneity in MAPK signaling state across cancer cells (Supplementary Figure 11). A single cell JNK activation score (high JNK, low ERK) was determined, using the major axis of phenotypic variation, due to the collinearity of gene expression between JNK and ERK genes. For each patient, average scRNAseq gene expression was calculated for subclonal cancer cell populations with different levels of JNK activation (n=40 levels).

### Relating CDK6 expression to JNK activation

We characterized the subclonal relationship between JNK activation and the expression of the key ribociclib target gene, CDK6, accounting for genetic differences in CDK6 copy number amplification status. Resistance phenotypes were examined specifically, by focusing on the transcription profiles of cells at end of treatment. Using generalized additive models, a nonlinear smooth function was fitted describing how the average CDK6 expression changed as the JNK activation of subclonal populations increased. A separate relationship was identified for each arm of therapy and for patients with and without baseline CDK6 genetic copy number amplification (as identified by CNA analysis). The significance of CDK6:JNK relationships in each group were assessed using likelihood ration tests comparing the full model to a null model without a CDK6:JNK association.

### Cell surface receptor activity analysis: Compensation for estrogen receptor loss

The Cell Surface Protein Atlas (47) provides a database of known cell receptor proteins. Genes encoding each protein were identified from the Ensembl database (107) (n=1406). For each patient, we identified genes with either: i) significantly higher/lower than average initial expression, or b) altered expression during treatment, using an ANOVA test. Receptor genes that were consistently identified across patients were determined by identifying those overlapping gene detected by our pathway analysis as well as being detectable in several patients using the ANOVA approach. To determine genes that were activated to compensate for the loss of estrogen signaling during treatment, we correlated estrogen pathway activity with alternative receptor gene expression for cells in each treatment arm and response group.

### Cell cycle reconstruction

We next assessed the cell cycle consequences of endocrine and combination therapy. As individual marker genes are insufficient to resolve cell cycle state (108), we reconstructed the cell cycle, using a set of cell cycle genes and the reCAT model of the cell cycle pseudo-time transitions (55). The cell cycle gene list was obtained from the Biocarta cell cycle pathway (48 genes). This signature was repeatedly detected in the pathway analysis and its predicted changes in cell cycle activity mirrored changes in KI-67 antigen expression and tumor size dynamics.

Discrete cell cycle states were first identified by applying a Gaussian mixture model to the cell cycle gene set. The reconstruction of cell cycle transitions was then formulated as a traveling salesman problem and the shortest cycle that passes through each cell cycle stage was identified based on subclonal distances on a UMAP, using the arbitrary insertion traveling salesman algorithm (109) (Supplementary Figure 16).

Fluctuations in gene expression throughout the cell (including non-cell cycle genes) was recovered using cyclical generalized additive models. The expected expression of each gene, as cells transition through the cell cycle, was described using a smooth and cyclical cubic spline function relating gene expression to cell stage. Three distinct phases of the cell cycle (G0, G1, S/G2) were identified by reclassifying cell states, by applying a Gaussian mixture model to the reconstructed gene expression of cell cycle genes in each stage. Differences in the frequency of cells from a sample found to be in the proliferative S/G2 phase was assessed using a quasi-binomial generalized additive model (106).

Treatment induced differences in the frequency of cells at different phases of the cell cycle were assessed by calculating the proportion of cells in each cell cycle phase and comparing the fraction of cells in a given phase between arms and over time, using logistic regression.

## Author Contributions

J.I.G. contributed study design and coordination, evaluated patient response to therapies, analyzed tumor heterogeneity, identified response related phenotypes using scRNA-Seq ssGSEA pathway analysis, quantified resistance phenotypes using mathematical models, linked genetic copy number alterations to phenotypes, reconstructed cancer cell cycle transition and gene expression, and wrote manuscript. J.C. conducted the bioinformatics pipelines to process DNA and scRNAseq data, performed normalization and cell type classification, conducted structural variation analysis of WGS, determined subclonal tumor structure, analysed WES data, and wrote manuscript. P.C. performed scRNA and WGS experiments. A.O.D, P.S, C.M., M.T., K.K, K.B.W, R.O.R, I.M., L.M.S, A.B. contributed patient samples and contributed to writing the manuscript. F.R.A. developed analyses and models and contributed to writing the manuscript. J.T.C. developed bioinformatics pipelines, performed data management and curation, conducted data analysis and wrote the manuscript. A.L.C. contributed data analysis and study design, provided clinical insight, and contributed to writing the manuscript. Q.K., conceived and coordinated the clinical trial, contributed clinical support and infrastructure and provided clinical data and patient samples as well as contributed to writing the manuscript. A.H.B. designed the research project and analyses, performed scRNA experiments and data analysis, coordinated genomic and mathematical/statistical analyses, and wrote the manuscript.

## Acknowledgements

We thank the anonymous patients from the trial that made this study possible. The High-Throughput Genomics Shared Resource was supported by the NIH Award Number P30CA042014. The content is solely the authors responsibility and does not necessarily represent the official views of the NIH. J.T.C. was supported by a Cancer Prevention Research Institute of Texas Core Facility Support Award (RP170668). A.H.B, J.G., J.C., J.T.C, P.C., and F.A. were supported by the National Cancer Institute of the National Institutes of Health under award number U54CA209978.

## Data availability

Raw single cell RNA-seq data has been deposited in GEO under accession code GSE158724. The DNA-Seq data is available from dbGaP at phs002287.v1.p1 at https://www.ncbi.nlm.nih.gov/projects/gap/cgi-bin/study.cgi?study_id=phs002287.v1.p1. Custom code used in analyses are available on GitHub at https://github.com/U54Bioinformatics/FELINE_project.

## References

1. Harbeck N, Gnant M. Breast cancer. Lancet. 2017;389(10074):1134–50. Epub 2016/11/21. doi: 10.1016/s0140-6736(16)31891-8. PubMed PMID: 27865536.

2. Rani A, Stebbing J, Giamas G, Murphy J. Endocrine Resistance in Hormone Receptor Positive Breast Cancer-From Mechanism to Therapy. Front Endocrinol (Lausanne). 2019;10:245. Epub 2019/06/11. doi: 10.3389/fendo.2019.00245. PubMed PMID: 31178825; PMCID: PMC6543000.

3. Brady SW, McQuerry JA, Qiao Y, Piccolo SR, Shrestha G, Jenkins DF, Layer RM, Pedersen BS, Miller RH, Esch A, Selitsky SR, Parker JS, Anderson LA, Dalley BK, Factor RE, Reddy CB, Boltax JP, Li DY, Moos PJ, Gray JW, Heiser LM, Buys SS, Cohen AL, Johnson WE, Quinlan AR, Marth G, Werner TL, Bild AH. Combating subclonal evolution of resistant cancer phenotypes. Nat Commun. 2017;8(1):1231. Epub 2017/11/03. doi: 10.1038/s41467-017-01174-3. PubMed PMID: 29093439; PMCID: PMC5666005.

4. Zhao H, Zhou L, Shangguan AJ, Bulun SE. Aromatase expression and regulation in breast and endometrial cancer. J Mol Endocrinol. 2016;57(1):R19–33. Epub 2016/04/14. doi: 10.1530/jme-15-0310. PubMed PMID: 27067638; PMCID: PMC5519084.

5. Osborne CK. Tamoxifen in the treatment of breast cancer. N Engl J Med. 1998;339(22):1609-18. Epub 1998/11/26. doi: 10.1056/nejm199811263392207. PubMed PMID: 9828250.

6. Effects of chemotherapy and hormonal therapy for early breast cancer on recurrence and 15-year survival: an overview of the randomised trials. Lancet. 2005;365(9472):1687–717. Epub 2005/05/17. doi: 10.1016/s0140-6736(05)66544-0. PubMed PMID: 15894097.

7. Company ELa. Endocrine Therapy With or Without Abemaciclib (LY2835219) Following Surgery in Participants With Breast Cancer (monarchE) 2017 [updated September 18, 2020; cited 2020 October 13]. Available from: https://www.clinicaltrials.gov/ct2/show/NCT03155997.

8. Pharmaceuticals N. A Trial to Evaluate Efficacy and Safety of Ribociclib With Endocrine Therapy as Adjuvant Treatment in Patients With HR+/HER2-Early Breast Cancer (NATALEE) 2018 [updated October 2, 2020; cited 2020 October 13]. Available from: https://clinicaltrials.gov/ct2/show/NCT03701334.

9. Bertagnolli M ME, DeMichele A, Gnant M. PALbociclib CoLlaborative Adjuvant Study (PALLAS) 2015 [updated March 9, 2020; cited 2020 October 2013]. Available from: https://clinicaltrials.gov/ct2/show/NCT02513394.

10. PENELOPE-B TRIAL OF IBRANCE® (PALBOCICLIB) IN EARLY BREAST CANCER DID NOT MEET PRIMARY ENDPOINT: Pfizer; 2020 [updated October 9, 2020; cited 2020 October 13]. Available from: https://www.pfizer.com/news/press-release/press-release-detail/penelope-b-trial-ibrancer-palbociclib-early-breast-cancer.

11. Johnston S, Puhalla S, Wheatley D, Ring A, Barry P, Holcombe C, Boileau JF, Provencher L, Robidoux A, Rimawi M, McIntosh SA, Shalaby I, Stein RC, Thirlwell M, Dolling D, Morden J, Snowdon C, Perry S, Cornman C, Batten LM, Jeffs LK, Dodson A, Martins V, Modi A, Osborne CK, Pogue-Geile KL, Cheang MCU, Wolmark N, Julian TB, Fisher K, MacKenzie M, Wilcox M, Huang Bartlett C, Koehler M, Dowsett M, Bliss JM, Jacobs SA. Randomized Phase II Study Evaluating Palbociclib in Addition to Letrozole as Neoadjuvant Therapy in Estrogen Receptor-Positive Early Breast Cancer: PALLET Trial. J Clin Oncol. 2019;37(3):178–89. Epub 2018/12/14. doi: 10.1200/JCO.18.01624. PubMed PMID: 30523750.

12. Johnston SRD, Harbeck N, Hegg R, Toi M, Martin M, Shao ZM, Zhang QY, Martinez Rodriguez JL, Campone M, Hamilton E, Sohn J, Guarneri V, Okada M, Boyle F, Neven P, Cortés J, Huober J, Wardley A, Tolaney SM, Cicin I, Smith IC, Frenzel M, Headley D, Wei R, San Antonio B, Hulstijn M, Cox J, O’Shaughnessy J, Rastogi P. Abemaciclib Combined With Endocrine Therapy for the Adjuvant Treatment of HR+, HER2-, Node-Positive, High-Risk, Early Breast Cancer (monarchE). J Clin Oncol. 2020;38(34):3987–98. Epub 2020/09/22. doi: 10.1200/jco.20.02514. PubMed PMID: 32954927; PMCID: PMC7768339.

13. Krzyzanowska MK, Julian JA, Powis M, Howell D, Earle CC, Enright KA, Mittmann N, Trudeau ME, Grunfeld E. Ambulatory Toxicity Management (AToM) in patients receiving adjuvant or neo-adjuvant chemotherapy for early stage breast cancer - a pragmatic cluster randomized trial protocol. BMC Cancer. 2019;19(1):884. Epub 2019/09/07. doi: 10.1186/s12885-019-6099-x. PubMed PMID: 31488084; PMCID: PMC6729066.

14. Weintraub SJ, Chow KN, Luo RX, Zhang SH, He S, Dean DC. Mechanism of active transcriptional repression by the retinoblastoma protein. Nature. 1995;375(6534):812-5. Epub 1995/06/29. doi: 10.1038/375812a0. PubMed PMID: 7596417.

15. Finn RS, Aleshin A, Slamon DJ. Targeting the cyclin-dependent kinases (CDK) 4/6 in estrogen receptor-positive breast cancers. Breast Cancer Res. 2016;18(1):17. Epub 2016/02/10. doi: 10.1186/s13058-015-0661-5. PubMed PMID: 26857361; PMCID: PMC4746893.

16. Portman N, Alexandrou S, Carson E, Wang S, Lim E, Caldon CE. Overcoming CDK4/6 inhibitor resistance in ER-positive breast cancer. Endocr Relat Cancer. 2019;26(1):R15–R30. Epub 2018/11/06. doi: 10.1530/ERC-18-0317. PubMed PMID: 30389903.

17. Sabbah M, Courilleau D, Mester J, Redeuilh G. Estrogen induction of the cyclin D1 promoter: Involvement of a cAMP response-like element. Proceedings of the National Academy of Sciences. 1999;96(20):11217–22. doi: 10.1073/pnas.96.20.11217.

18. Razavi P, Chang MT, Xu G, Bandlamudi C, Ross DS, Vasan N, Cai Y, Bielski CM, Donoghue MTA, Jonsson P, Penson A, Shen R, Pareja F, Kundra R, Middha S, Cheng ML, Zehir A, Kandoth C, Patel R, Huberman K, Smyth LM, Jhaveri K, Modi S, Traina TA, Dang C, Zhang W, Weigelt B, Li BT, Ladanyi M, Hyman DM, Schultz N, Robson ME, Hudis C, Brogi E, Viale A, Norton L, Dickler MN, Berger MF, Iacobuzio-Donahue CA, Chandarlapaty S, Scaltriti M, Reis-Filho JS, Solit DB, Taylor BS, Baselga J. The Genomic Landscape of Endocrine-Resistant Advanced Breast Cancers. Cancer Cell. 2018;34(3):427–38 e6. Epub 2018/09/12. doi: 10.1016/j.ccell.2018.08.008. PubMed PMID: 30205045; PMCID: PMC6327853.

19. Formisano L, Lu Y, Servetto A, Hanker AB, Jansen VM, Bauer JA, Sudhan DR, Guerrero-Zotano AL, Croessmann S, Guo Y, Ericsson PG, Lee K-m, Nixon MJ, Schwarz LJ, Sanders ME, Dugger TC, Cruz MR, Behdad A, Cristofanilli M, Bardia A, O’Shaughnessy J, Nagy RJ, Lanman RB, Solovieff N, He W, Miller M, Su F, Shyr Y, Mayer IA, Balko JM, Arteaga CL. Aberrant FGFR signaling mediates resistance to CDK4/6 inhibitors in ER+ breast cancer. Nature Communications. 2019;10(1):1373. doi: 10.1038/s41467-019-09068-2.

20. Kato S, Endoh H, Masuhiro Y, Kitamoto T, Uchiyama S, Sasaki H, Masushige S, Gotoh Y, Nishida E, Kawashima H, Metzger D, Chambon P. Activation of the estrogen receptor through phosphorylation by mitogen-activated protein kinase. Science. 1995;270(5241):1491-4. Epub 1995/12/01. doi: 10.1126/science.270.5241.1491. PubMed PMID: 7491495.

21. Filardo EJ, Quinn JA, Bland KI, Frackelton AR, Jr. Estrogen-induced activation of Erk-1 and Erk-2 requires the G protein-coupled receptor homolog, GPR30, and occurs via trans-activation of the epidermal growth factor receptor through release of HB-EGF. Mol Endocrinol. 2000;14(10):1649–60. Epub 2000/10/24. doi: 10.1210/mend.14.10.0532. PubMed PMID: 11043579.

22. Bi R, Foy MR, Vouimba R-M, Thompson RF, Baudry M. Cyclic changes in estradiol regulate synaptic plasticity through the MAP kinase pathway. Proceedings of the National Academy of Sciences. 2001;98(23):13391–5. doi: 10.1073/pnas.241507698.

23. Khan Q. Letrozole Plus Ribociclib or Placebo as Neo-adjuvant Therapy in ER-positive, HER2-negative Early Breast Cancer (FELINE) 2016 [updated August 29, 2019; cited 2020 September 11]. Available from: https://clinicaltrials.gov/ct2/show/NCT02712723.

24. Griffiths JI, Chen J, O’Dea A, Sharma P, Winblad O, Trivedi M, Kalinsky K, Wisinski K, O’Regan R, Makhoul I, Spring L, Bardia A, Yuan Y, Jahanzeb M, Adler F, Cohen A, Bild A, Khan Q. Reconstructing tumor trajectories during therapy through integration of multiple measurement modalities. bioRxiv. 2021:2021.01.14.426737. doi: 10.1101/2021.01.14.426737.

25. Berg WA, Blume JD, Cormack JB, Mendelson EB, Lehrer D, Böhm-Vélez M, Pisano ED, Jong RA, Evans WP, Morton MJ, Mahoney MC, Larsen LH, Barr RG, Farria DM, Marques HS, Boparai K. Combined screening with ultrasound and mammography vs mammography alone in women at elevated risk of breast cancer. Jama. 2008;299(18):2151–63. Epub 2008/05/15. doi: 10.1001/jama.299.18.2151. PubMed PMID: 18477782; PMCID: PMC2718688.

26. Marinovich ML, Houssami N, Macaskill P, Sardanelli F, Irwig L, Mamounas EP, von Minckwitz G, Brennan ME, Ciatto S. Meta-analysis of magnetic resonance imaging in detecting residual breast cancer after neoadjuvant therapy. J Natl Cancer Inst. 2013;105(5):321–33. Epub 2013/01/09. doi: 10.1093/jnci/djs528. PubMed PMID: 23297042.

27. Risso D, Perraudeau F, Gribkova S, Dudoit S, Vert JP. A general and flexible method for signal extraction from single-cell RNA-seq data. Nat Commun. 2018;9(1):284. Epub 2018/01/20. doi: 10.1038/s41467-017-02554-5. PubMed PMID: 29348443; PMCID: PMC5773593.

28. Stuart T, Butler A, Hoffman P, Hafemeister C, Papalexi E, Mauck WM, 3rd, Hao Y, Stoeckius M, Smibert P, Satija R. Comprehensive Integration of Single-Cell Data. Cell. 2019;177(7):1888–902 e21. Epub 2019/06/11. doi: 10.1016/j.cell.2019.05.031. PubMed PMID: 31178118; PMCID: PMC6687398.

29. Tickle TI GC, Brown M, Haas B. inferCNV of the Trinity CTAT Project 2019 [cited 2020 September 14]. Available from: https://github.com/broadinstitute/inferCNV.

30. van der Maaten L, Hinton G. Visualizing Data using t-SNE. Journal of Machine Learning Research. 2008;9:2579–605.

31. Aran D, Looney AP, Liu L, Wu E, Fong V, Hsu A, Chak S, Naikawadi RP, Wolters PJ, Abate AR, Butte AJ, Bhattacharya M. Reference-based analysis of lung single-cell sequencing reveals a transitional profibrotic macrophage. Nature Immunology. 2019;20(2):163–72. doi: 10.1038/s41590-018-0276-y.

32. Roth A, Khattra J, Yap D, Wan A, Laks E, Biele J, Ha G, Aparicio S, Bouchard-Côté A, Shah SP. PyClone: statistical inference of clonal population structure in cancer. Nat Methods. 2014;11(4):396–8. Epub 2014/03/19. doi: 10.1038/nmeth.2883. PubMed PMID: 24633410; PMCID: PMC4864026.

33. Kim C, Gao R, Sei E, Brandt R, Hartman J, Hatschek T, Crosetto N, Foukakis T, Navin NE. Chemoresistance Evolution in Triple-Negative Breast Cancer Delineated by Single-Cell Sequencing. Cell. 2018;173(4):879–93 e13. Epub 2018/04/24. doi: 10.1016/j.cell.2018.03.041. PubMed PMID: 29681456; PMCID: PMC6132060.

34. Morris EK, Caruso T, Buscot F, Fischer M, Hancock C, Maier TS, Meiners T, Müller C, Obermaier E, Prati D, Socher SA, Sonnemann I, Wäschke N, Wubet T, Wurst S, Rillig MC. Choosing and using diversity indices: insights for ecological applications from the German Biodiversity Exploratories. Ecol Evol. 2014;4(18):3514–24. Epub 2014/12/06. doi: 10.1002/ece3.1155. PubMed PMID: 25478144; PMCID: PMC4224527.

35. Barbie DA, Tamayo P, Boehm JS, Kim SY, Moody SE, Dunn IF, Schinzel AC, Sandy P, Meylan E, Scholl C, Fröhling S, Chan EM, Sos ML, Michel K, Mermel C, Silver SJ, Weir BA, Reiling JH, Sheng Q, Gupta PB, Wadlow RC, L.H, Hoersch S, Wittner BS, Ramaswamy S, Livingston DM, Sabatini DM, Meyerson M, Thomas RK, Lander ES, Mesirov JP, Root DE, Gilliland DG, Jacks T, Hahn WC. Systematic RNA interference reveals that oncogenic KRAS-driven cancers require TBK1. Nature. 2009;462(7269):108–12. Epub 2009/10/23. doi: 10.1038/nature08460. PubMed PMID: 19847166; PMCID: PMC2783335.

36. Liberzon A, Subramanian A, Pinchback R, Thorvaldsdóttir H, Tamayo P, Mesirov JP. Molecular signatures database (MSigDB) 3.0. Bioinformatics. 2011;27(12):1739–40. Epub 2011/05/07. doi: 10.1093/bioinformatics/btr260. PubMed PMID: 21546393; PMCID: PMC3106198.

37. Chen SY, Feng Z, Yi X. A general introduction to adjustment for multiple comparisons. J Thorac Dis. 2017;9(6):1725–9. Epub 2017/07/26. doi: 10.21037/jtd.2017.05.34. PubMed PMID: 28740688; PMCID: PMC5506159.

38. Iwase H, Greenman JM, Barnes DM, Bobrow L, Hodgson S, Mathew CG. Loss of heterozygosity of the oestrogen receptor gene in breast cancer. Br J Cancer. 1995;71(3):448-50. Epub 1995/03/01. doi: 10.1038/bjc.1995.91. PubMed PMID: 7880722; PMCID: PMC2033633.

39. Herynk MH, Fuqua SA. Estrogen receptor mutations in human disease. Endocr Rev. 2004;25(6):869–98. Epub 2004/12/08. doi: 10.1210/er.2003-0010. PubMed PMID: 15583021.

40. Reddy KB, Mangold GL, Tandon AK, Yoneda T, Mundy GR, Zilberstein A, Osborne CK. Inhibition of Breast Cancer Cell Growth <em>in Vitro</em> by a Tyrosine Kinase Inhibitor. Cancer Research. 1992;52(13):3636–41.

41. Kahlert S, Nuedling S, van Eickels M, Vetter H, Meyer R, Grohe C. Estrogen receptor alpha rapidly activates the IGF-1 receptor pathway. J Biol Chem. 2000;275(24):18447–53. Epub 2000/04/06. doi: 10.1074/jbc.M910345199. PubMed PMID: 10749889.

42. Dupont J, Karas M, LeRoith D. The potentiation of estrogen on insulin-like growth factor I action in MCF-7 human breast cancer cells includes cell cycle components. J Biol Chem. 2000;275(46):35893–901. Epub 2000/09/01. doi: 10.1074/jbc.M006741200. PubMed PMID: 10967123.

43. Segars JH, Driggers PH. Estrogen action and cytoplasmic signaling cascades. Part I: membrane-associated signaling complexes. Trends Endocrinol Metab. 2002;13(8):349–54. Epub 2002/09/10. doi: 10.1016/s1043-2760(02)00633-1. PubMed PMID: 12217492; PMCID: PMC4137481.

44. Arpino G, Wiechmann L, Osborne CK, Schiff R. Crosstalk between the estrogen receptor and the HER tyrosine kinase receptor family: molecular mechanism and clinical implications for endocrine therapy resistance. Endocr Rev. 2008;29(2):217–33. Epub 2008/01/25. doi: 10.1210/er.2006-0045. PubMed PMID: 18216219; PMCID: PMC2528847.

45. Thomas LW, Lam C, Edwards SW. Mcl-1; the molecular regulation of protein function. FEBS Letters. 2010;584(14):2981–9. doi: 10.1016/j.febslet.2010.05.061.

46. Hirata Y, Sugie A, Matsuda A, Matsuda S, Koyasu S. TAK1–JNK Axis Mediates Survival Signal through Mcl1 Stabilization in Activated T Cells. The Journal of Immunology. 2013;190(9):4621–6. doi: 10.4049/jimmunol.1202809.

47. Bausch-Fluck D, Hofmann A, Bock T, Frei AP, Cerciello F, Jacobs A, Moest H, Omasits U, Gundry RL, Yoon C, Schiess R, Schmidt A, Mirkowska P, Hartlova A, Van Eyk JE, Bourquin JP, Aebersold R, Boheler KR, Zandstra P, Wollscheid B. A mass spectrometric-derived cell surface protein atlas. PLoS One. 2015;10(3):e0121314. Epub 2015/04/22. doi: 10.1371/journal.pone.0121314. PubMed PMID: 25894527; PMCID: PMC4404347.

48. Zhu Y, Sullivan LL, Nair SS, Williams CC, Pandey AK, Marrero L, Vadlamudi RK, Jones FE. Coregulation of estrogen receptor by ERBB4/HER4 establishes a growth-promoting autocrine signal in breast tumor cells. Cancer Res. 2006;66(16):7991–8. Epub 2006/08/17. doi: 10.1158/0008-5472.Can-05-4397. PubMed PMID: 16912174.

49. Tiong KH, Mah LY, Leong CO. Functional roles of fibroblast growth factor receptors (FGFRs) signaling in human cancers. Apoptosis. 2013;18(12):1447–68. Epub 2013/08/01. doi: 10.1007/s10495-013-0886-7. PubMed PMID: 23900974; PMCID: PMC3825415.

50. Brewer JR, Mazot P, Soriano P. Genetic insights into the mechanisms of Fgf signaling. Genes Dev. 2016;30(7):751–71. Epub 2016/04/03. doi: 10.1101/gad.277137.115. PubMed PMID: 27036966; PMCID: PMC4826393.

51. Odawara H, Iwasaki T, Horiguchi J, Rokutanda N, Hirooka K, Miyazaki W, Koibuchi Y, Shimokawa N, Iino Y, Takeyoshi I, Koibuchi N. Activation of aromatase expression by retinoic acid receptor-related orphan receptor (ROR) alpha in breast cancer cells: identification of a novel ROR response element. J Biol Chem. 2009;284(26):17711–9. Epub 2009/05/15. doi: 10.1074/jbc.M109.009241. PubMed PMID: 19439415; PMCID: PMC2719410.

52. Malumbres M, Barbacid M. Cell cycle, CDKs and cancer: a changing paradigm. Nat Rev Cancer. 2009;9(3):153–66. Epub 2009/02/25. doi: 10.1038/nrc2602. PubMed PMID: 19238148.

53. Lapenna S, Giordano A. Cell cycle kinases as therapeutic targets for cancer. Nat Rev Drug Discov. 2009;8(7):547–66. Epub 2009/07/02. doi: 10.1038/nrd2907. PubMed PMID: 19568282.

54. Vermeulen K, Van Bockstaele DR, Berneman ZN. The cell cycle: a review of regulation, deregulation and therapeutic targets in cancer. Cell Proliferation. 2003;36(3):131–49. doi: 10.1046/j.1365-2184.2003.00266.x.

55. Liu Z, Lou H, Xie K, Wang H, Chen N, Aparicio OM, Zhang MQ, Jiang R, Chen T. Reconstructing cell cycle pseudo time-series via single-cell transcriptome data. Nat Commun. 2017;8(1):22. Epub 2017/06/21. doi: 10.1038/s41467-017-00039-z. PubMed PMID: 28630425; PMCID: PMC5476636.

56. Hortobagyi GN, Stemmer SM, Burris HA, Yap YS, Sonke GS, Paluch-Shimon S, Campone M, Petrakova K, Blackwell KL, Winer EP, Janni W, Verma S, Conte P, Arteaga CL, Cameron DA, Mondal S, Su F, Miller M, Elmeliegy M, Germa C, O’Shaughnessy J. Updated results from MONALEESA-2, a phase III trial of first-line ribociclib plus letrozole versus placebo plus letrozole in hormone receptor-positive, HER2-negative advanced breast cancer. Ann Oncol. 2018;29(7):1541–7. Epub 2018/05/03. doi: 10.1093/annonc/mdy155. PubMed PMID: 29718092.

57. Finn RS, Martin M, Rugo HS, Jones S, Im SA, Gelmon K, Harbeck N, Lipatov ON, Walshe JM, Moulder S, Gauthier E, Lu DR, Randolph S, Diéras V, Slamon DJ. Palbociclib and Letrozole in Advanced Breast Cancer. N Engl J Med. 2016;375(20):1925–36. Epub 2016/12/14. doi: 10.1056/NEJMoa1607303. PubMed PMID: 27959613.

58. Goetz MP, Toi M, Campone M, Sohn J, Paluch-Shimon S, Huober J, Park IH, Trédan O, Chen SC, Manso L, Freedman OC, Garnica Jaliffe G, Forrester T, Frenzel M, Barriga S, Smith IC, Bourayou N, Di Leo A. MONARCH 3: Abemaciclib As Initial Therapy for Advanced Breast Cancer. J Clin Oncol. 2017;35(32):3638–46. Epub 2017/10/03. doi: 10.1200/jco.2017.75.6155. PubMed PMID: 28968163.

59. Osborne CK, Schiff R. Mechanisms of endocrine resistance in breast cancer. Annu Rev Med. 2011;62:233–47. Epub 2010/10/05. doi: 10.1146/annurev-med-070909-182917. PubMed PMID: 20887199; PMCID: PMC3656649.

60. Ichijo H, Nishida E, Irie K, ten Dijke P, Saitoh M, Moriguchi T, Takagi M, Matsumoto K, Miyazono K, Gotoh Y. Induction of apoptosis by ASK1, a mammalian MAPKKK that activates SAPK/JNK and p38 signaling pathways. Science. 1997;275(5296):90-4. Epub 1997/01/03. doi: 10.1126/science.275.5296.90. PubMed PMID: 8974401.

61. Tobiume K, Matsuzawa A, Takahashi T, Nishitoh H, Morita K, Takeda K, Minowa O, Miyazono K, Noda T, Ichijo H. ASK1 is required for sustained activations of JNK/p38 MAP kinases and apoptosis. EMBO Rep. 2001;2(3):222–8. Epub 2001/03/27. doi: 10.1093/embo-reports/kve046. PubMed PMID: 11266364; PMCID: PMC1083842.

62. Jacinto E, Facchinetti V, Liu D, Soto N, Wei S, Jung SY, Huang Q, Qin J, Su B. SIN1/MIP1 Maintains rictor-mTOR Complex Integrity and Regulates Akt Phosphorylation and Substrate Specificity. Cell. 2006;127(1):125–37. doi: https://doi.org/10.1016/j.cell.2006.08.033.

63. Makino C, Sano Y, Shinagawa T, Millar JBA, Ishii S. Sin1 binds to both ATF-2 and p38 and enhances ATF-2-dependent transcription in an SAPK signaling pathway. Genes Cells. 2006;11(11):1239–51. doi: 10.1111/j.1365-2443.2006.01016.x. PubMed PMID: 17054722.

64. Zeke A, Misheva M, Reményi A, Bogoyevitch MA. JNK Signaling: Regulation and Functions Based on Complex Protein-Protein Partnerships. Microbiol Mol Biol Rev. 2016;80(3):793–835. Epub 2016/07/29. doi: 10.1128/mmbr.00043-14. PubMed PMID: 27466283; PMCID: PMC4981676.

65. Dhanasekaran DN, Reddy EP. JNK-signaling: A multiplexing hub in programmed cell death. Genes Cancer. 2017;8(9-10):682–94. Epub 2017/12/14. doi: 10.18632/genesandcancer.155. PubMed PMID: 29234486; PMCID: PMC5724802.

66. La Marca JE, Richardson HE. Two-Faced: Roles of JNK Signalling During Tumourigenesis in the Drosophila Model. Frontiers in Cell and Developmental Biology. 2020;8(42). doi: 10.3389/fcell.2020.00042.

67. Pinal N, Calleja M, Morata G. Pro-apoptotic and pro-proliferation functions of the JNK pathway of Drosophila: roles in cell competition, tumorigenesis and regeneration. Open Biology. 2019;9(3):180256. doi: doi:10.1098/rsob.180256.

68. Colleoni B, Paternot S, Pita JM, Bisteau X, Coulonval K, Davis RJ, Raspé E, Roger PP. JNKs function as CDK4-activating kinases by phosphorylating CDK4 and p21. Oncogene. 2017;36(30):4349–61. Epub 2017/04/04. doi: 10.1038/onc.2017.7. PubMed PMID: 28368408; PMCID: PMC5537611.

69. Zhang JY, Tao S, Kimmel R, Khavari PA. CDK4 regulation by TNFR1 and JNK is required for NF-κB–mediated epidermal growth control. Journal of Cell Biology. 2005;168(4):561–6. doi: 10.1083/jcb.200411060.

70. Khiem D, Cyster JG, Schwarz JJ, Black BL. A p38 MAPK-MEF2C pathway regulates B-cell proliferation. Proceedings of the National Academy of Sciences. 2008;105(44):17067–72. doi: 10.1073/pnas.0804868105.

71. Thornton TM, Rincon M. Non-classical p38 map kinase functions: cell cycle checkpoints and survival. Int J Biol Sci. 2009;5(1):44–51. Epub 2009/01/23. doi: 10.7150/ijbs.5.44. PubMed PMID: 19159010; PMCID: PMC2610339.

72. Hafner M, Mills CE, Subramanian K, Chen C, Chung M, Boswell SA, Everley RA, Liu C, Walmsley CS, Juric D, Sorger PK. Multiomics Profiling Establishes the Polypharmacology of FDA-Approved CDK4/6 Inhibitors and the Potential for Differential Clinical Activity. Cell Chem Biol. 2019;26(8):1067-80.e8. Epub 2019/06/11. doi: 10.1016/j.chembiol.2019.05.005. PubMed PMID: 31178407; PMCID: PMC6936329.

73. Piccart-Gebhart M, Holmes E, Baselga J, de Azambuja E, Dueck AC, Viale G, Zujewski JA, Goldhirsch A, Armour A, Pritchard KI, McCullough AE, Dolci S, McFadden E, Holmes AP, Tonghua L, Eidtmann H, Dinh P, Di Cosimo S, Harbeck N, Tjulandin S, Im YH, Huang CS, Diéras V, Hillman DW, Wolff AC, Jackisch C, Lang I, Untch M, Smith I, Boyle F, Xu B, Gomez H, Suter T, Gelber RD, Perez EA. Adjuvant Lapatinib and Trastuzumab for Early Human Epidermal Growth Factor Receptor 2-Positive Breast Cancer: Results From the Randomized Phase III Adjuvant Lapatinib and/or Trastuzumab Treatment Optimization Trial. J Clin Oncol. 2016;34(10):1034–42. Epub 2015/11/26. doi: 10.1200/jco.2015.62.1797. PubMed PMID: 26598744; PMCID: PMC4872016 online at www.jco.org. Author contributions are found at the end of this article.

74. von Minckwitz G, Procter M, de Azambuja E, Zardavas D, Benyunes M, Viale G, Suter T, Arahmani A, Rouchet N, Clark E, Knott A, Lang I, Levy C, Yardley DA, Bines J, Gelber RD, Piccart M, Baselga J. Adjuvant Pertuzumab and Trastuzumab in Early HER2-Positive Breast Cancer. N Engl J Med. 2017;377(2):122–31. Epub 2017/06/06. doi: 10.1056/NEJMoa1703643. PubMed PMID: 28581356; PMCID: PMC5538020.

75. Ruíz-Borrego M, Guerrero-Zotano A, Bermejo B, Ramos M, Cruz J, Baena-Cañada JM, Cirauqui B, Rodríguez-Lescure Á, Alba E, Martínez-Jáñez N, Muñoz M, Antolín S, Álvarez I, Del Barco S, Sevillano E, Chacón JI, Antón A, Escudero MJ, Ruiz V, Carrasco E, Martín M. Phase III evaluating the addition of fulvestrant (F) to anastrozole (A) as adjuvant therapy in postmenopausal women with hormone receptor-positive HER2-negative (HR+/HER2-) early breast cancer (EBC): results from the GEICAM/2006-10 study. Breast Cancer Res Treat. 2019;177(1):115–25. Epub 2019/06/04. doi: 10.1007/s10549-019-05296-8. PubMed PMID: 31152327.

76. Dowsett M, Smith IE, Ebbs SR, Dixon JM, Skene A, Griffith C, Boeddinghaus I, Salter J, Detre S, Hills M, Ashley S, Francis S, Walsh G. Short-term changes in Ki-67 during neoadjuvant treatment of primary breast cancer with anastrozole or tamoxifen alone or combined correlate with recurrence-free survival. Clin Cancer Res. 2005;11(2 Pt 2):951s–8s. Epub 2005/02/11. PubMed PMID: 15701892.

77. Zaman K, Winterhalder R, Mamot C, Hasler-Strub U, Rochlitz C, Mueller A, Berset C, Wiliders H, Perey L, Rudolf CB, Hawle H, Rondeau S, Neven P. Fulvestrant with or without selumetinib, a MEK 1/2 inhibitor, in breast cancer progressing after aromatase inhibitor therapy: a multicentre randomised placebo-controlled double-blind phase II trial, SAKK 21/08. Eur J Cancer. 2015;51(10):1212–20. Epub 2015/04/22. doi: 10.1016/j.ejca.2015.03.016. PubMed PMID: 25892646.

78. Cicenas J, Zalyte E, Rimkus A, Dapkus D, Noreika R, Urbonavicius S. JNK, p38, ERK, and SGK1 Inhibitors in Cancer. Cancers (Basel). 2017;10(1). Epub 2017/12/22. doi: 10.3390/cancers10010001. PubMed PMID: 29267206; PMCID: PMC5789351.

79. Navid S, Fan C, P OF-V, Generali D, Li Y. The Fibroblast Growth Factor Receptors in Breast Cancer: from Oncogenesis to Better Treatments. Int J Mol Sci. 2020;21(6). Epub 2020/03/20. doi: 10.3390/ijms21062011. PubMed PMID: 32188012; PMCID: PMC7139621.

80. Roskoski R. Small molecule inhibitors targeting the EGFR/ErbB family of protein-tyrosine kinases in human cancers. Pharmacological Research. 2019;139:395–411. doi: https://doi.org/10.1016/j.phrs.2018.11.014.

81. Chen X, Chang JT. Planning bioinformatics workflows using an expert system. Bioinformatics. 2017;33(8):1210–5. Epub 2017/01/06. doi: 10.1093/bioinformatics/btw817. PubMed PMID: 28052928; PMCID: PMC5860174.

82. Bolger AM, Lohse M, Usadel B. Trimmomatic: a flexible trimmer for Illumina sequence data. Bioinformatics. 2014;30(15):2114–20. Epub 2014/04/04. doi: 10.1093/bioinformatics/btu170. PubMed PMID: 24695404; PMCID: PMC4103590.

83. Li H, Durbin R. Fast and accurate short read alignment with Burrows-Wheeler transform. Bioinformatics. 2009;25(14):1754–60. Epub 2009/05/20. doi: 10.1093/bioinformatics/btp324. PubMed PMID: 19451168; PMCID: PMC2705234.

84. Li H, Durbin R. Fast and accurate long-read alignment with Burrows-Wheeler transform. Bioinformatics. 2010;26(5):589–95. Epub 2010/01/19. doi: 10.1093/bioinformatics/btp698. PubMed PMID: 20080505; PMCID: PMC2828108.

85. Li H. Aligning sequence reads, clone sequences and assembly contigs with BWA-MEM. arXiv preprint arXiv:13033997. 2013.

86. Picard Toolkit: Broad Institute, GitHub Repository; 2019 [cited 2020 September 18]. Available from: http://broadinstitute.github.io/picard/.

87. McKenna A, Hanna M, Banks E, Sivachenko A, Cibulskis K, Kernytsky A, Garimella K, Altshuler D, Gabriel S, Daly M, DePristo MA. The Genome Analysis Toolkit: a MapReduce framework for analyzing next-generation DNA sequencing data. Genome Res. 2010;20(9):1297–303. Epub 2010/07/21. doi: 10.1101/gr.107524.110. PubMed PMID: 20644199; PMCID: PMC2928508.

88. Kim S, Scheffler K, Halpern AL, Bekritsky MA, Noh E, Källberg M, Chen X, Kim Y, Beyter D, Krusche P, Saunders CT. Strelka2: fast and accurate calling of germline and somatic variants. Nat Methods. 2018;15(8):591–4. Epub 2018/07/18. doi: 10.1038/s41592-018-0051-x. PubMed PMID: 30013048.

89. Koboldt DC, Zhang Q, Larson DE, Shen D, McLellan MD, Lin L, Miller CA, Mardis ER, Ding L, Wilson RK. VarScan 2: somatic mutation and copy number alteration discovery in cancer by exome sequencing. Genome Res. 2012;22(3):568–76. Epub 2012/02/04. doi: 10.1101/gr.129684.111. PubMed PMID: 22300766; PMCID: PMC3290792.

90. Wang K, Li M, Hakonarson H. ANNOVAR: functional annotation of genetic variants from high-throughput sequencing data. Nucleic Acids Res. 2010;38(16):e164. Epub 2010/07/06. doi: 10.1093/nar/gkq603. PubMed PMID: 20601685; PMCID: PMC2938201.

91. Shen R, Seshan VE. FACETS: allele-specific copy number and clonal heterogeneity analysis tool for high-throughput DNA sequencing. Nucleic Acids Res. 2016;44(16):e131. Epub 2016/06/09. doi: 10.1093/nar/gkw520. PubMed PMID: 27270079; PMCID: PMC5027494.

92. Favero F, Joshi T, Marquard AM, Birkbak NJ, Krzystanek M, Li Q, Szallasi Z, Eklund AC. Sequenza: allele-specific copy number and mutation profiles from tumor sequencing data. Ann Oncol. 2015;26(1):64–70. Epub 2014/10/17. doi: 10.1093/annonc/mdu479. PubMed PMID: 25319062; PMCID: PMC4269342.

93. Dang HX, White BS, Foltz SM, Miller CA, Luo J, Fields RC, Maher CA. ClonEvol: clonal ordering and visualization in cancer sequencing. Ann Oncol. 2017;28(12):3076–82. Epub 2017/09/28. doi: 10.1093/annonc/mdx517. PubMed PMID: 28950321; PMCID: PMC5834020.

94. Miller CA, McMichael J, Dang HX, Maher CA, Ding L, Ley TJ, Mardis ER, Wilson RK. Visualizing tumor evolution with the fishplot package for R. BMC Genomics. 2016;17(1):880. Epub 2016/11/09. doi: 10.1186/s12864-016-3195-z. PubMed PMID: 27821060; PMCID: PMC5100182.

95. Dobin A, Davis CA, Schlesinger F, Drenkow J, Zaleski C, Jha S, Batut P, Chaisson M, Gingeras TR. STAR: ultrafast universal RNA-seq aligner. Bioinformatics. 2013;29(1):15–21. Epub 2012/10/30. doi: 10.1093/bioinformatics/bts635. PubMed PMID: 23104886; PMCID: PMC3530905.

96. Liao Y, Smyth GK, Shi W. featureCounts: an efficient general purpose program for assigning sequence reads to genomic features. Bioinformatics. 2014;30(7):923–30. Epub 2013/11/15. doi: 10.1093/bioinformatics/btt656. PubMed PMID: 24227677.

97. Müllner D. fastcluster: Fast Hierarchical, Agglomerative Clustering Routines for R and Pythoin. Journal of Statistical Software. 2013;53(no 9):1–18.

98. Gu Z, Eils R, Schlesner M. Complex heatmaps reveal patterns and correlations in multidimensional genomic data. Bioinformatics. 2016;32(18):2847–9. Epub 2016/05/22. doi: 10.1093/bioinformatics/btw313. PubMed PMID: 27207943.

99. Paradis E, Claude J, Strimmer K. APE: Analyses of Phylogenetics and Evolution in R language. Bioinformatics. 2004;20(2):289–90. Epub 2004/01/22. doi: 10.1093/bioinformatics/btg412. PubMed PMID: 14734327.

100. Asnicar F, Weingart G, Tickle TL, Huttenhower C, Segata N. Compact graphical representation of phylogenetic data and metadata with GraPhlAn. PeerJ. 2015;3:e1029. Epub 2015/07/15. doi: 10.7717/peerj.1029. PubMed PMID: 26157614; PMCID: PMC4476132.

101. Seurat - Guided Clustering Tutorial 2020 [updated September 9, 2020; cited 2020 September 18]. Available from: https://satijalab.org/seurat/v3.2/pbmc3k_tutorial.html.

102. Gendoo DM, Ratanasirigulchai N, Schröder MS, Paré L, Parker JS, Prat A, Haibe-Kains B. Genefu: an R/Bioconductor package for computation of gene expression-based signatures in breast cancer. Bioinformatics. 2016;32(7):1097–9. Epub 2015/11/27. doi: 10.1093/bioinformatics/btv693. PubMed PMID: 26607490; PMCID: PMC6410906.

103. Hänzelmann S, Castelo R, Guinney J. GSVA: gene set variation analysis for microarray and RNA-seq data. BMC Bioinformatics. 2013;14:7. Epub 2013/01/18. doi: 10.1186/1471-2105-14-7. PubMed PMID: 23323831; PMCID: PMC3618321.

104. Bolker BM, Brooks ME, Clark CJ, Geange SW, Poulsen JR, Stevens MH, White JS. Generalized linear mixed models: a practical guide for ecology and evolution. Trends Ecol Evol. 2009;24(3):127–35. Epub 2009/02/03. doi: 10.1016/j.tree.2008.10.008. PubMed PMID: 19185386.

105. Kuznetsova A, Brockhoff PB, Christensen RHB. lmerTest Package: Tests in Linear Mixed Effects Models. 2017. 2017;82(13):26. Epub 2017-11-29. doi: 10.18637/jss.v082.i13.

106. Pedersen EJ, Miller DL, Simpson GL, Ross N. Hierarchical generalized additive models in ecology: an introduction with mgcv. PeerJ. 2019;7:e6876. Epub 2019/06/11. doi: 10.7717/peerj.6876. PubMed PMID: 31179172; PMCID: PMC6542350.

107. Yates AD, Achuthan P, Akanni W, Allen J, Allen J, Alvarez-Jarreta J, Amode MR, Armean IM, Azov AG, Bennett R, Bhai J, Billis K, Boddu S, Marugán JC, Cummins C, Davidson C, Dodiya K, Fatima R, Gall A, Giron CG, Gil L, Grego T, Haggerty L, Haskell E, Hourlier T, Izuogu OG, Janacek SH, Juettemann T, Kay M, Lavidas I, Le T, Lemos D, Martinez JG, Maurel T, McDowall M, McMahon A, Mohanan S, Moore B, Nuhn M, Oheh DN, Parker A, Parton A, Patricio M, Sakthivel MP, Abdul Salam AI, Schmitt BM, Schuilenburg H, Sheppard D, Sycheva M, Szuba M, Taylor K, Thormann A, Threadgold G, Vullo A, Walts B, Winterbottom A, Zadissa A, Chakiachvili M, Flint B, Frankish A, Hunt SE, G II, Kostadima M, Langridge N, Loveland JE, Martin FJ, Morales J, Mudge JM, Muffato M, Perry E, Ruffier M, Trevanion SJ, Cunningham F, Howe KL, Zerbino DR, Flicek P. Ensembl 2020. Nucleic Acids Res. 2020;48(D1):D682–d8. Epub 2019/11/07. doi: 10.1093/nar/gkz966. PubMed PMID: 31691826; PMCID: PMC7145704.

108. Buettner F, Natarajan KN, Casale FP, Proserpio V, Scialdone A, Theis FJ, Teichmann SA, Marioni JC, Stegle O. Computational analysis of cell-to-cell heterogeneity in single-cell RNA-sequencing data reveals hidden subpopulations of cells. Nat Biotechnol. 2015;33(2):155–60. Epub 2015/01/20. doi: 10.1038/nbt.3102. PubMed PMID: 25599176.

109. Hahsler M, Hornik K. TSP—Infrastructure for the Traveling Salesperson Problem. 2007. 2007;23(2):21. Epub 2007-11-21. doi: 10.18637/jss.v023.i02.

