## Supplementary Materials for "Convergent evolution of resistance pathways during early stage breast cancer treatment with combination cell cycle (CDK) and endocrine inhibitors"

**Affiliations:**

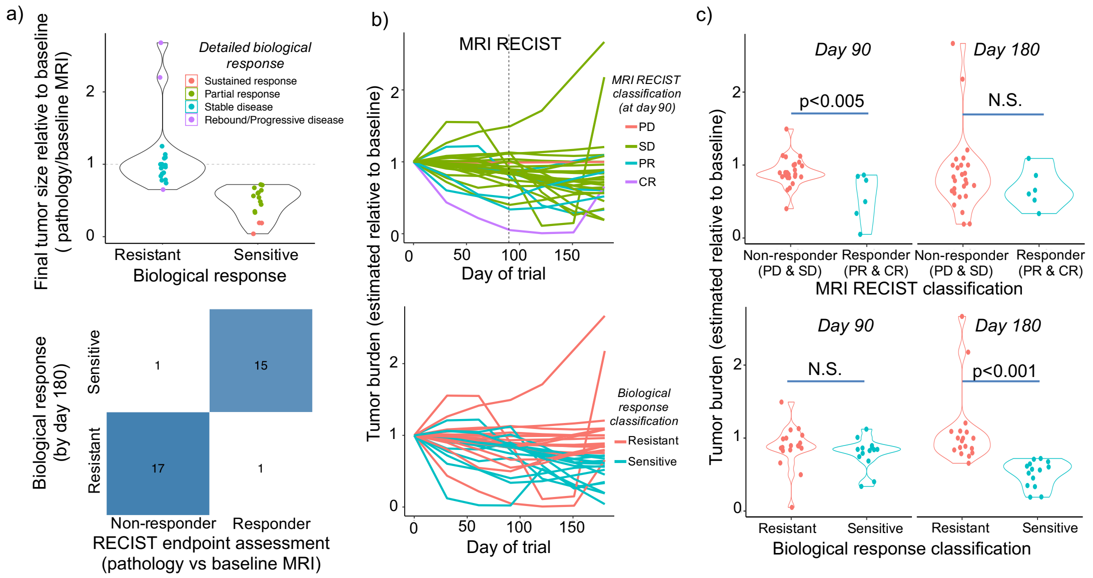


**Supplementary Figure 1. Classification of patient tumors as sensitive or resistant to treatment, reflecting changes in tumor size observed at pathology relative to baseline. Reconstructed trajectories of tumor burden are consistent with results of RECIST 1.1 MRI assessment at day 90 and allow sensitive and resistant tumors to be distinguished at end of treatment (day 180). a,** Changes in tumor size during therapy for tumors classified as sensitive or resistant. Tumor growth (y-axis) calculated directly from data as the proportion tumor remaining at end of trial (final observed tumor size at pathology/baseline MRI tumor measurement). Values <1 indicate tumor shrinkage, whilst values>1 indicate an increase in size (Dashed horizontal line = no change in size during trial). A detailed biological response classification was determined by classifying tumors with similar trajectories using a Gaussian mixture model (colors). Sustained or partial responses were grouped and defined as sensitive tumors, whilst those with stable, progressive or rebound disease were classified as resistant tumors. The changes in tumor size is highly significantly different between resistance categories (t=4.45, p<0.001). Violins show the distinct distribution of tumor growth observed across patients. Heatmap shows the strong agreement in the end of treatment classification obtained by classifying trajectories of tumor growth vs simple pathology/baseline MRI RECIST assessment of change in size during trial. **b,** Spiderplots show the reconstructed trajectories of tumor size (relative to day 0) during the trial, as inferred using all available clinical measurements of patients' tumor size. Predicted tumor sizes at day 90 match the RECIST assessments of tumor response (top panels) whilst trajectories of tumor burden distinguish sensitive (shrinking) and resistant (persistent) tumor through to the end of the trial (bottom panels). **c,** Inferred change in tumor size between the start-midpoint (left panel) or start-end (right panel) of the trial, in patient response groups classified by either RECIST assessment at trial midpoint (top row) or the biological response classification from tumor trajectories (bottom row). RECIST assessments distinguish response/non-response at day 90 but not day 180, whilst the biological response classification does distinguish resistance or sensitivity at day 180.

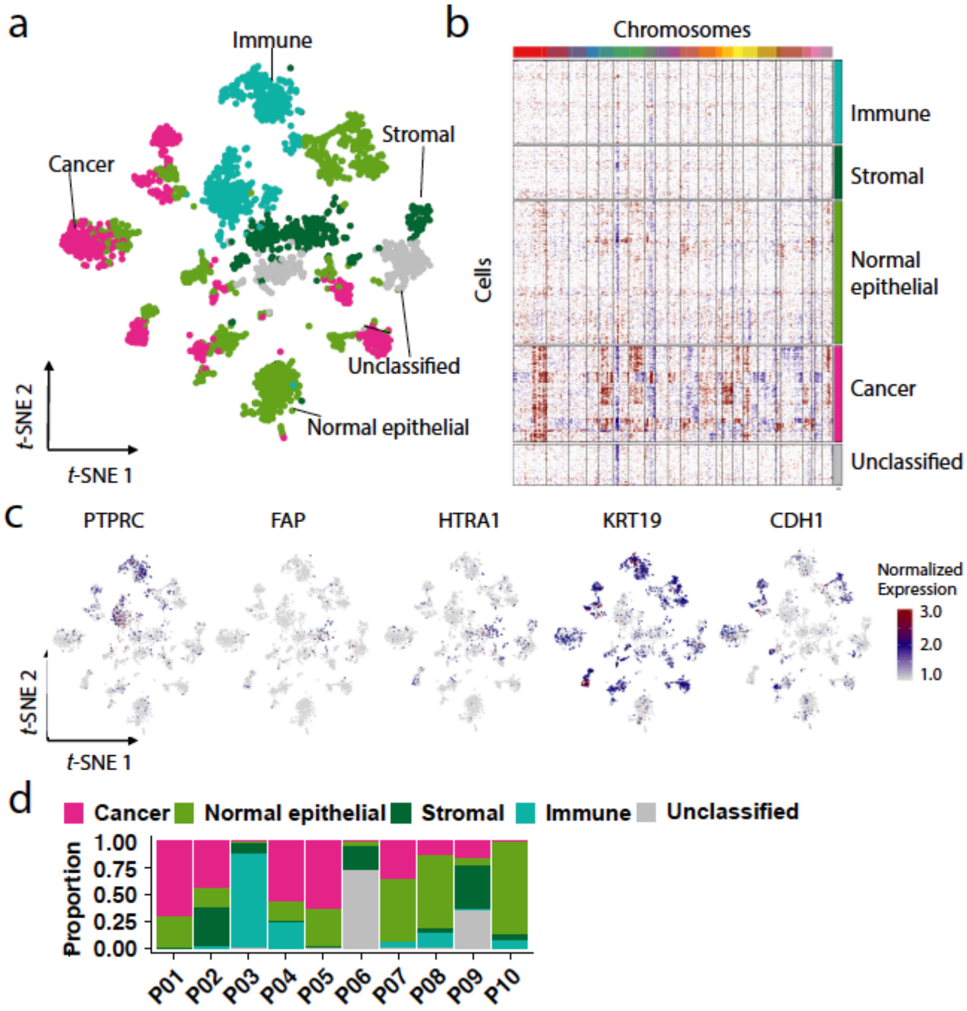


**Supplementary Figure 2. Landscape of tumor and macroenvironment of 10 patients with single nucleus isolated by ICELL8 platform**. **a,** *t*-SNE plot of 3,484 cells. Cells were classified into cancer cells, normal epithelial cells, immune cells, stroma cells, and unclassified cells, which are indicated by colors and labels. **b,** Gene copy number profile in cancer cells and neighboring normal cells. Blue color indicates copy number loss and red color indicates copy number gain. **c,** Expression of marker genes of cancer cells and normal epithelial cells (KRT19, CDH1), stromal cells (FAP, HTRA1), and immune cells (PTPRC). **d,** Proportion of cancer cells and neighboring normal cells in each patient.

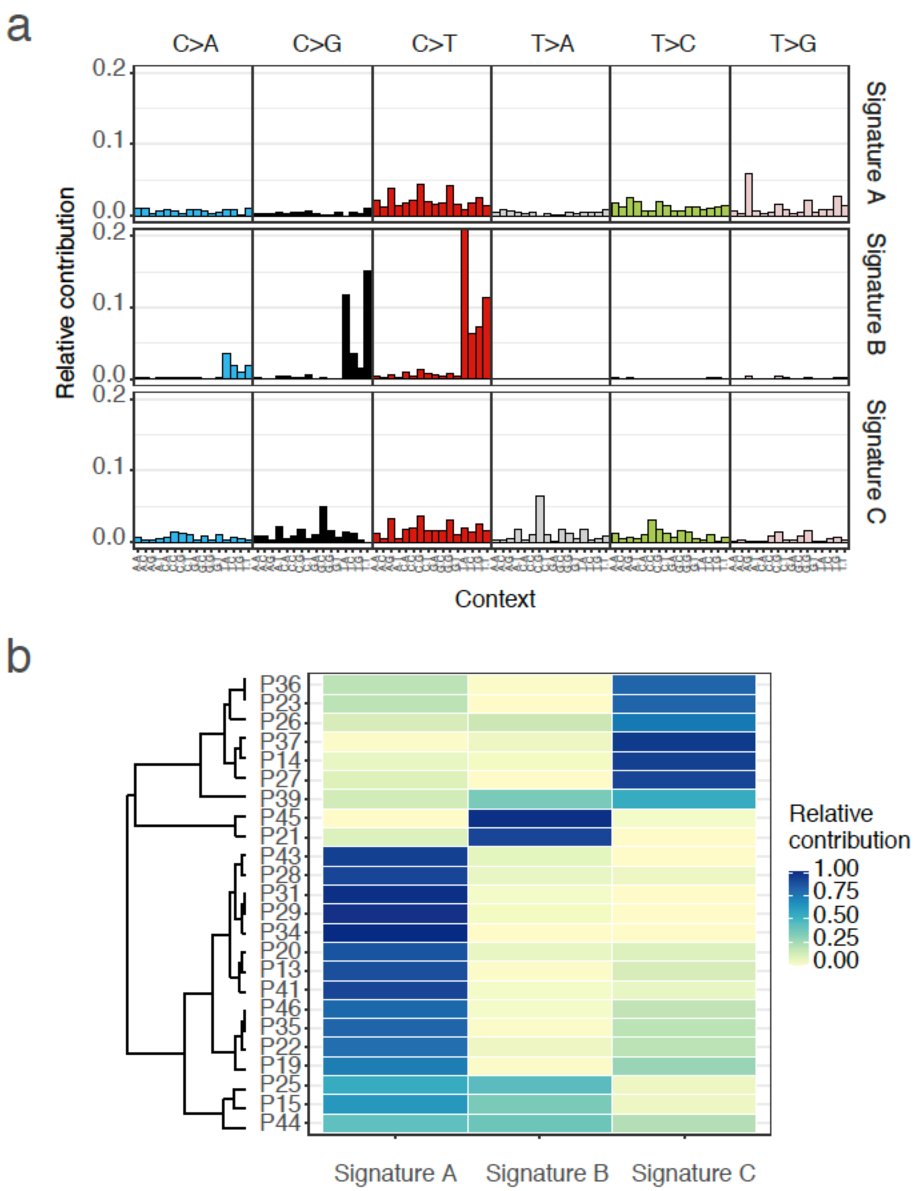


**Supplementary Figure 3. Mutational signature in 24 patients with whole-exome sequencing data. a,** Relative contribution of trinucleotide changes to three *de novo* mutational signatures identified in 24 patients. **b,** Relative contribution of each mutational signature to mutations in each patient.

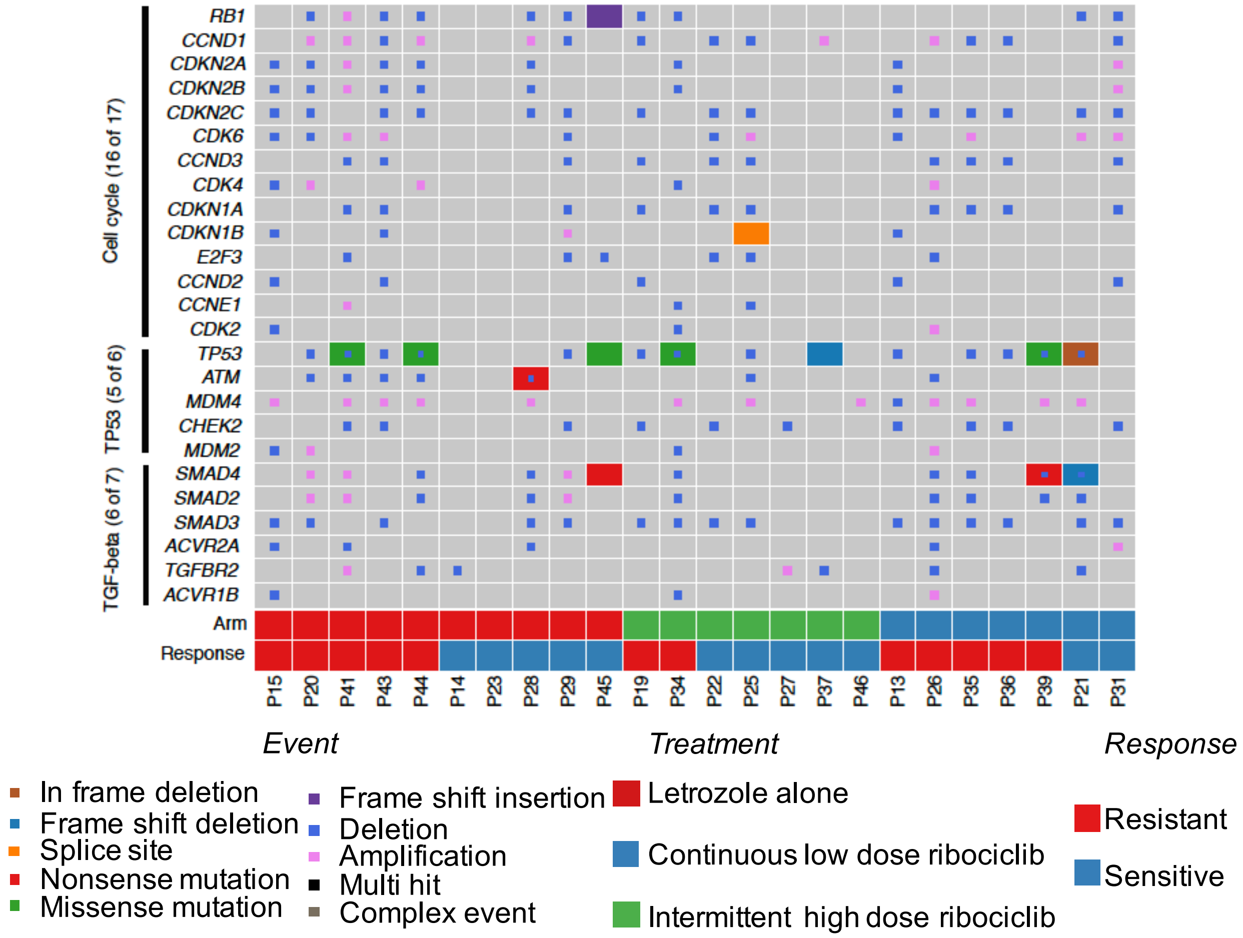


**Supplementary Figure 4. Mutated genes in three frequently altered oncogenic pathways.** Genes are grouped by oncogenic pathways. Presence of gene mutations in each patient is colored as indicated in the legend. Treatment arm and clinical response (Response: sensitive, resistant) are indicated in final two rows of the plot (colors indicated in legend).

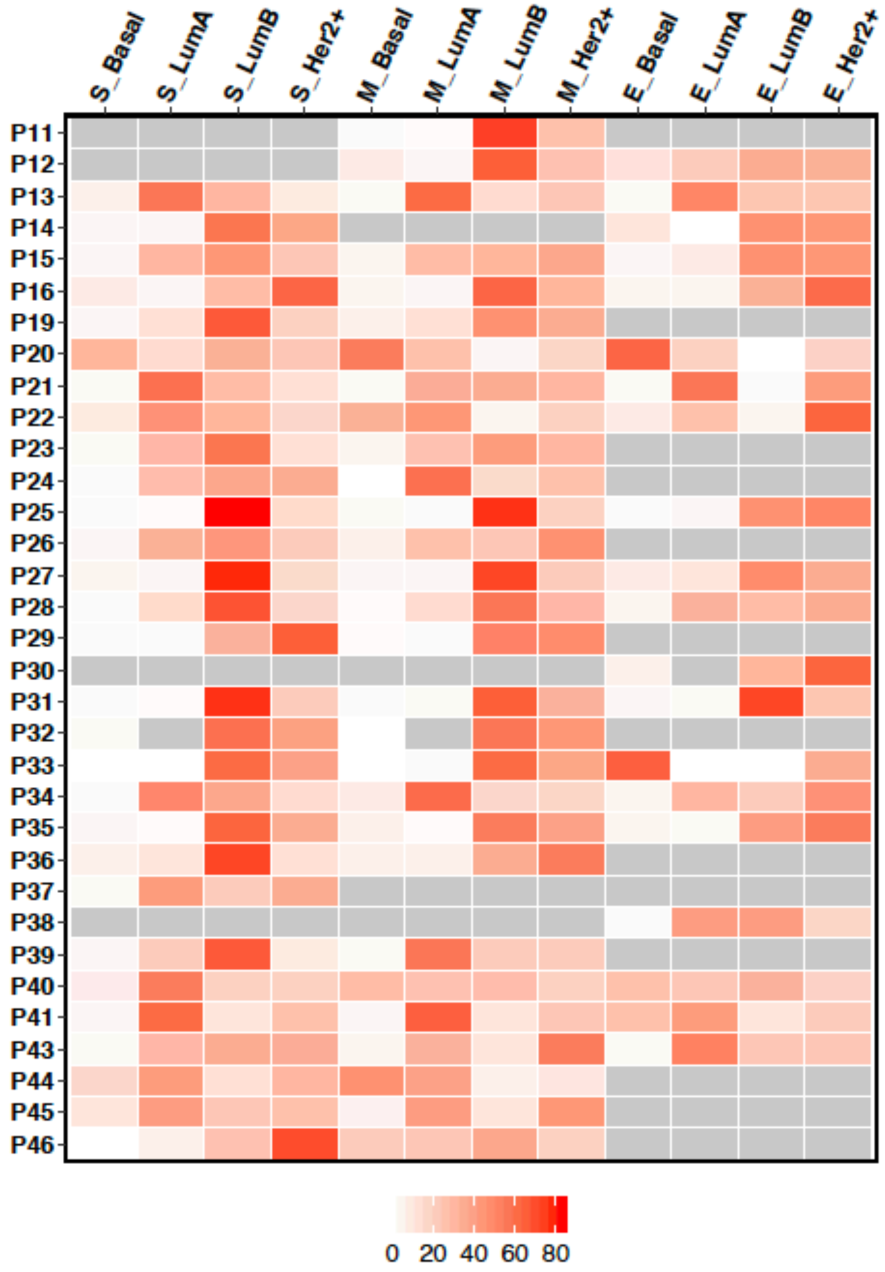


**Supplementary Figure 5. Intrinsic subtype of 35 patients with single nucleus isolated by 10x genomics platform.** Each row represents a patient and each column represents an intrinsic subtype at three timepoints. The proportion of cancer cells in each intrinsic subtype was indicated by colors ranging from 0 to 85. Patient samples without cancer cells were indicated by gray.

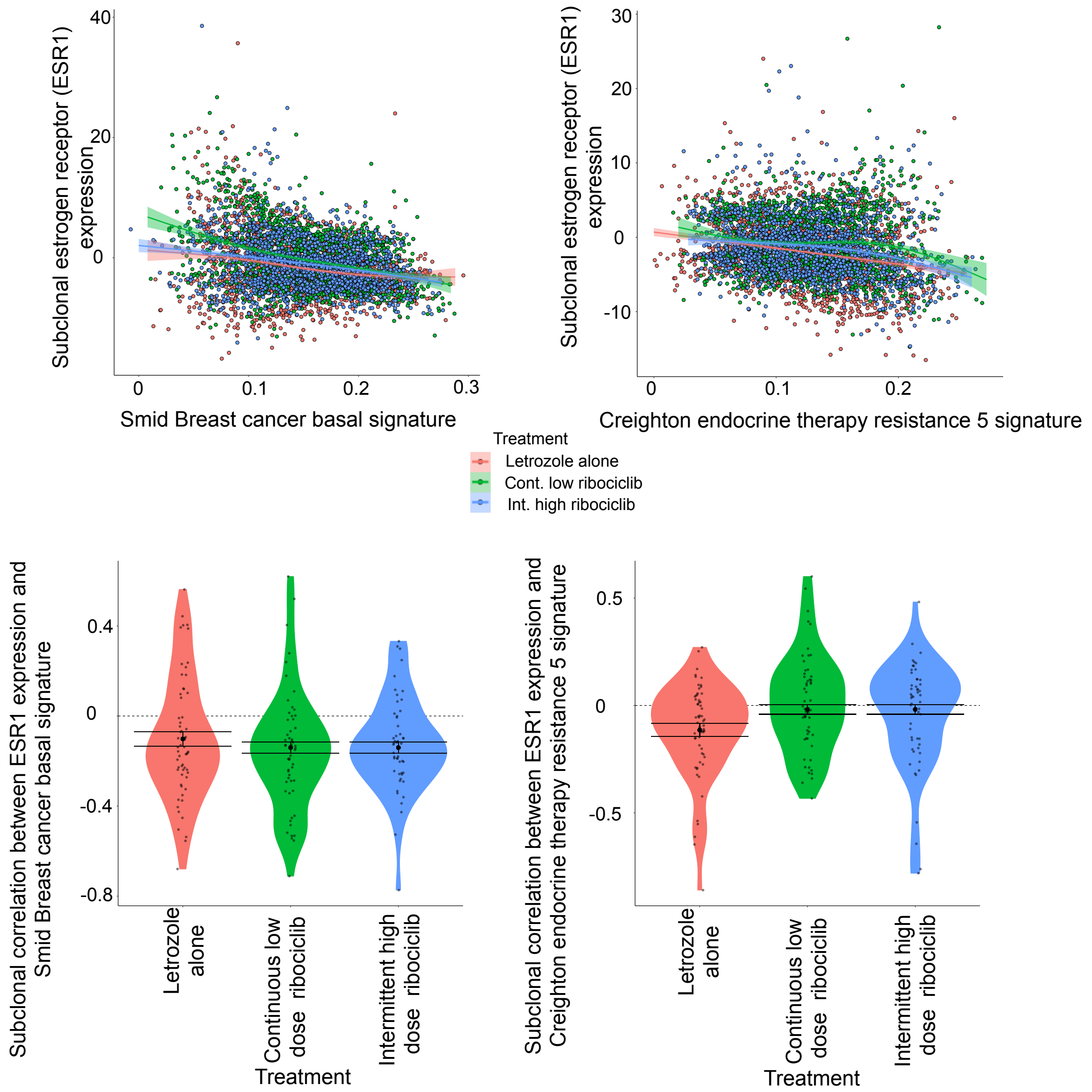


**Supplementary Figure 6. Reduced subclonal estrogen receptor (ESR1) expression at end of therapy correlates with increased basal-like pathway and Creighton endocrine therapy resistance signatures, independent of treatment.** Top row shows the ESR1 expression and basal-like (left) and endocrine resistance (right) pathway signatures across subclonal cancer populations with differing MAPK activation (points) and the coloration signifies the treatment received. Fitted lines show the overall trend between ESR1 expression and pathway activity (shaded regions show 95% confidence bands). Bottom row shows the correlation between ESR1 expression and basal-like (left) and endocrine resistance (right) pathway signatures for each cancer subclone present at end of trial, in patients treated with different therapies (colors). Black points and error bars signifies the mean and confidence interval for the correlation between ESR1 and pathway activity under each treatment.

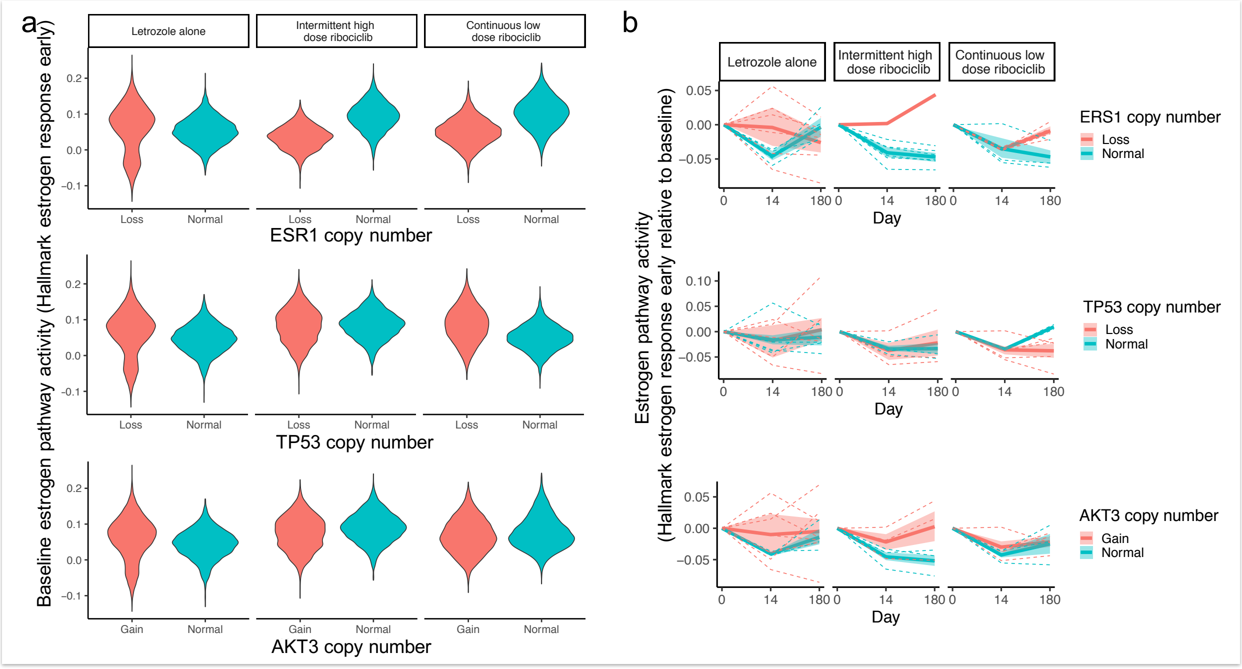


**Supplementary Figure 7. Minor impact of copy number loss in ESR1 and p53 or gain in AKT3 (rows) on estrogen pathway activity during therapy (columns). a,** Distribution of single cell estrogen pathway activity (Hallmark estrogen response early) at baseline, in patients with altered (red) or normal (blue) copy number status for the genes ESR1 (top), TP53 (middle) and AKT3 (bottom). b, Trajectories of change in estrogen pathway activity during treatment in patients with altered (red) or normal (blue) copy number genetic status for ESR1, TP53 and AKT3. Pathway trends determined across patients using hierarchical regression (solid lines). Inter-patient variability in pathway activity shown by dashed lines indicating patient specific responses and shaded regions showing confidence intervals of model estimates.

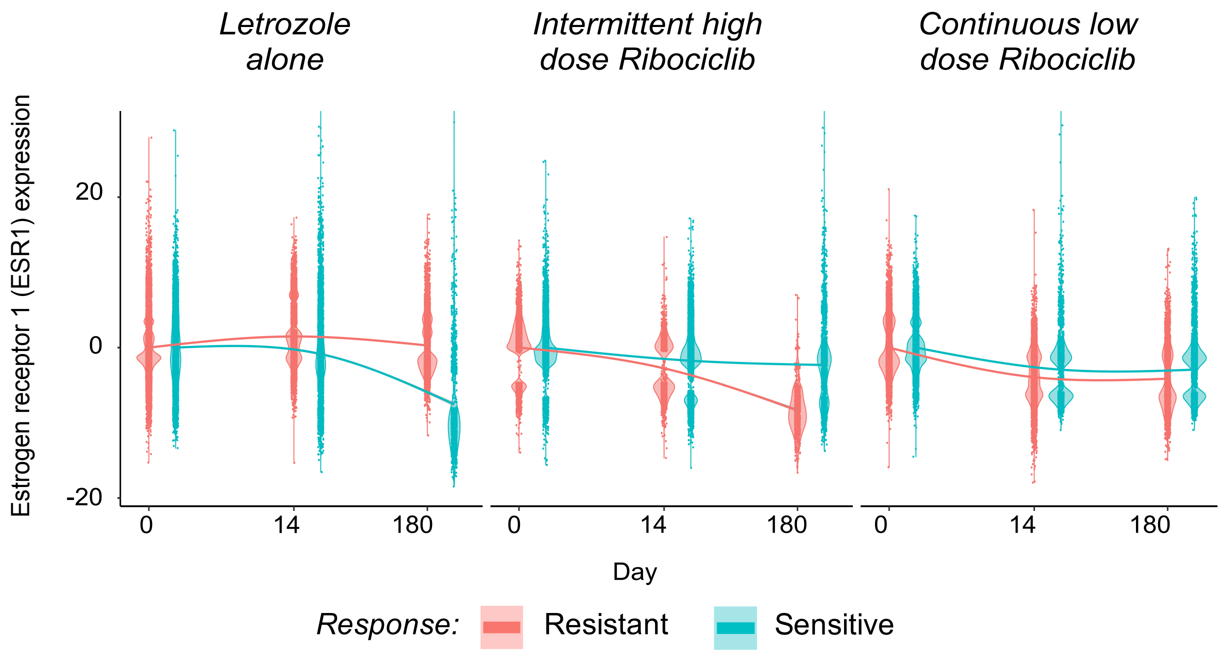


**Supplementary Figure 8. Reduction of estrogen receptor (ESR1) expression during different treatments (columns) for sensitive and resistant tumors (color).** Violin curves show, for each timepoint and response category, the distribution of ESR1 expression, with expression standardized relative to each patients average expression at baseline. Fitted lines show the temoral trend in expression as modelled using a random effects generalized additive model accouting for initial differences in patients' ESR1 expression.

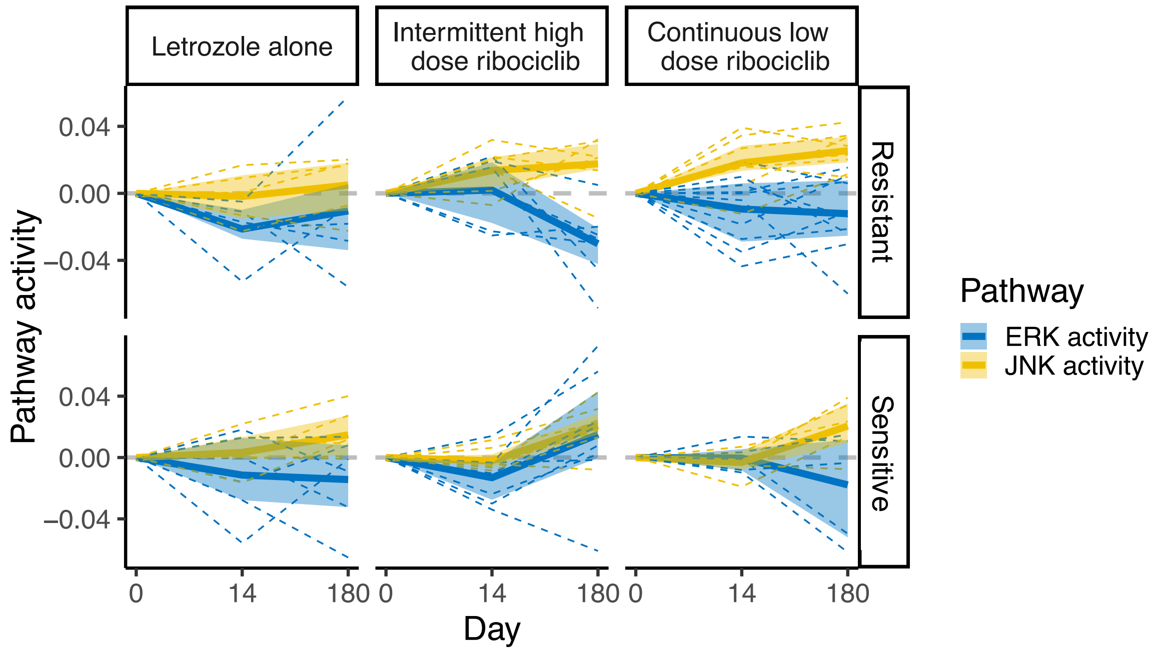


**Supplementary Figure 9. Divergence of JNK and ERK signalling pathway activity during treatment with combination therapy, especially in resistant tumors.**
JNK and ERK expression (color=pathway) during treatment (columns) in sensitive and resistant tumors (rows). Pathway trends determined across patients using hierarchical regression (solid lines). Inter-patient variability in pathway activity shown by dashed lines indicating patient specific responses and shaded regions showing confidence intervals of model estimates (JNK ssGSEA pathway=St JNK MAPK and ERK pathway=Biocarta ERK).

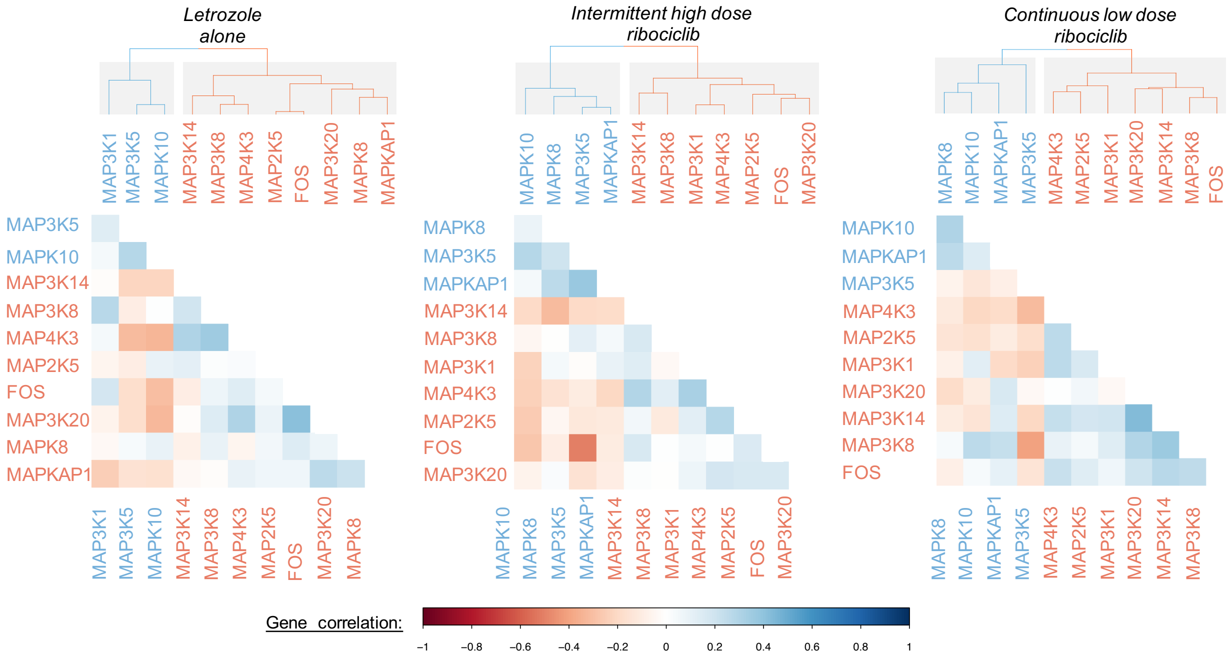


**Supplementary Figure 10. Heatmaps of the correlation between MAPK gene expression in each treatment arm (columns), showing the dichotomy between JNK and ERK activating genes across treatments**. Dendrograms show the collinearity of MAPK gene expression following each endocrine or combination therapies (columns).

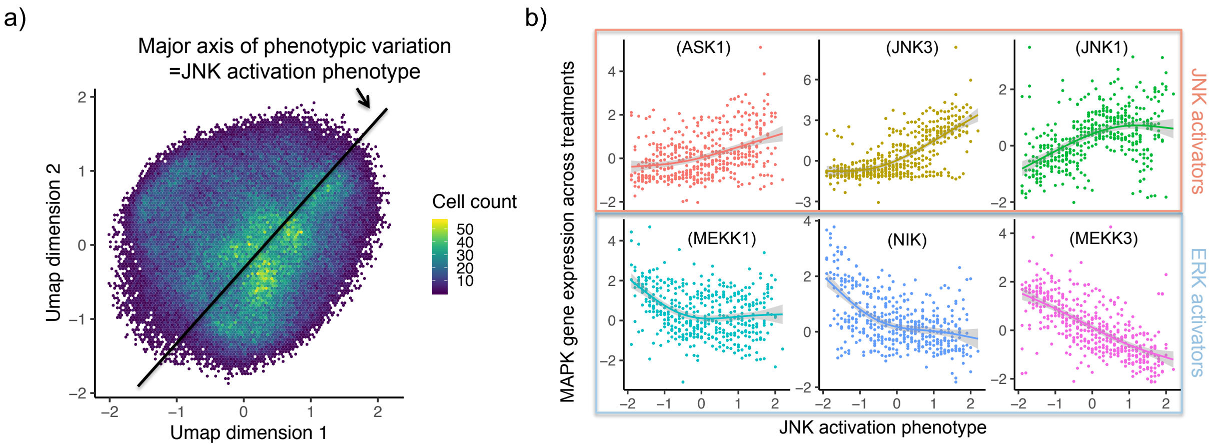


**Supplementary Figure 11. Construction of the overall JNK activation phenotype score, utilizing this collinearity of gene expression between ERK and JNK genes.** **a,** UMAP dimension reduction of MAPK genes, showing the bivariate Gaussian distribution of UMAP values, centered around the major axis of phenotypic variation (black line). The frequency of cells found in different parts of the UMAP phenotype space is shown by the color gradient. The major axis of phenotypic variation, (the JNK activation phenotype) is identified as the first principle component in the UMAP phenotype space. **b,** Relationship between the JNK activation phenotype and expression of MAPK genes that are known a JNK activators (red) or ERK activators (blue) across subclonal cancer populations. Loess smooths are added showing the positive relationship between the JNK phenotype score and key JNK activators and the negative association between ERK activators and the JNK phenotype.

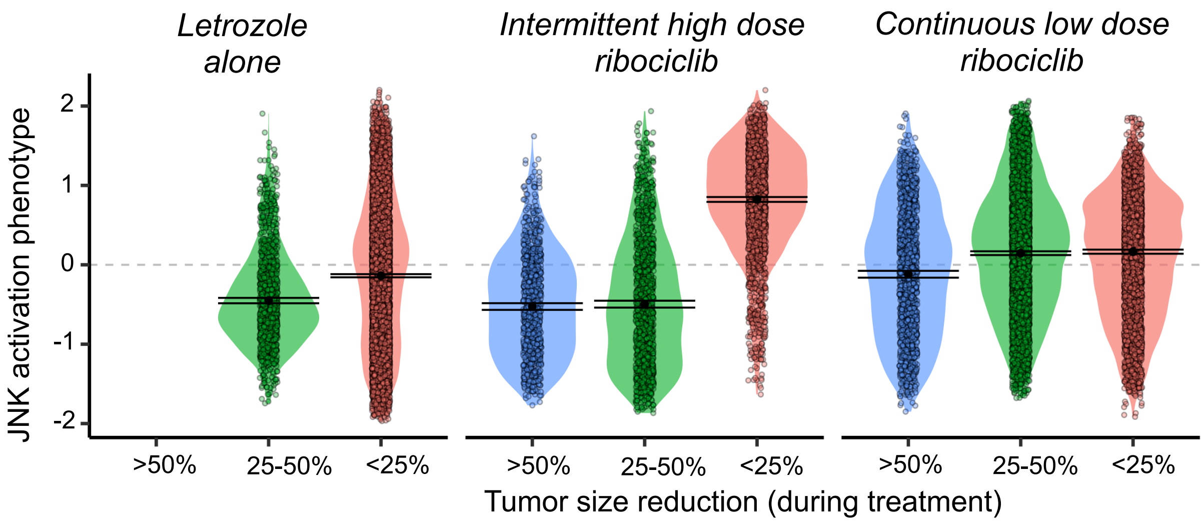


**Supplementary Figure 12. Distribution of single cell JNK activation scores at end of treatment, in patients with differing levels of reduction in tumor size (color) during different therapies (column).** Violin curves show the distribution of single cell JNK activation scores within each response category and the mean and confidence intervals bars are shown by the black dots and error bars.

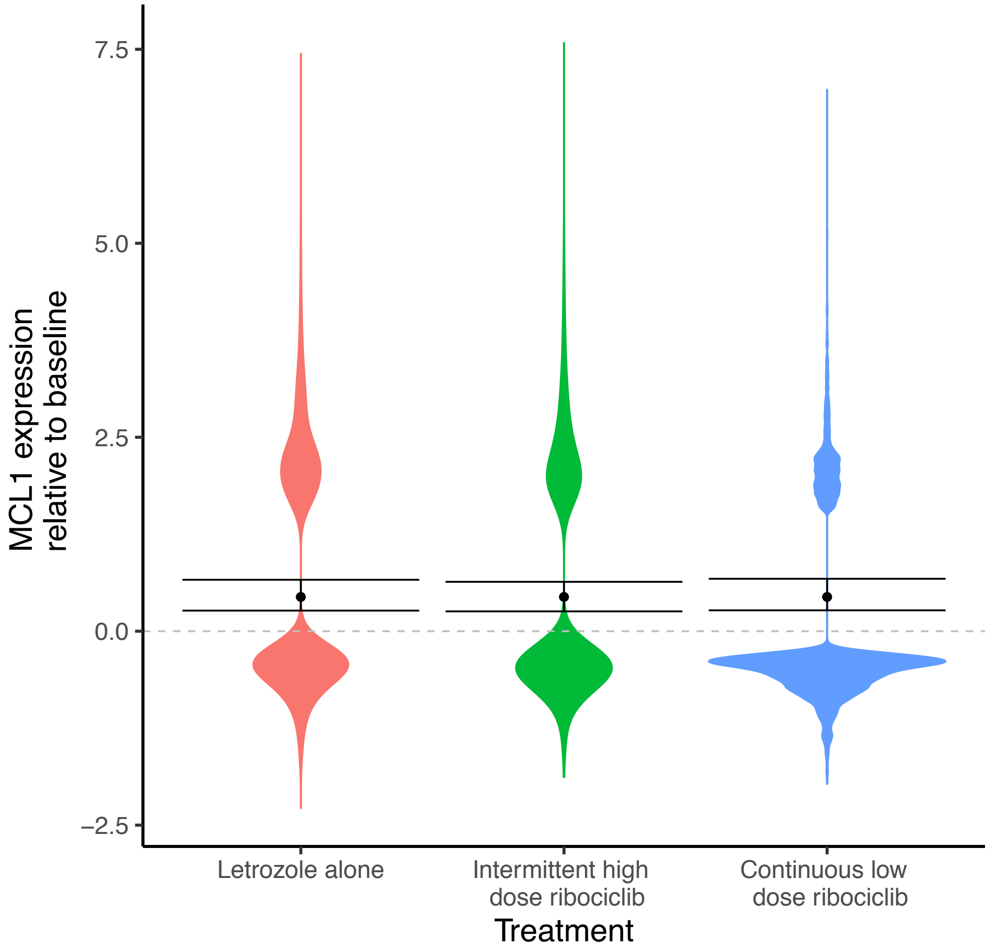


**Supplementary Figure 13. Upregulation of the BCL2 family member and apoptosis regulator, Myeloid Cell Leukemia Sequence 1 (MCL1) across treatment arms.** The distribution of single cell MCL1 expression at end of treatment, relative to the average expression at baseline, is shown by violin curves for patients given endocrine or combination therapy. Horizontal dashed grey line shows average expression at baseline. The increase in MCL1 expression across patient tumors following each treatment was determined using hierarchical regression (black point). Model indicated by error bars showing the 95% confidence interval of model estimates.

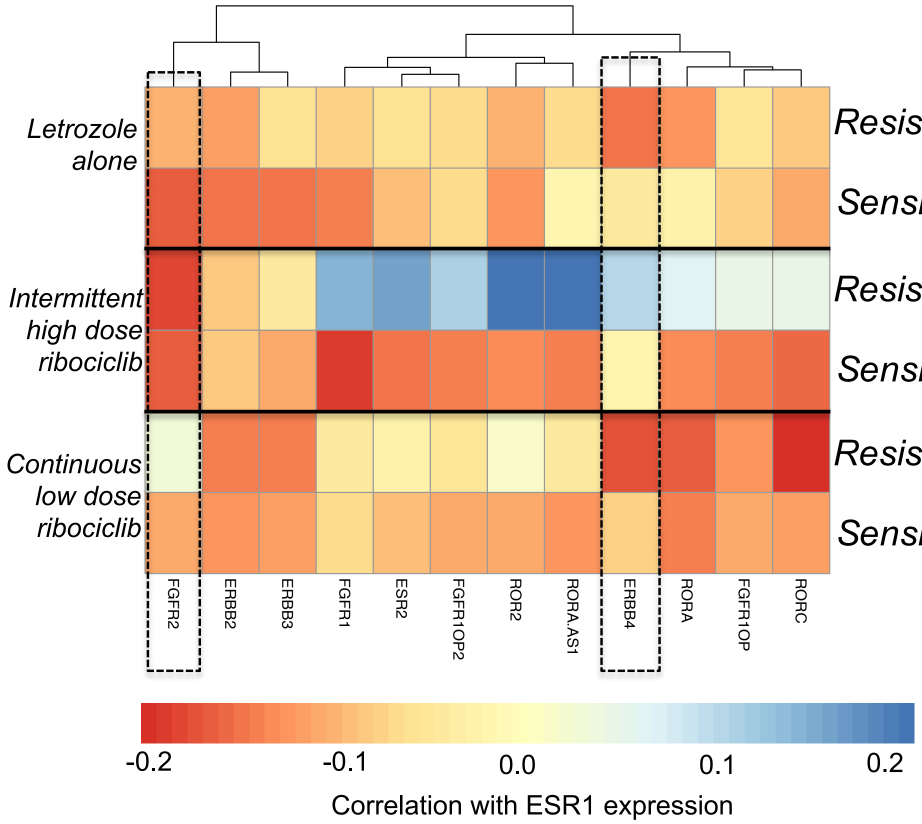


**Supplementary Figure 14. Correlation of growth factor receptors expression with estrogen pathway activity (Hallmark estrogen response early) in cancer cells from sensitive and resistant tumors under each therapy.** Strong negative correlations identify genes that are upregulated as estrogen signaling is lost . Specifically, tumors resistant to intermittent high dose and continuous low dose show compensatory activation of FGFR2 and ERBB4 respectively.

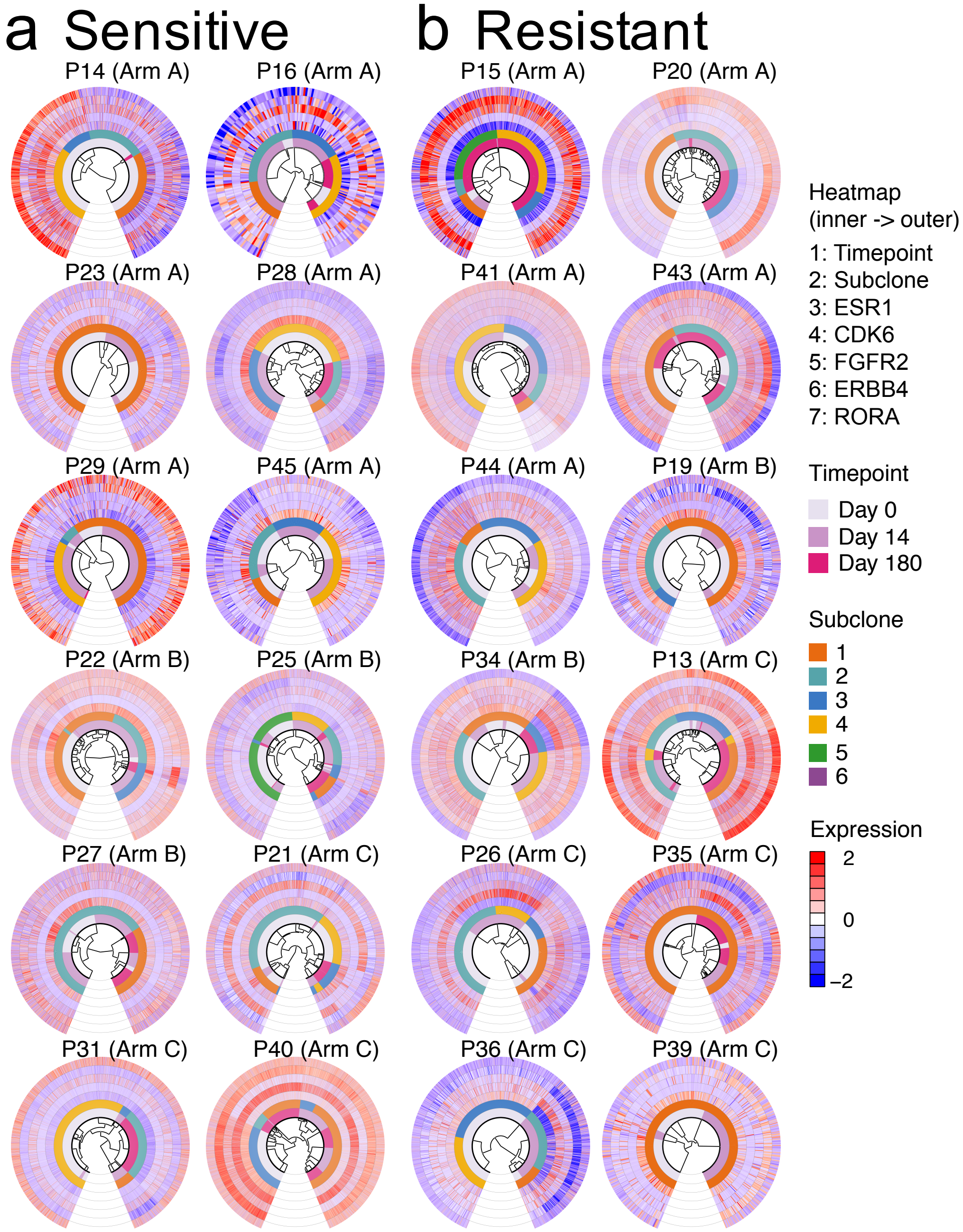


**Supplementary Figure 15. Transcriptional heterogeneity of key resistent genes. a,** sensitive and **b,** resistant tumors. For each patient's tumor cells, a single-cell phylogenetic tree is shown at the center of circos plot. Cell annotaiton (timepoint and subclone) as well as expression of key resistent genes (ESR1, CDK6, FGFR2, ERBB4, RORA) are shown as heatmap. Phylogenetic tree of cells were contructed based on the distance between cell gene copy number profile. Subclones were infered based on gene copy number profile. Zinbwave normalized gene expression were centered and scaled.

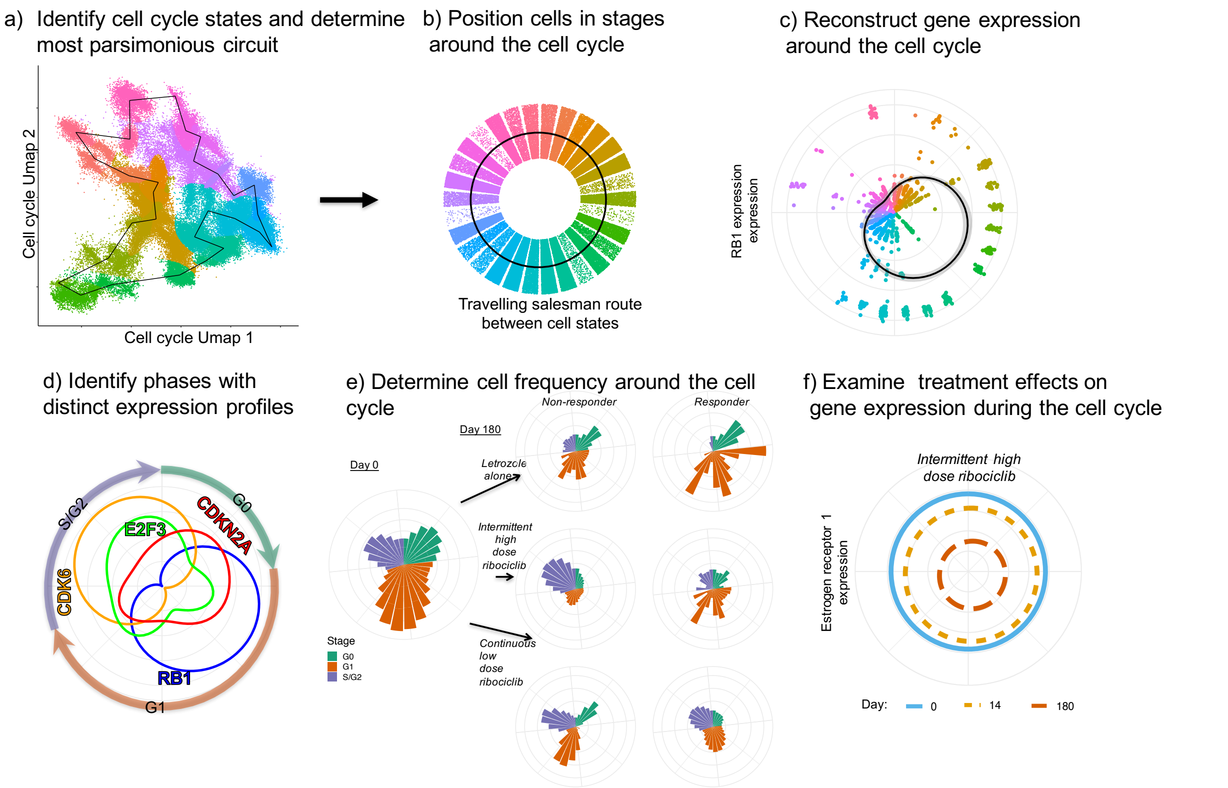


**Supplementary Figure 16. Reconstruction of cell cycle, fluctuations in gene expression during the cell cycle, distinct cell cycle phases, frequencies of cells throughout the cell cycle and shifts in gene expression within the cell cycle during therapy. a,** Single cell RNA seq gene expression profiles of cell cycle genes are extracted and used to perform dimension reduction with the UMAP algorithm. Cell cycle states (colors) with differing expression were identified using a gausian mixture model and the transitions between these states determined by the sortest distance to travel through each state and return to the original (Traveling salesman route=black line). **b,** Cells states ordered along the traveling salesman route. **c,** Example of fluctuations in gene expression of cells around the cell cycle (distance of points from origin=RB1 expression; colors=cell cycle state) Reconstruction of the fluctuation in average gene expression is predicted using a cyclical generalised additive model (black line with shaded confidence bands). **d,** Reconstructed fluctuations (coloured curves) in expression of genes around the cell cycle are used to classify distinct phases of the cell cylce (annotated by arrows around). Here we show four examples of key cell cycle genes which influence the classification of cell cycle phases (G0, G1, S/G2). **e,** The frequncy of cells in each stage of the cell cycle (height of bars) was counted and used to examine changes in the fraction of sampled cells in each phases cell cycle phase over time and between treatment and response groups. **f,** During treatment, the changes in gene expression fluctuations around the cell cycle were examined. Distance of the curve from the origin indicates gene expression and colored curves shows expression at different timepoints.

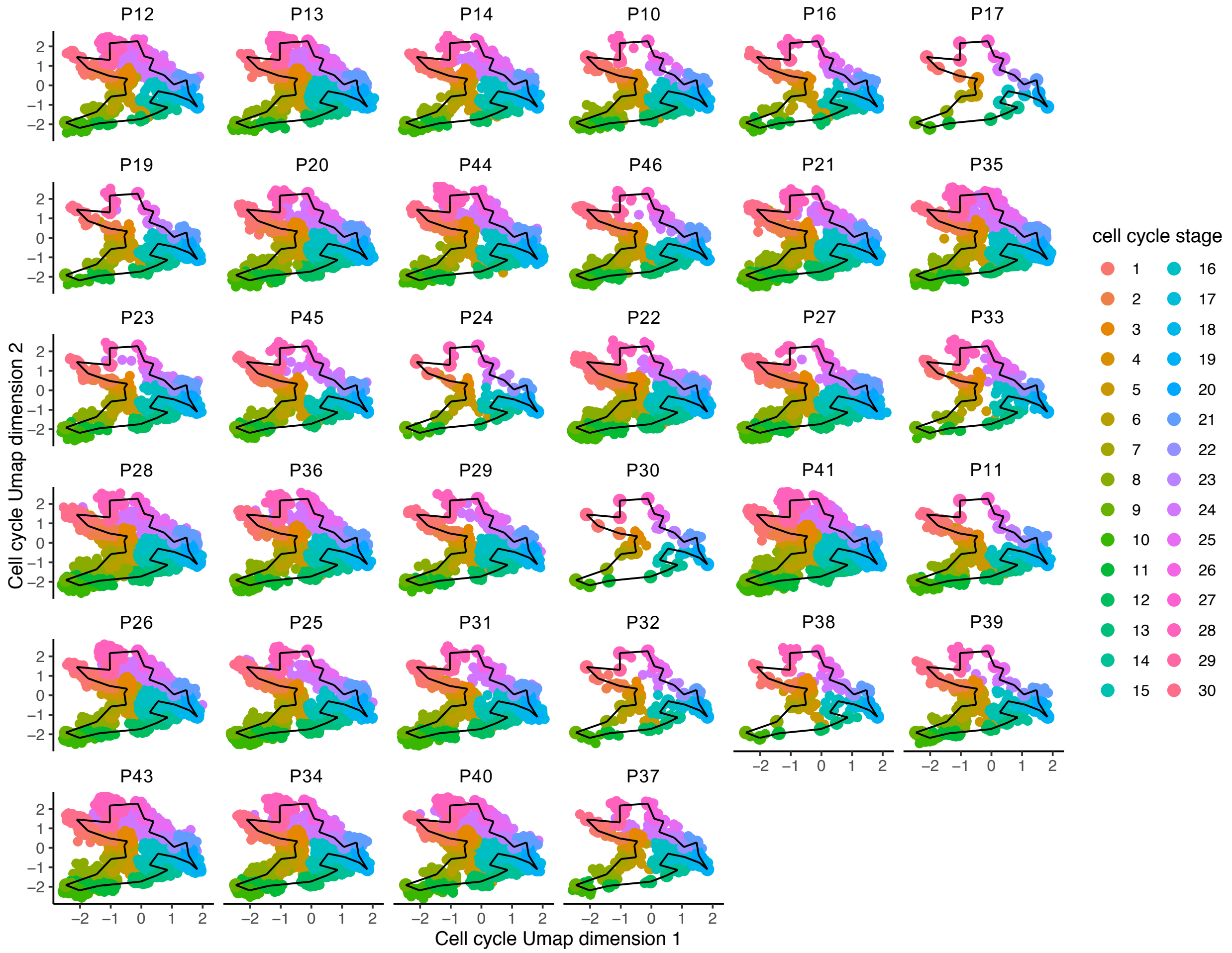


**Supplementary Figure 17. Consistent cell cycle stages present across patients.** For each patient (subpanel), single cell RNA seq gene expression profiles for cell cycle genes were extracted and the fitted UMAP model used to project cells onto the lower dimensional cell phenotype space (UMAP dimensions 1 and 2). Cell cycle stages (colors) with differing expression, identified using the Gaussian mixture model, were overlaid, showing that all patients have cells that are distributed across the cell cycle phenotype space. The traveling salesman route (black line) shows the transitions between these stages, as determined by the shortest distance to travel through each state and return to the original.
