## Supplementary Dataset 1 for "Convergent evolution of resistance pathways during early stage breast cancer treatment with combination cell cycle (CDK) and endocrine inhibitors"

**Supplementary Dataset 1. Copy number analysis of whole-exome sequencing (WES) using FACETS.** In each figure, FACETS analysis of a pre-treatment sample (Day 0) and a post-treatment sample (Day 14 or 180) were shown for a patient. The top panel shows total copy number log-ratio (logR), which is computed from read counts in tumor compared to read counts in normal samples. The middle panel shows allele-specific log-odds-ratio (logOR), which is an unbiased estimate of allele-specific copy ratio. The bottom panel shows the estimated copy numbers with black colors indicating total copy number and red colors indicating minor copy number.

P13 Day 0

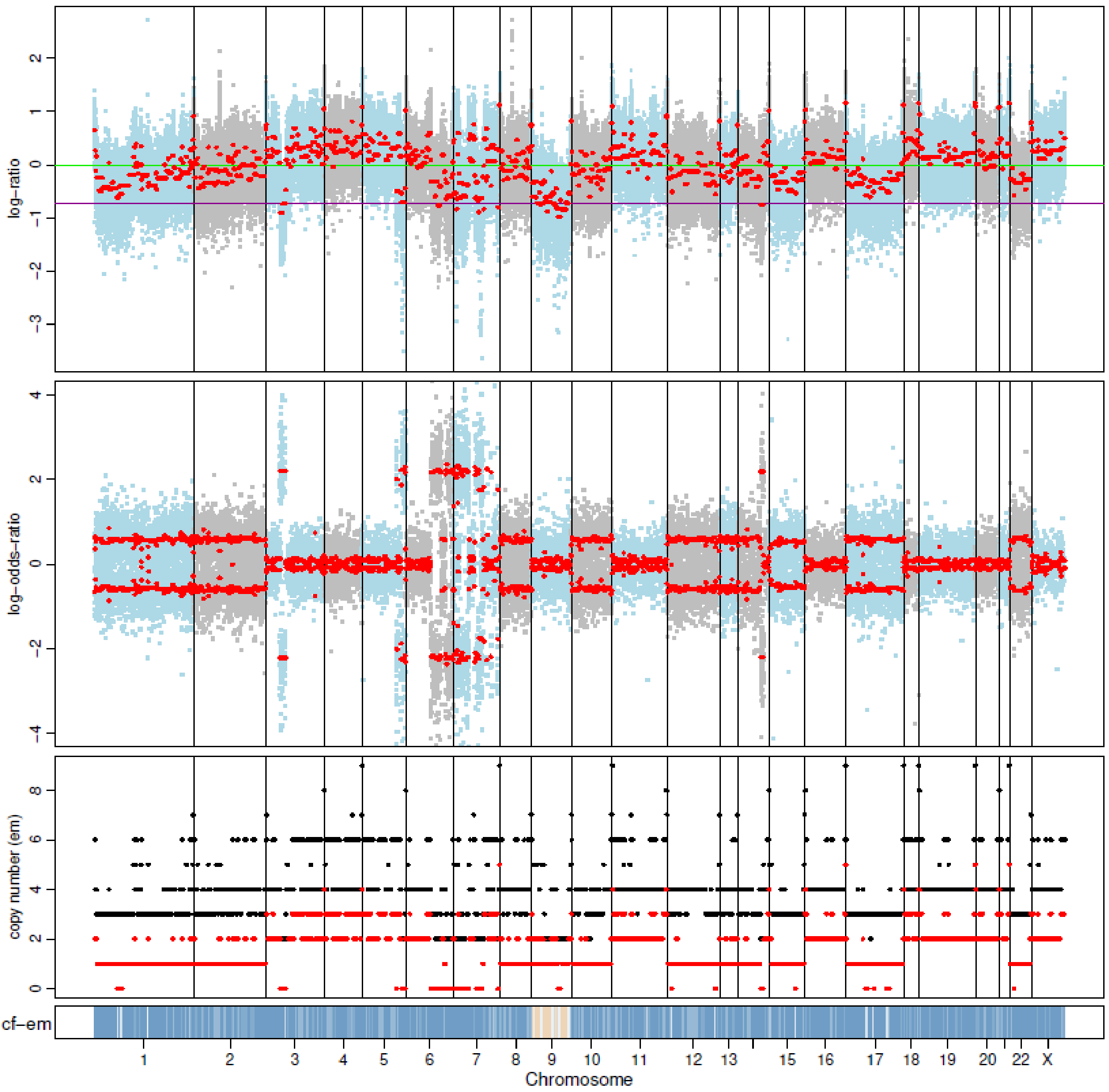


P13 Day 180

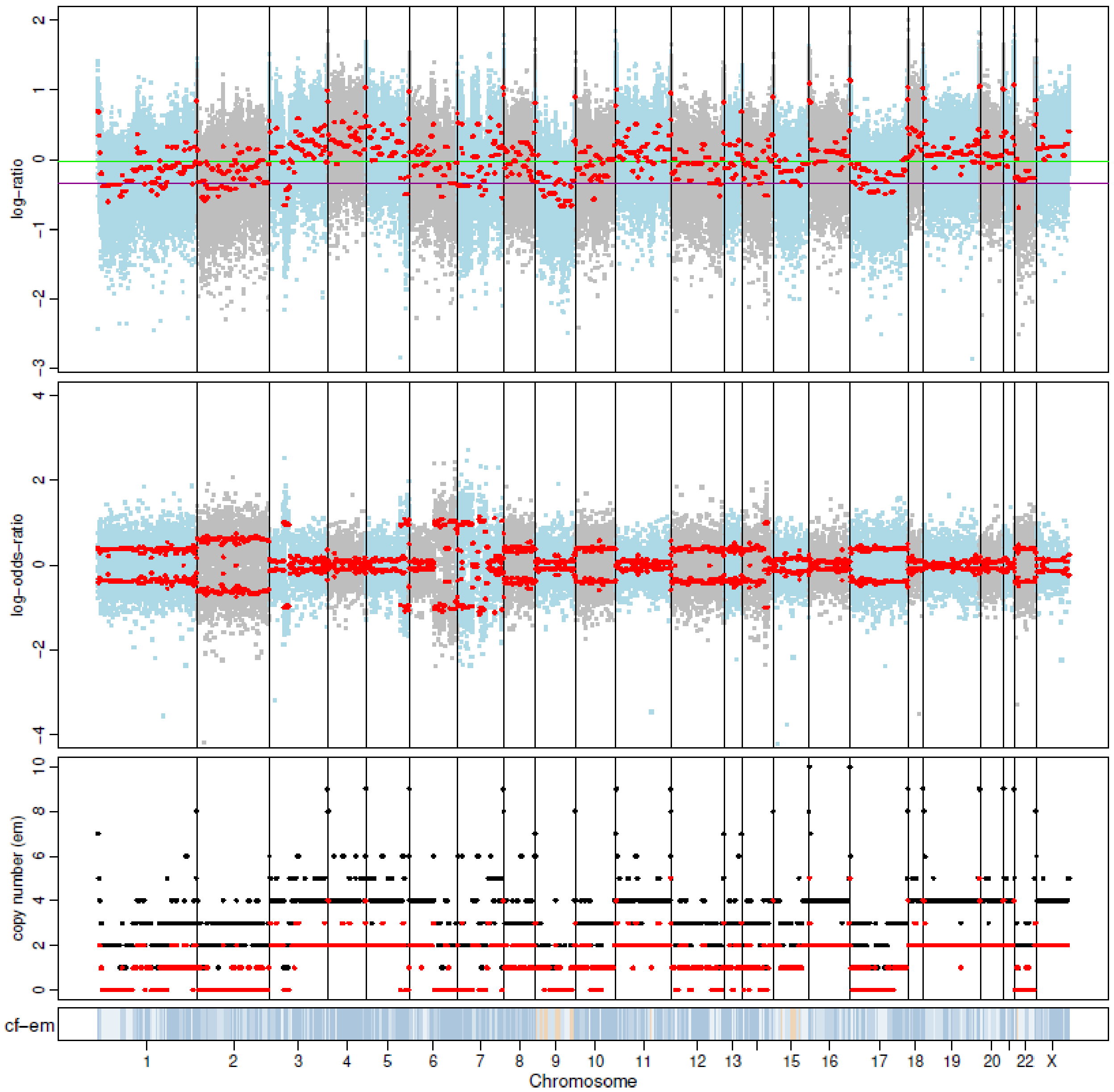

P14 Day 0

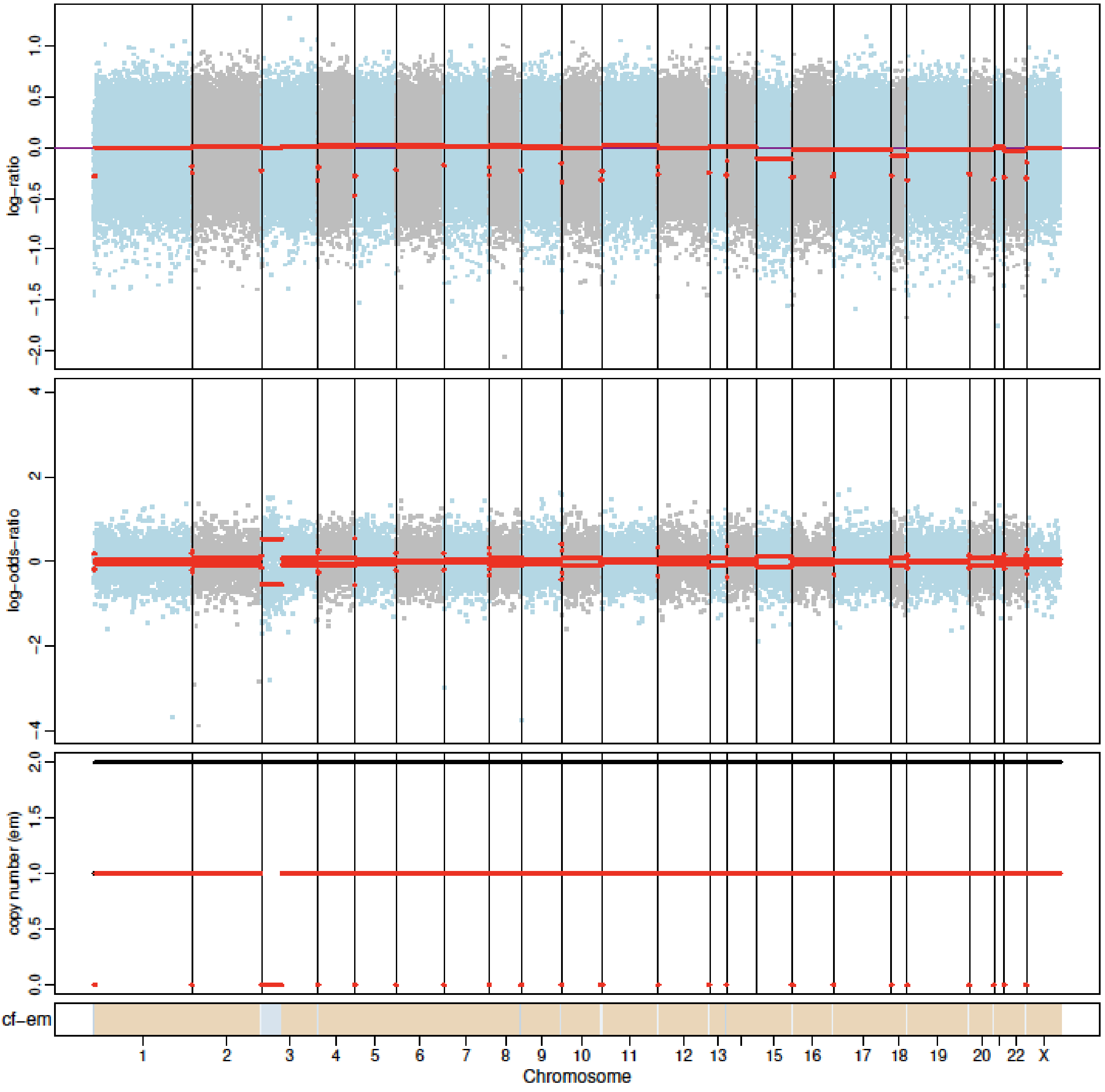


P14 Day 180

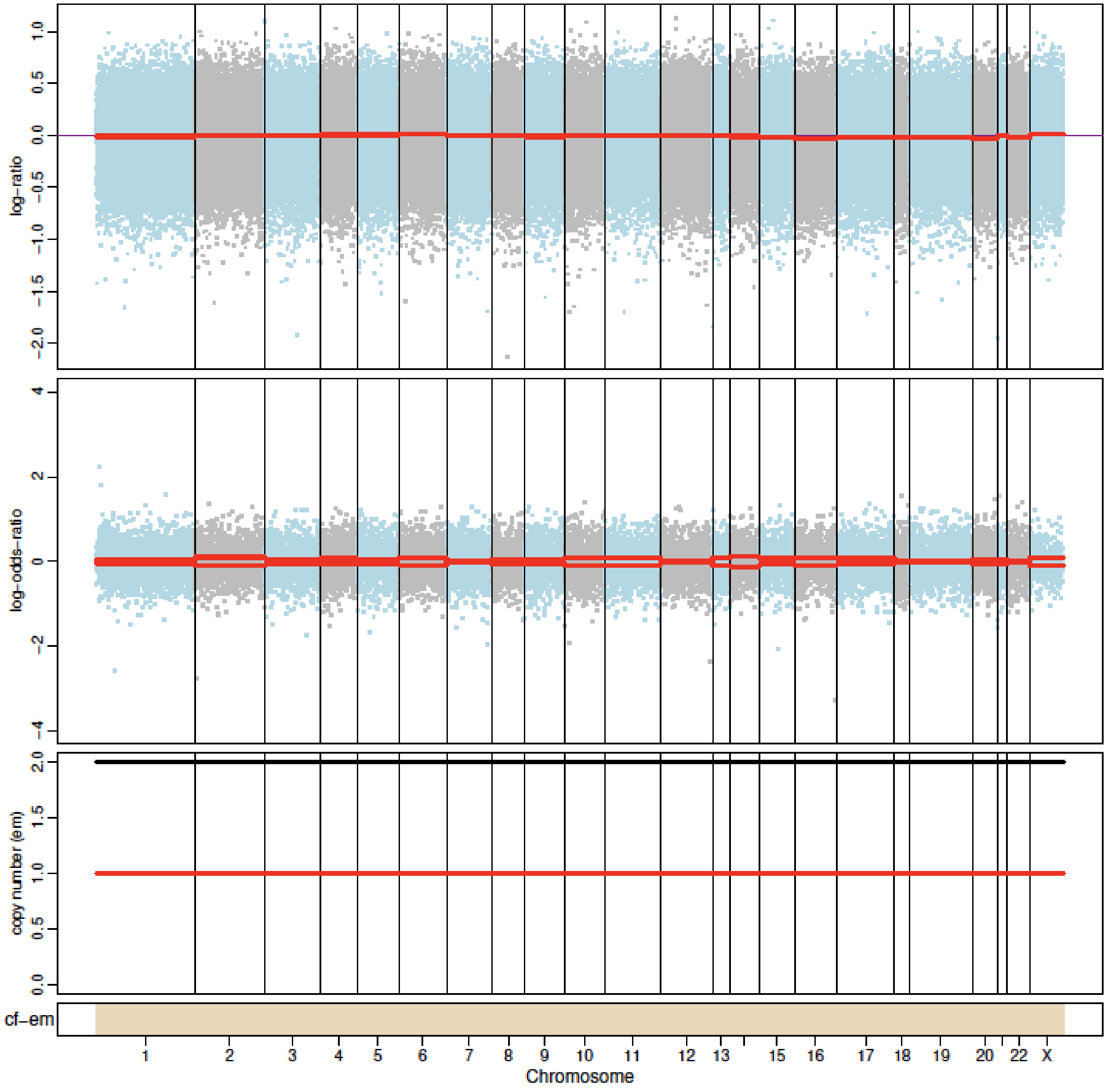

#### P15 Day 0

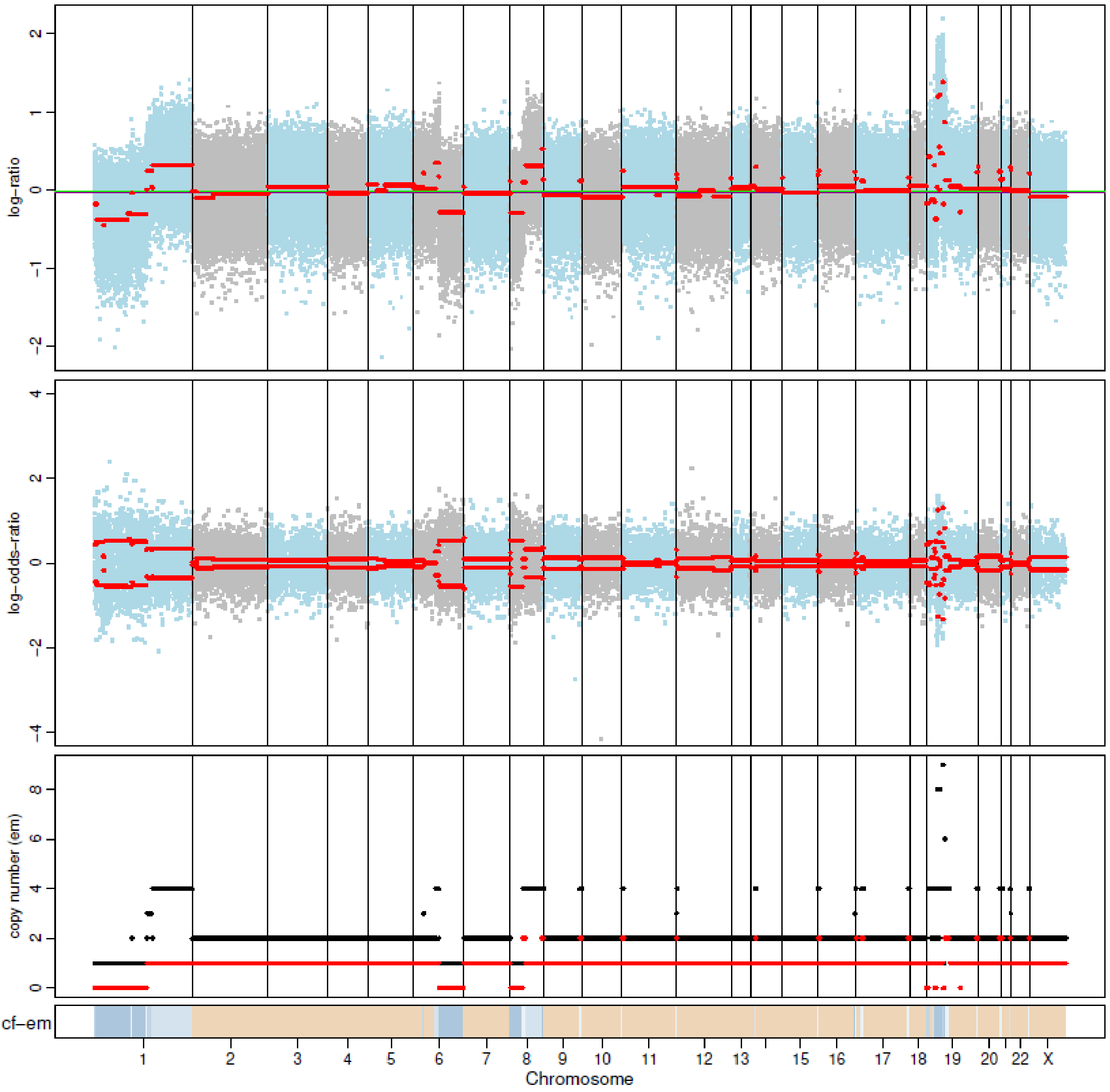


#### P15 Day 180

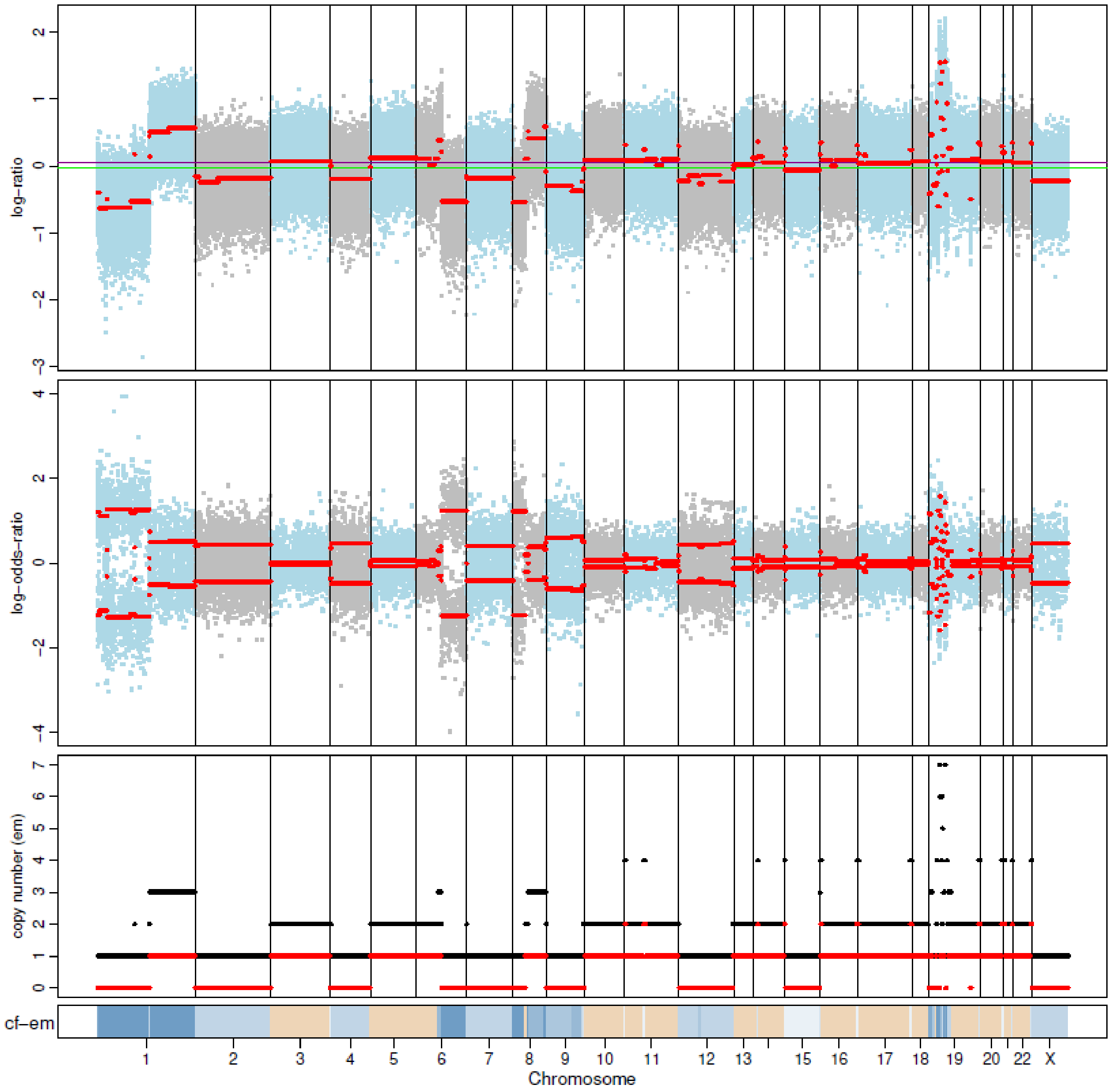

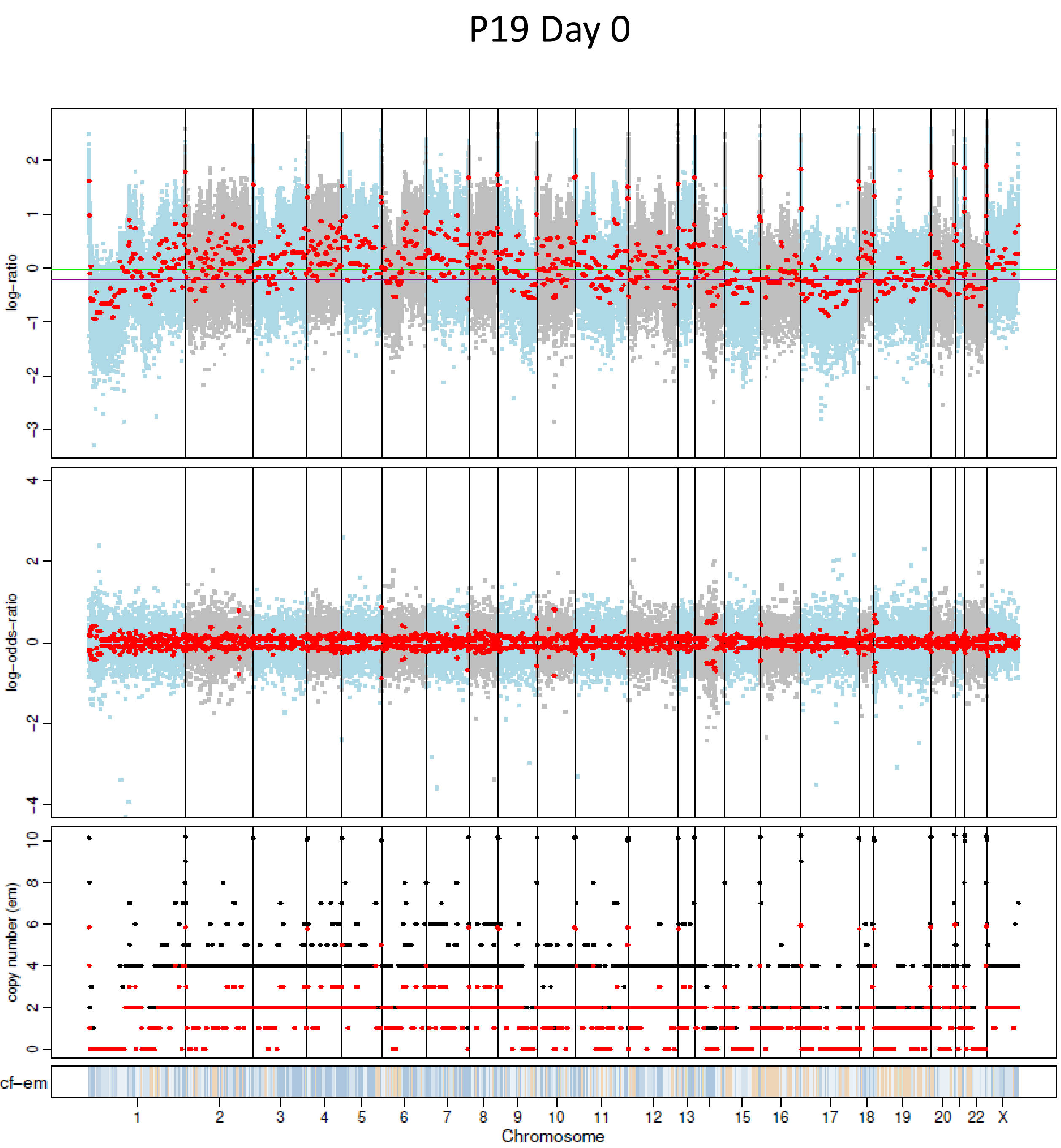
P19 Day 0
log-ratio
log-odds-ratio
copy number (em)
cf-em
Chromosome

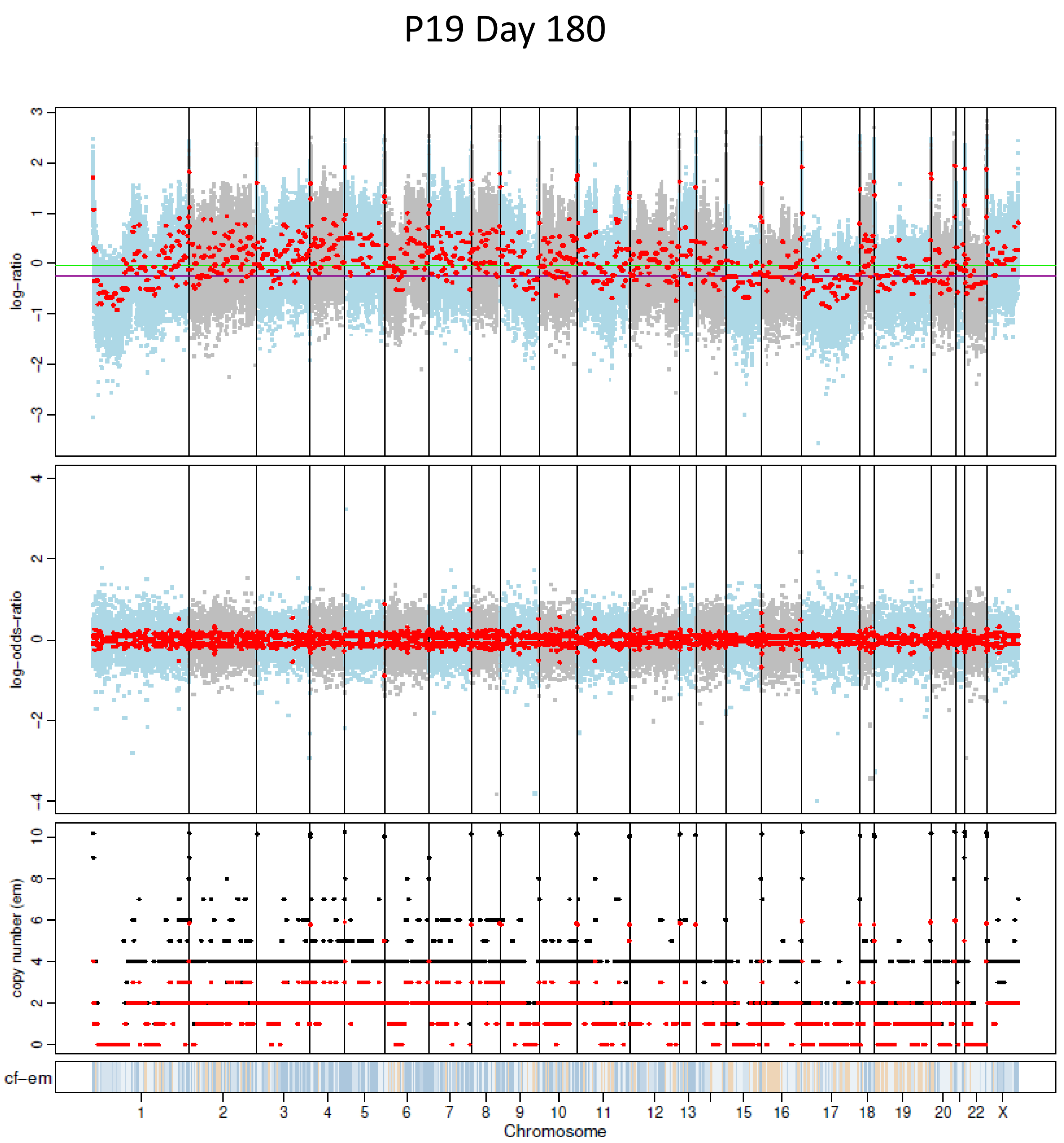
P19 Day 180
log-ratio
log-odds-ratio
copy number (em)
cf-em
Chromosome

#### P20 Day 0

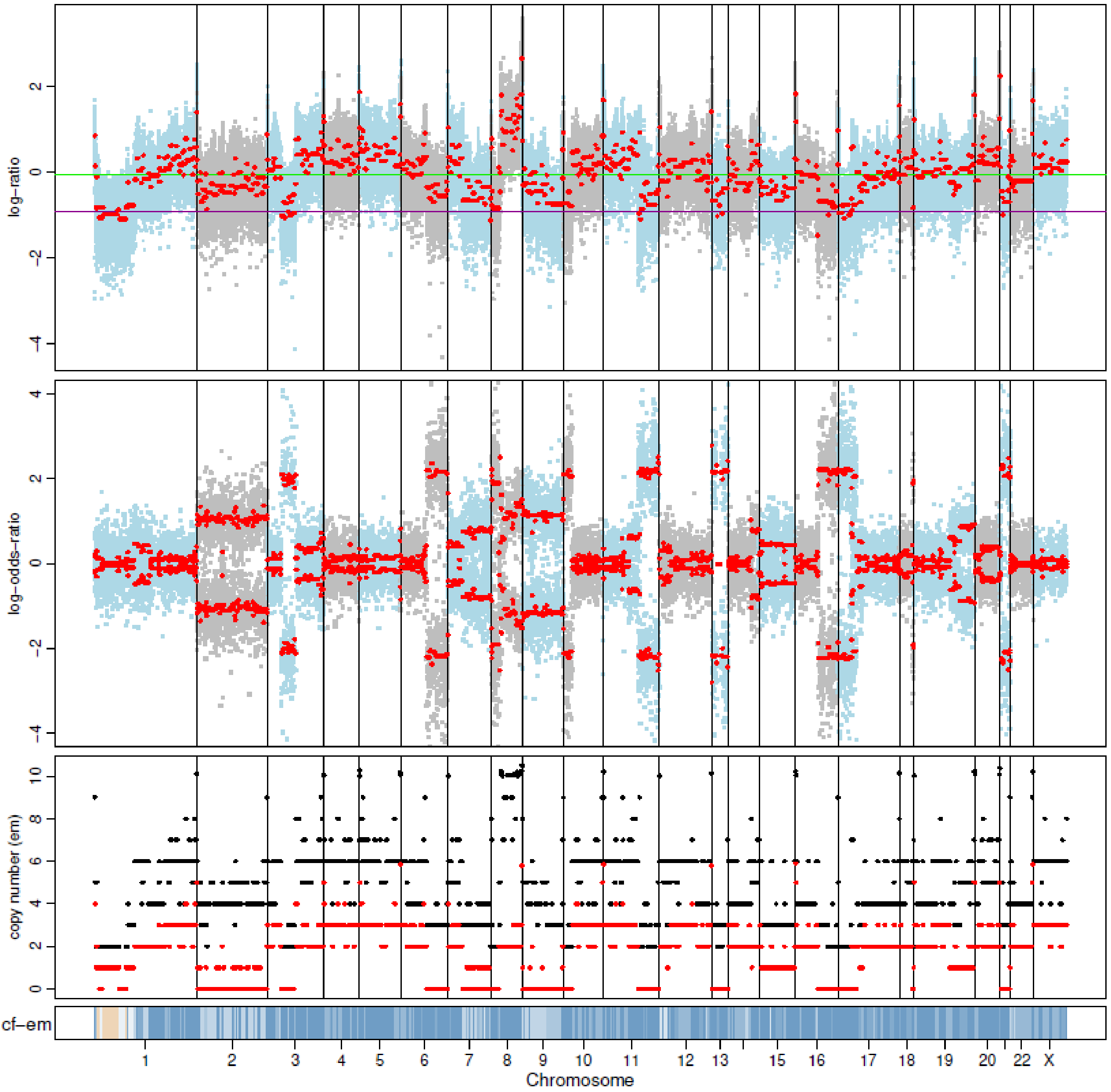


#### P20 Day 180

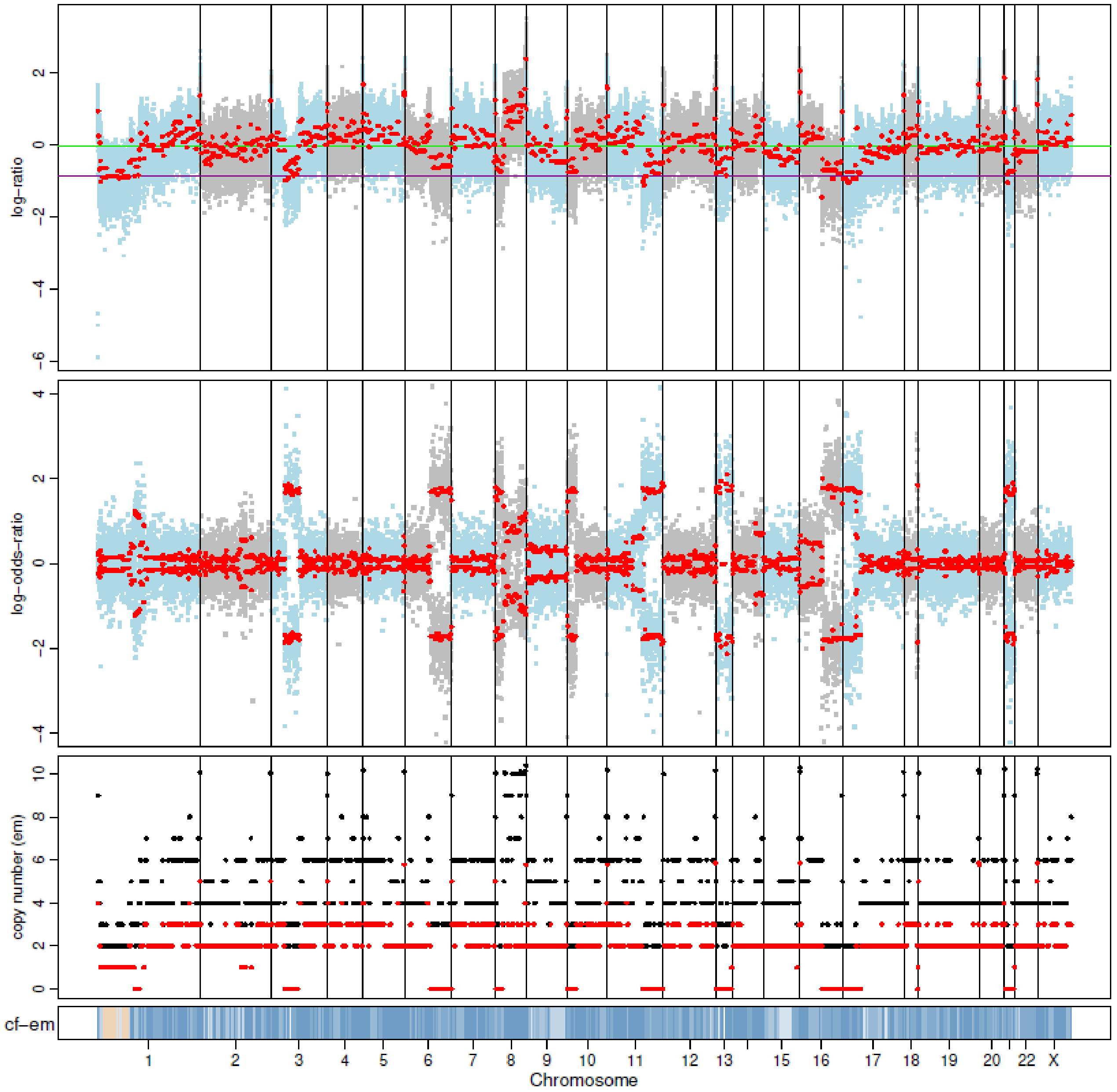

#### P21 Day 0

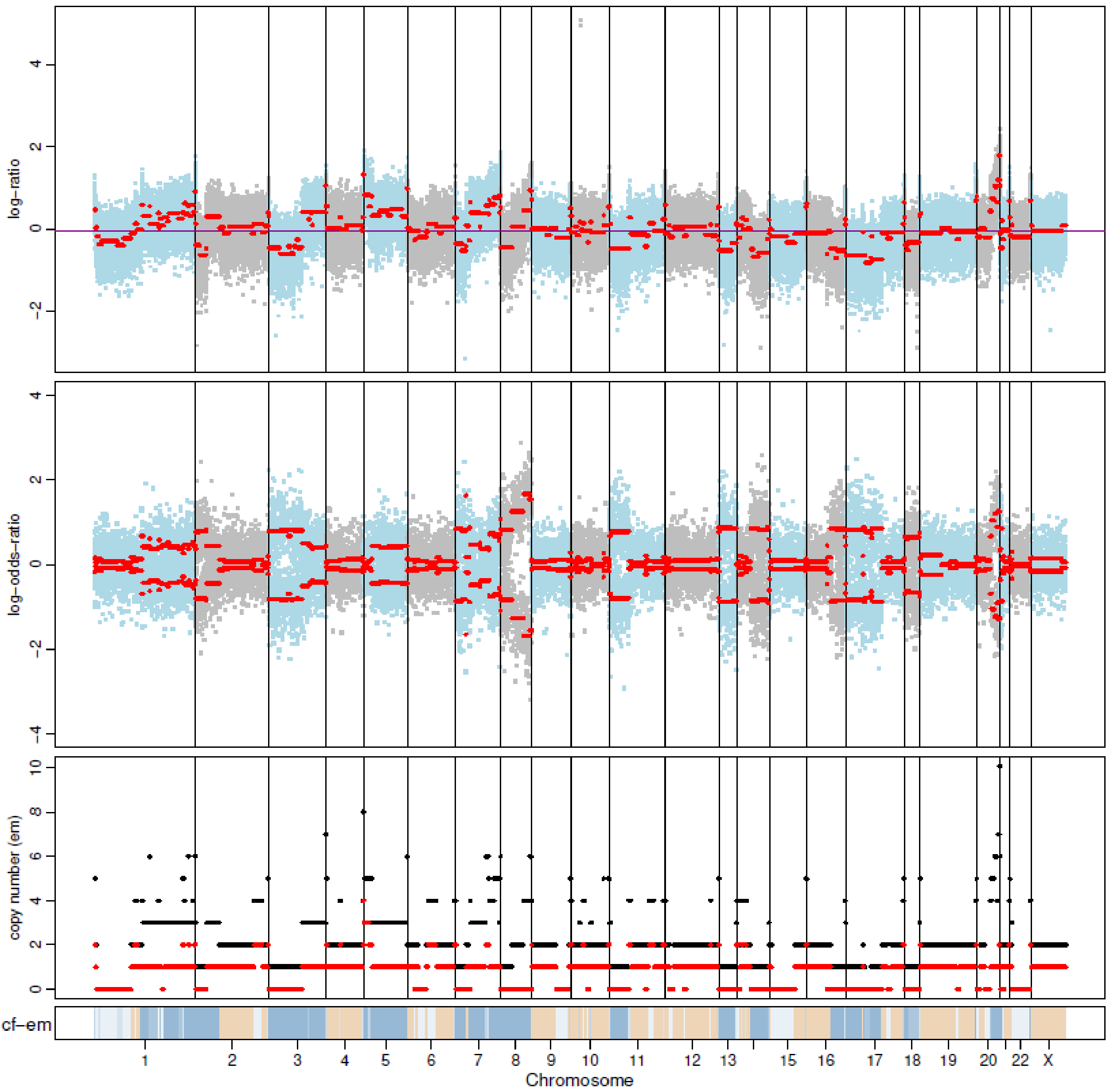


#### P21 Day 180

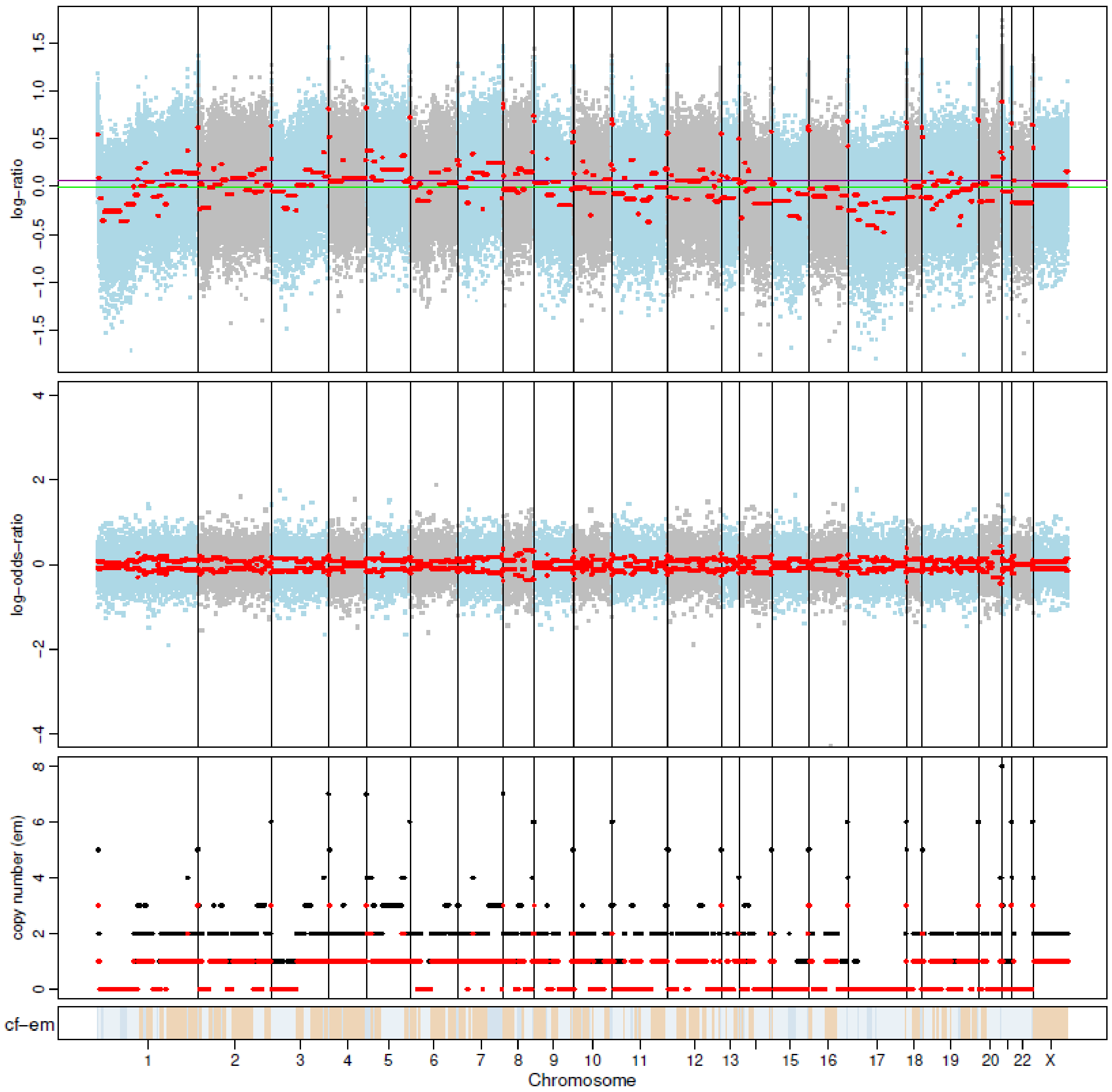

P22 Day 0

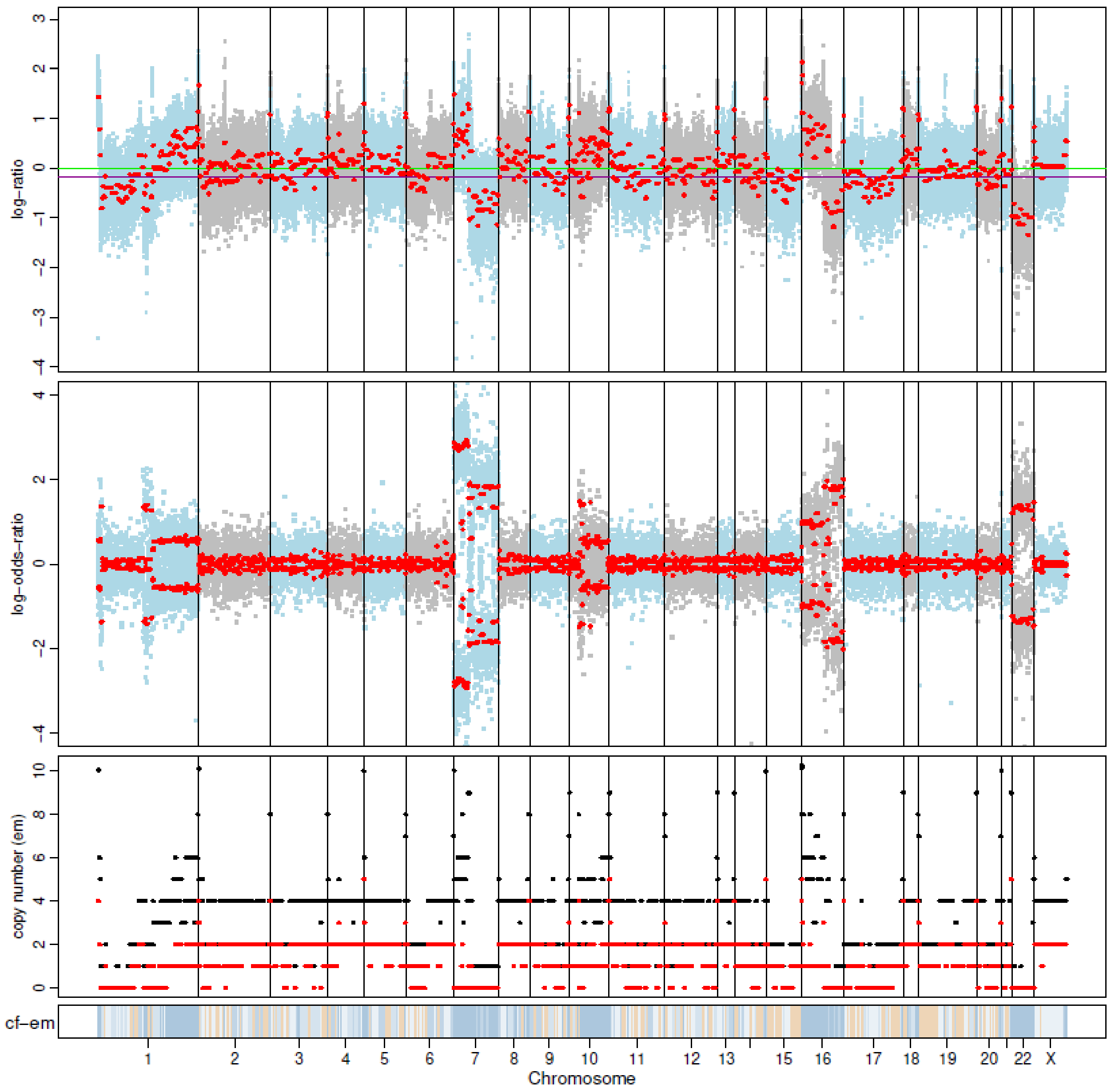


P22 Day 180

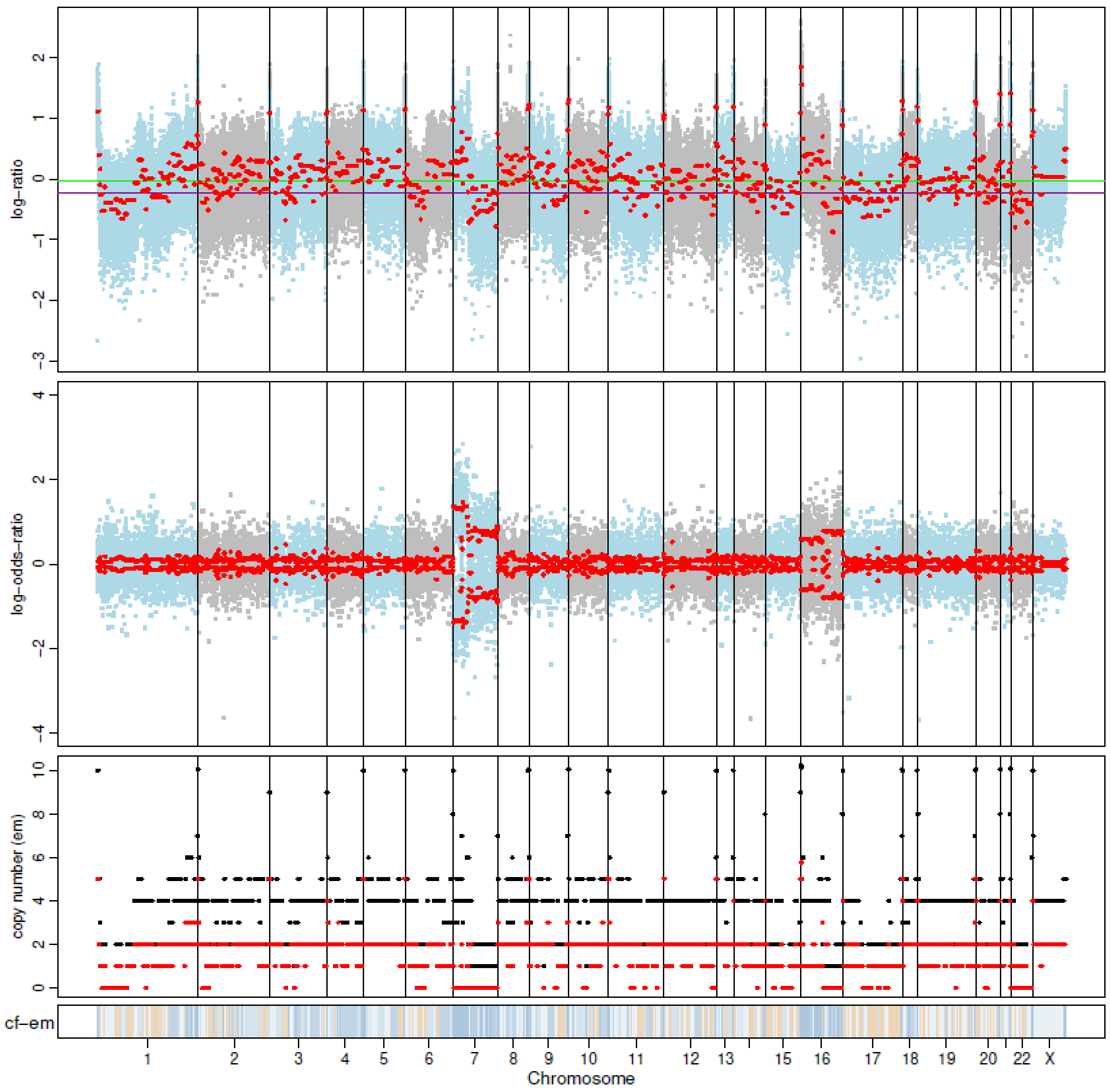

P23 Day 0

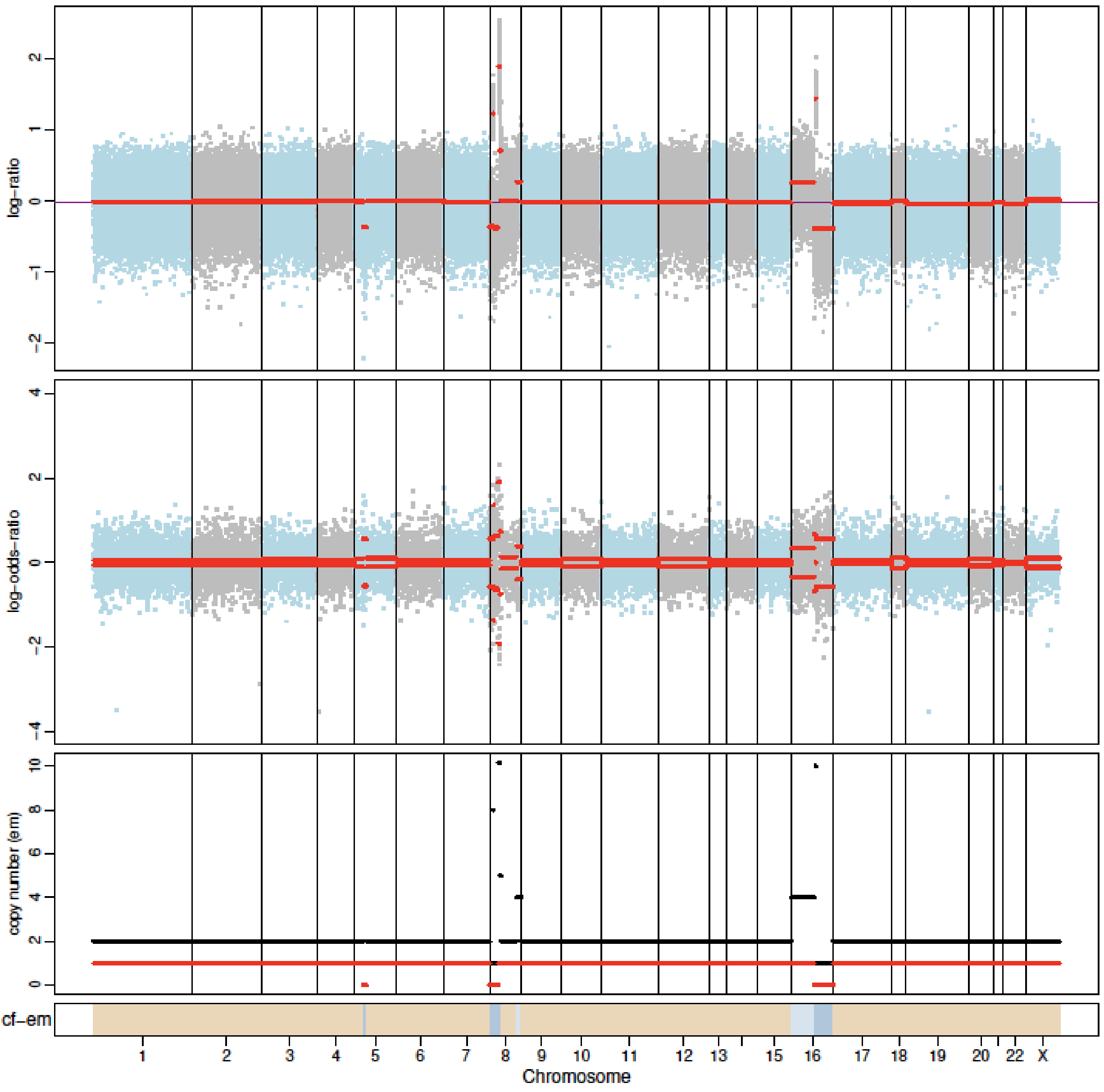


P23 Day 180

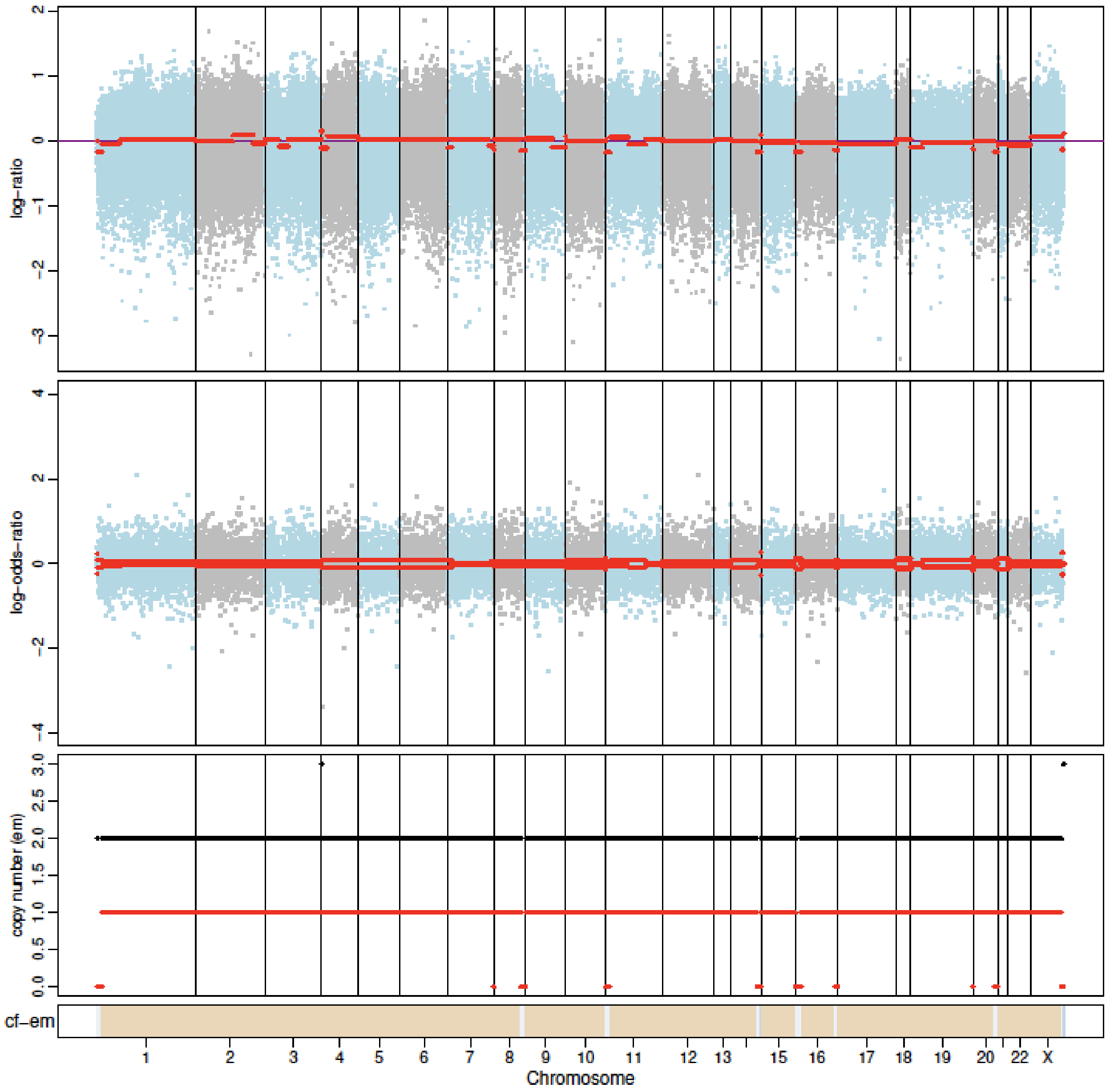

P25 Day 0

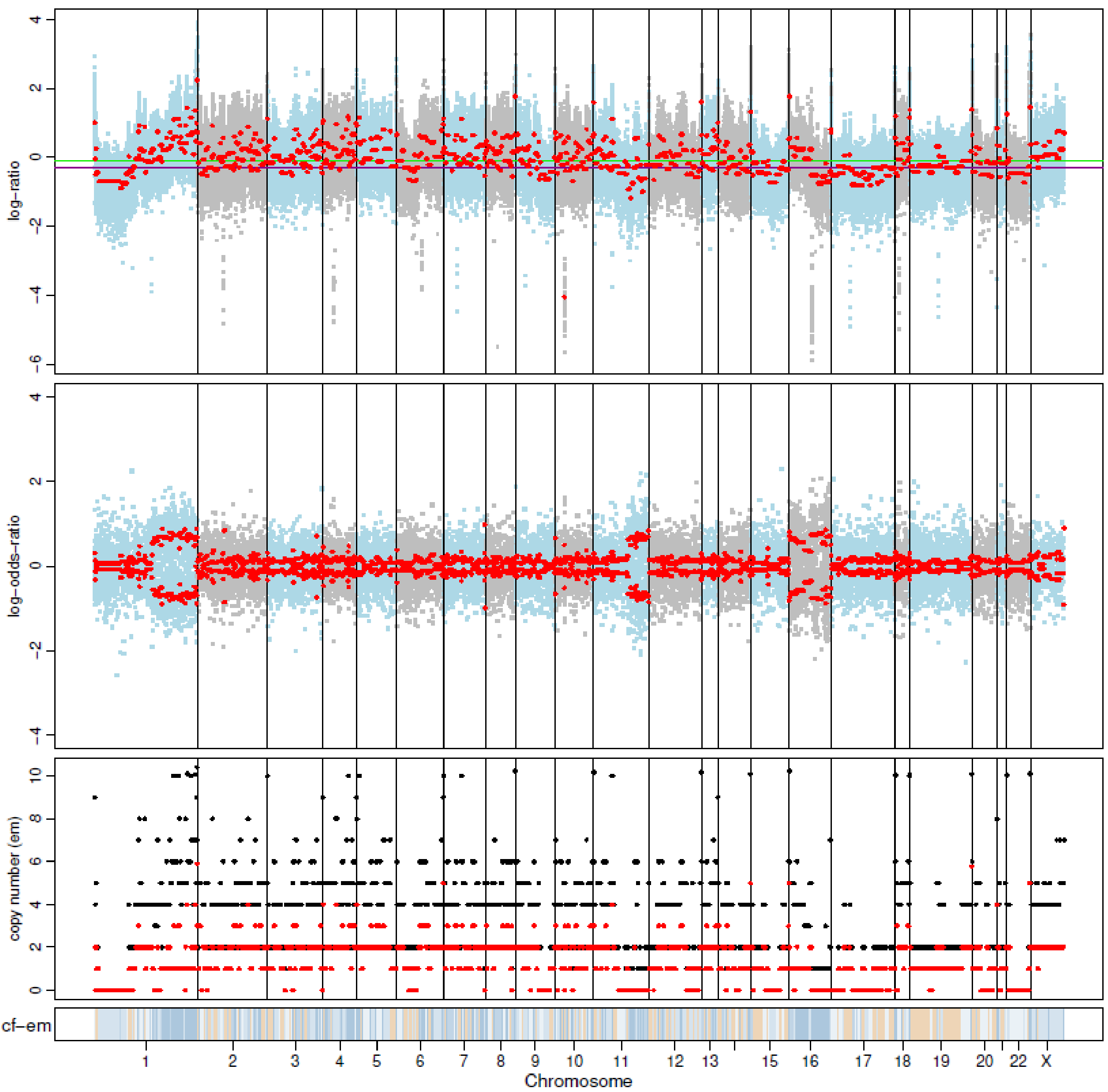


P25 Day 180

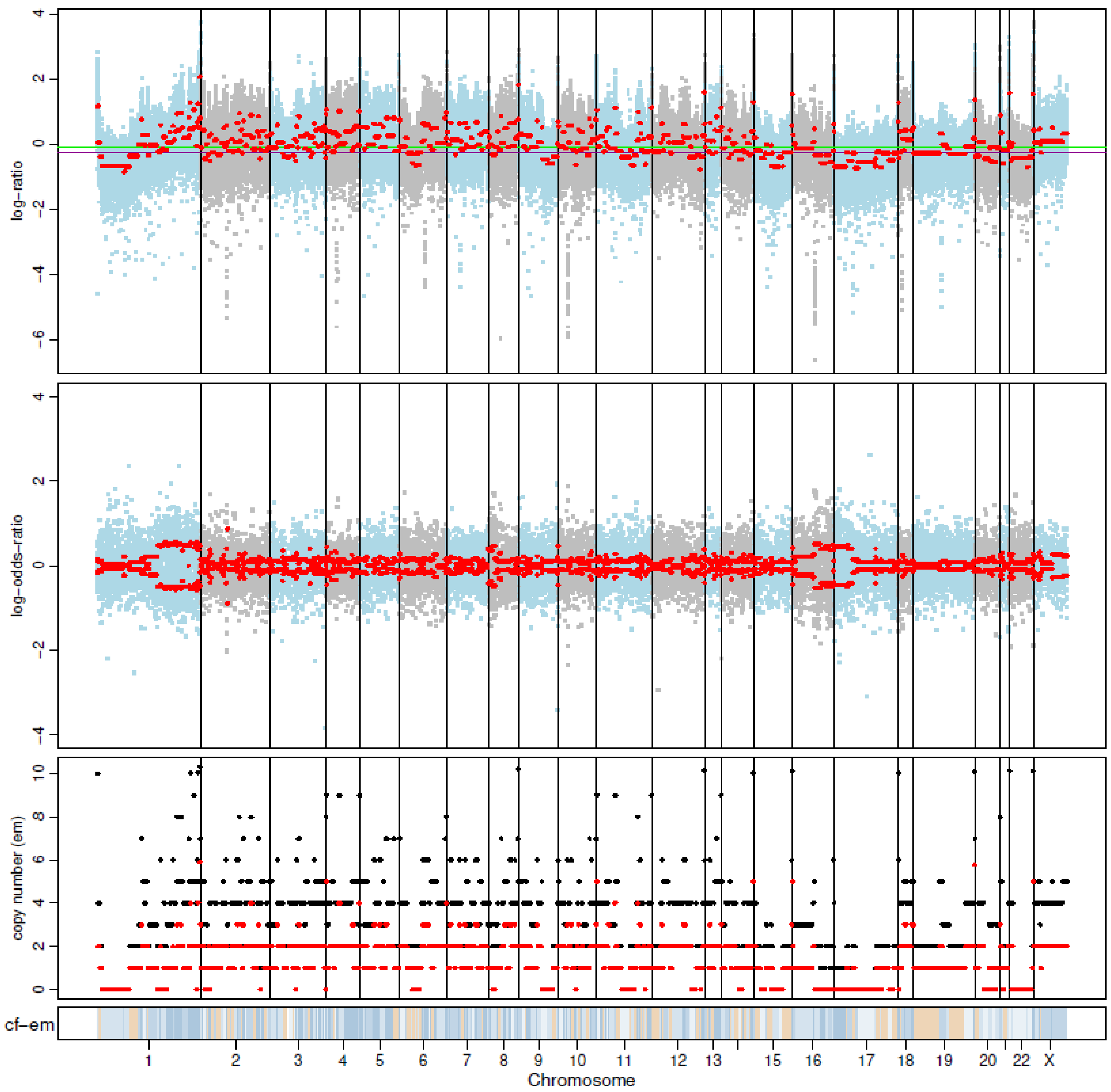

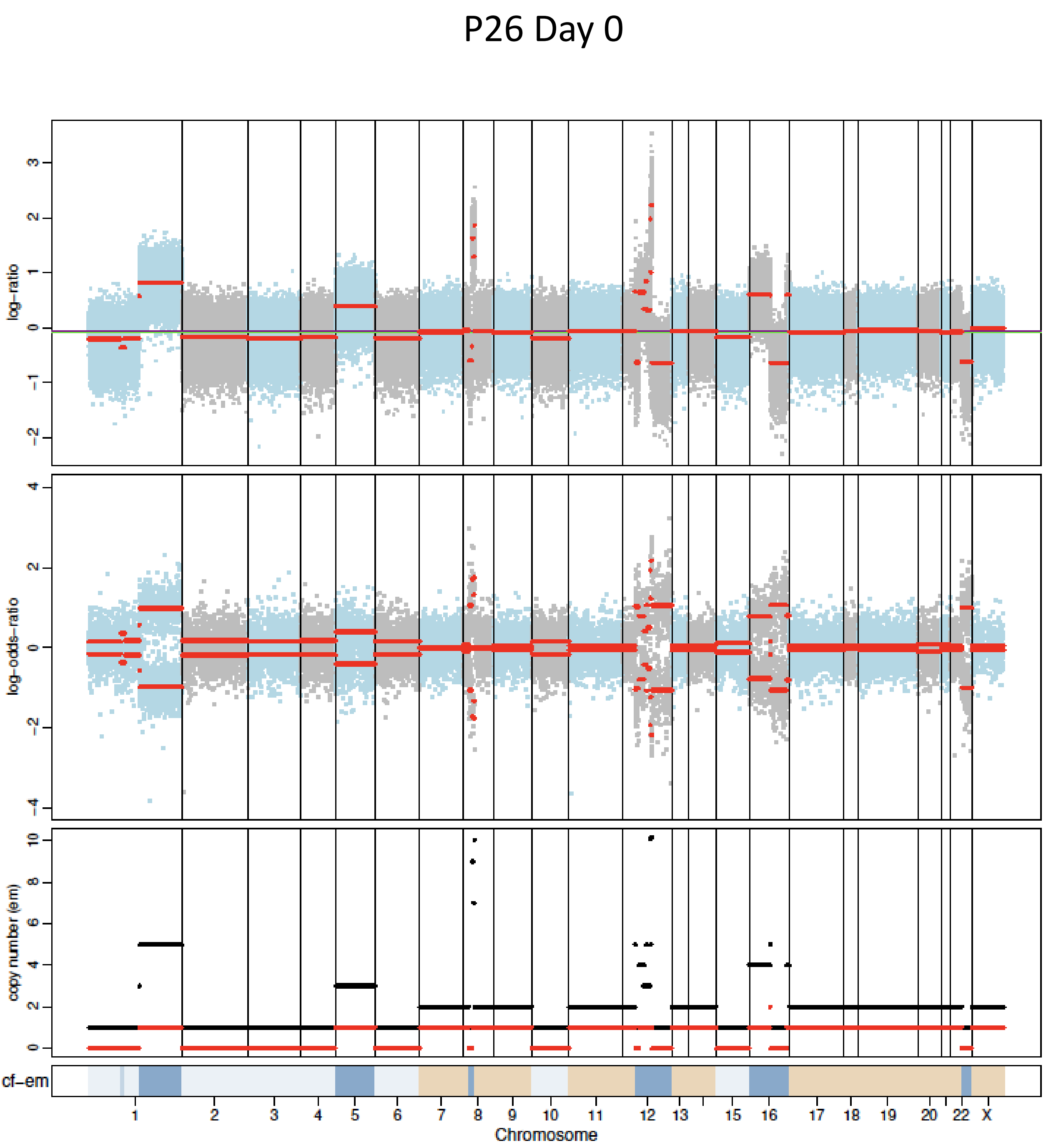
P26 Day 0
log-ratio
log-odds-ratio
copy number (em)
cf-em
Chromosome

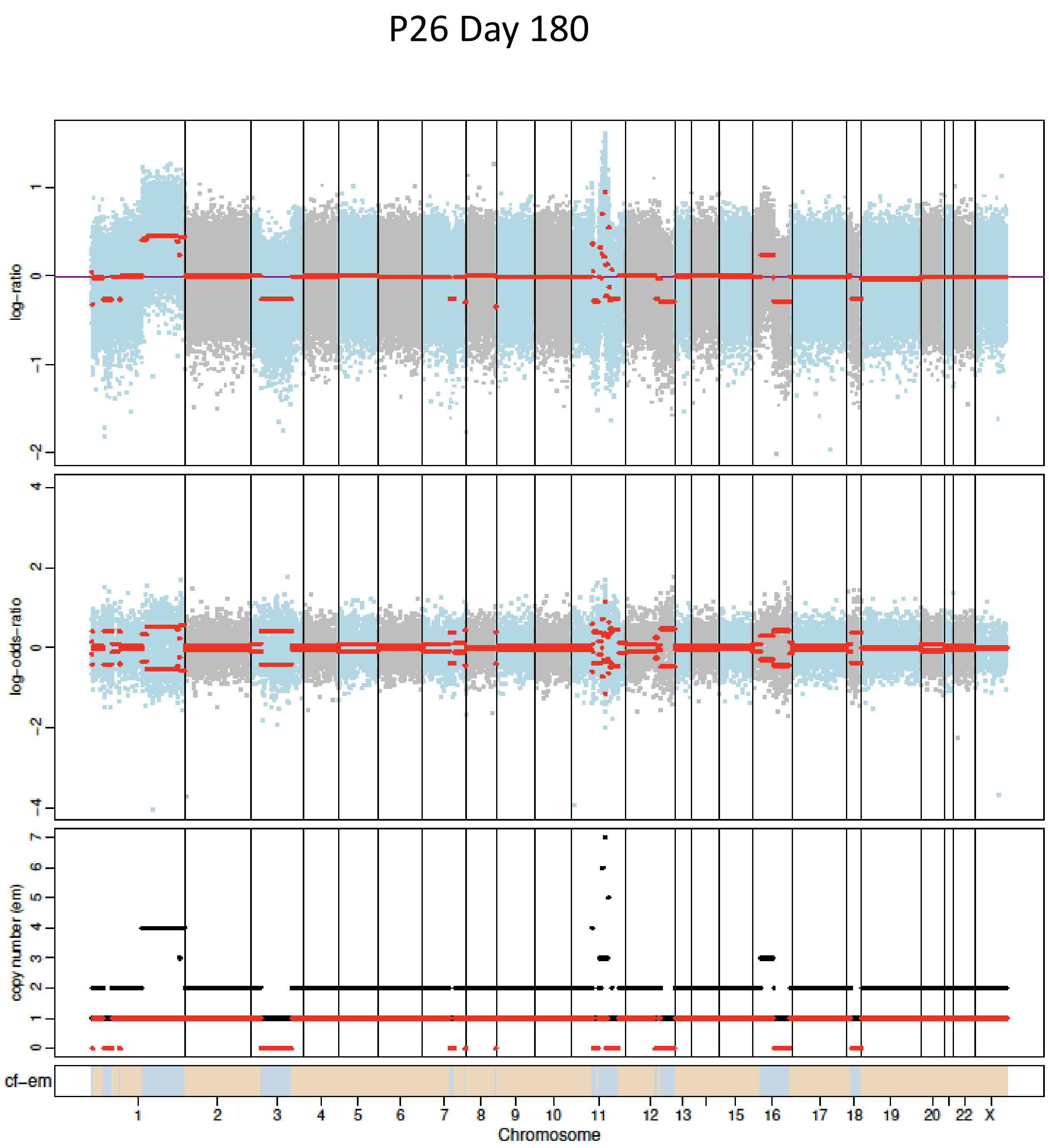
P26 Day 180
log-ratio
log-odds-ratio
copy number (em)
cf-em
Chromosome

P27 Day 0

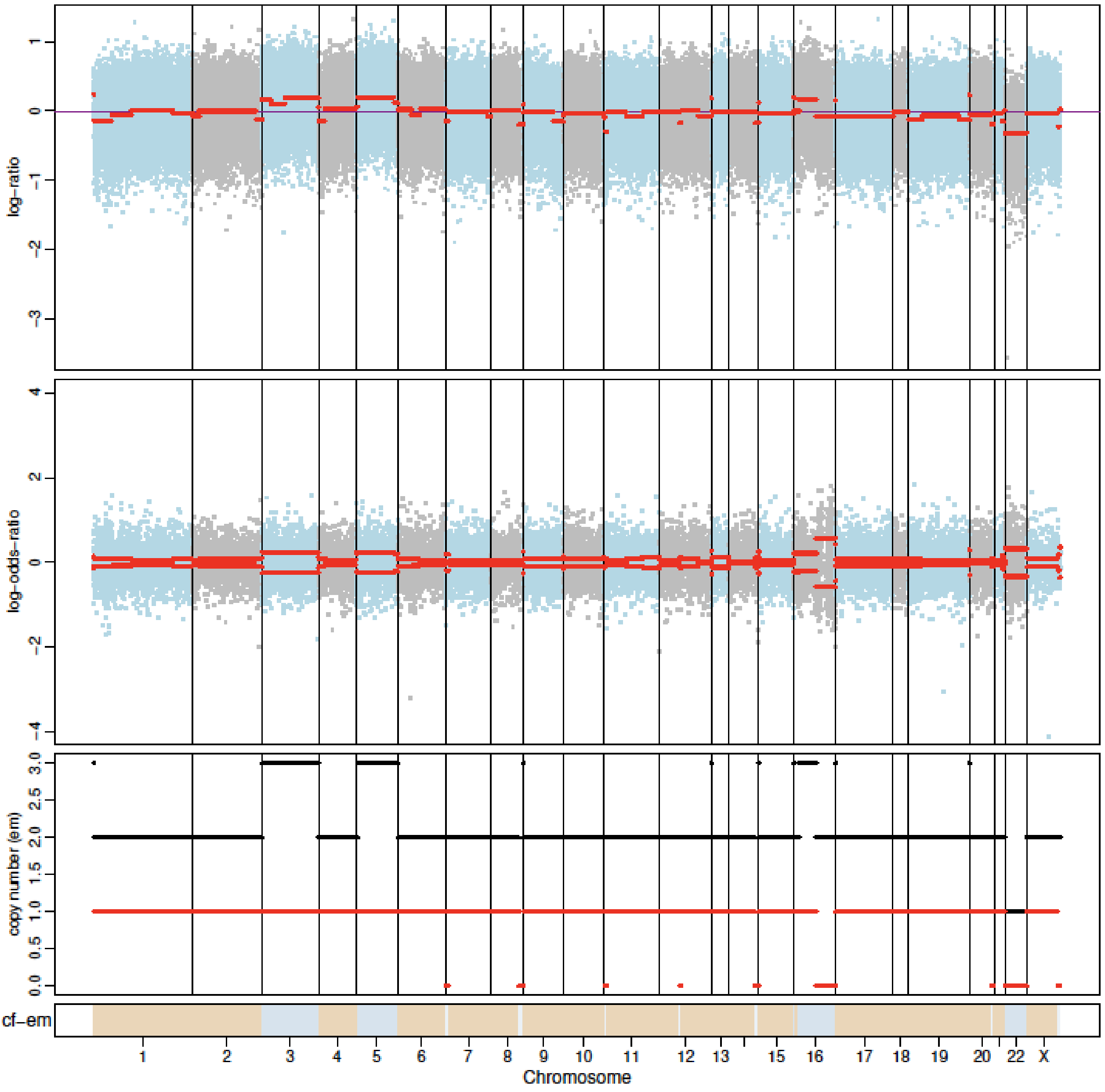


P27 Day 180

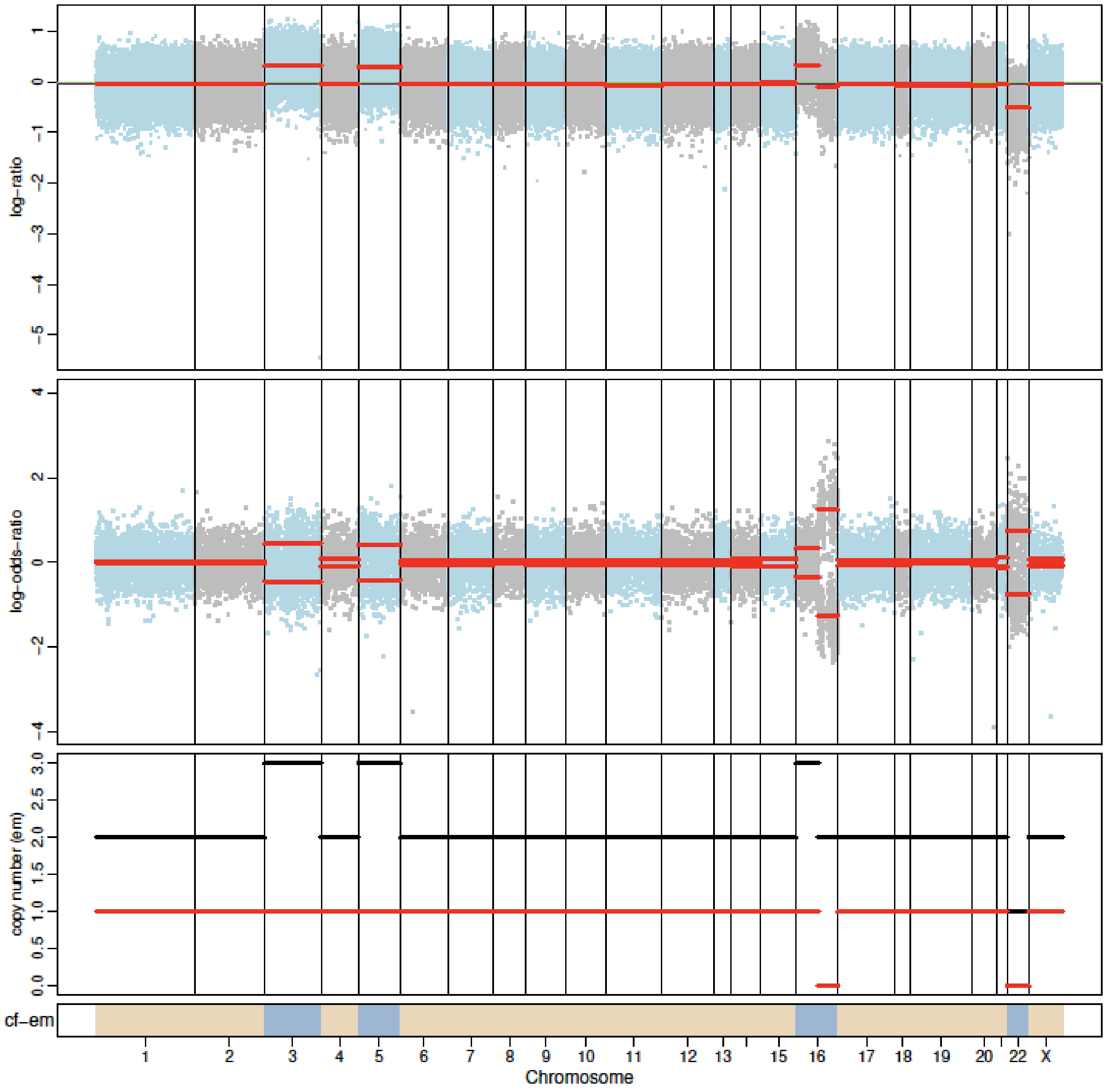

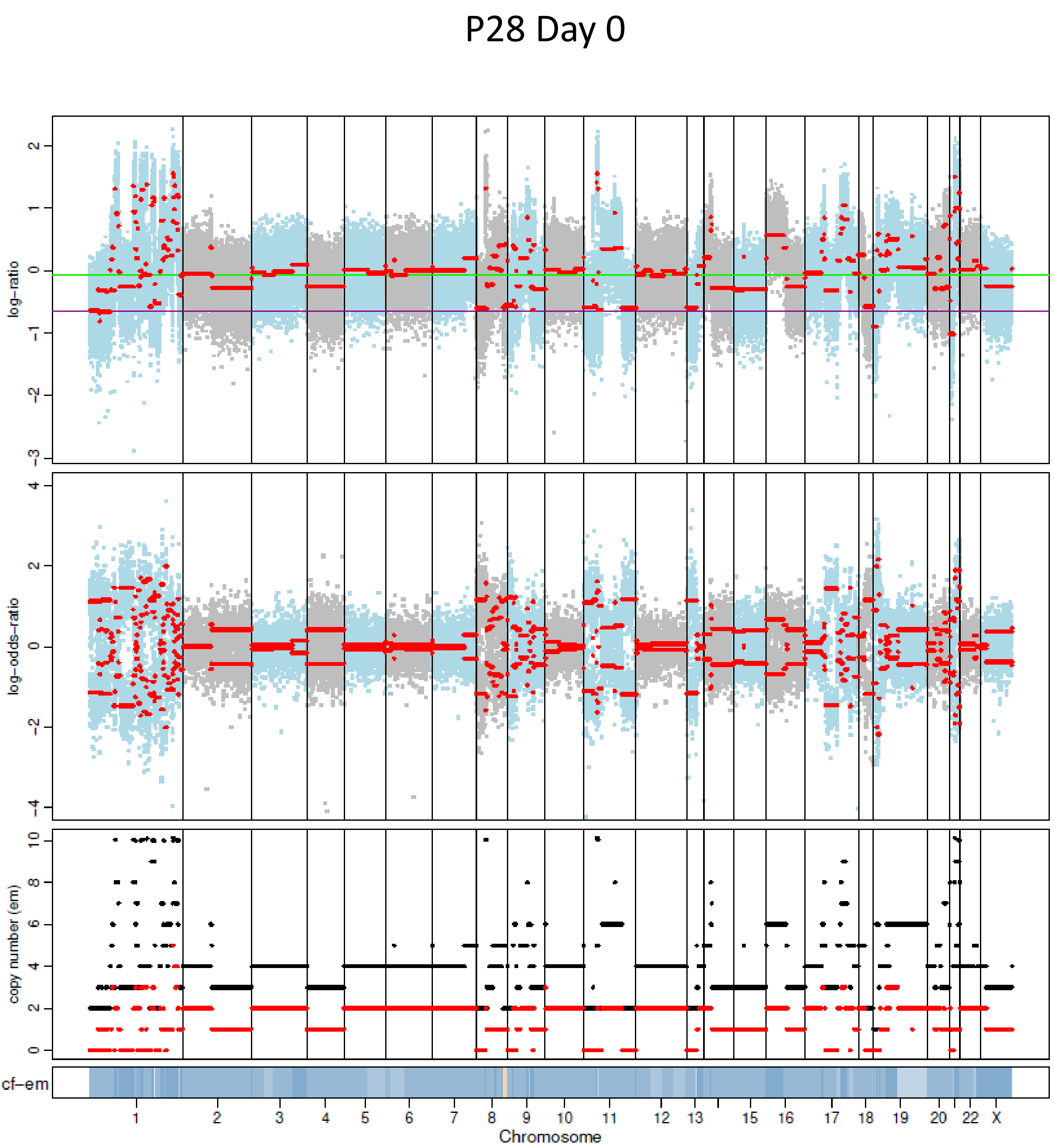
P28 Day 0
log-ratio
log-odds-ratio
copy number (em)
cf-em
Chromosome

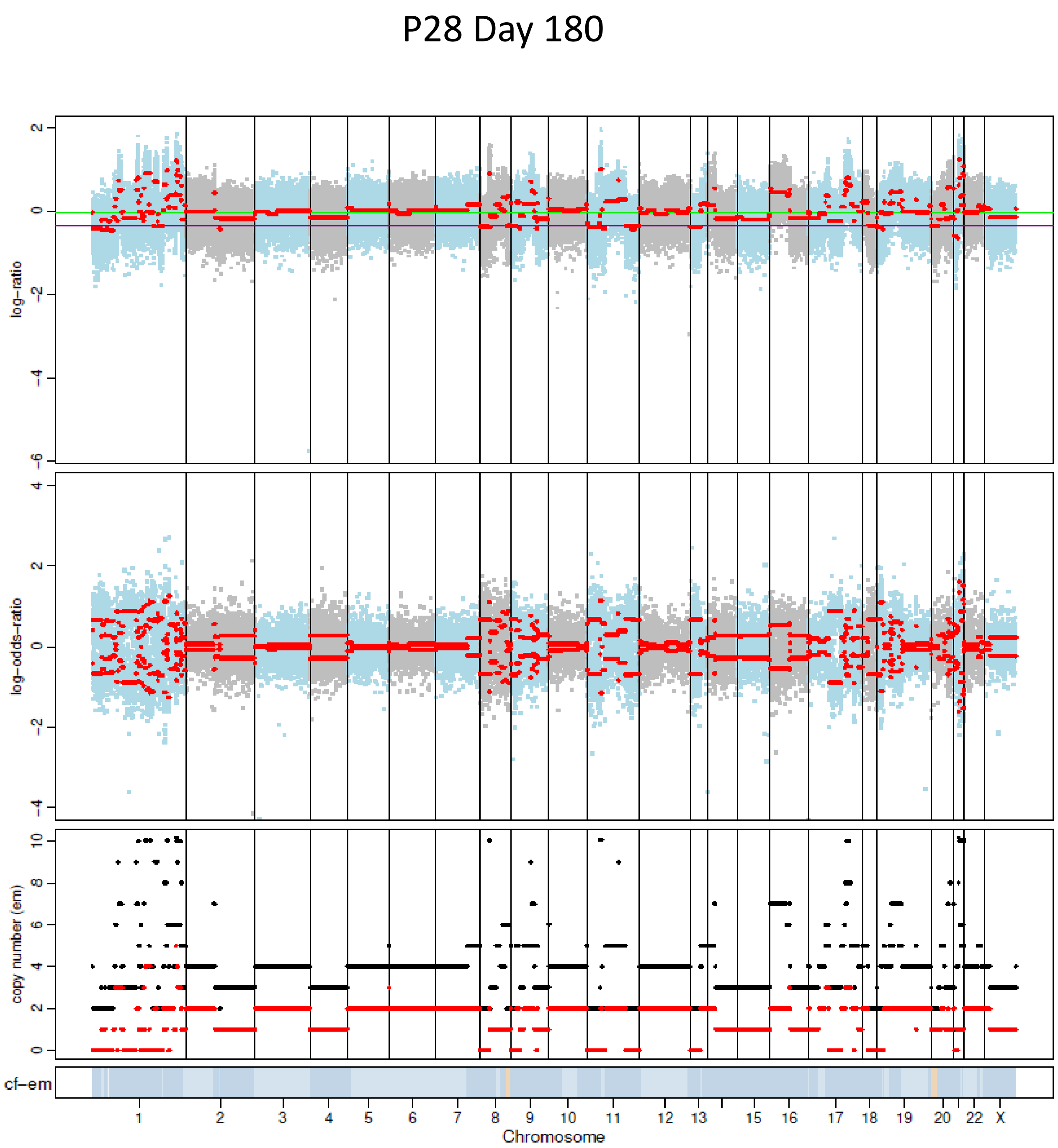
P28 Day 180
log-ratio
log-odds-ratio
copy number (em)
cf-em
Chromosome

### P29 Day 0

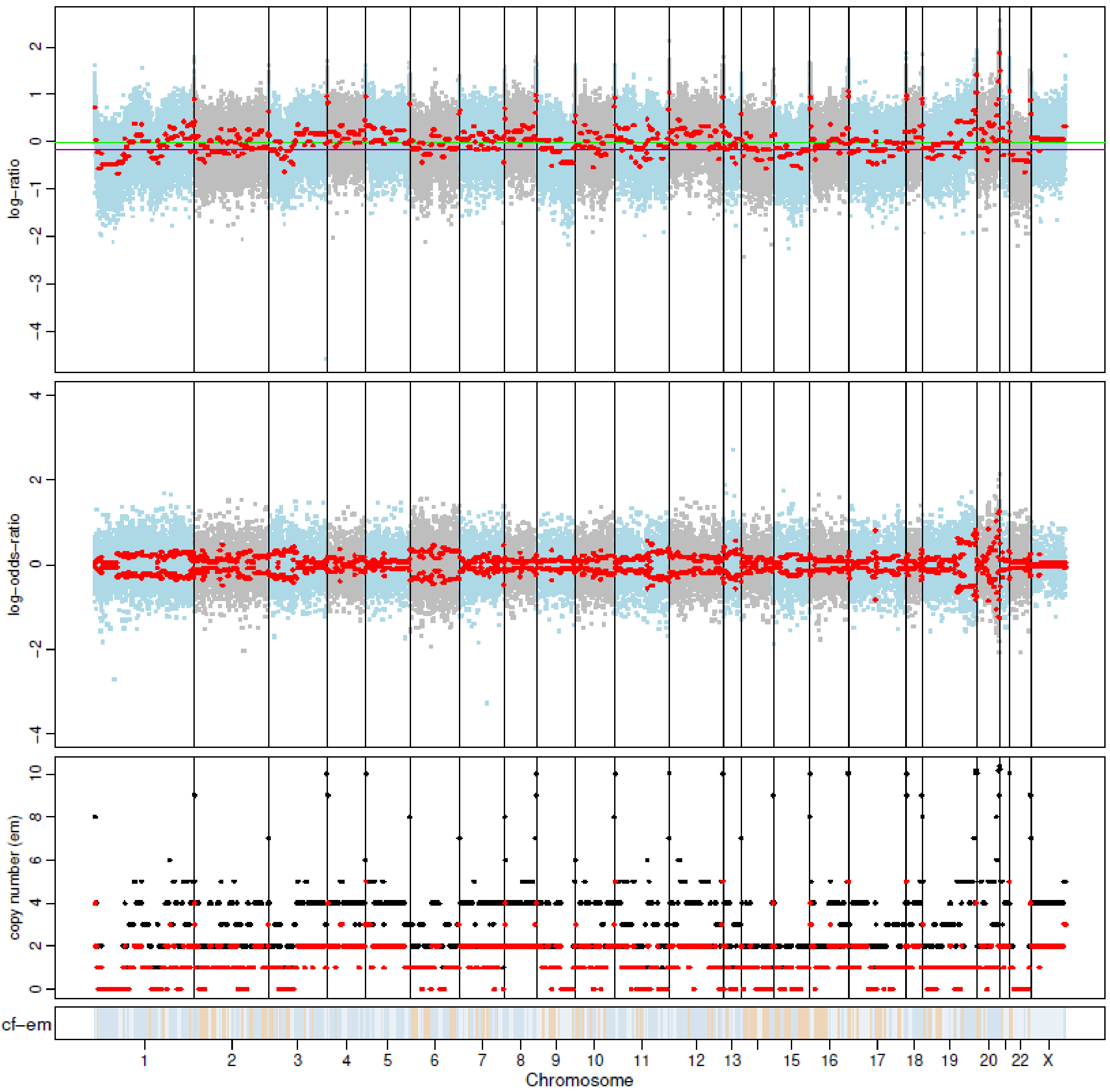


### P29 Day 180

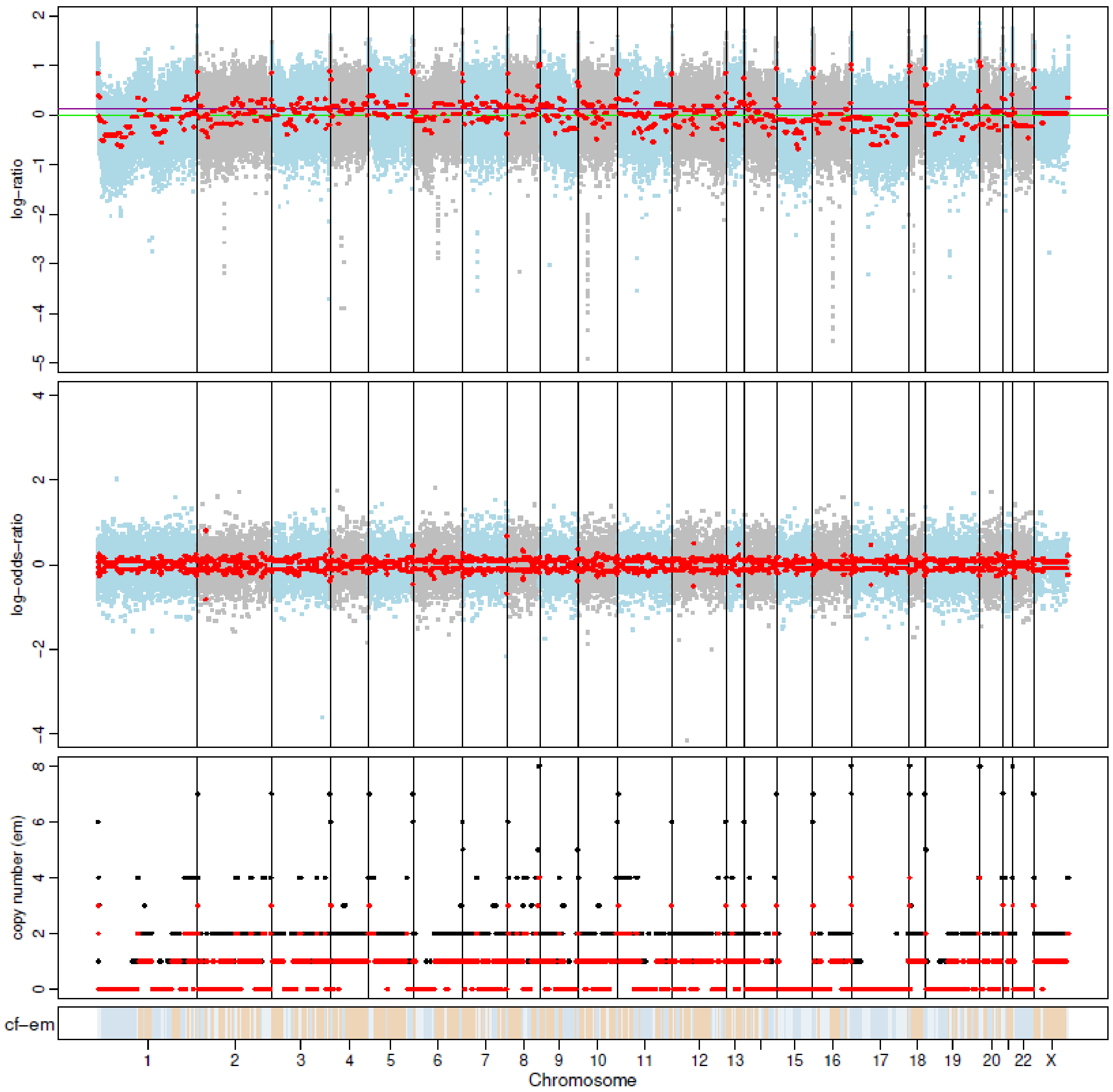

P31 Day 0

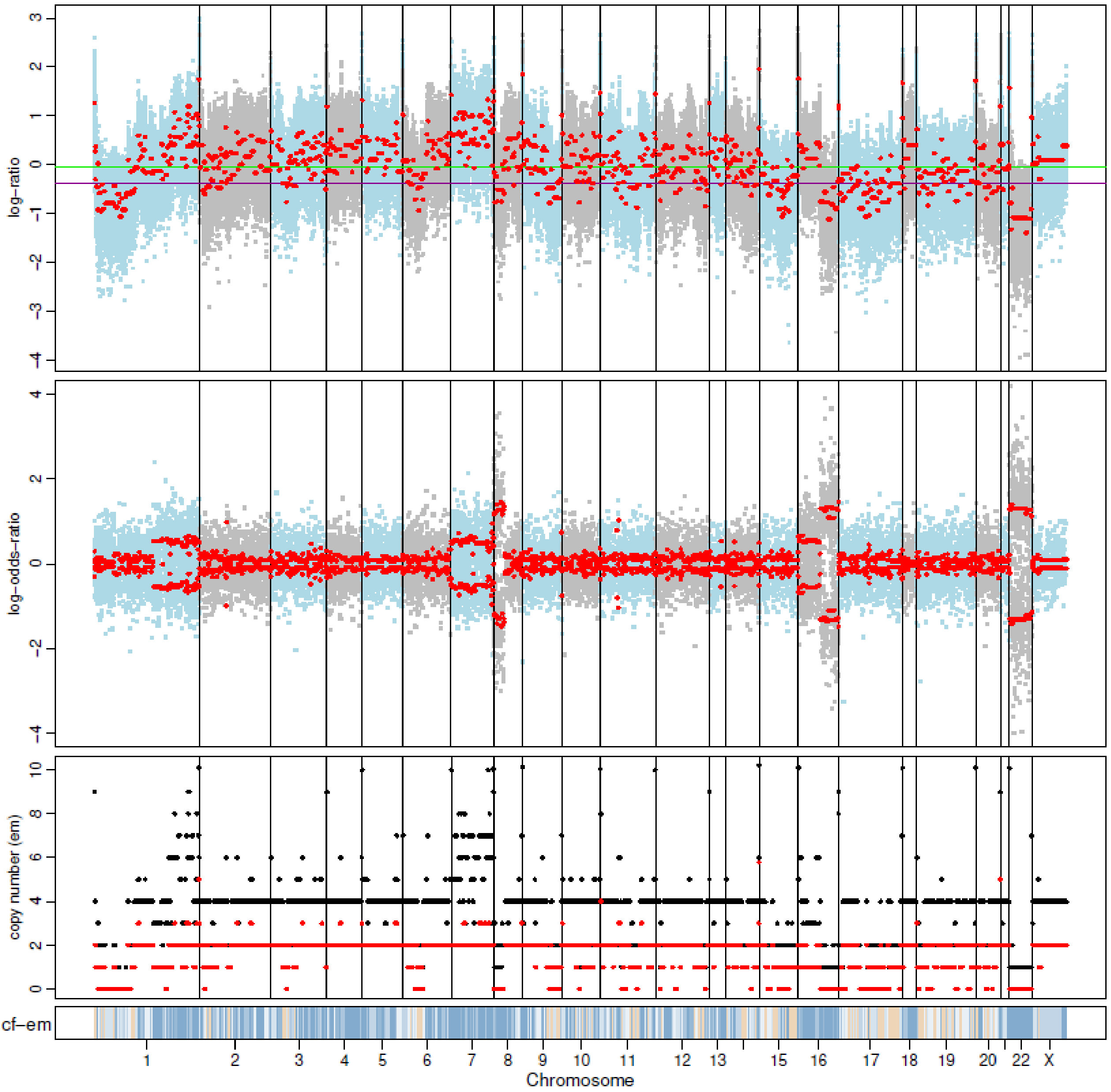


P31 Day 180

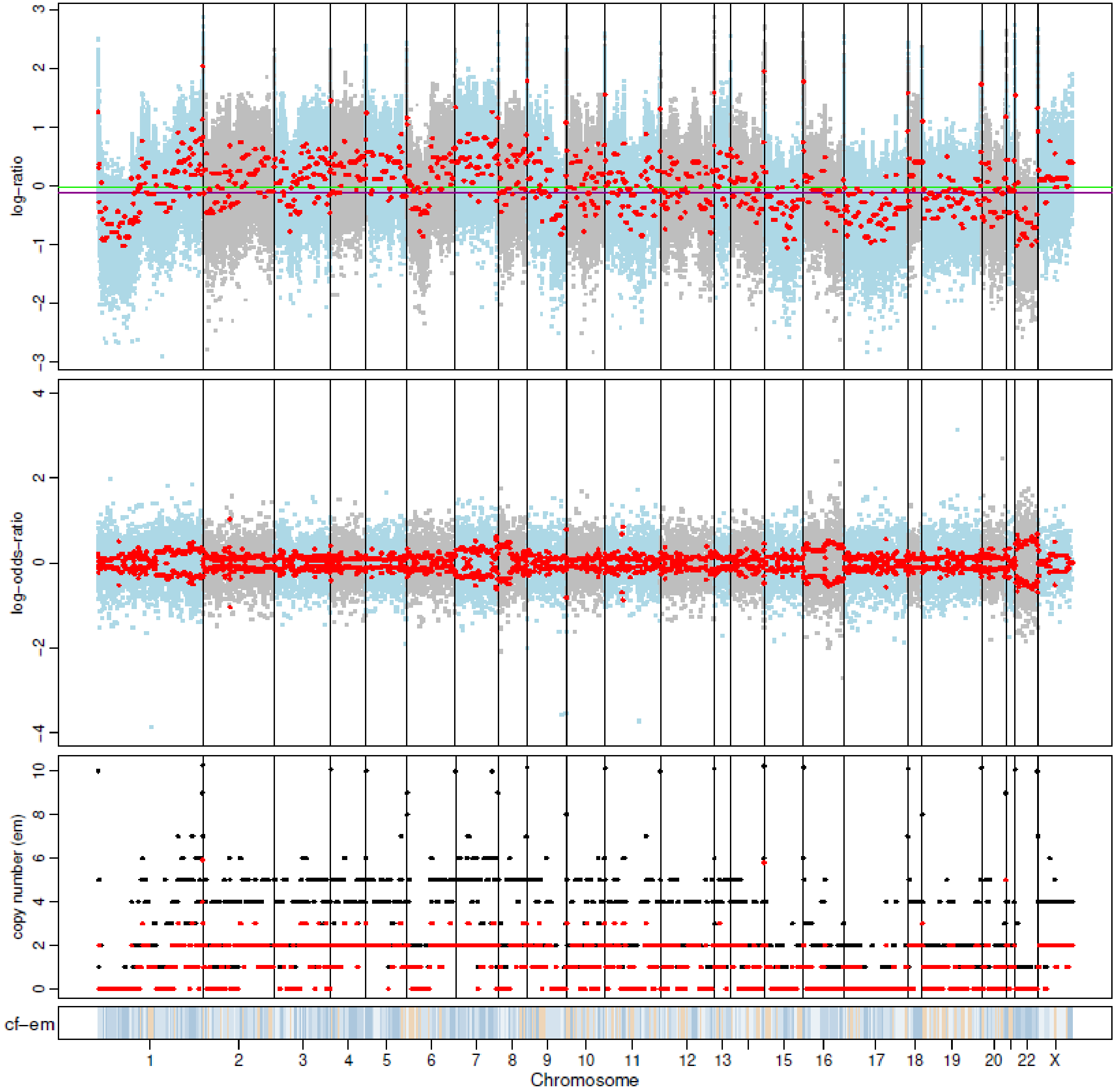

P34 Day 0

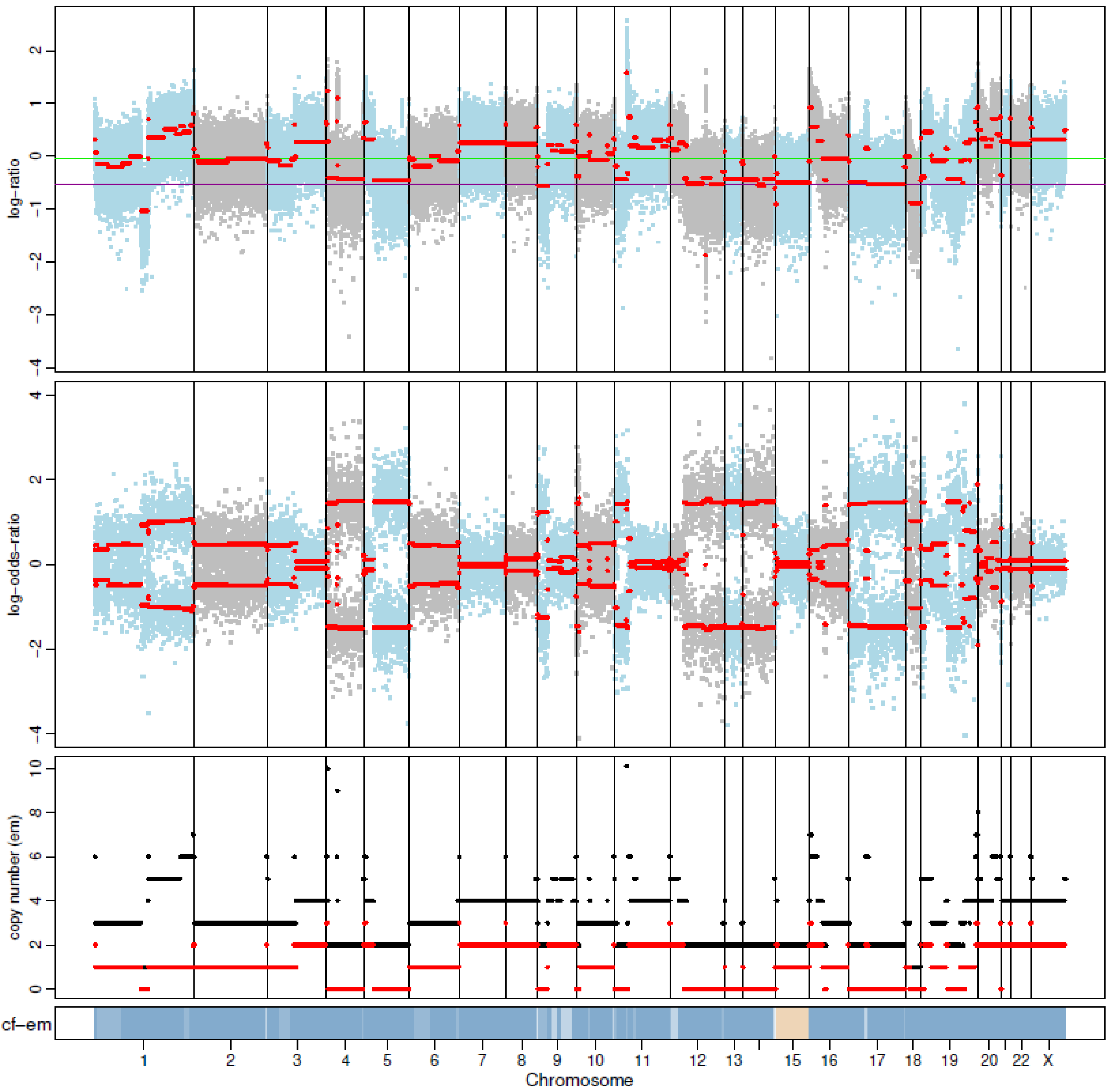


P34 Day 180

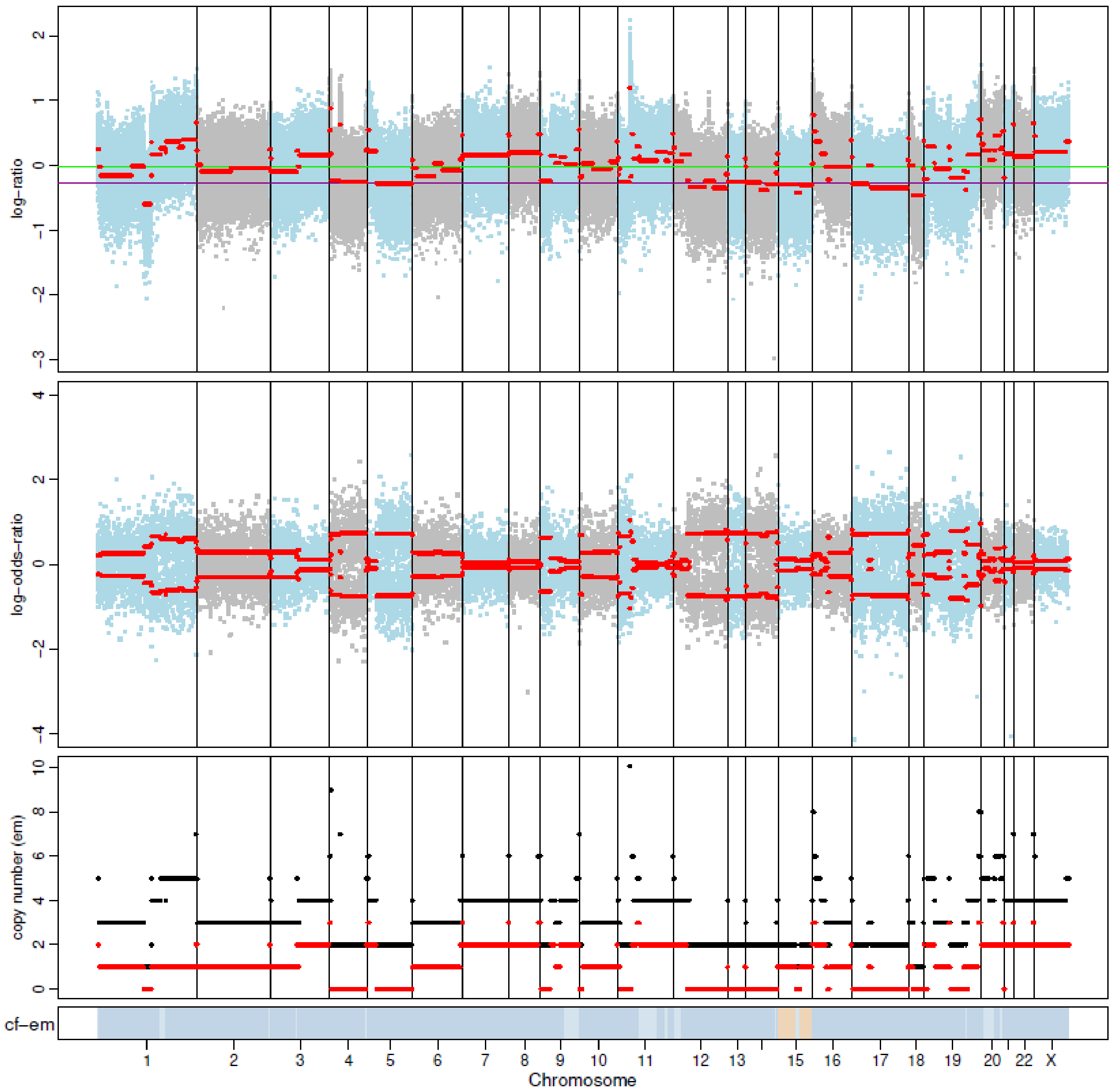

P35 Day 0

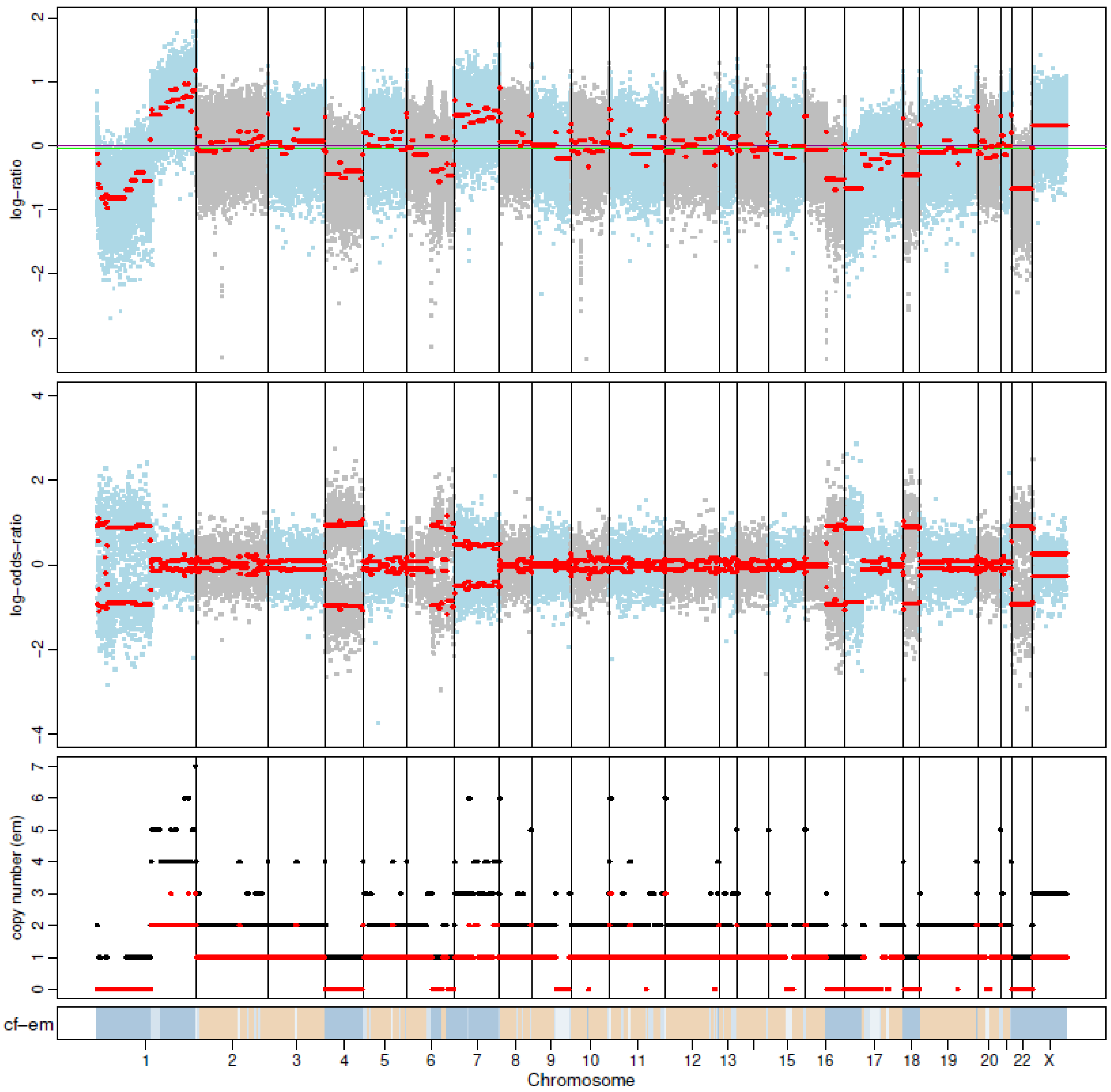


P35 Day 180

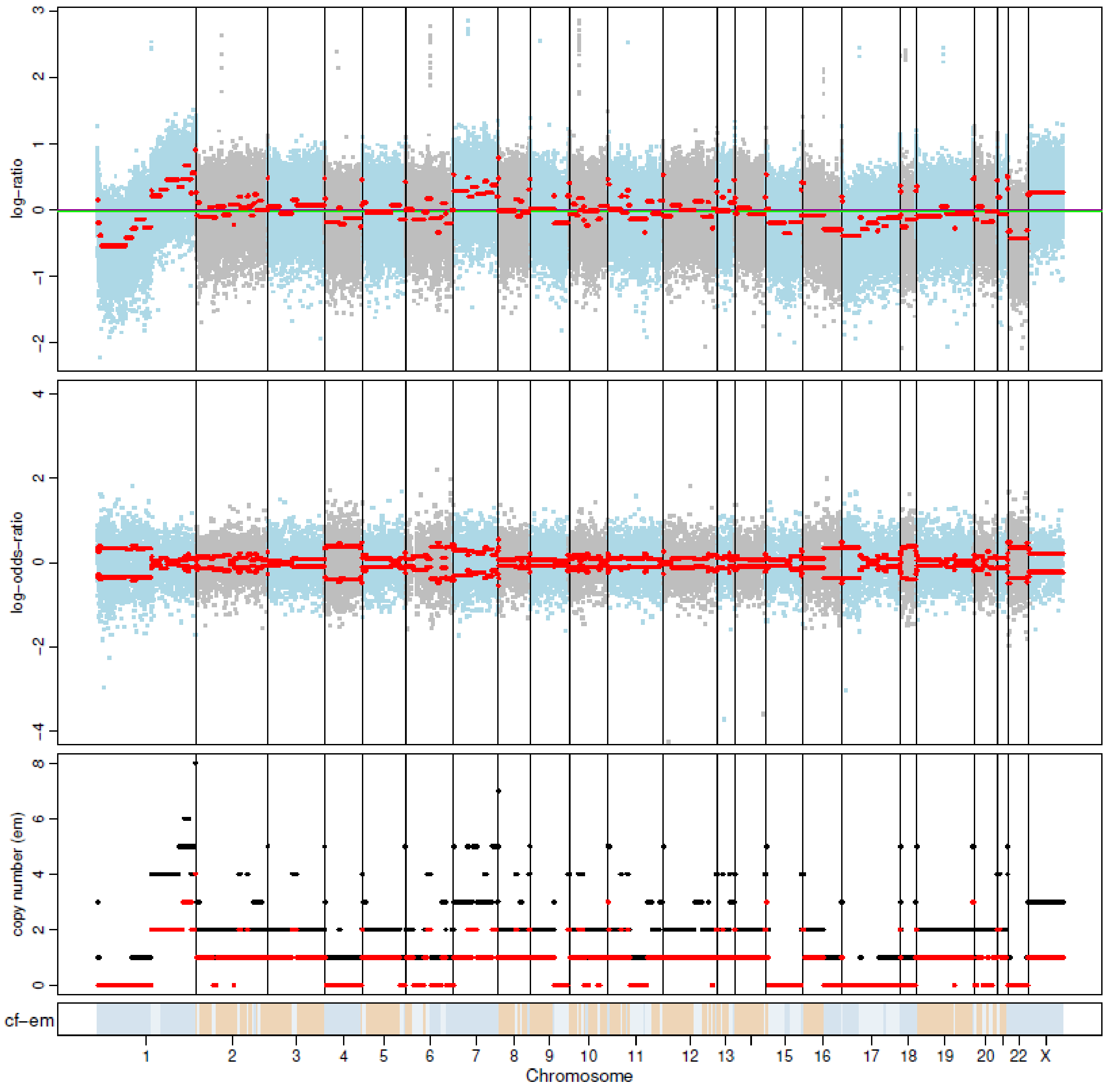

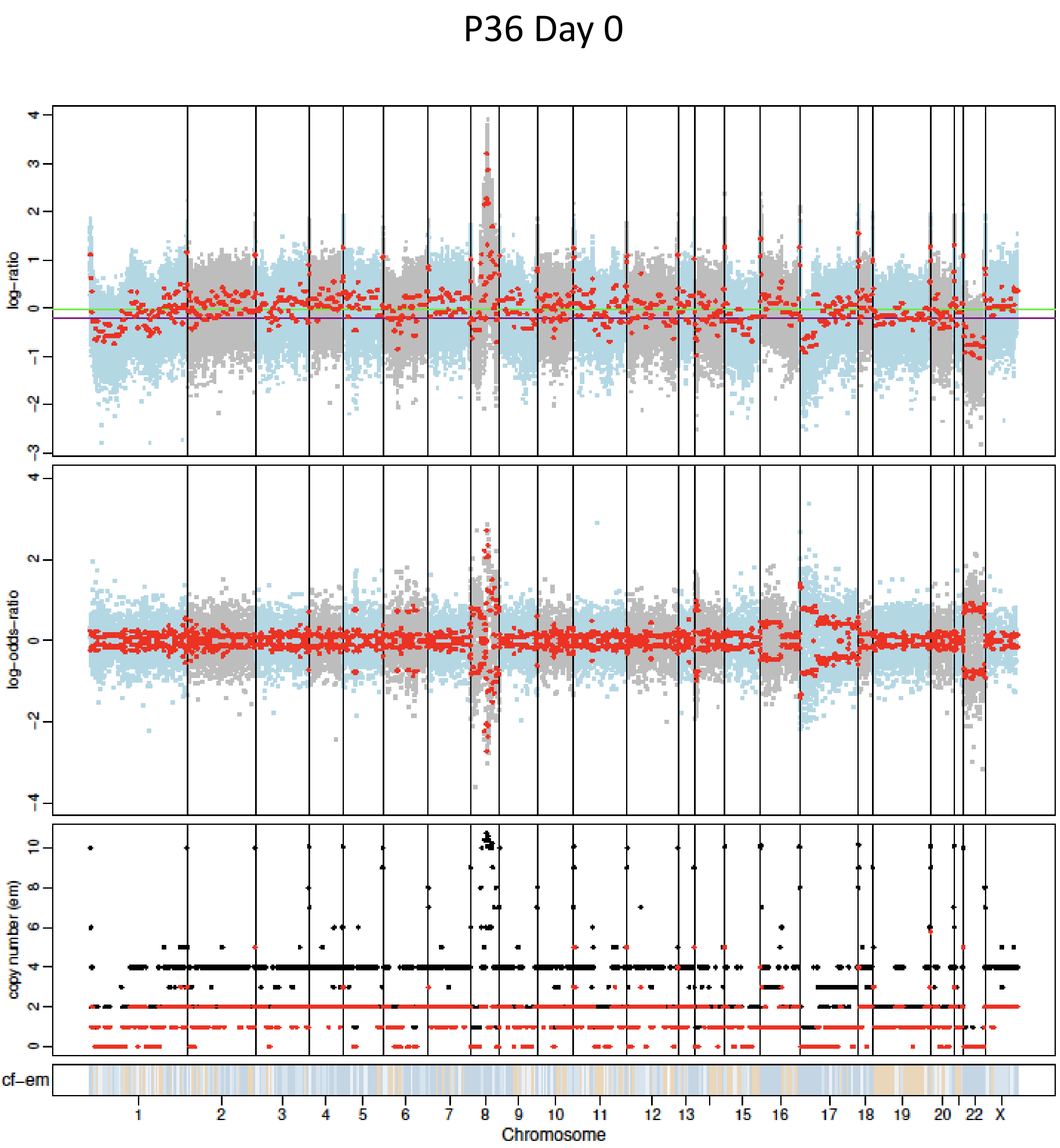
P36 Day 0
log-ratio
log-odds-ratio
copy number (em)
cf-em
1
2
3
4
5
6
7
8
9
10
11
12
13
15
16
17
18
19
20
22
X
Chromosome

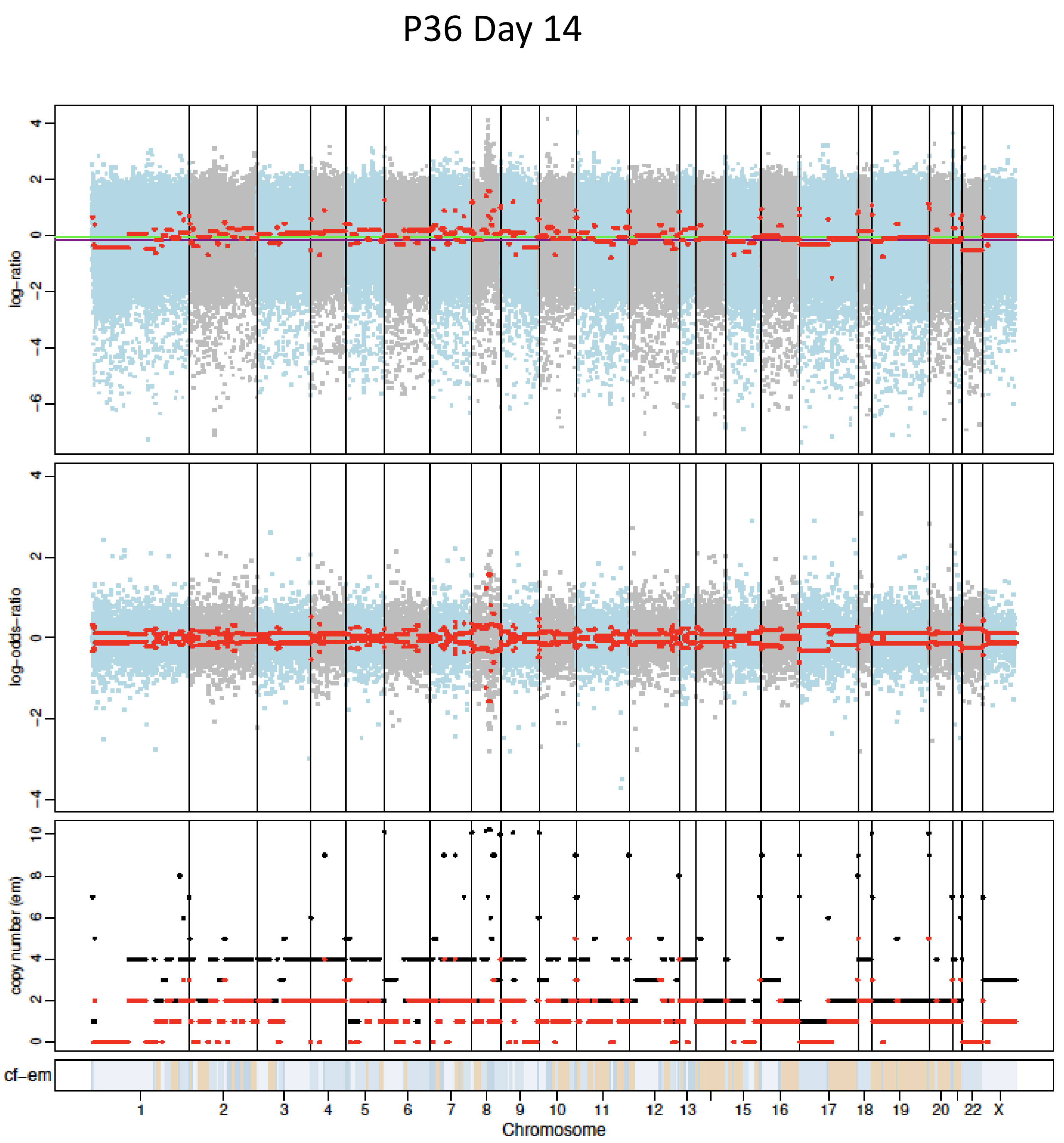
P36 Day 14
log-ratio
log-odds-ratio
copy number (em)
cf-em
1
2
3
4
5
6
7
8
9
10
11
12
13
15
16
17
18
19
20
22
X
Chromosome

### P37 Day 0

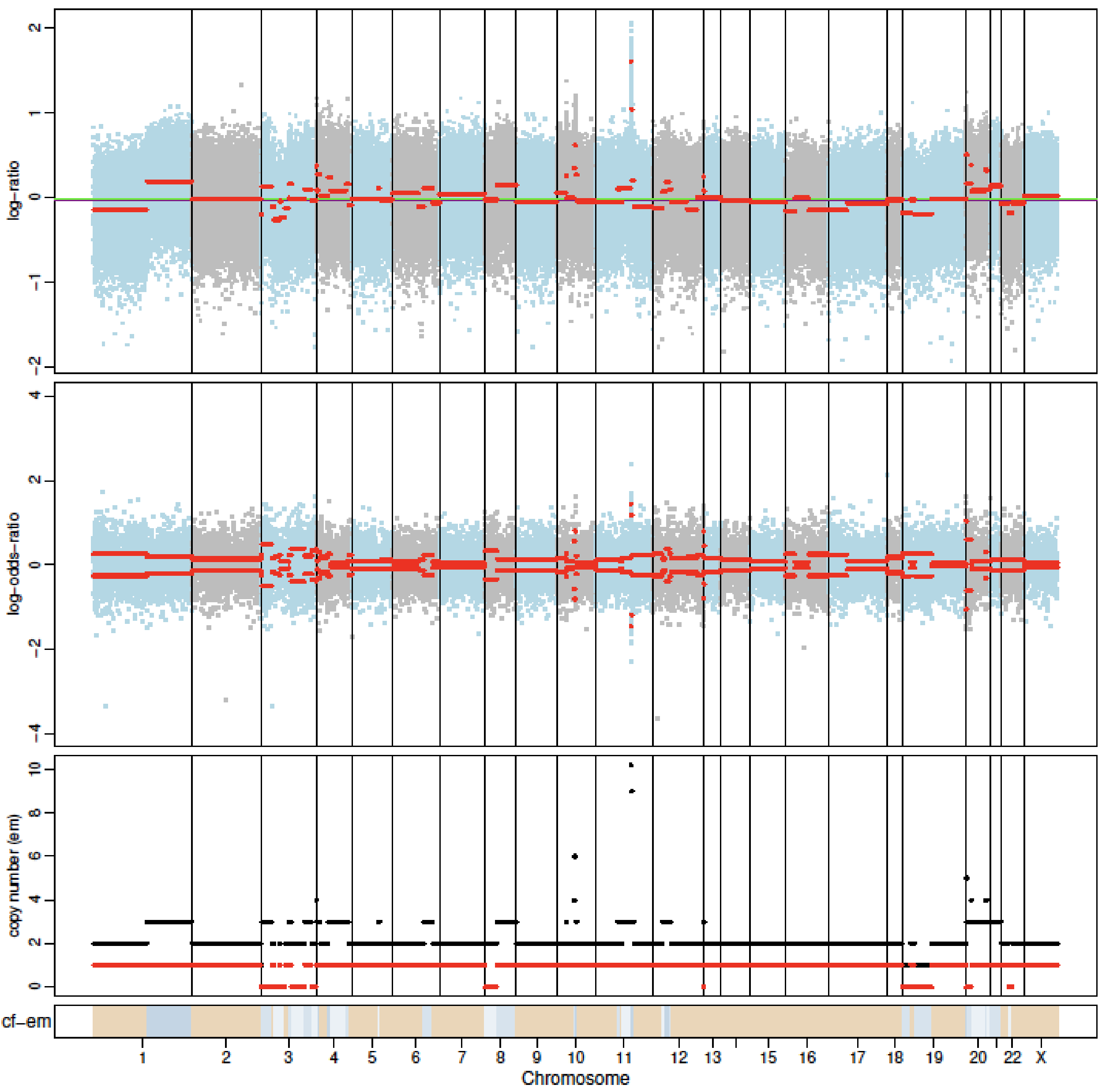


### P37 Day 180

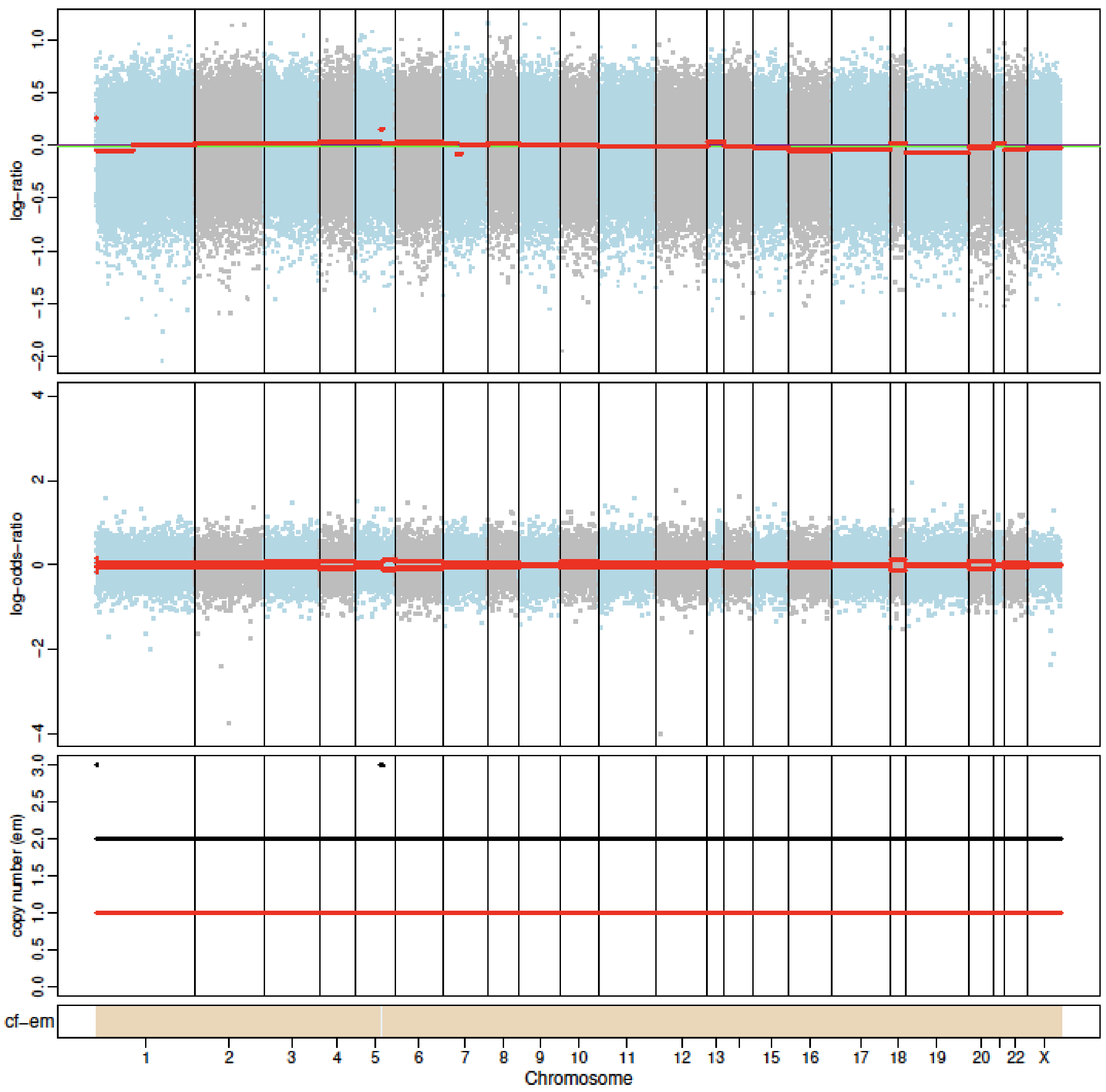

P39 Day 0

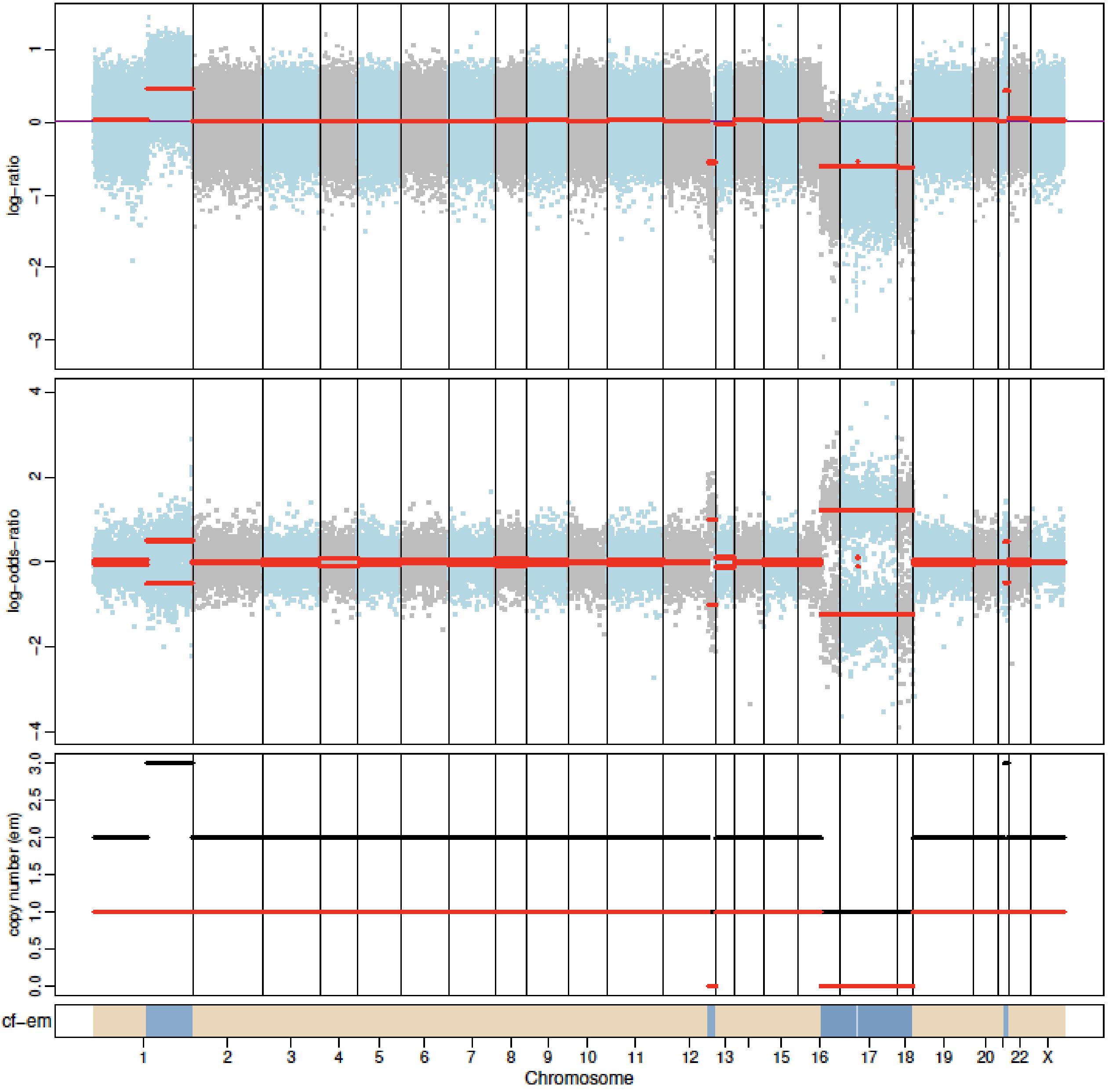


P39 Day 180

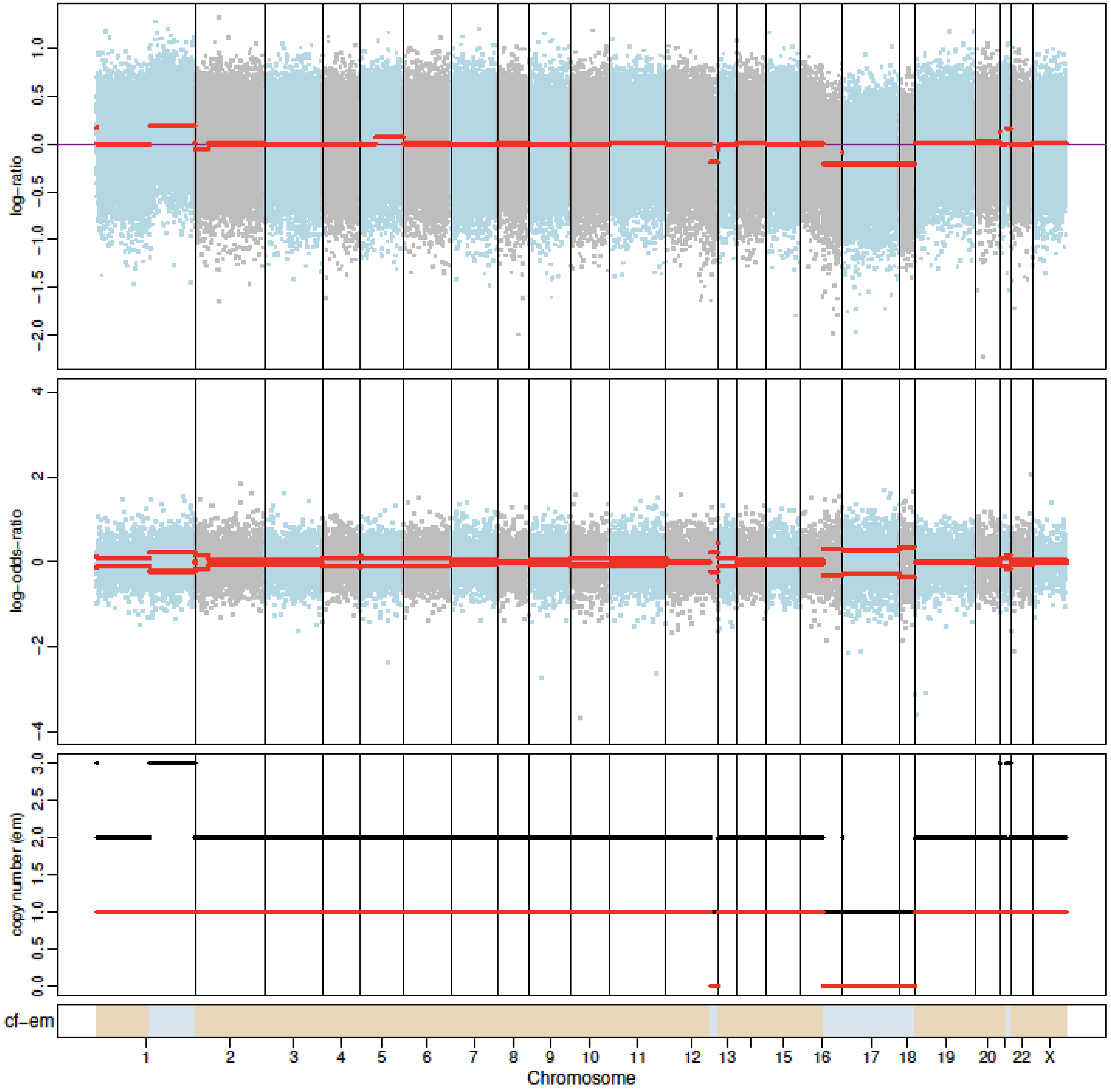

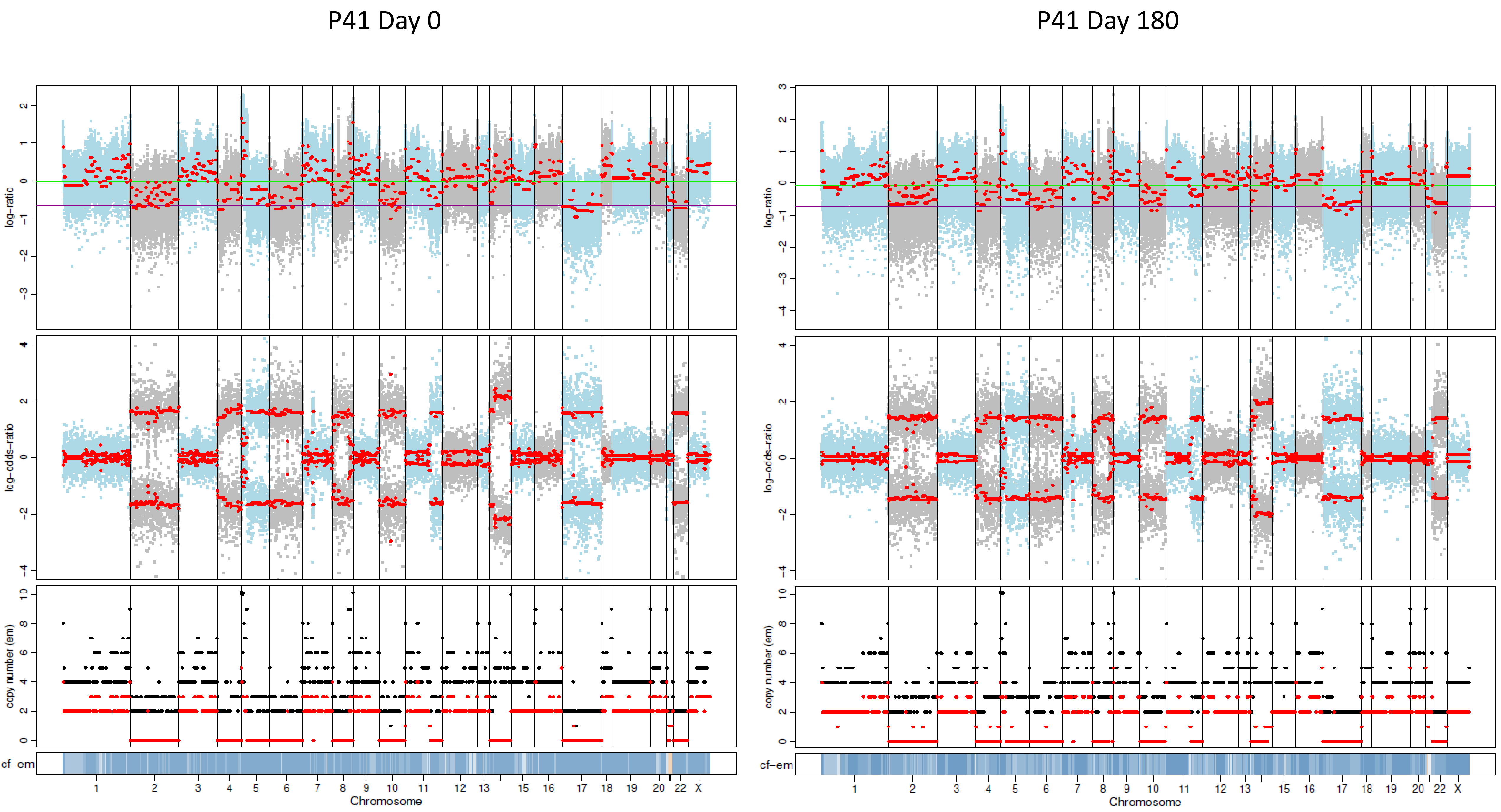

P41 Day 0
P41 Day 180
log–ratio
log–odds–ratio
copy number (em)
cf–em
Chromosome
1
2
3
4
5
6
7
8
9
10
11
12
13
15
16
17
18
19
20
22
X

P43 Day 0

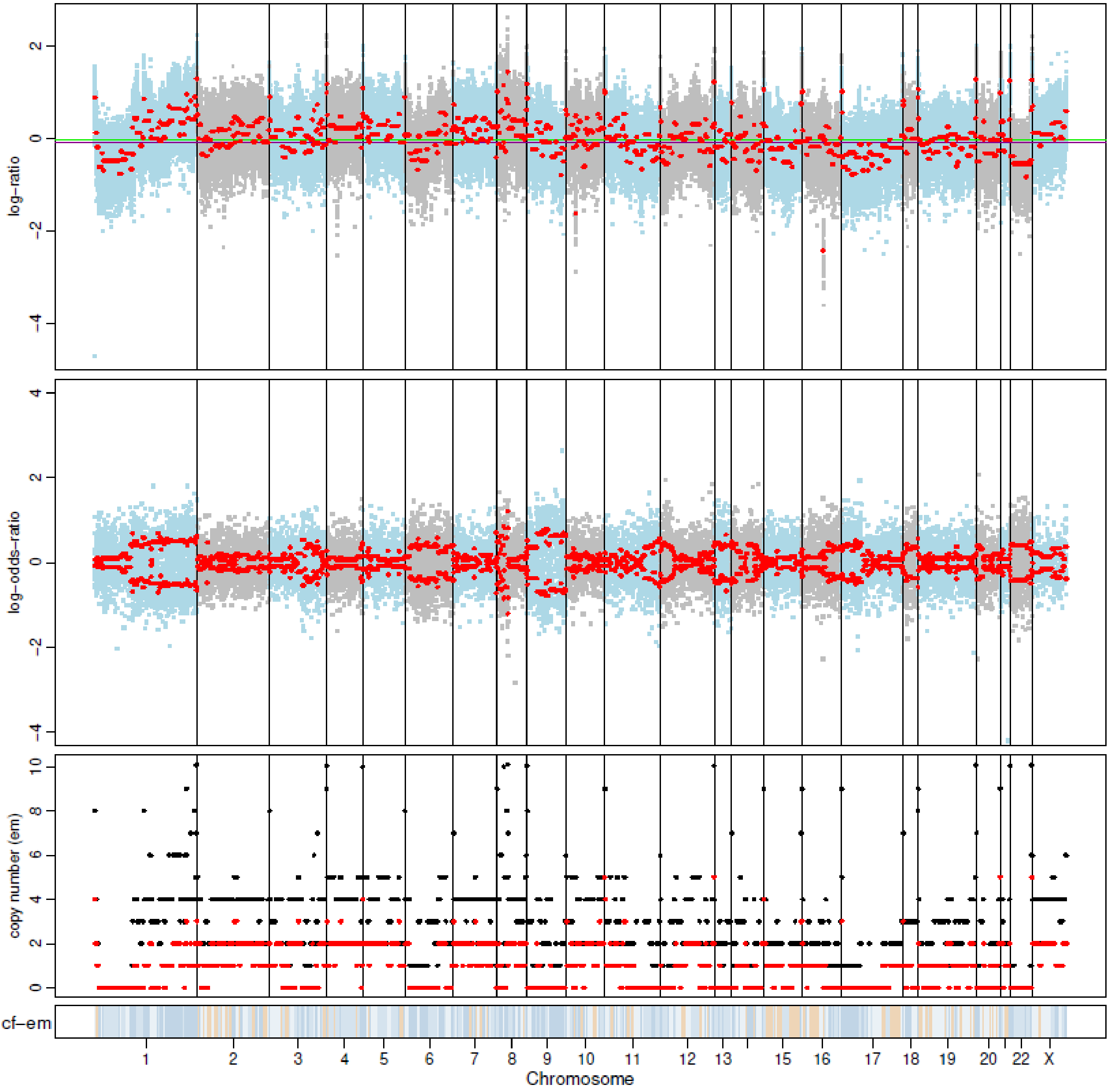


P43 Day 180

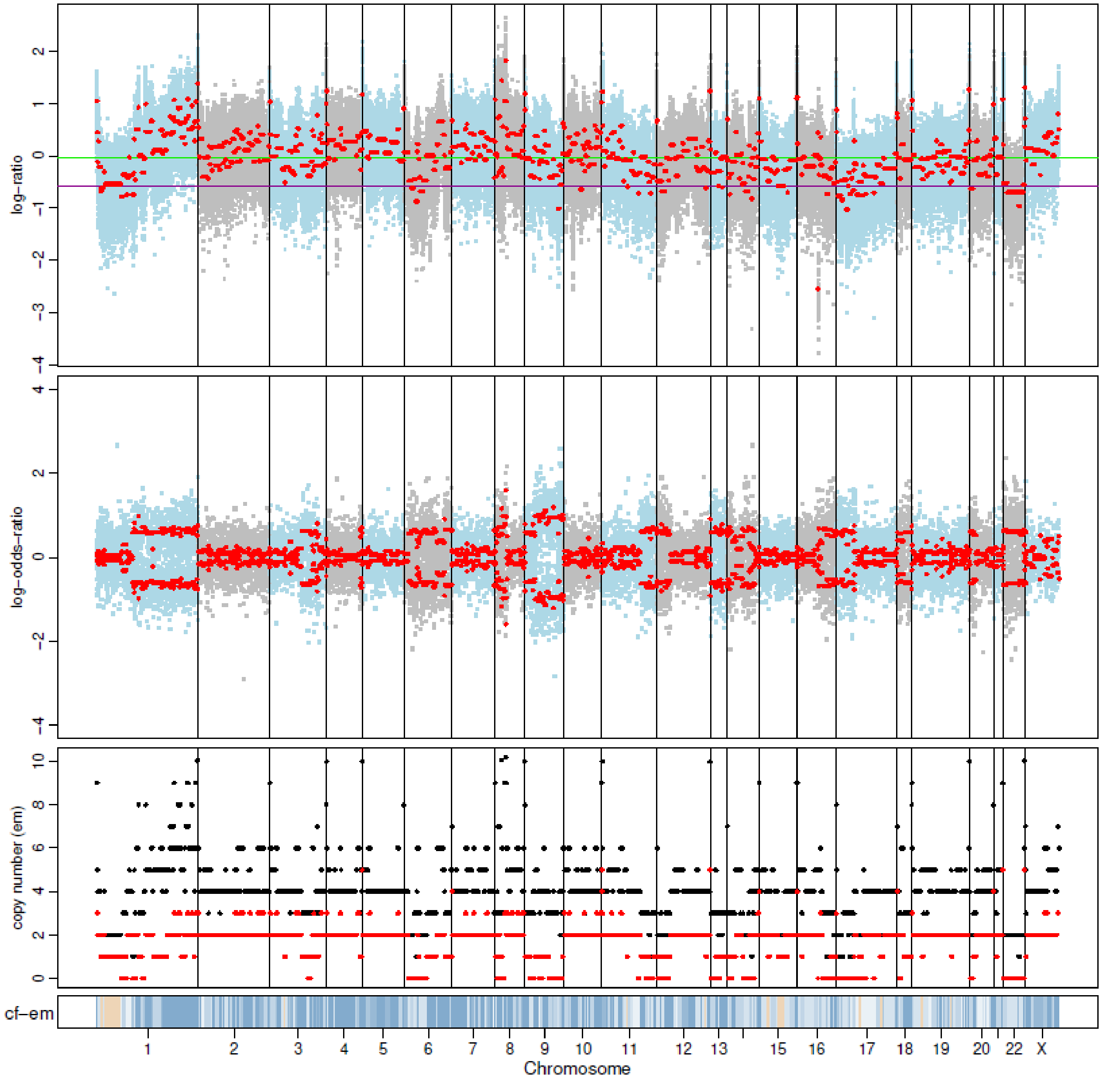

P44 Day 0

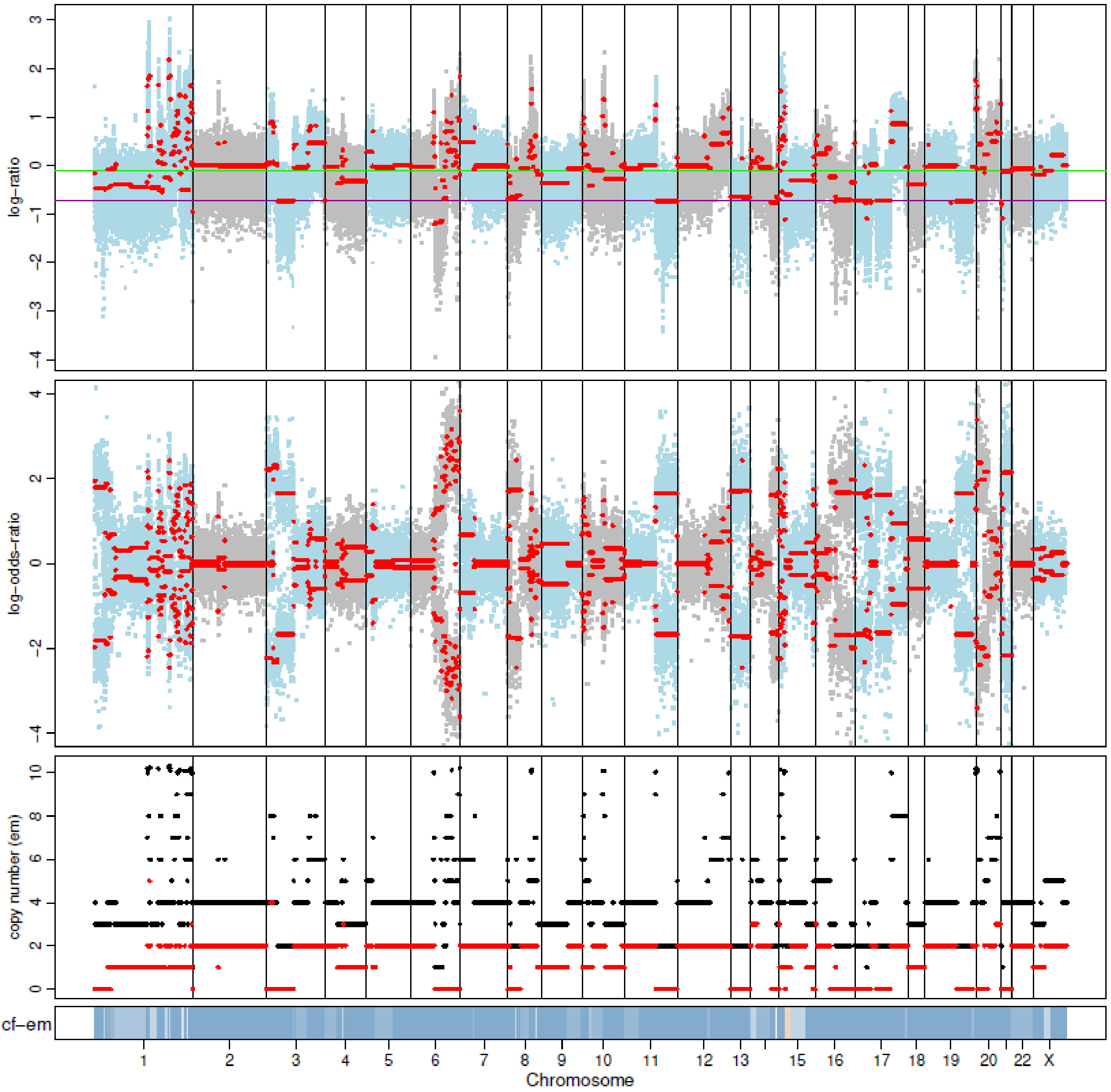


P44 Day 14

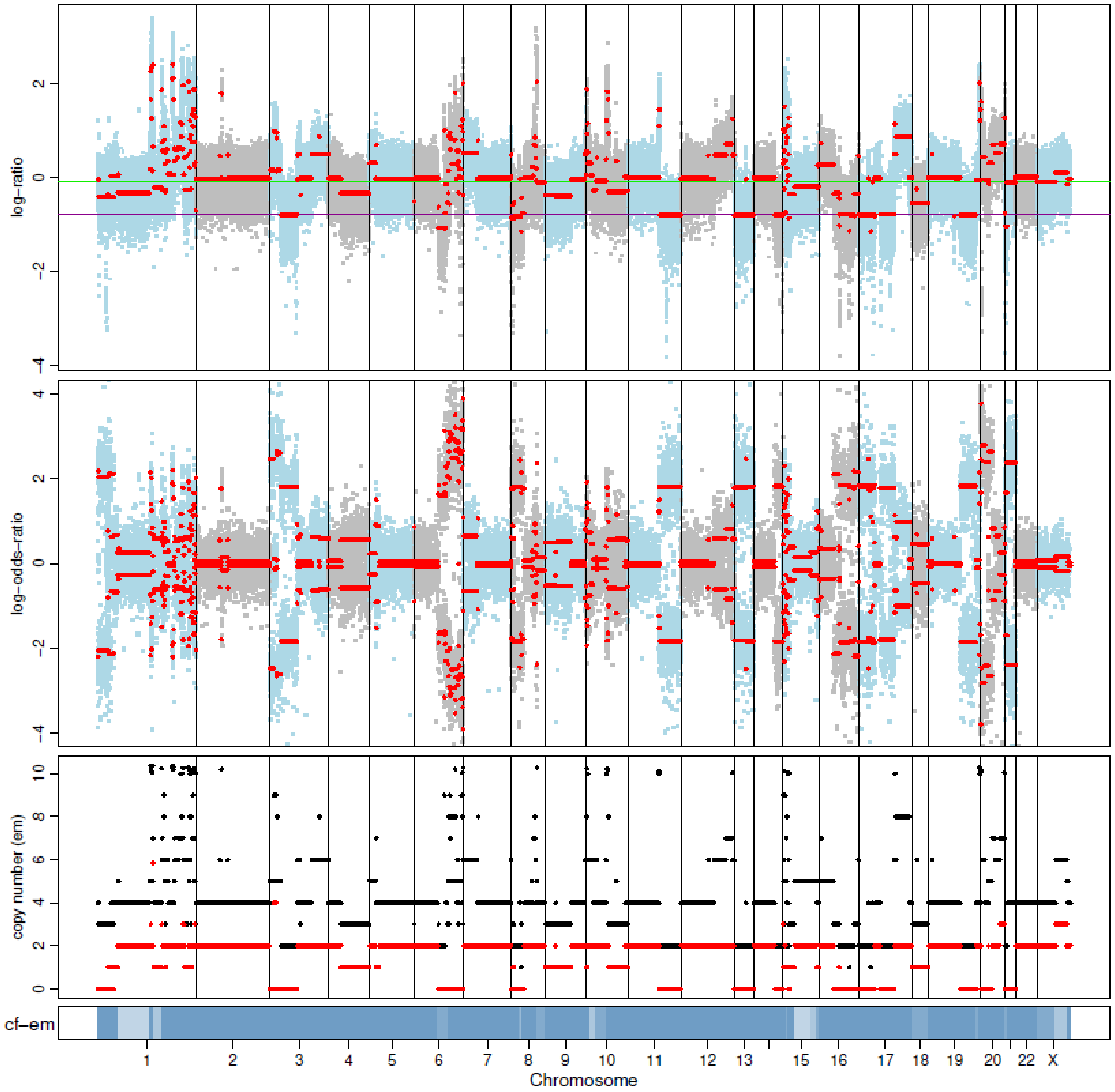

P45 Day 0

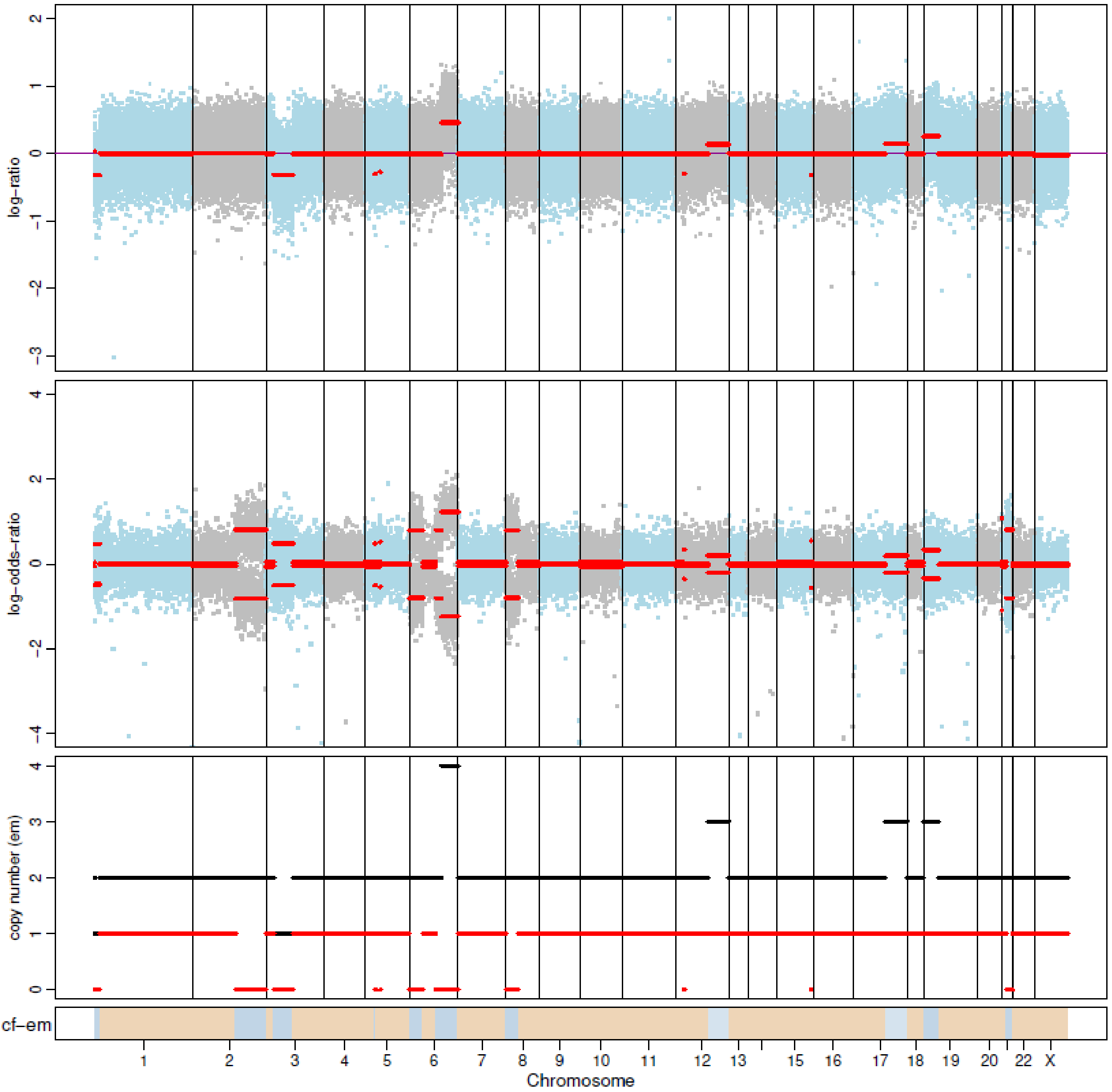


P45 Day 14

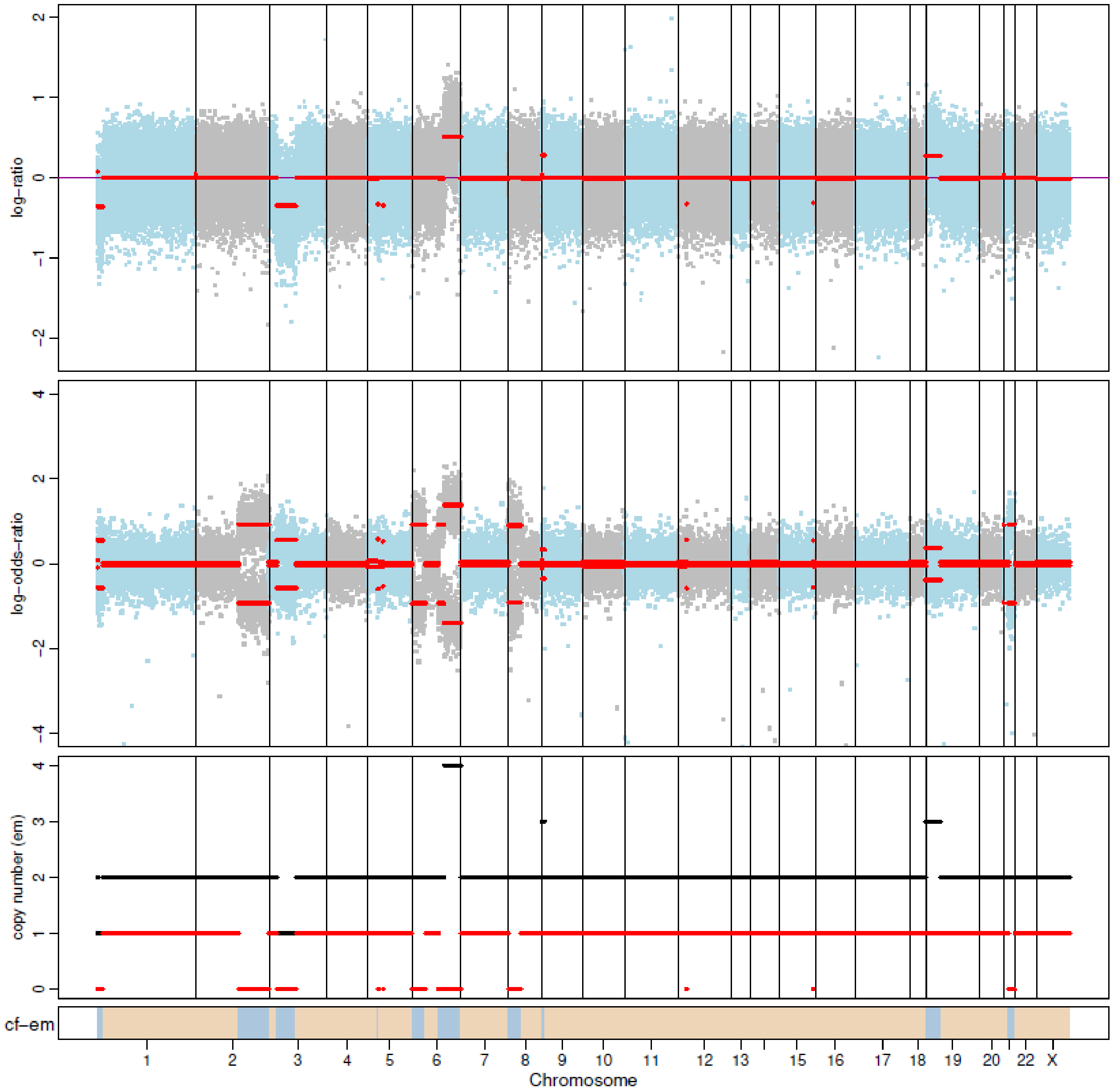

P46 Day 0

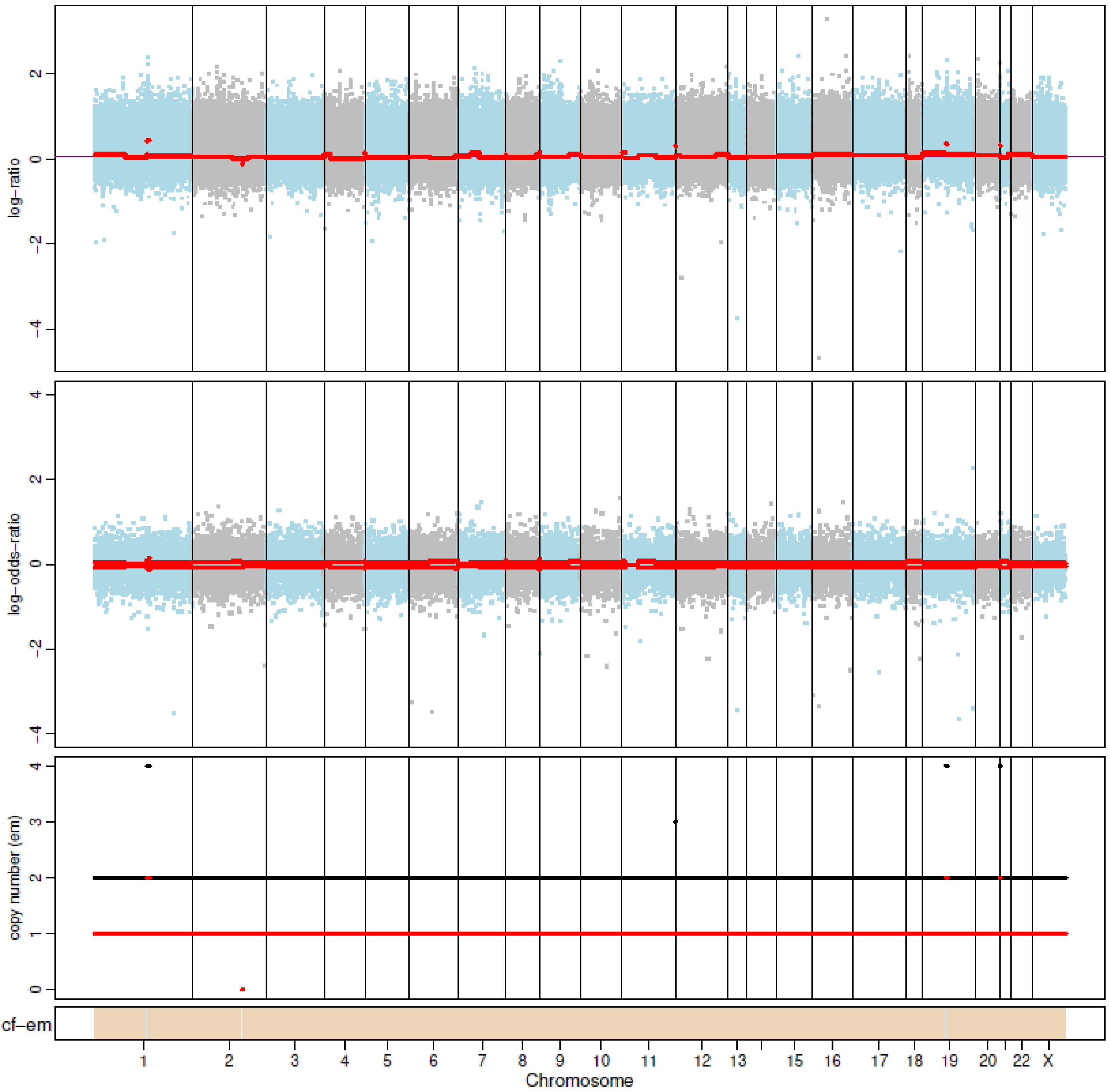


P46 Day 14

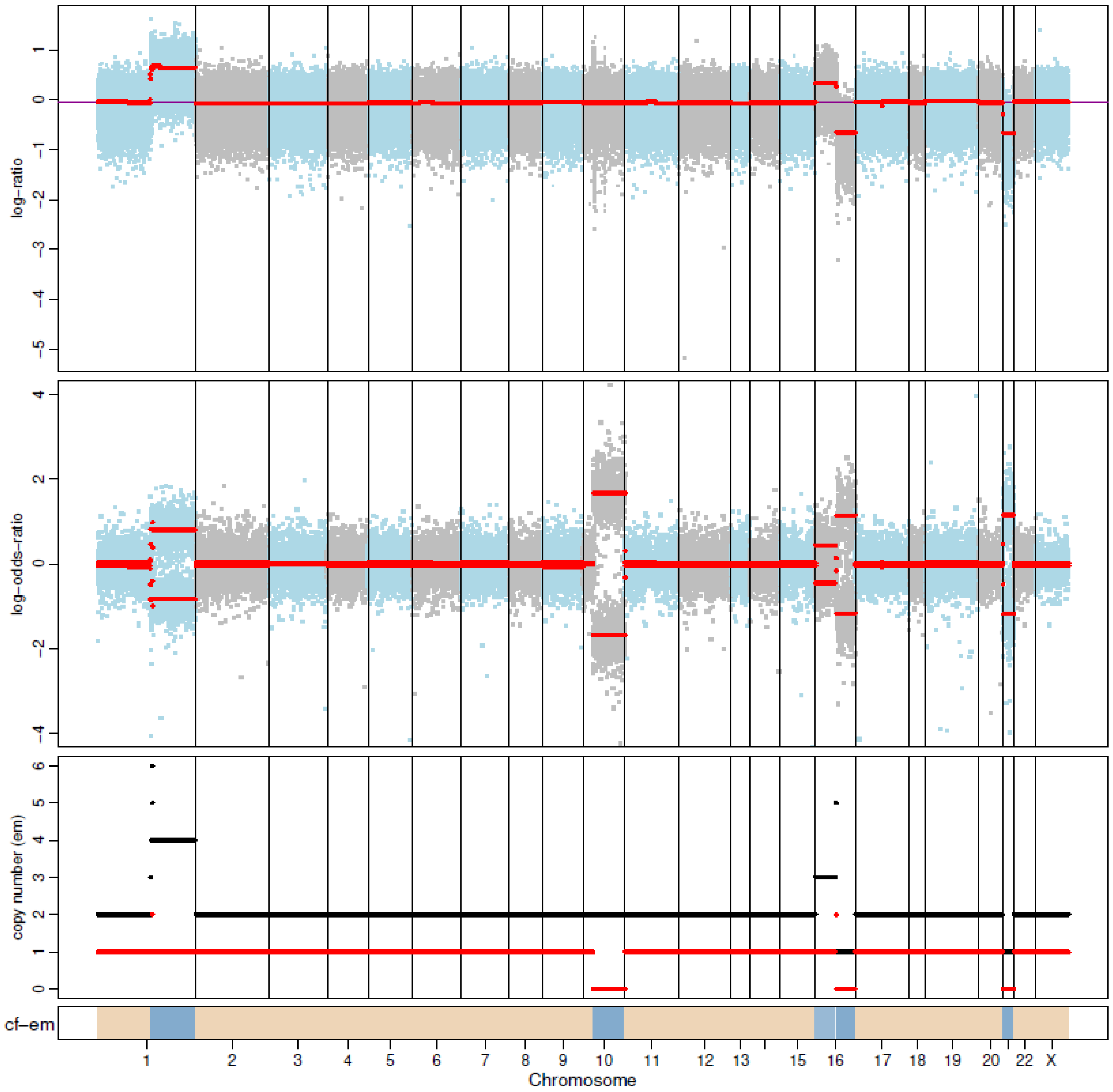
