## Supplementary Dataset 2 for "Convergent evolution of resistance pathways during early stage breast cancer treatment with combination cell cycle (CDK) and endocrine inhibitors"

**Supplementary Dataset 2. Single cell gene copy number profiles predicted by inferCNV in patient tumor cells.** In each figure, gene copy number profile estimated by i6 HMM model or denoising method (final) were shown. Red colors representing gene gains and blue colors representing gene losses. Single cells were clustered based on gene copy number profiles. Timepoints of patient samples (Day 0, 14, and 180) were annotated for each cell as gray, brown and green colors. Gene copy number profile were not shown for patients without sufficient number of tumor cells (P17, P30, and P42).

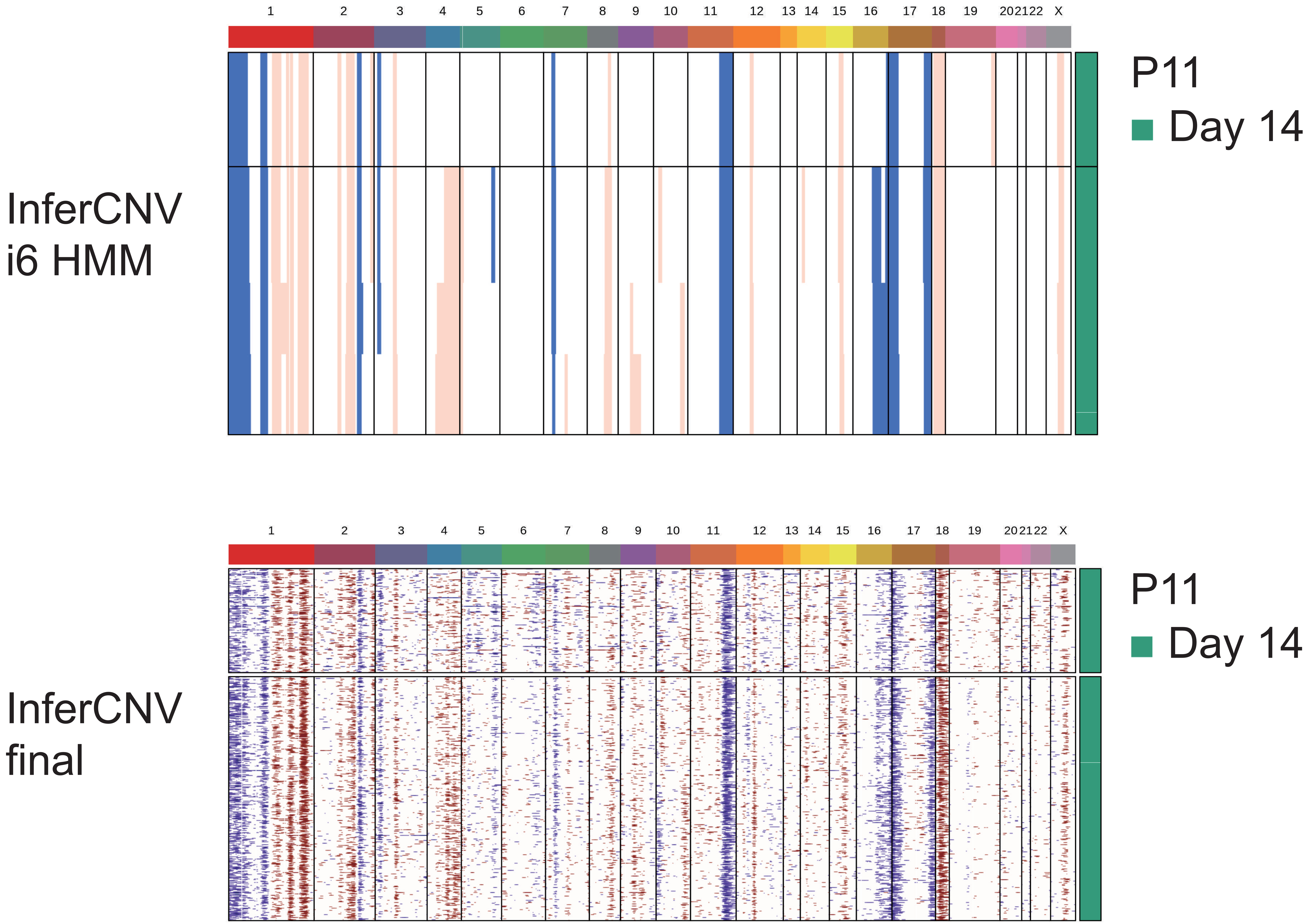
X
P11
Day 14
InferCNV
i6 HMM
InferCNV
final

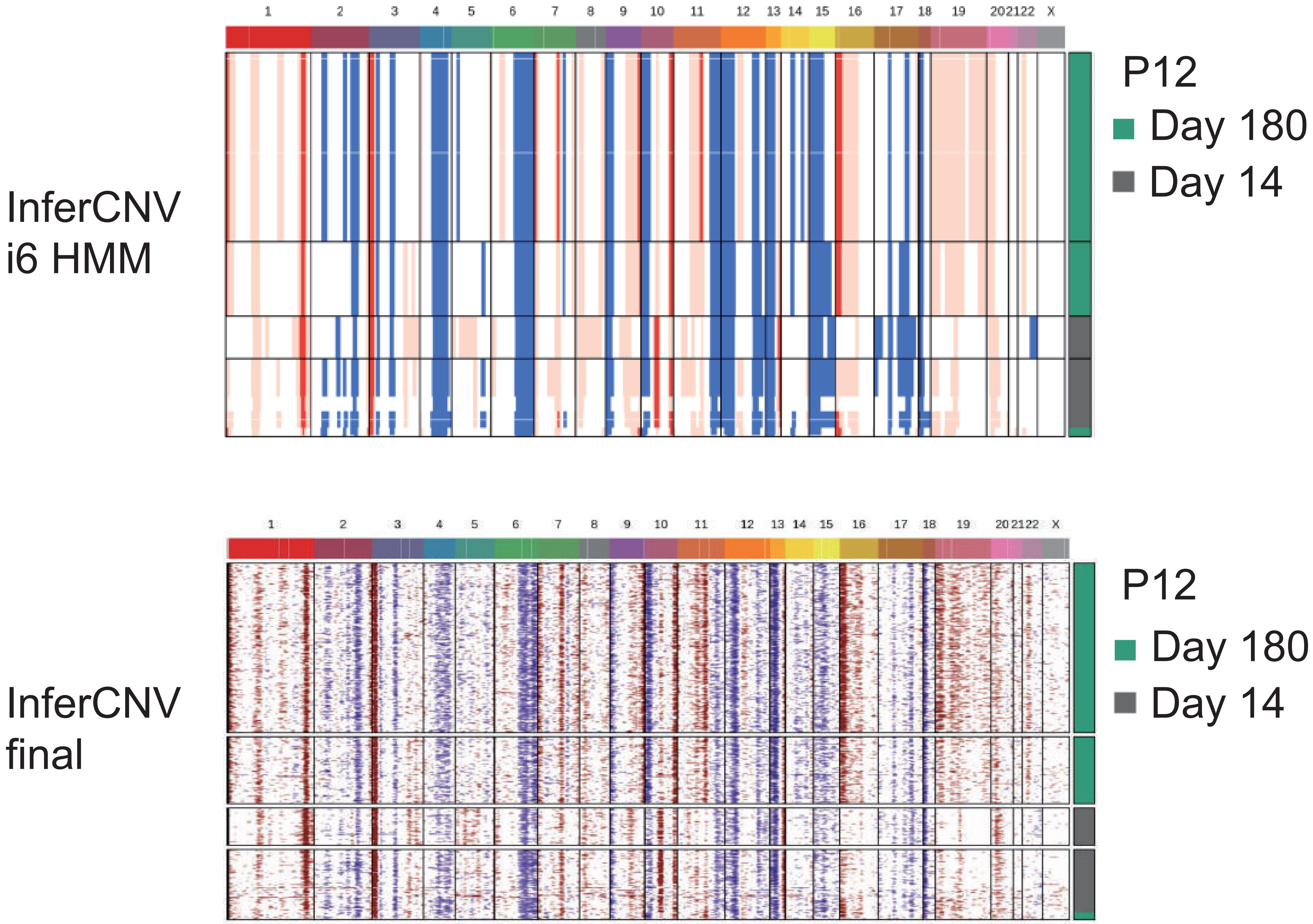
InferCNV
i6 HMM
P12
Day 180
Day 14
1 2 3 4 5 6 7 8 9 10 11 12 13 14 15 16 17 18 19 20 21 22 X
InferCNV
final
P12
Day 180
Day 14
1 2 3 4 5 6 7 8 9 10 11 12 13 14 15 16 17 18 19 20 21 22 X

InferCNV
i6 HMM

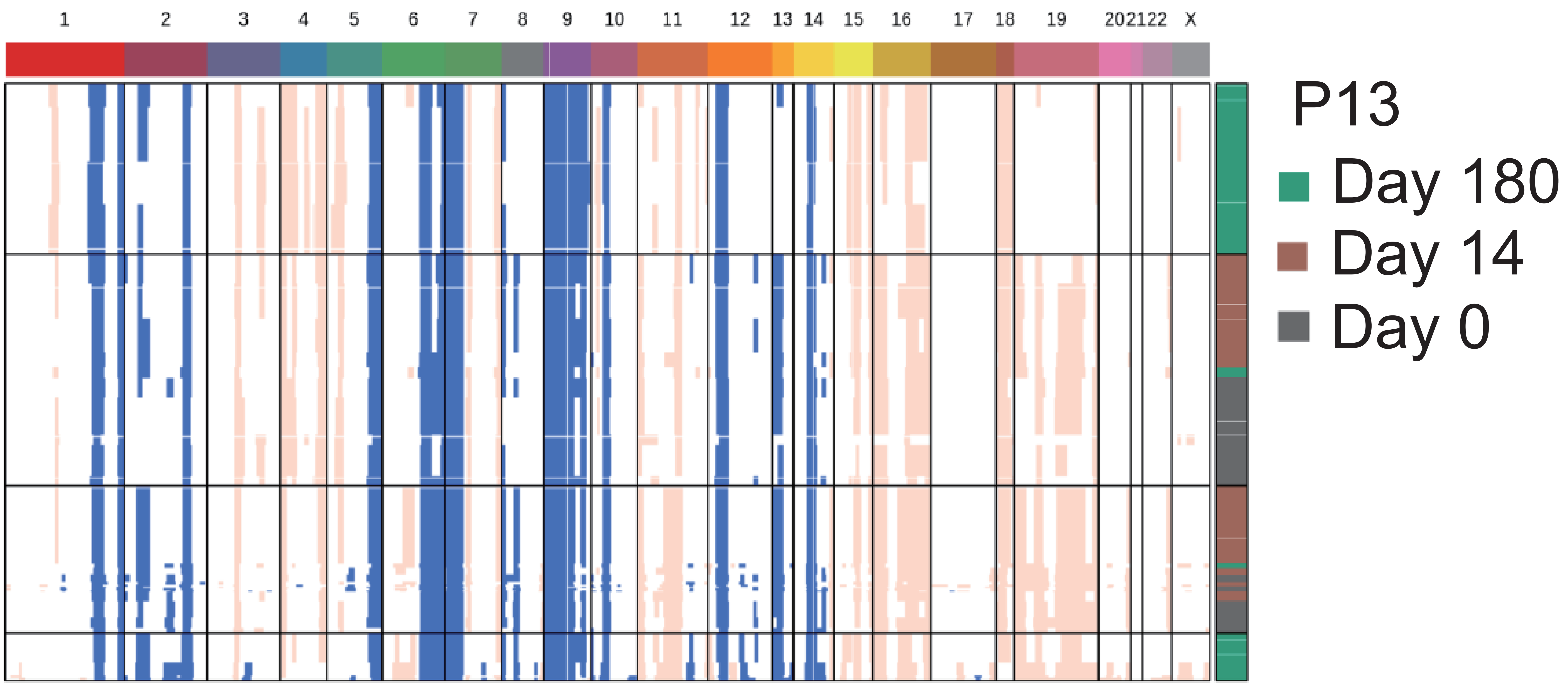

InferCNV
final

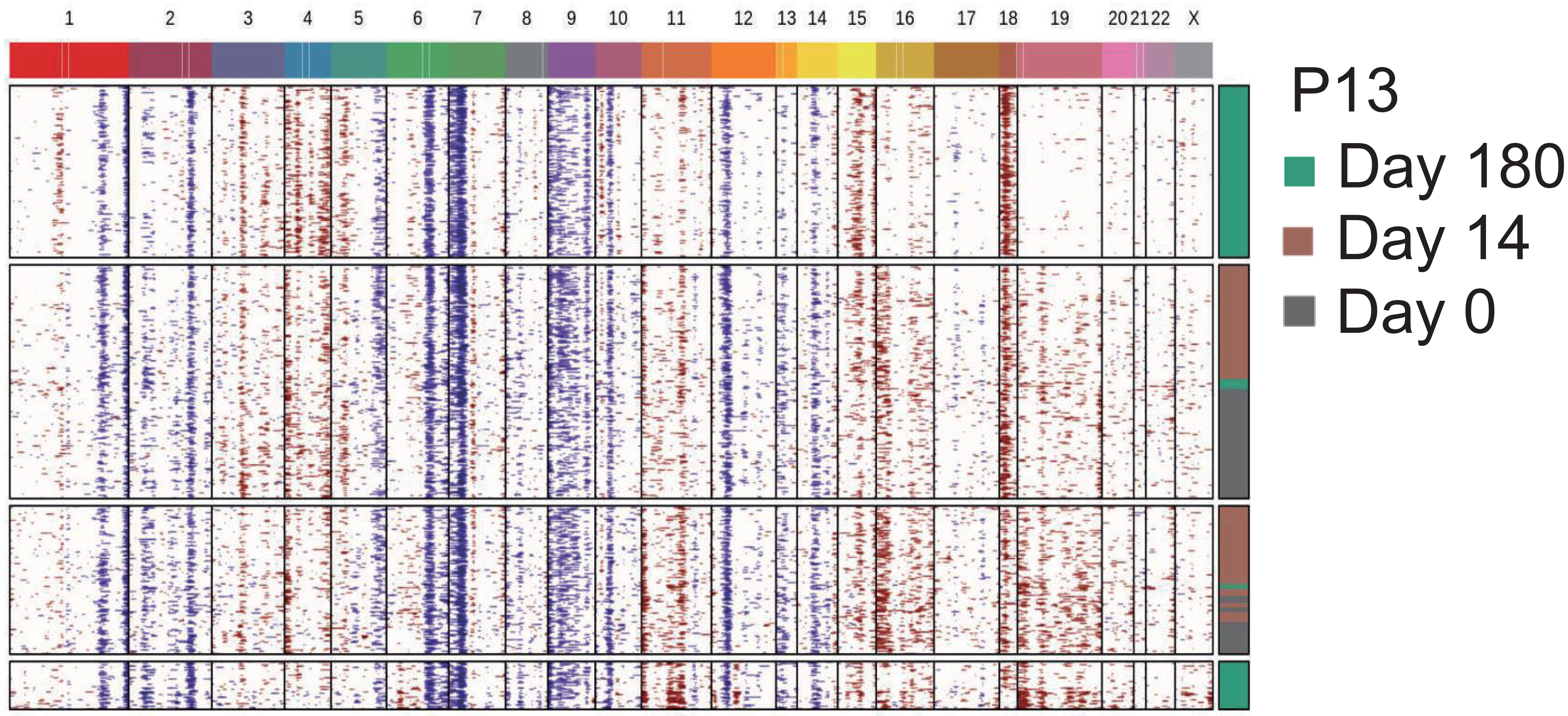

InferCNV
i6 HMM

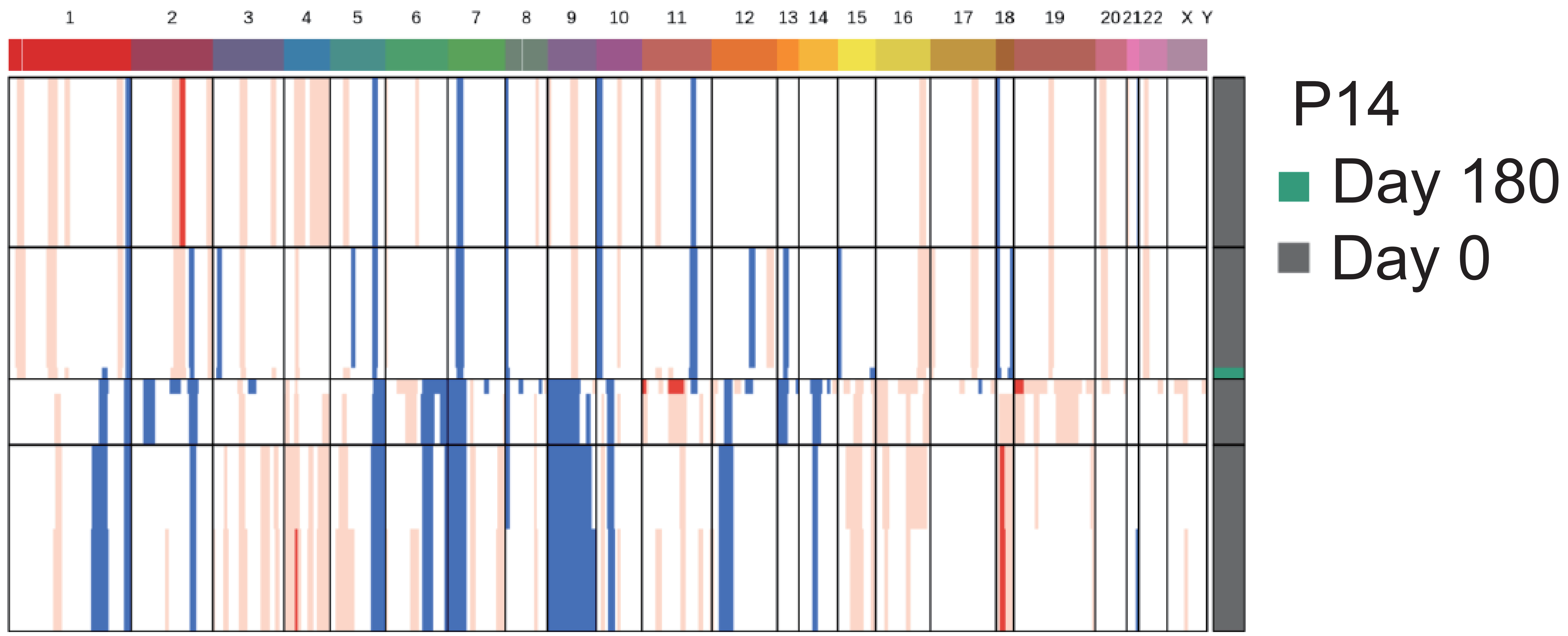

InferCNV
final

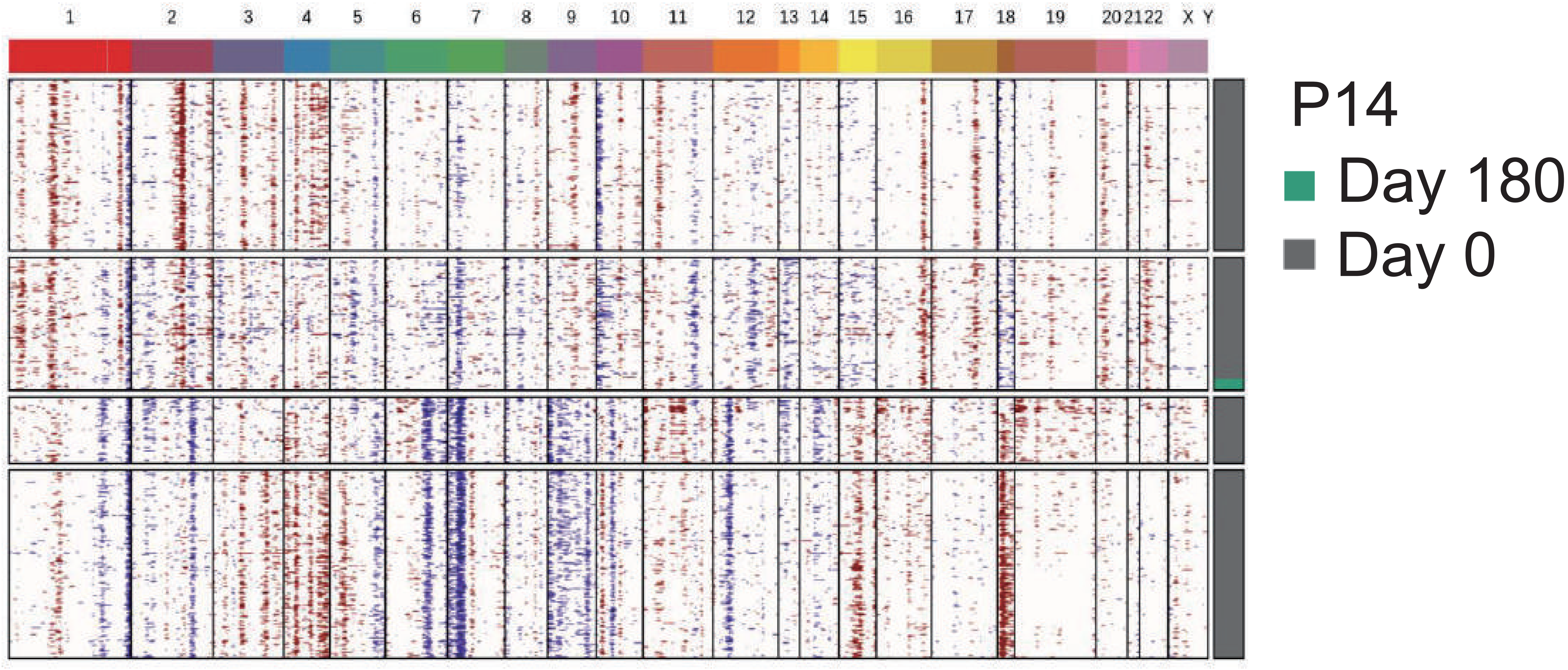

InferCNV
i6 HMM

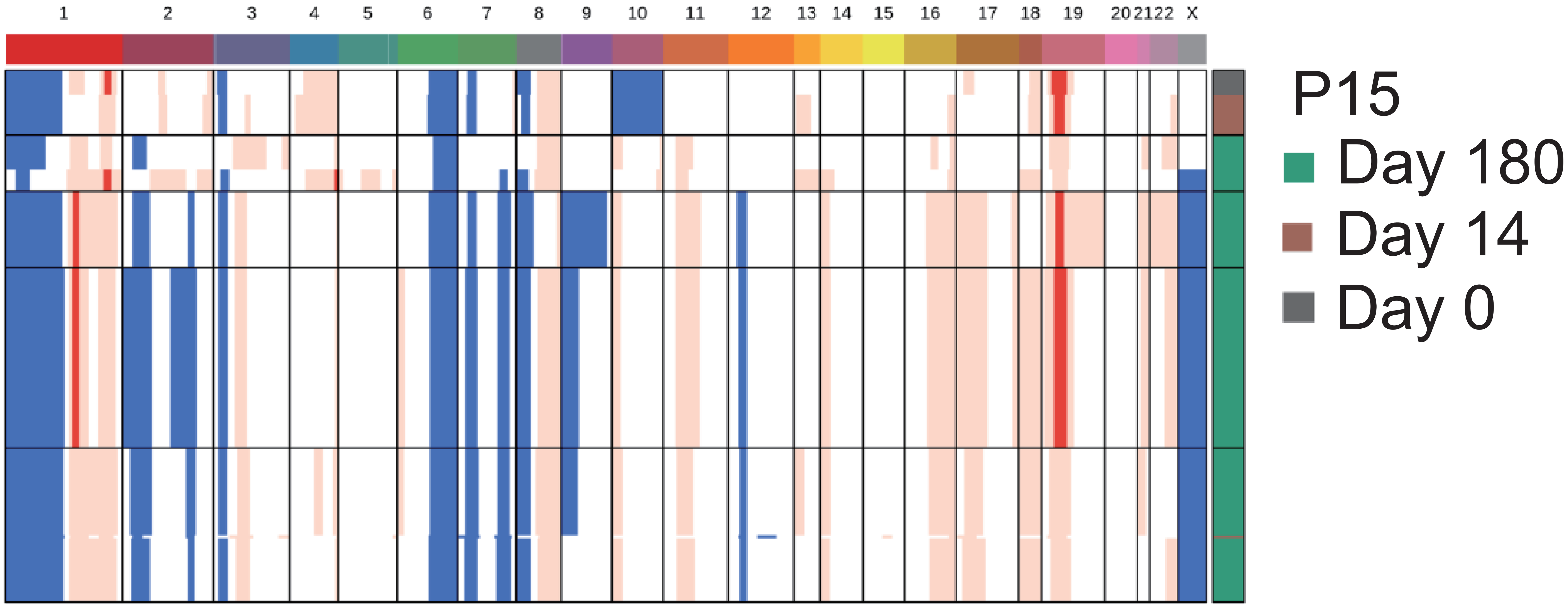

InferCNV
final

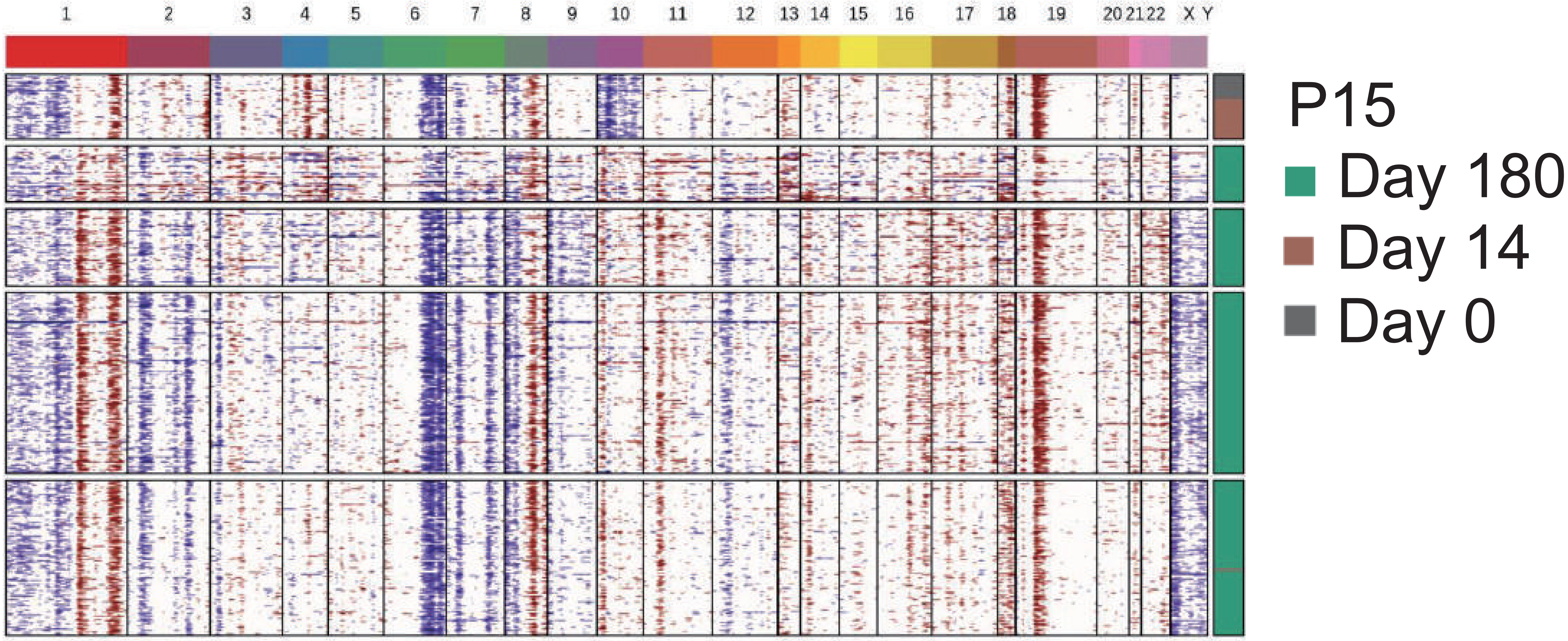

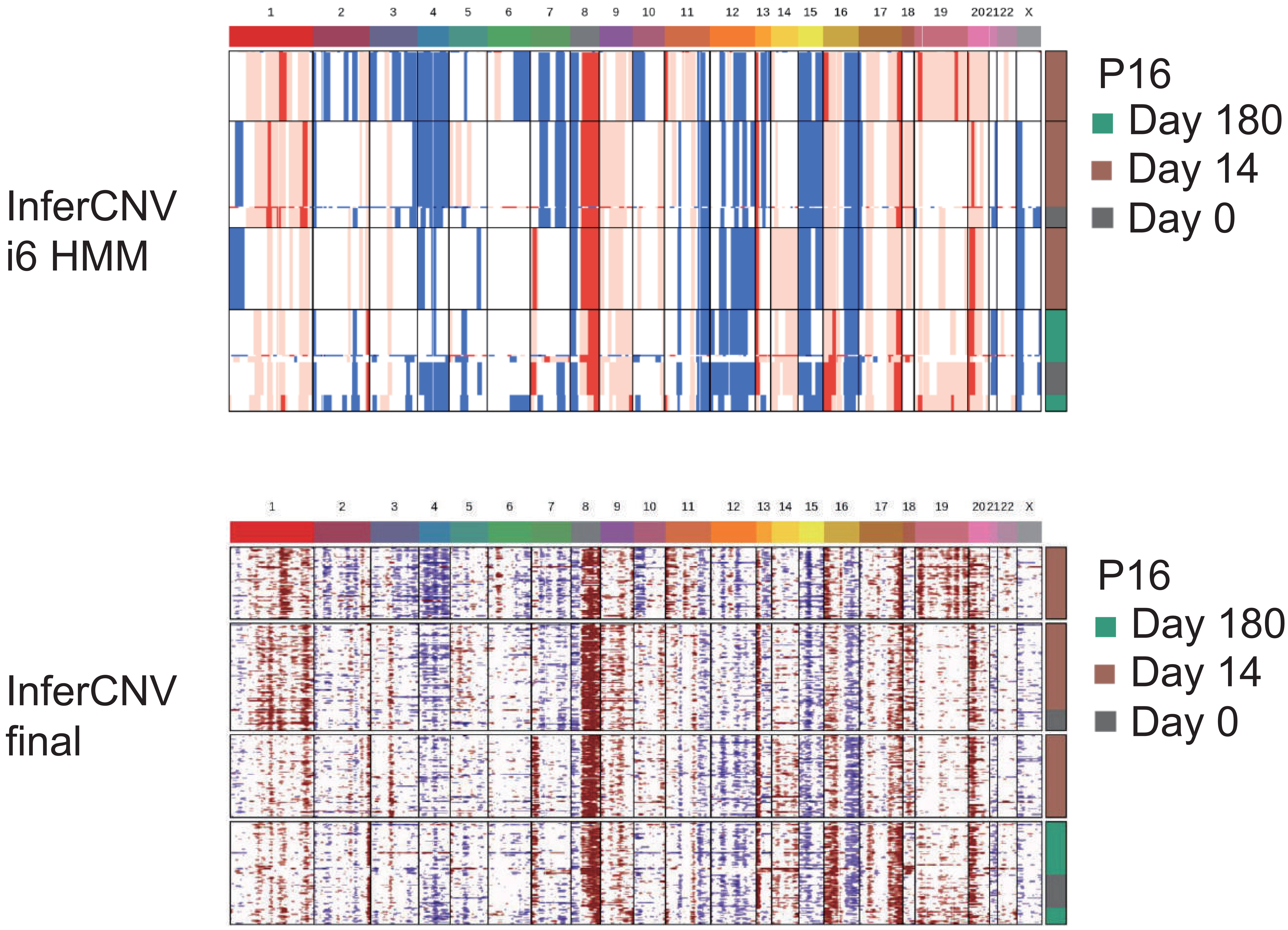
InferCNV
i6 HMM
1 2 3 4 5 6 7 8 9 10 11 12 13 14 15 16 17 18 19 20 21 22 X
P16
Day 180
Day 14
Day 0
InferCNV
final
1 2 3 4 5 6 7 8 9 10 11 12 13 14 15 16 17 18 19 20 21 22 X
P16
Day 180
Day 14
Day 0

InferCNV
i6 HMM

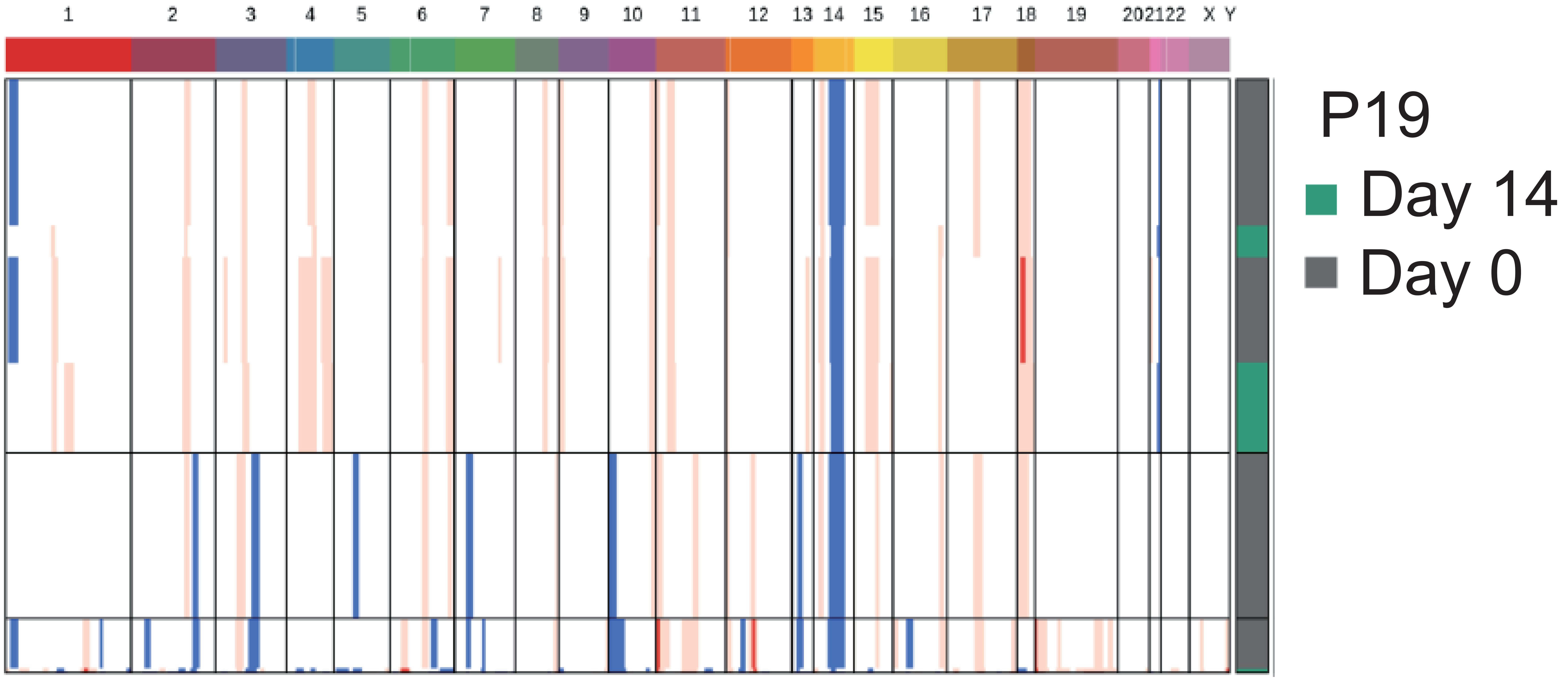

InferCNV
final

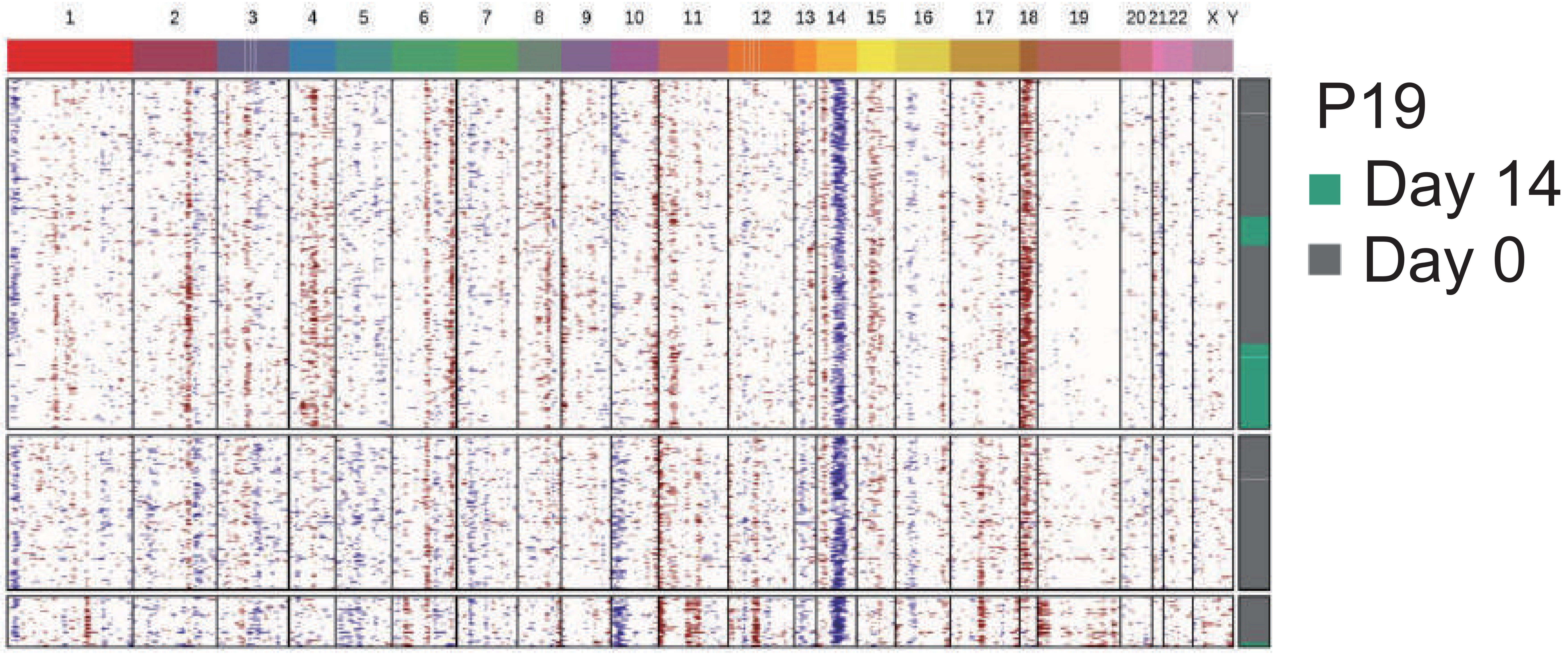

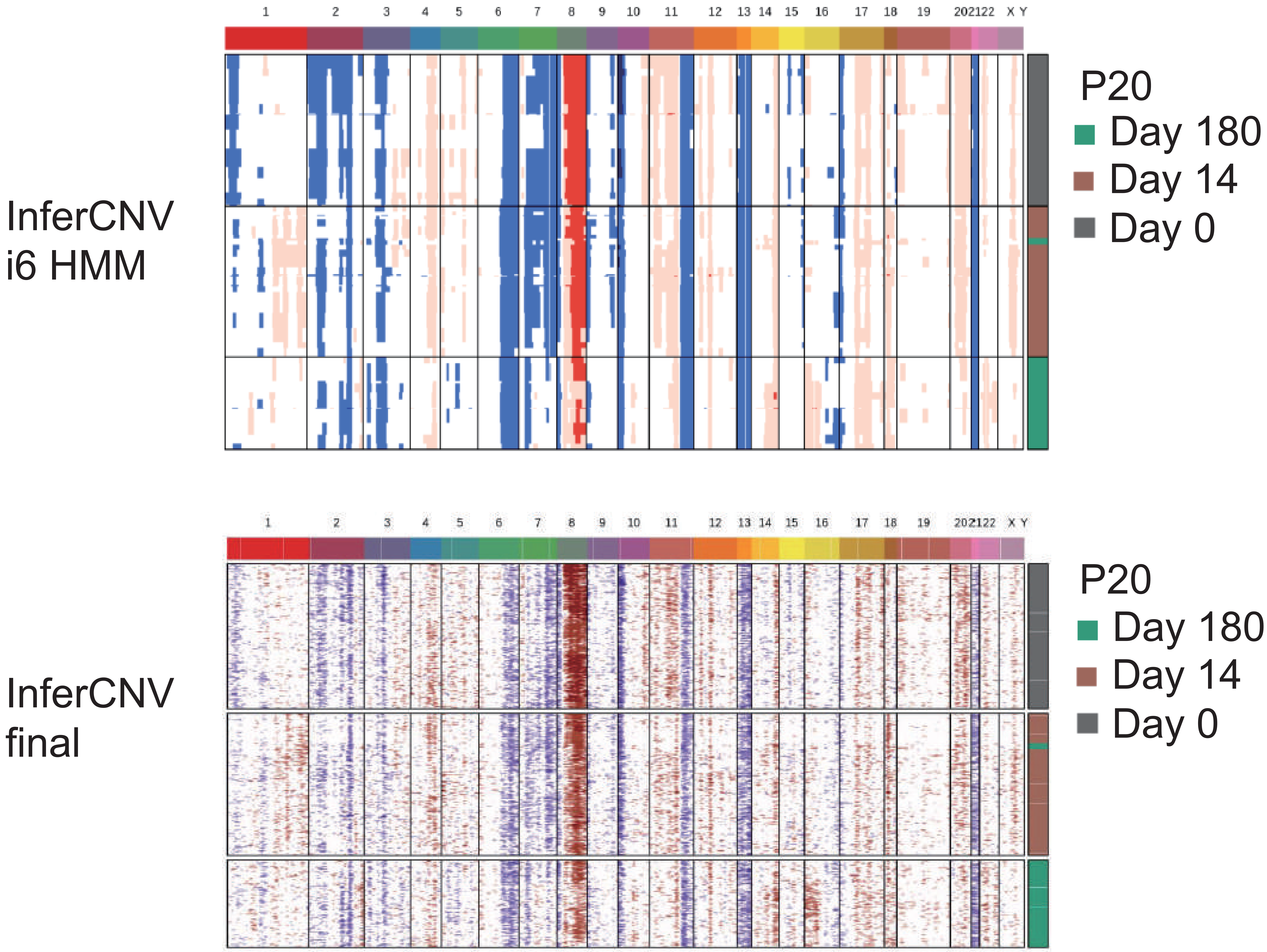

InferCNV
i6 HMM
1 2 3 4 5 6 7 8 9 10 11 12 13 14 15 16 17 18 19 20 21 22 X Y
P20
Day 180
Day 14
Day 0
InferCNV
final
1 2 3 4 5 6 7 8 9 10 11 12 13 14 15 16 17 18 19 20 21 22 X Y
P20
Day 180
Day 14
Day 0

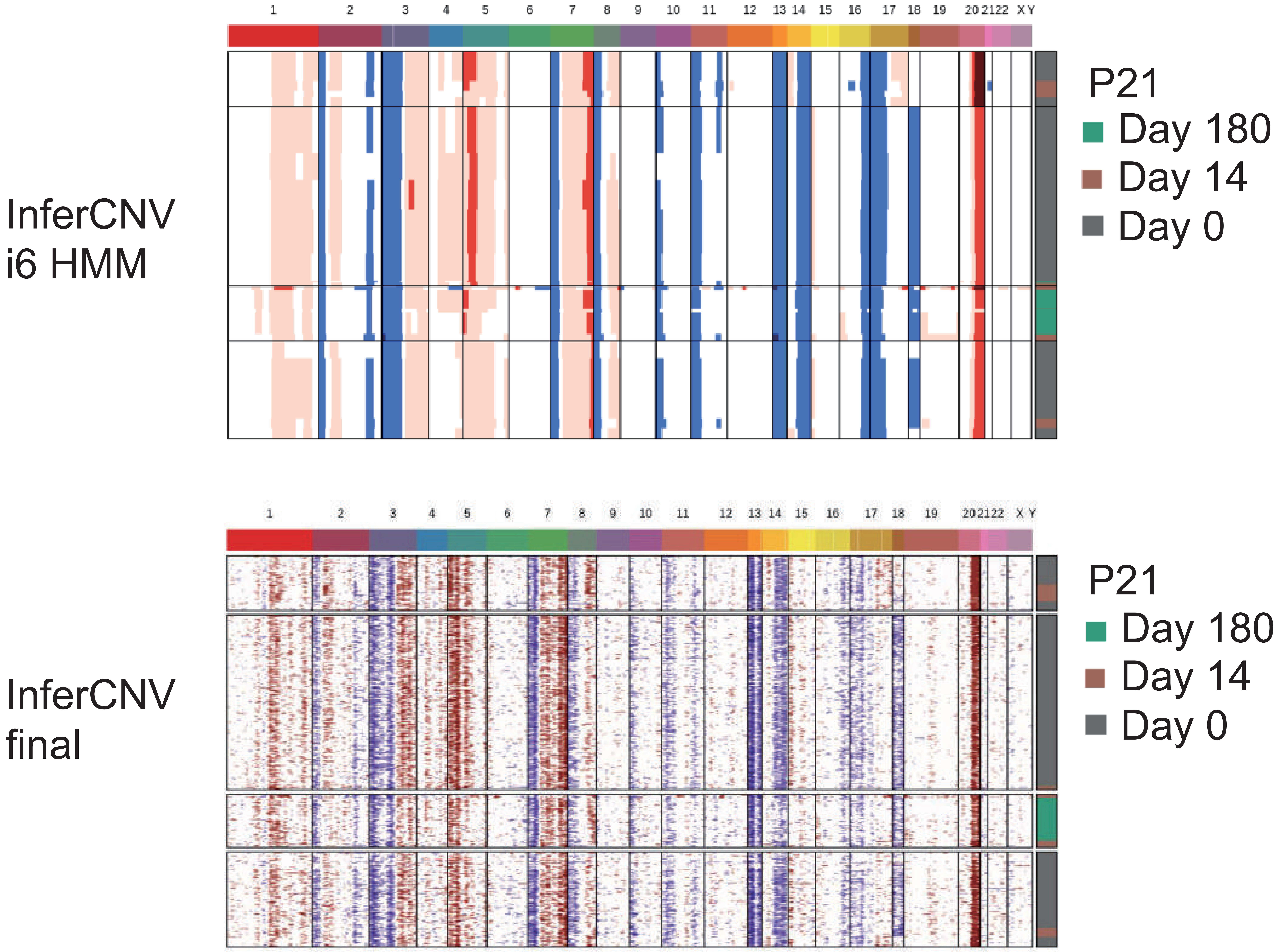
InferCNV
i6 HMM
1 2 3 4 5 6 7 8 9 10 11 12 13 14 15 16 17 18 19 20 21 22 X Y
P21
Day 180
Day 14
Day 0
InferCNV
final
1 2 3 4 5 6 7 8 9 10 11 12 13 14 15 16 17 18 19 20 21 22 X Y
P21
Day 180
Day 14
Day 0

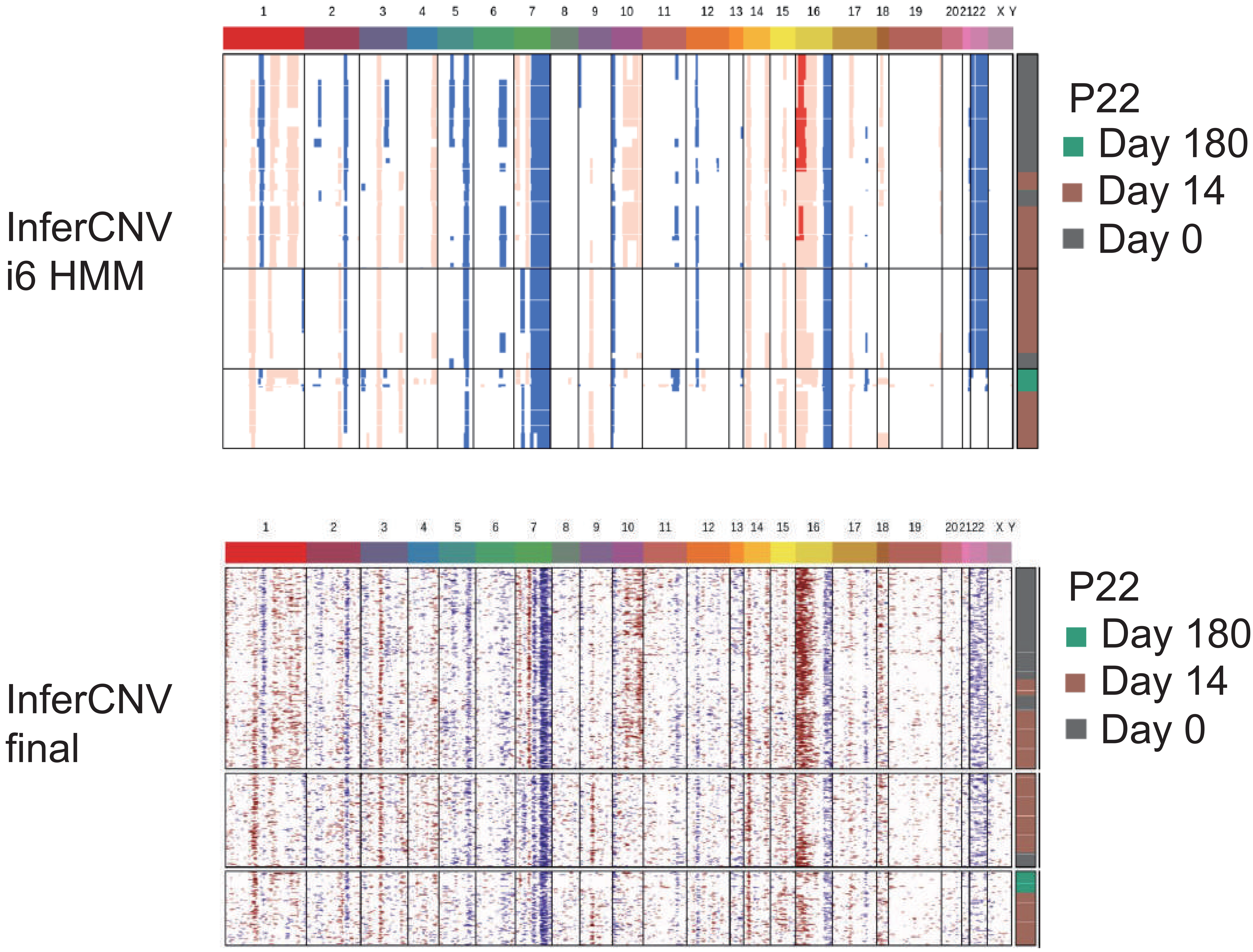
InferCNV
i6 HMM
InferCNV
final
1 2 3 4 5 6 7 8 9 10 11 12 13 14 15 16 17 18 19 20 21 22 X Y
P22
Day 180
Day 14
Day 0

InferCNV
i6 HMM

InferCNV
final

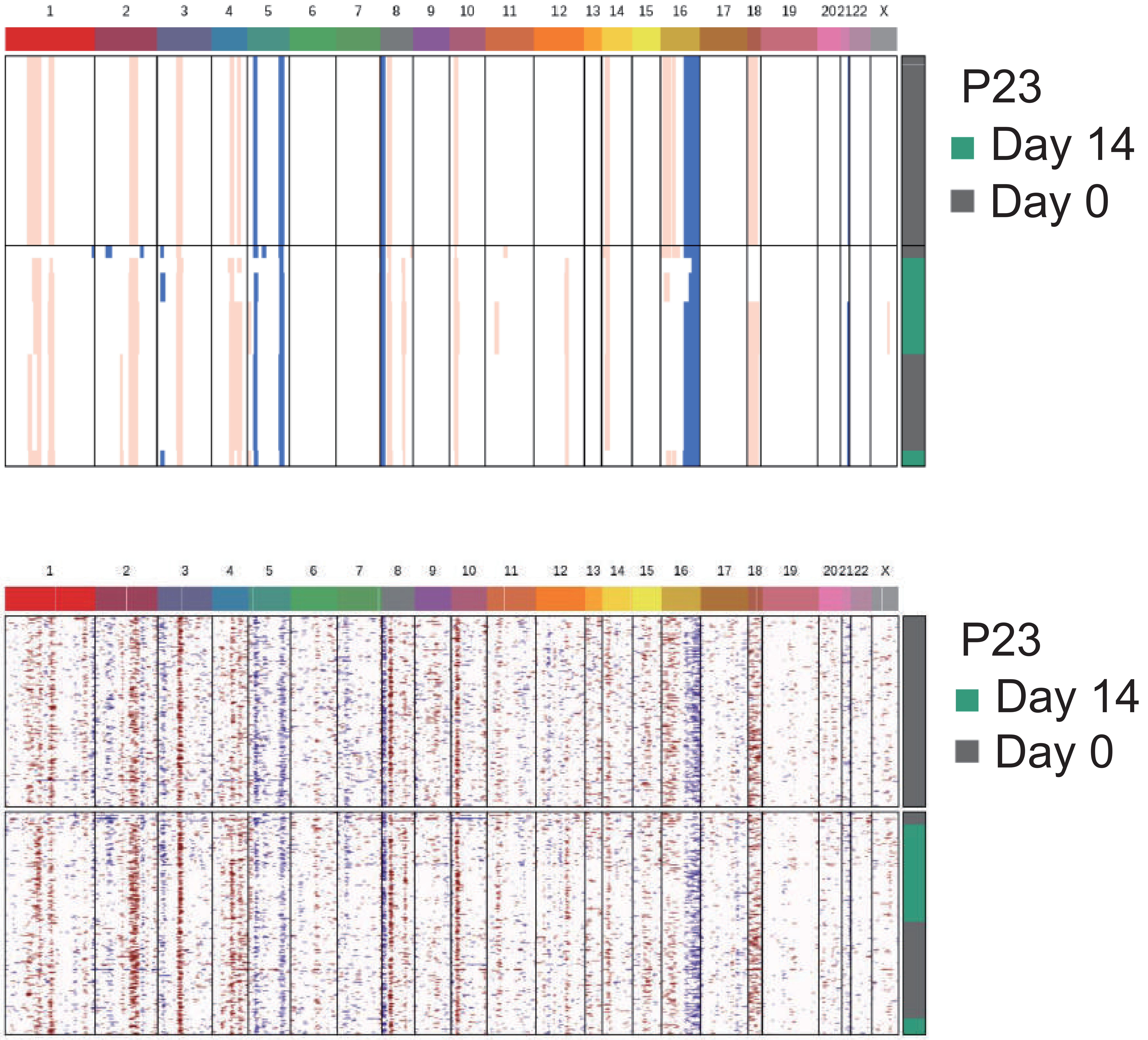

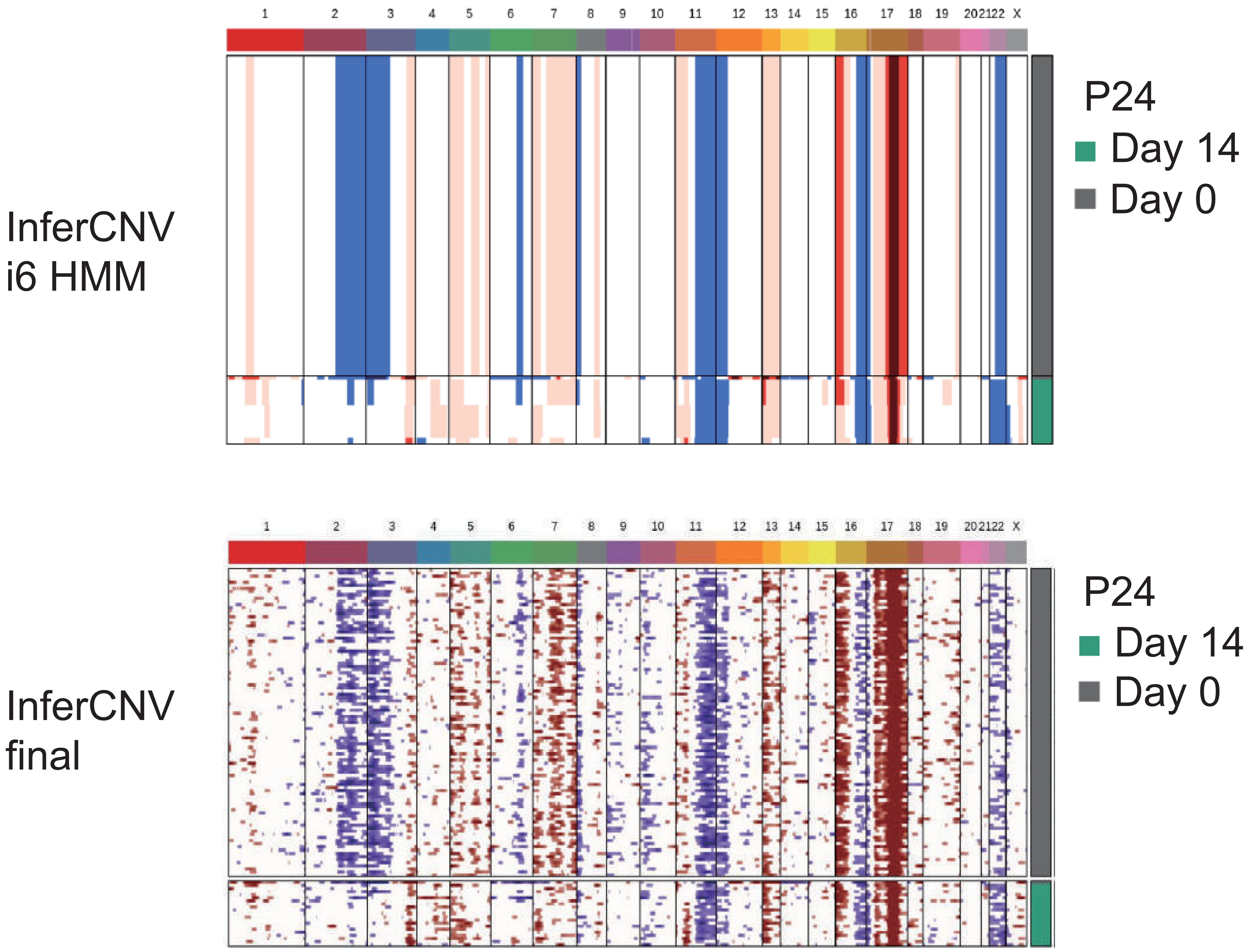
InferCNV
i6 HMM
1 2 3 4 5 6 7 8 9 10 11 12 13 14 15 16 17 18 19 20 21 22 X
P24
Day 14
Day 0
InferCNV
final
1 2 3 4 5 6 7 8 9 10 11 12 13 14 15 16 17 18 19 20 21 22 X
P24
Day 14
Day 0

InferCNV
i6 HMM

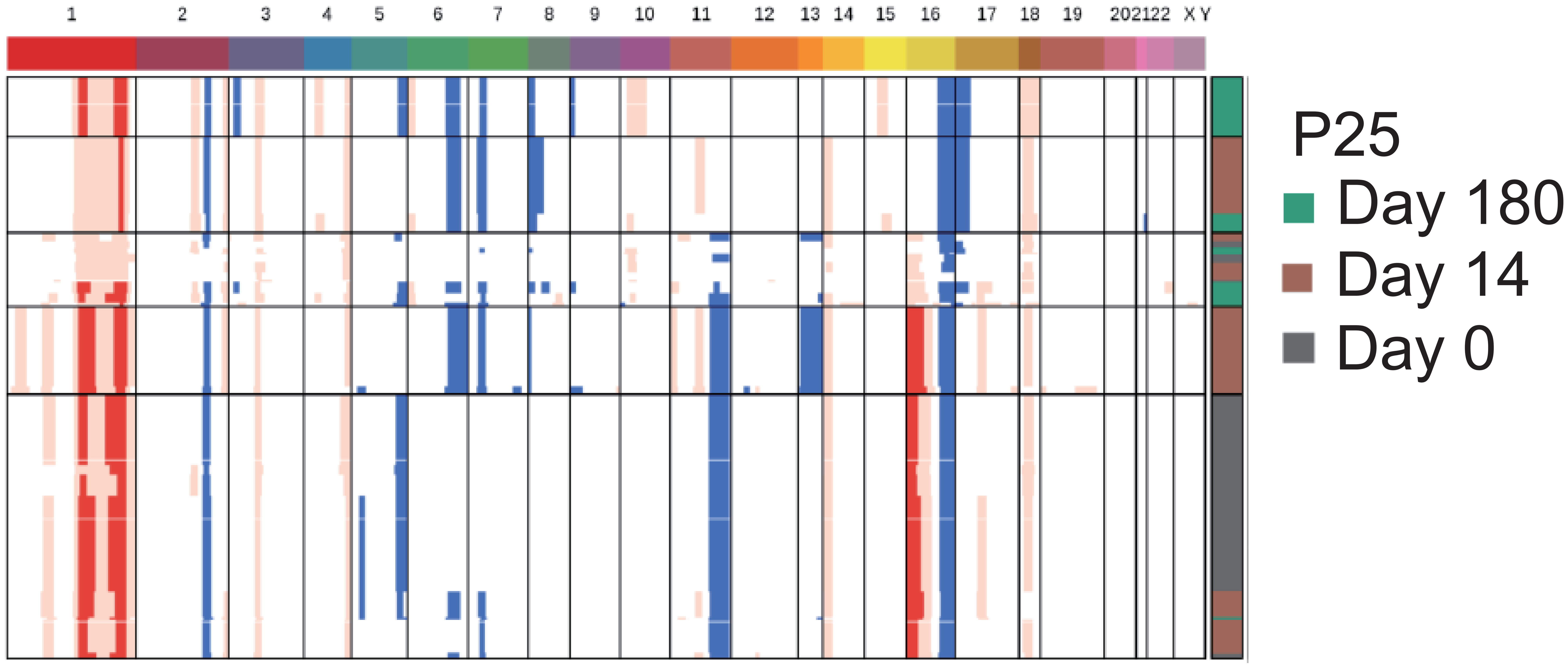

InferCNV
final

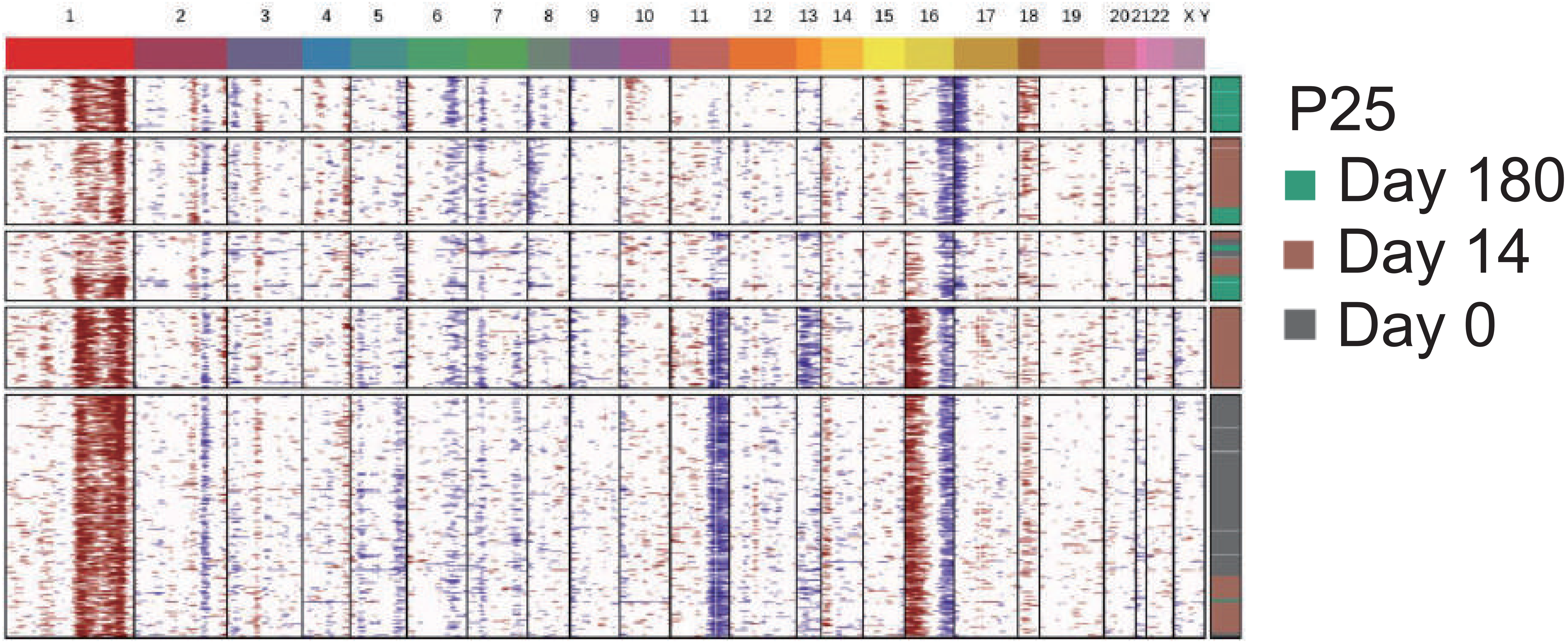

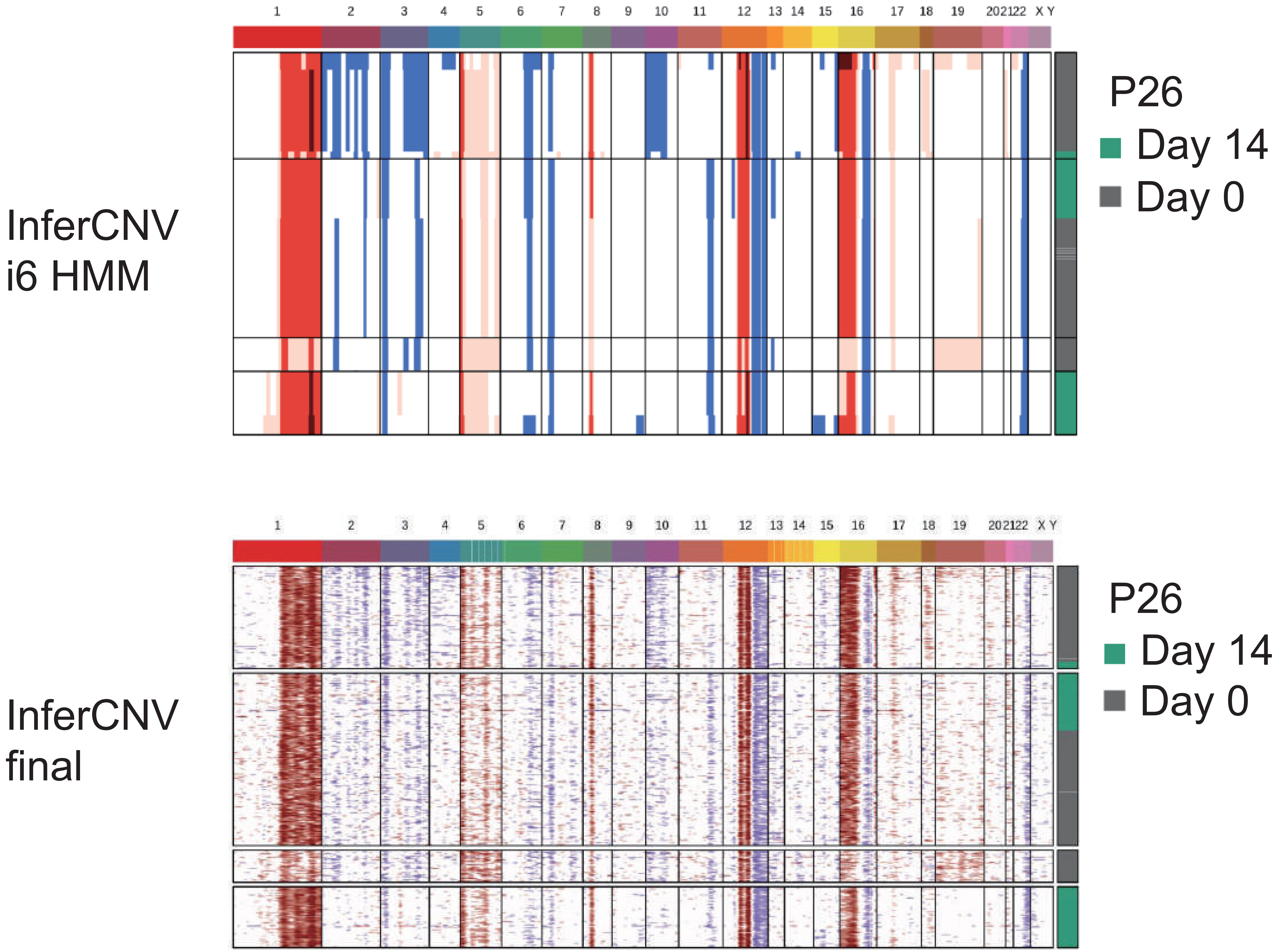
InferCNV
i6 HMM
1 2 3 4 5 6 7 8 9 10 11 12 13 14 15 16 17 18 19 20 21 22 X Y
P26
Day 14
Day 0
InferCNV
final
1 2 3 4 5 6 7 8 9 10 11 12 13 14 15 16 17 18 19 20 21 22 X Y
P26
Day 14
Day 0

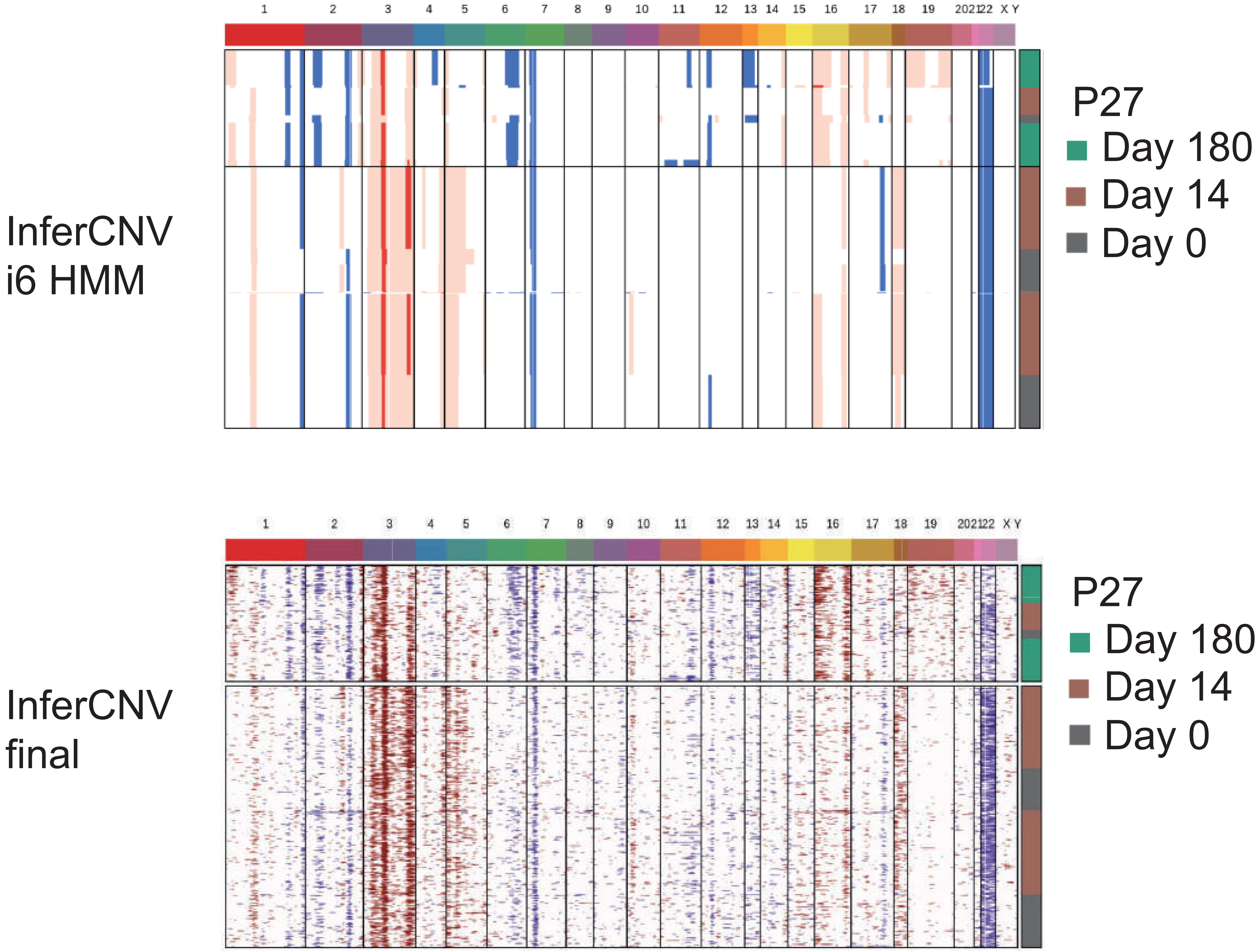
InferCNV
i6 HMM
P27
Day 180
Day 14
Day 0
1 2 3 4 5 6 7 8 9 10 11 12 13 14 15 16 17 18 19 20 21 22 X Y
InferCNV
final
P27
Day 180
Day 14
Day 0
1 2 3 4 5 6 7 8 9 10 11 12 13 14 15 16 17 18 19 20 21 22 X Y

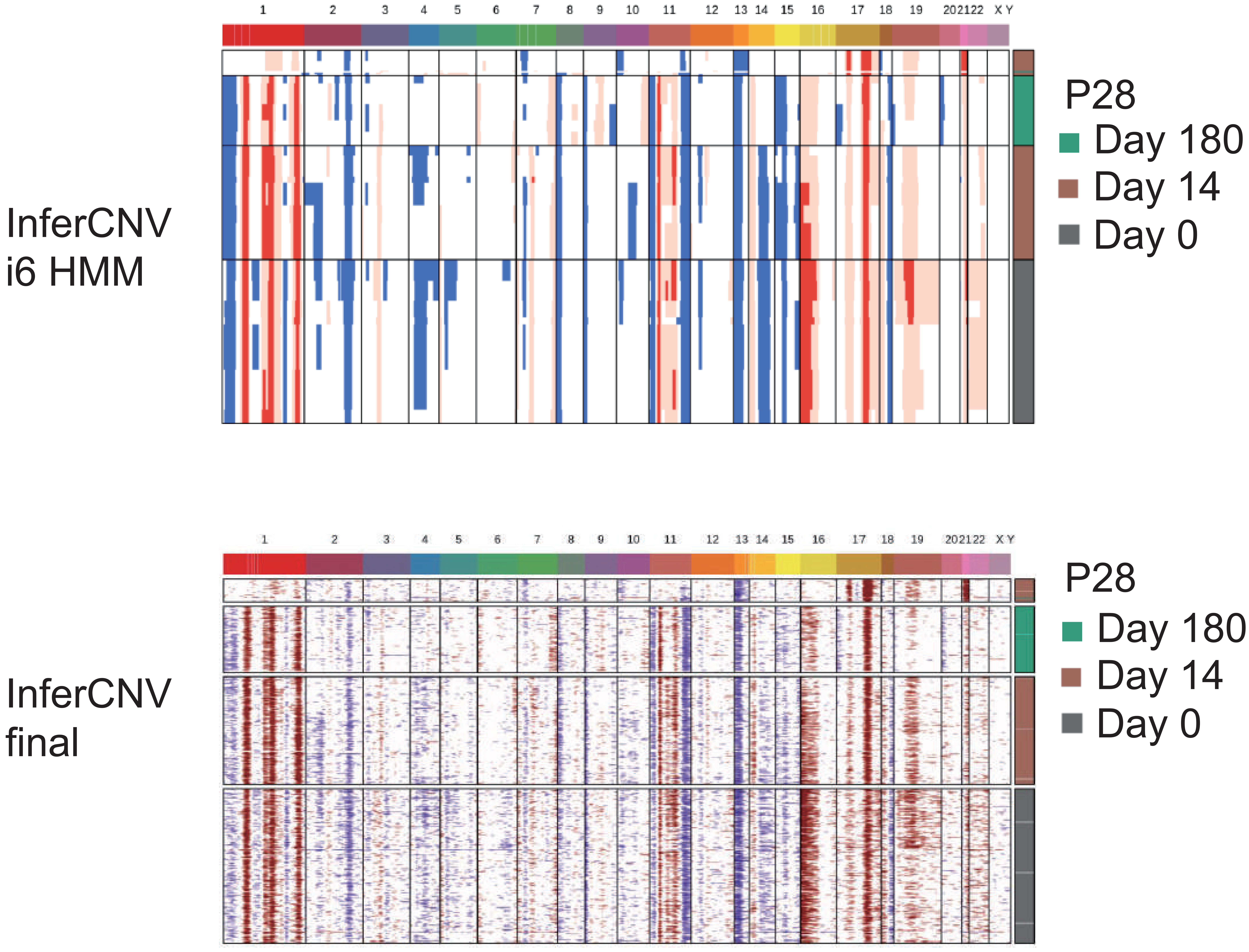

InferCNV
i6 HMM
1 2 3 4 5 6 7 8 9 10 11 12 13 14 15 16 17 18 19 20 21 22 X Y
P28
Day 180
Day 14
Day 0
InferCNV
final
1 2 3 4 5 6 7 8 9 10 11 12 13 14 15 16 17 18 19 20 21 22 X Y
P28
Day 180
Day 14
Day 0

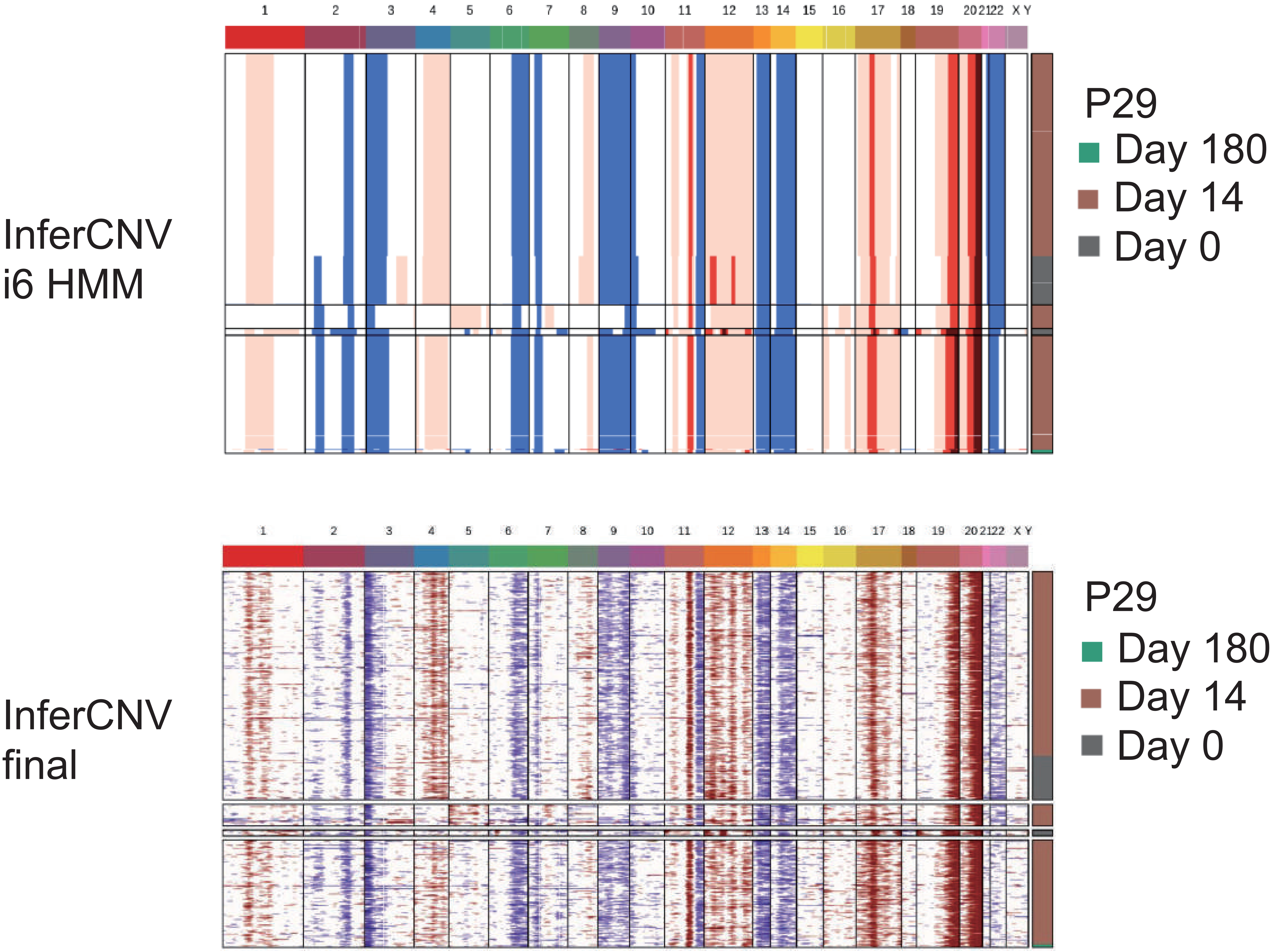
InferCNV
i6 HMM
InferCNV
final
1 2 3 4 5 6 7 8 9 10 11 12 13 14 15 16 17 18 19 20 21 22 X Y
P29
Day 180
Day 14
Day 0

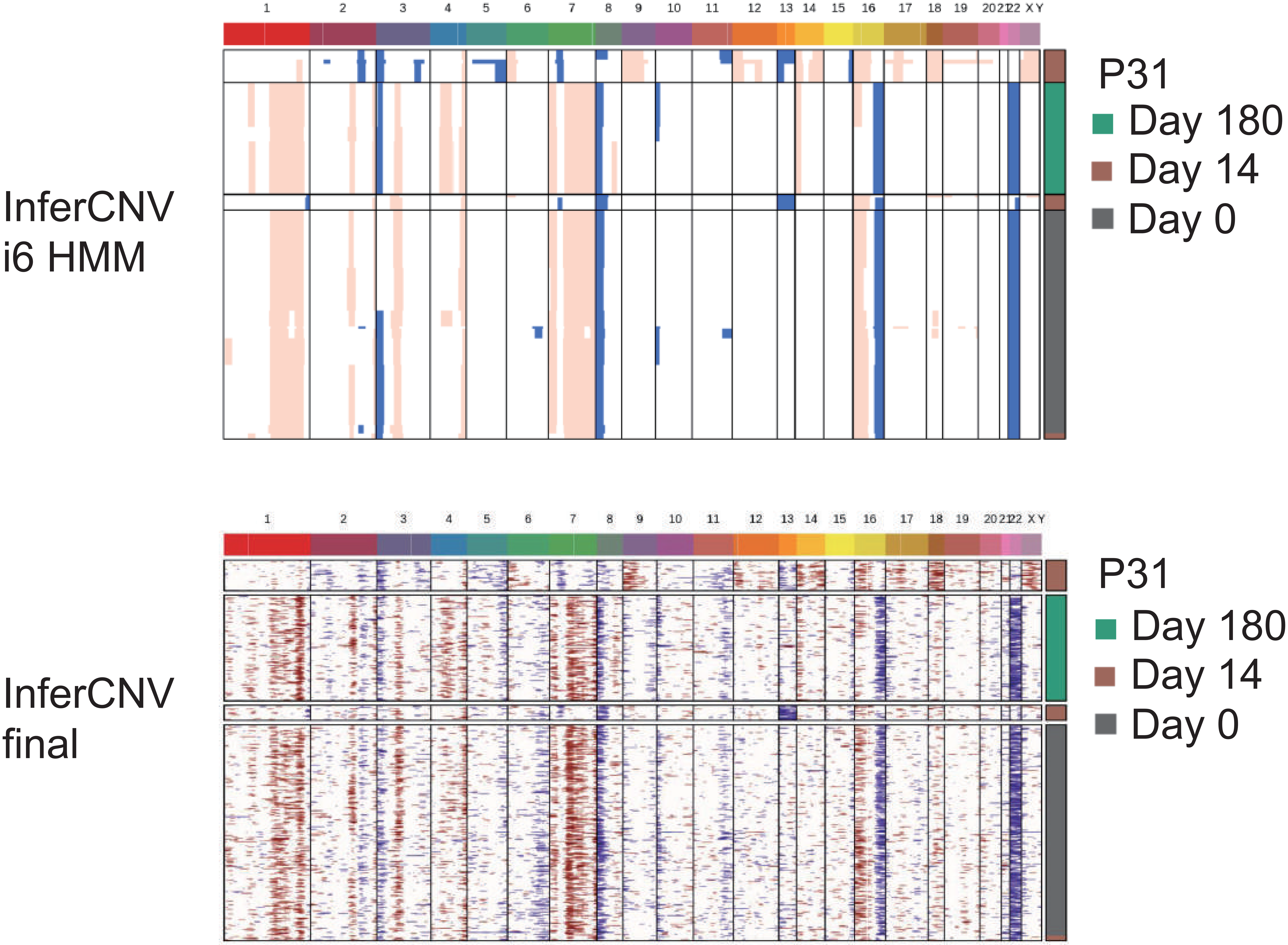
1 2 3 4 5 6 7 8 9 10 11 12 13 14 15 16 17 18 19 20 21 22 X Y
P31
Day 180
Day 14
Day 0
InferCNV
i6 HMM
1 2 3 4 5 6 7 8 9 10 11 12 13 14 15 16 17 18 19 20 21 22 X Y
P31
Day 180
Day 14
Day 0
InferCNV
final

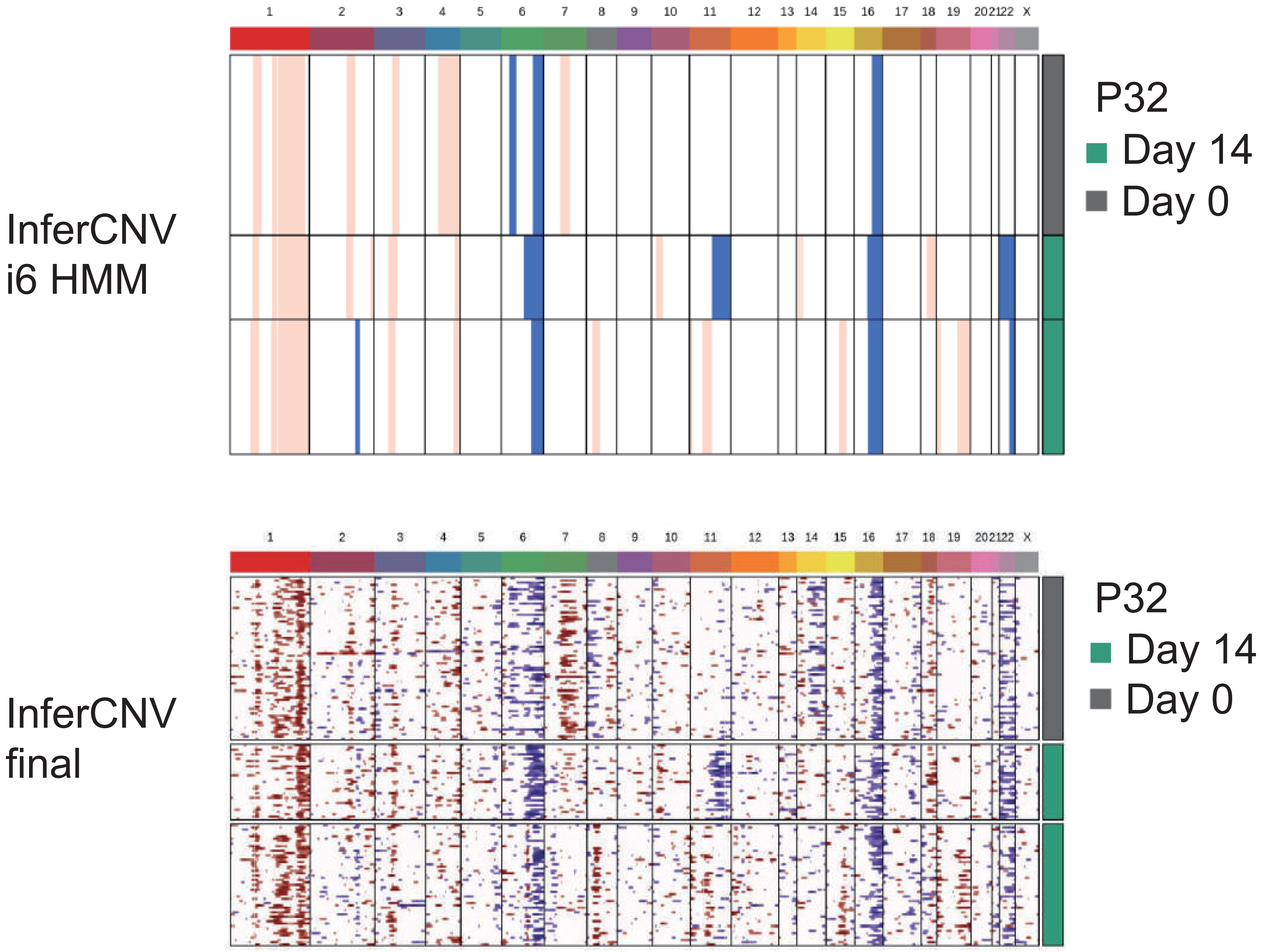
InferCNV
i6 HMM
InferCNV
final
P32
Day 14
Day 0
1 2 3 4 5 6 7 8 9 10 11 12 13 14 15 16 17 18 19 20 21 22 X

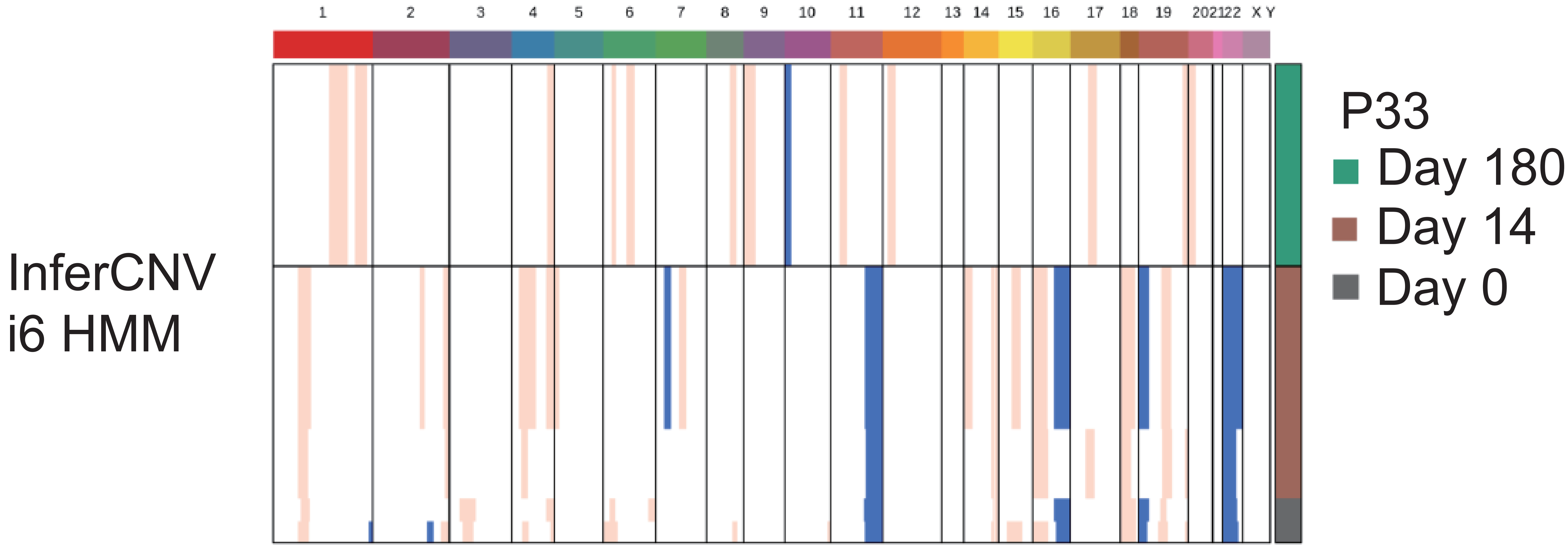
InferCNV
i6 HMM
1 2 3 4 5 6 7 8 9 10 11 12 13 14 15 16 17 18 19 202122 X Y
P33
Day 180
Day 14
Day 0

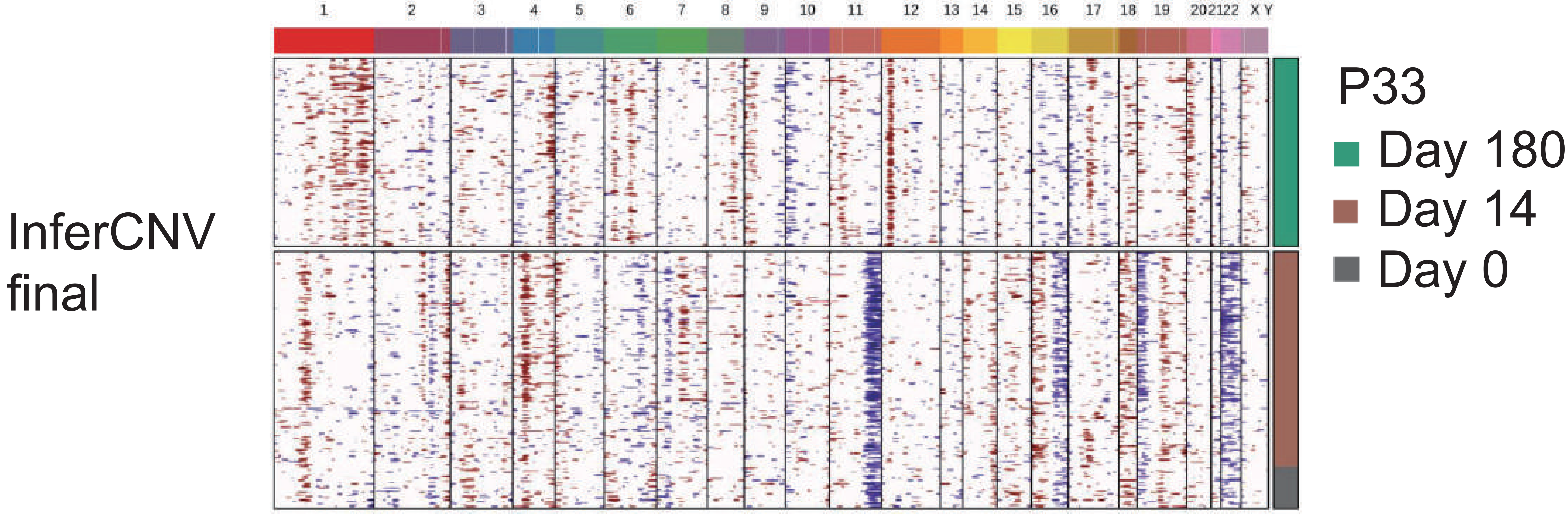
InferCNV
final
1 2 3 4 5 6 7 8 9 10 11 12 13 14 15 16 17 18 19 202122 X Y
P33
Day 180
Day 14
Day 0

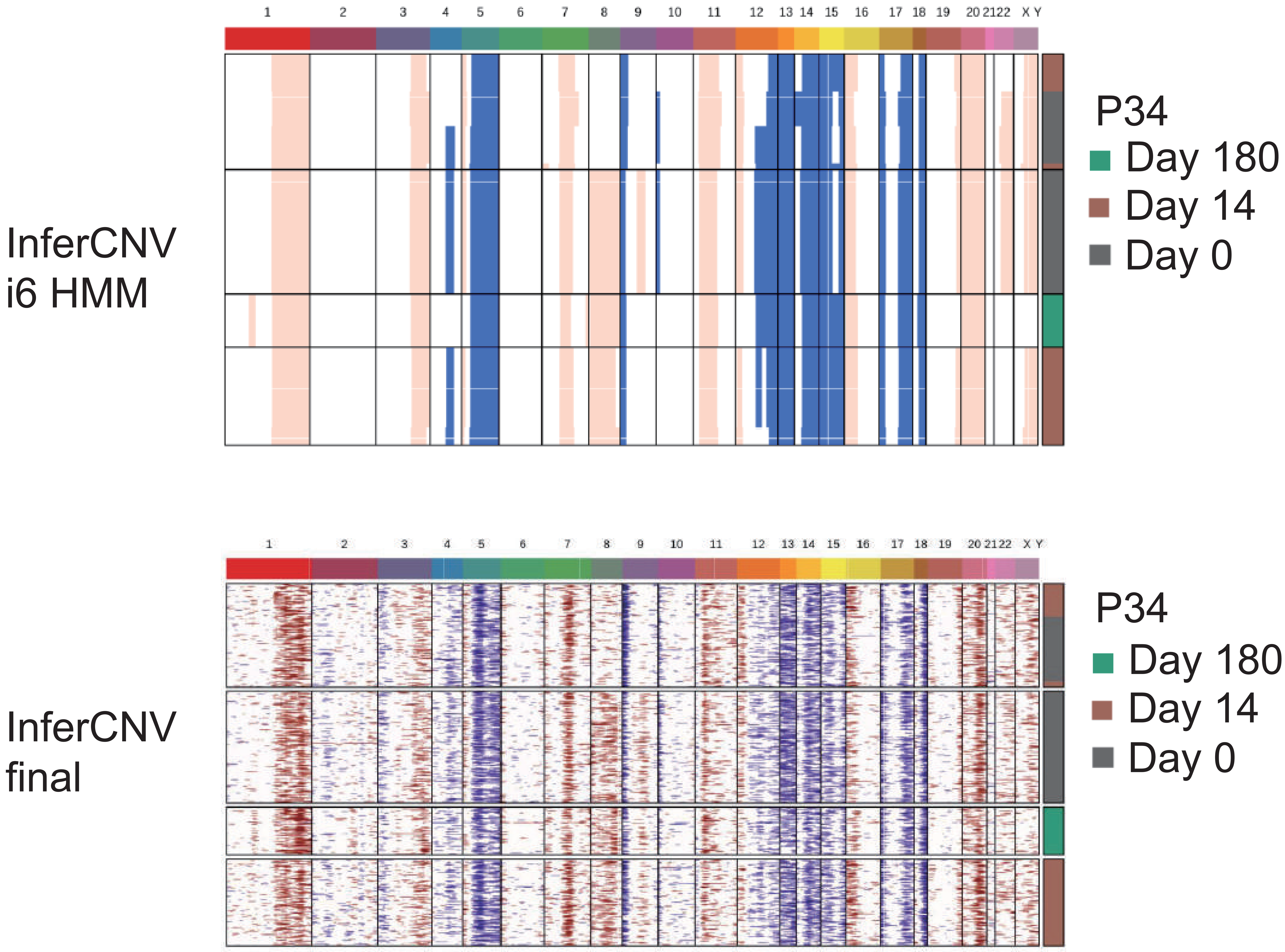

InferCNV
i6 HMM
1 2 3 4 5 6 7 8 9 10 11 12 13 14 15 16 17 18 19 20 21 22 X Y
P34
Day 180
Day 14
Day 0
InferCNV
final
1 2 3 4 5 6 7 8 9 10 11 12 13 14 15 16 17 18 19 20 21 22 X Y
P34
Day 180
Day 14
Day 0

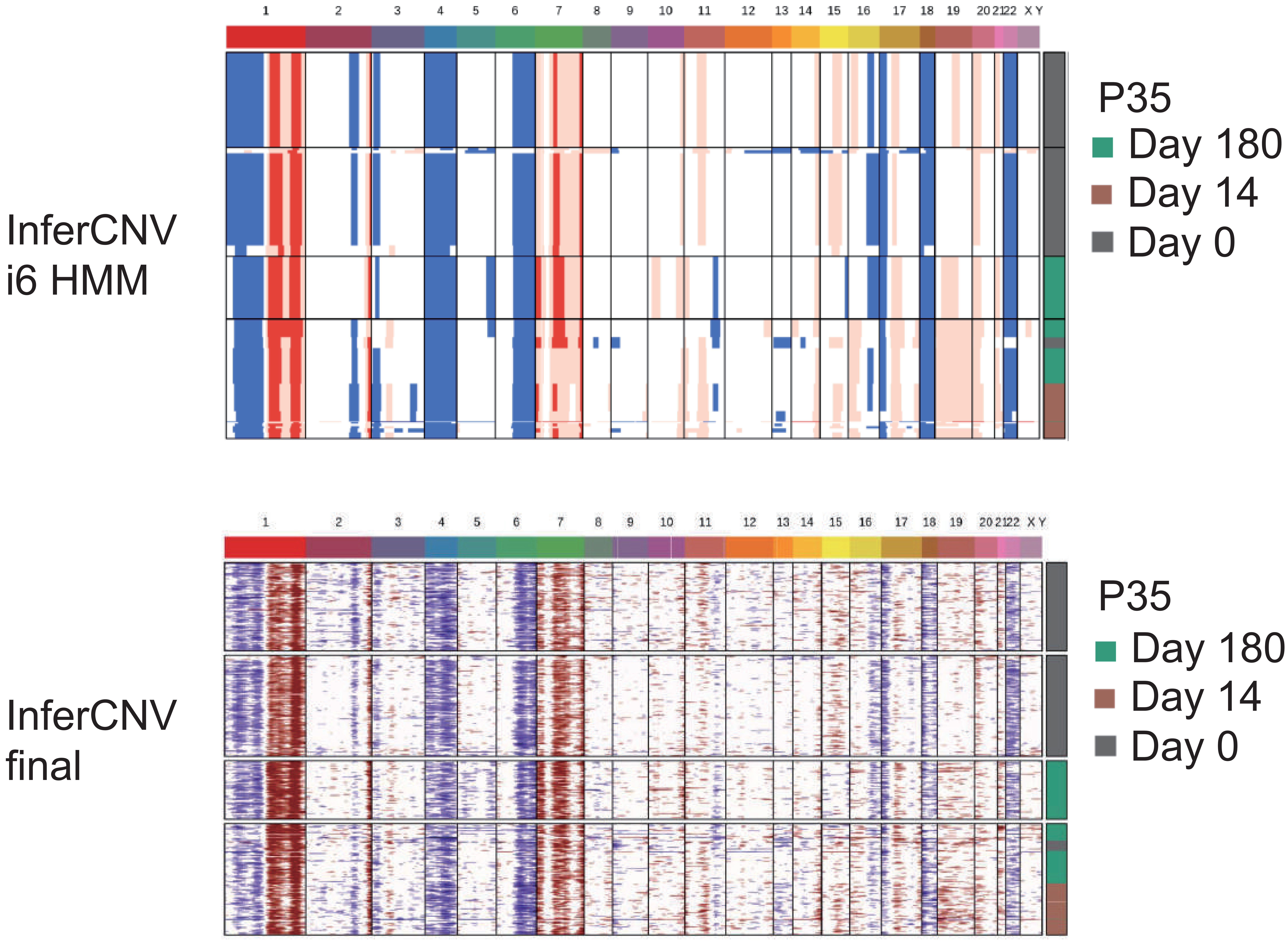

InferCNV
i6 HMM
1 2 3 4 5 6 7 8 9 10 11 12 13 14 15 16 17 18 19 20 21 22 X Y
P35
Day 180
Day 14
Day 0
InferCNV
final
1 2 3 4 5 6 7 8 9 10 11 12 13 14 15 16 17 18 19 20 21 22 X Y
P35
Day 180
Day 14
Day 0

InferCNV i6 HMM

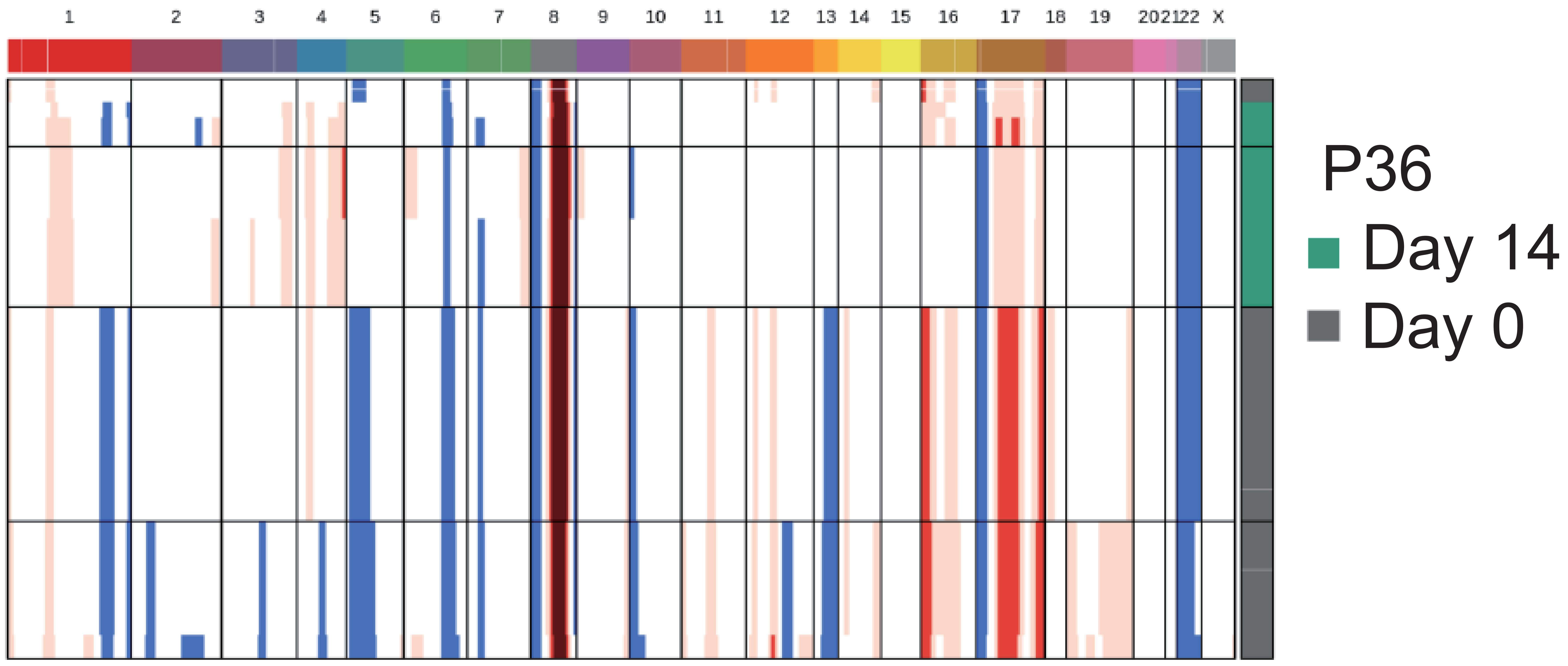

InferCNV final

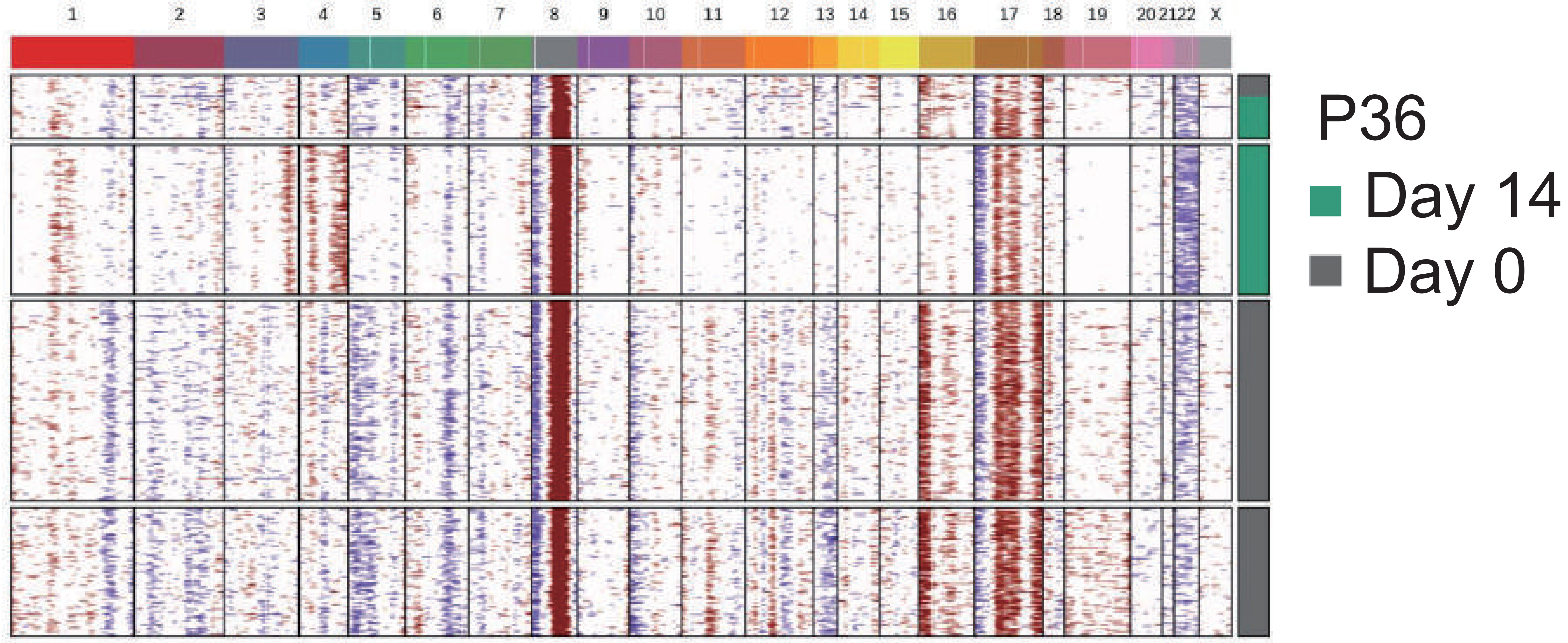

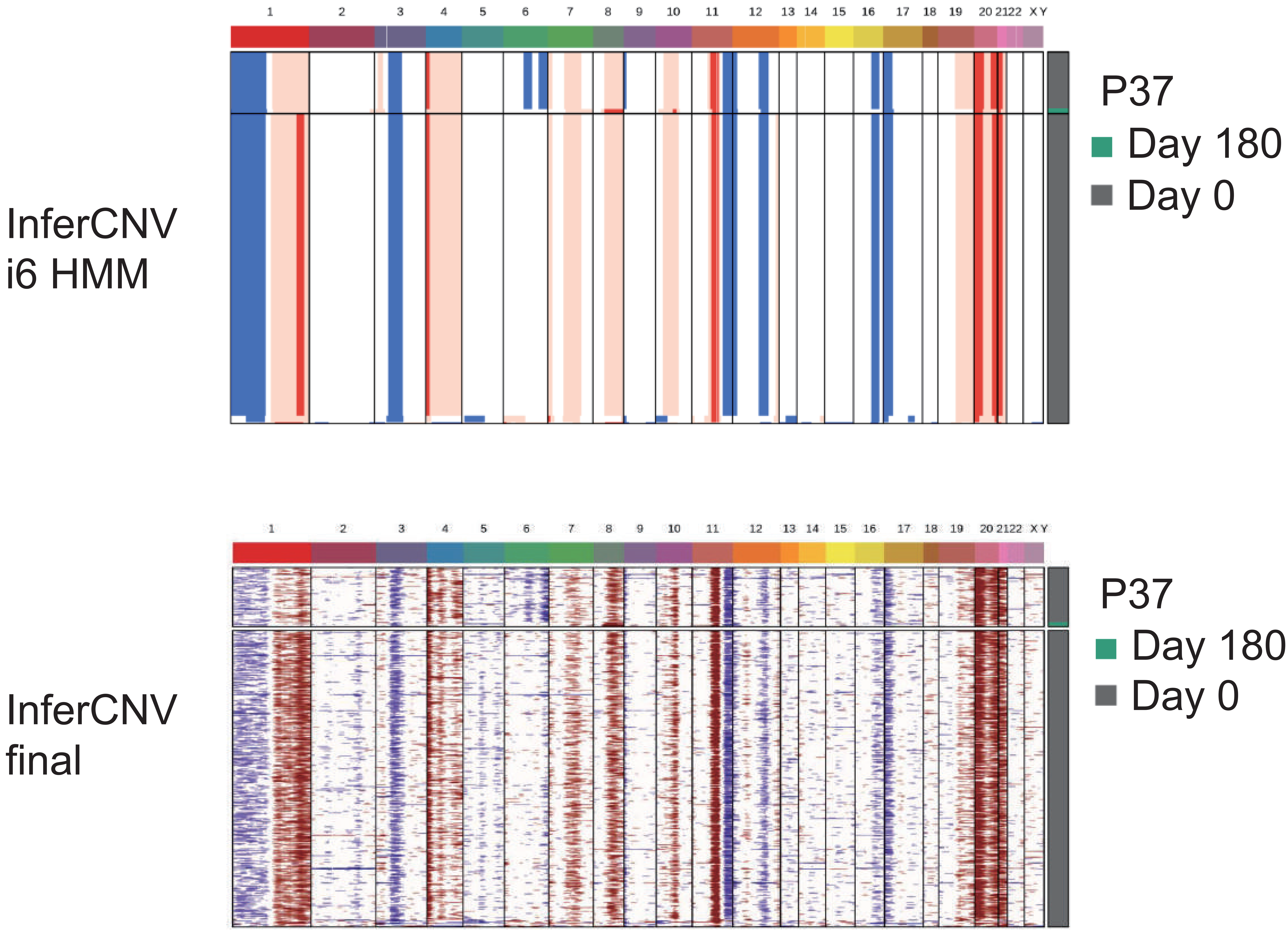
InferCNV
i6 HMM
InferCNV
final
P37
Day 180
Day 0
X
Y

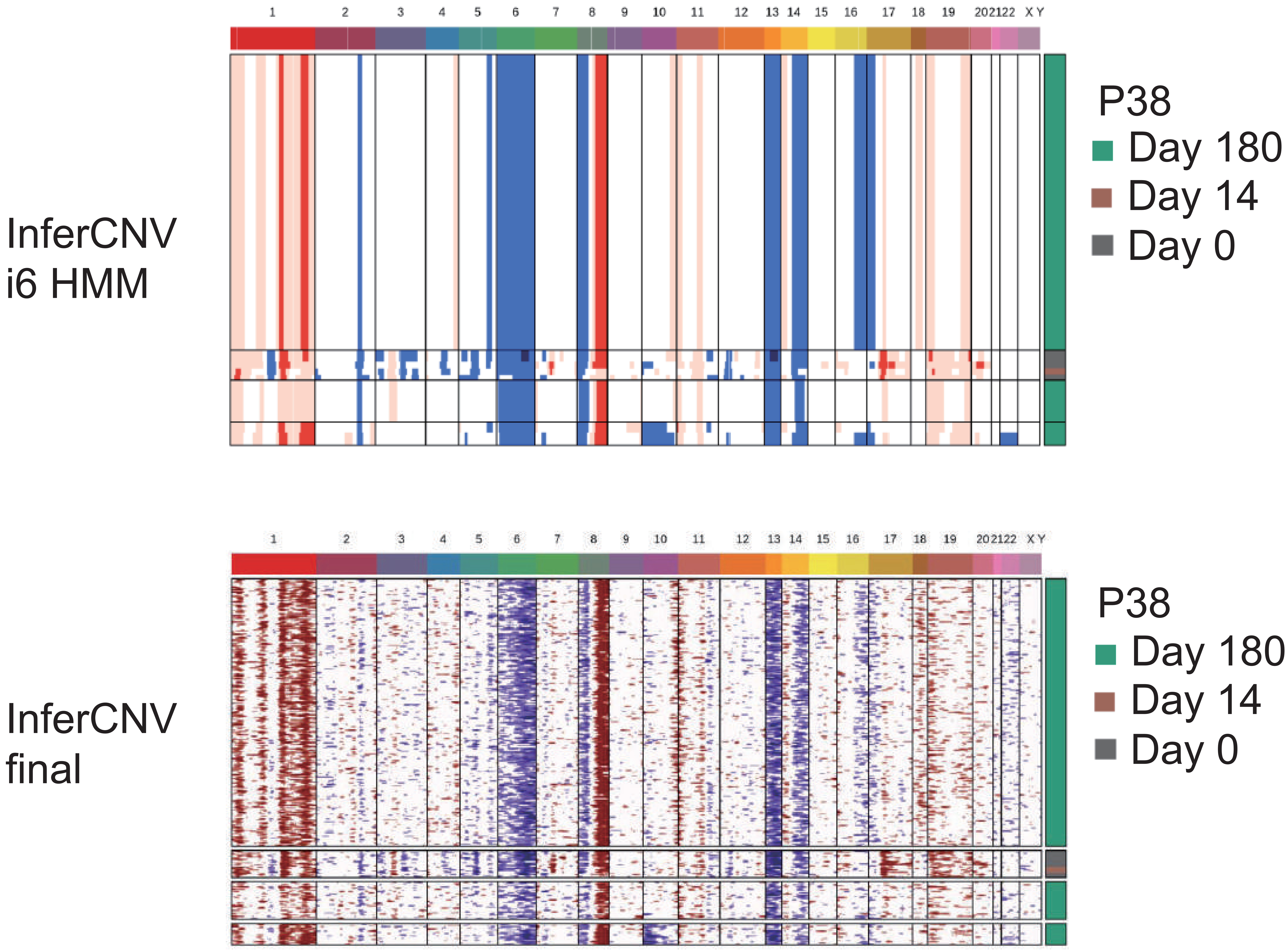
InferCNV
i6 HMM
P38
Day 180
Day 14
Day 0
1 2 3 4 5 6 7 8 9 10 11 12 13 14 15 16 17 18 19 20 21 22 X Y
InferCNV
final
P38
Day 180
Day 14
Day 0
1 2 3 4 5 6 7 8 9 10 11 12 13 14 15 16 17 18 19 20 21 22 X Y

InferCNV
i6 HMM

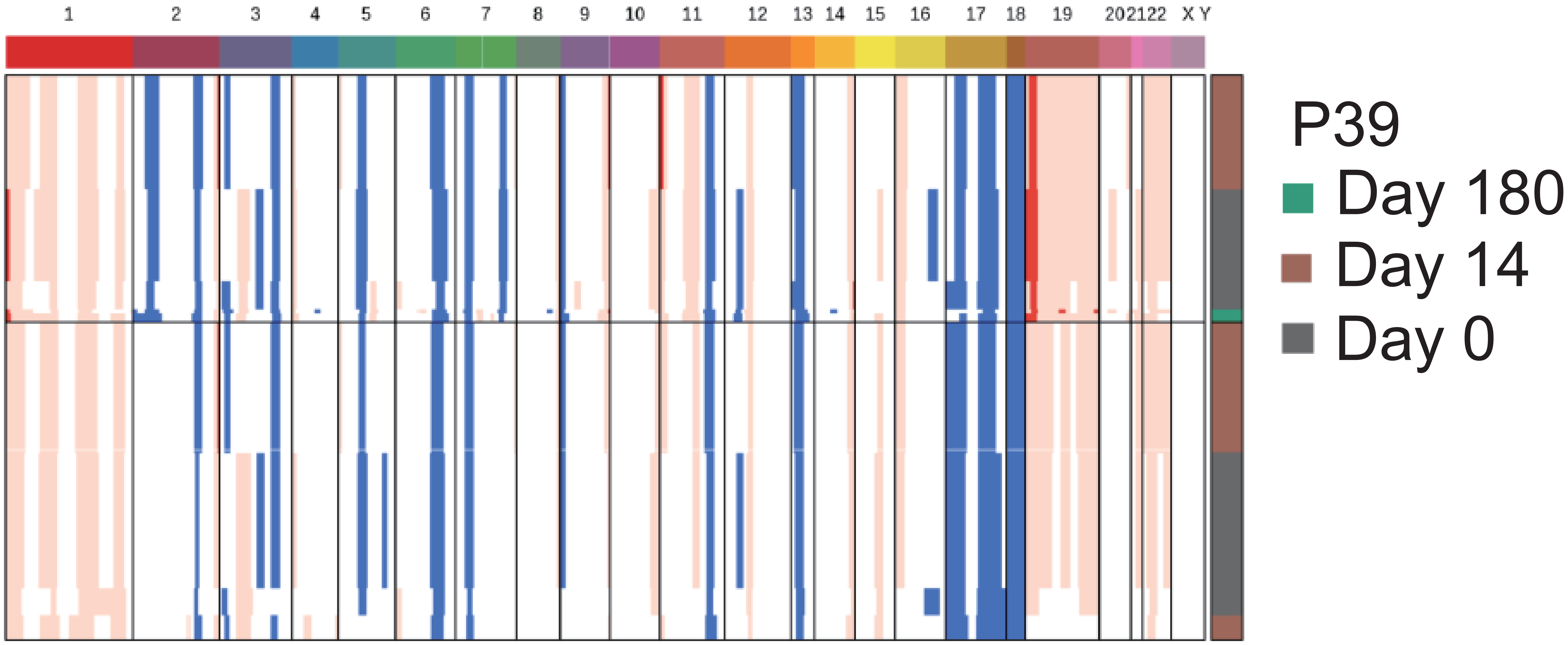

InferCNV
final

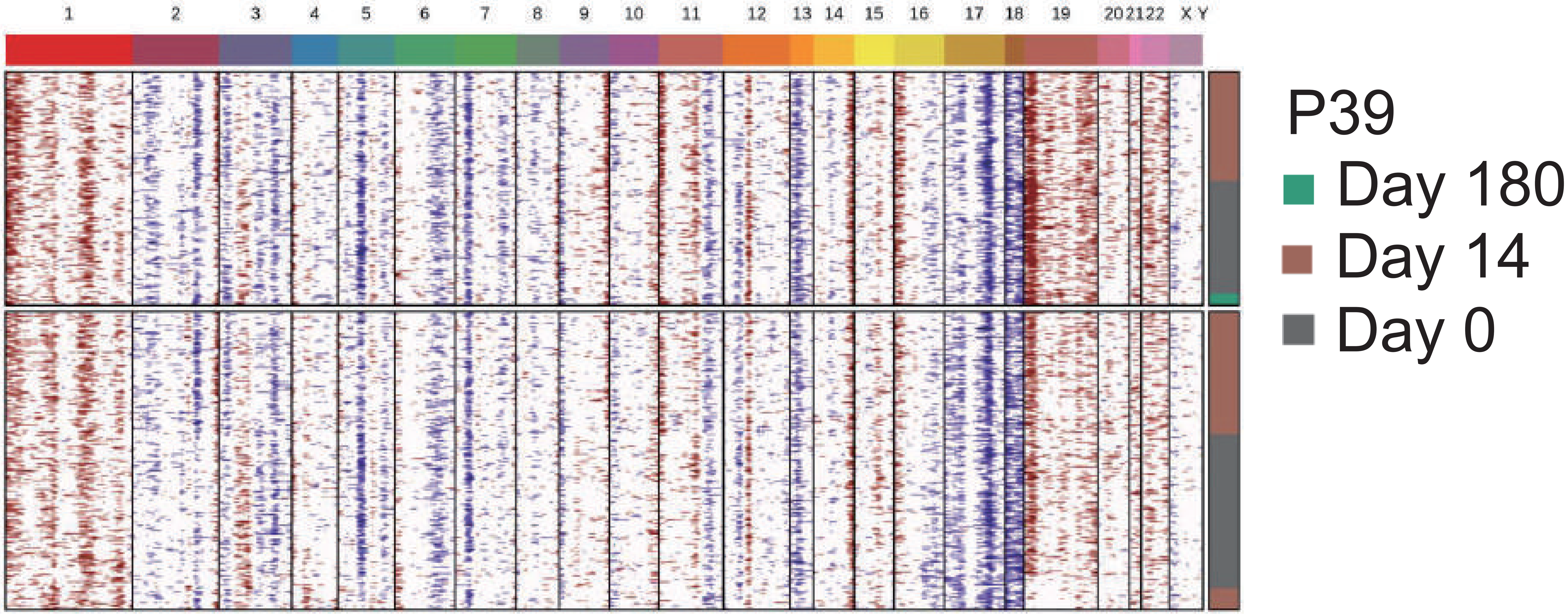

InferCNV
i6 HMM

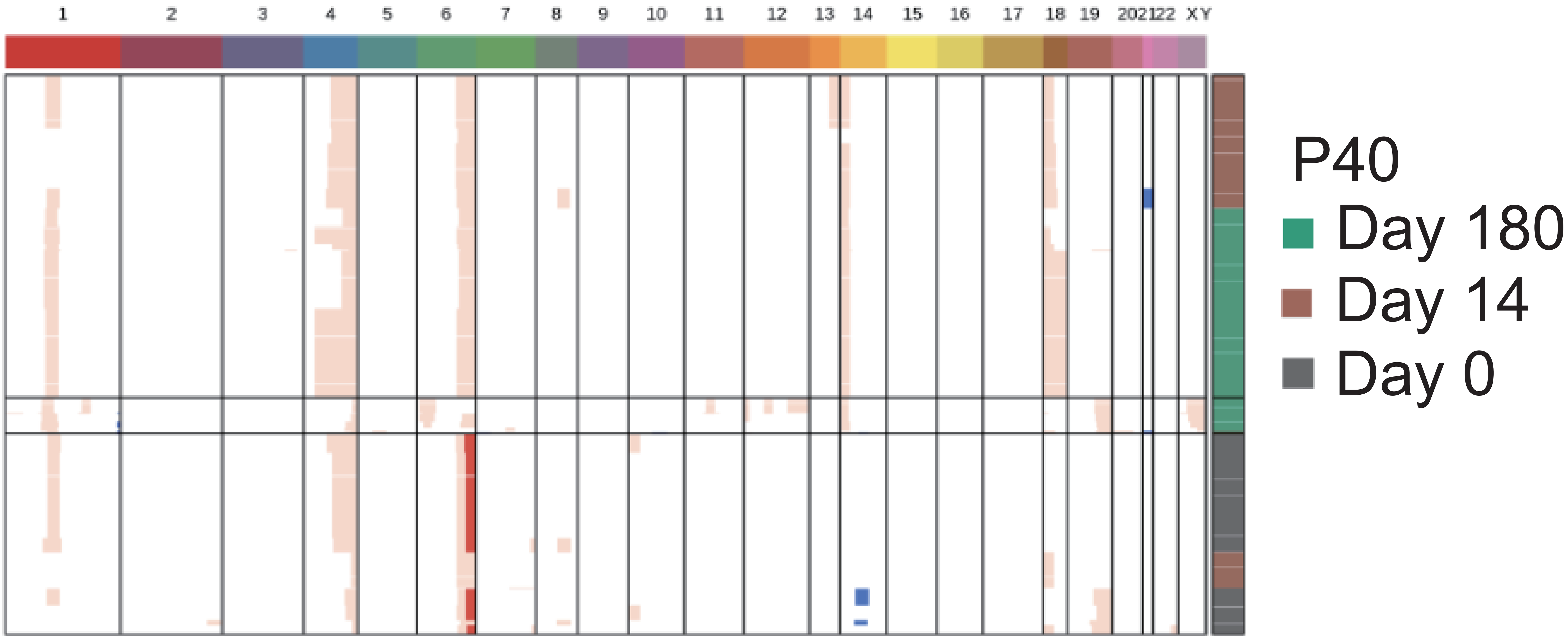

InferCNV
final

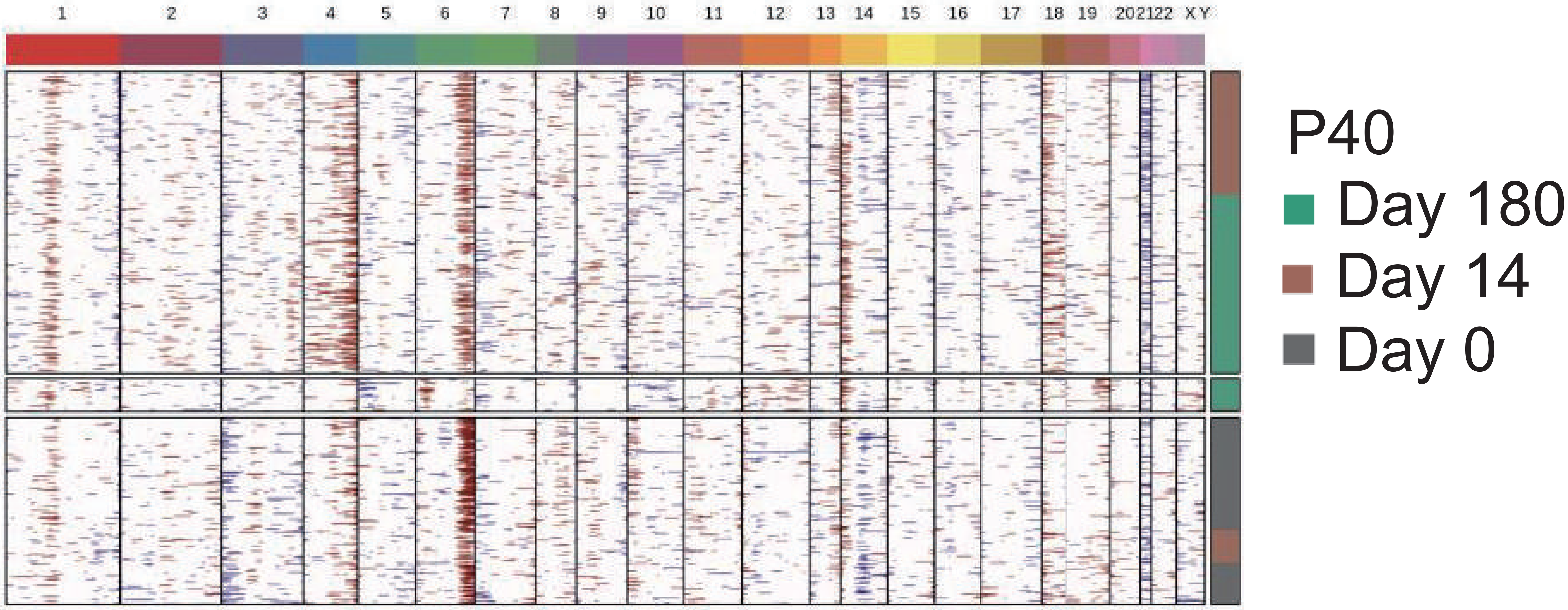

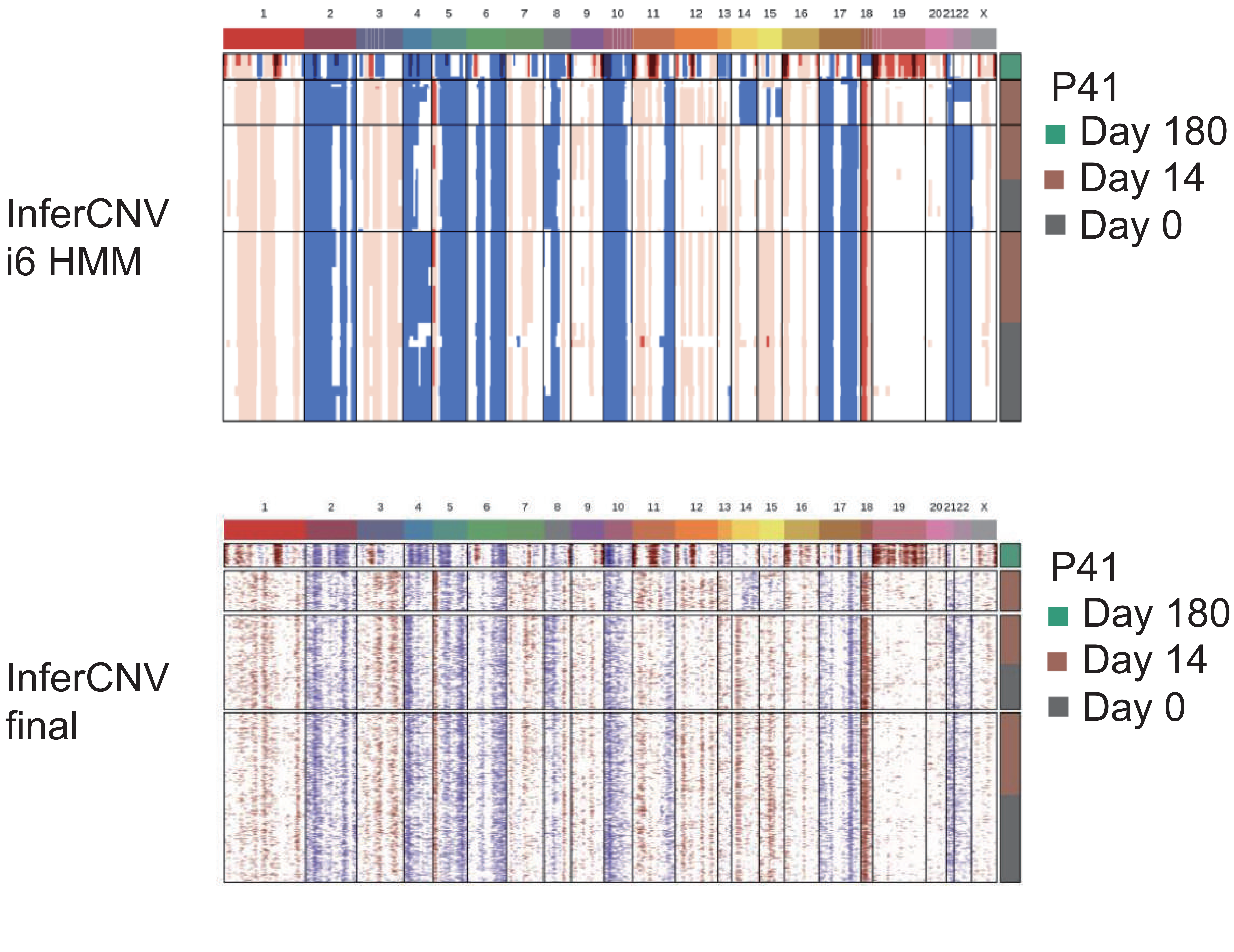

InferCNV
i6 HMM
1 2 3 4 5 6 7 8 9 10 11 12 13 14 15 16 17 18 19 20 21 22 X
P41
Day 180
Day 14
Day 0
InferCNV
final
1 2 3 4 5 6 7 8 9 10 11 12 13 14 15 16 17 18 19 20 21 22 X
P41
Day 180
Day 14
Day 0

InferCNV
i6 HMM

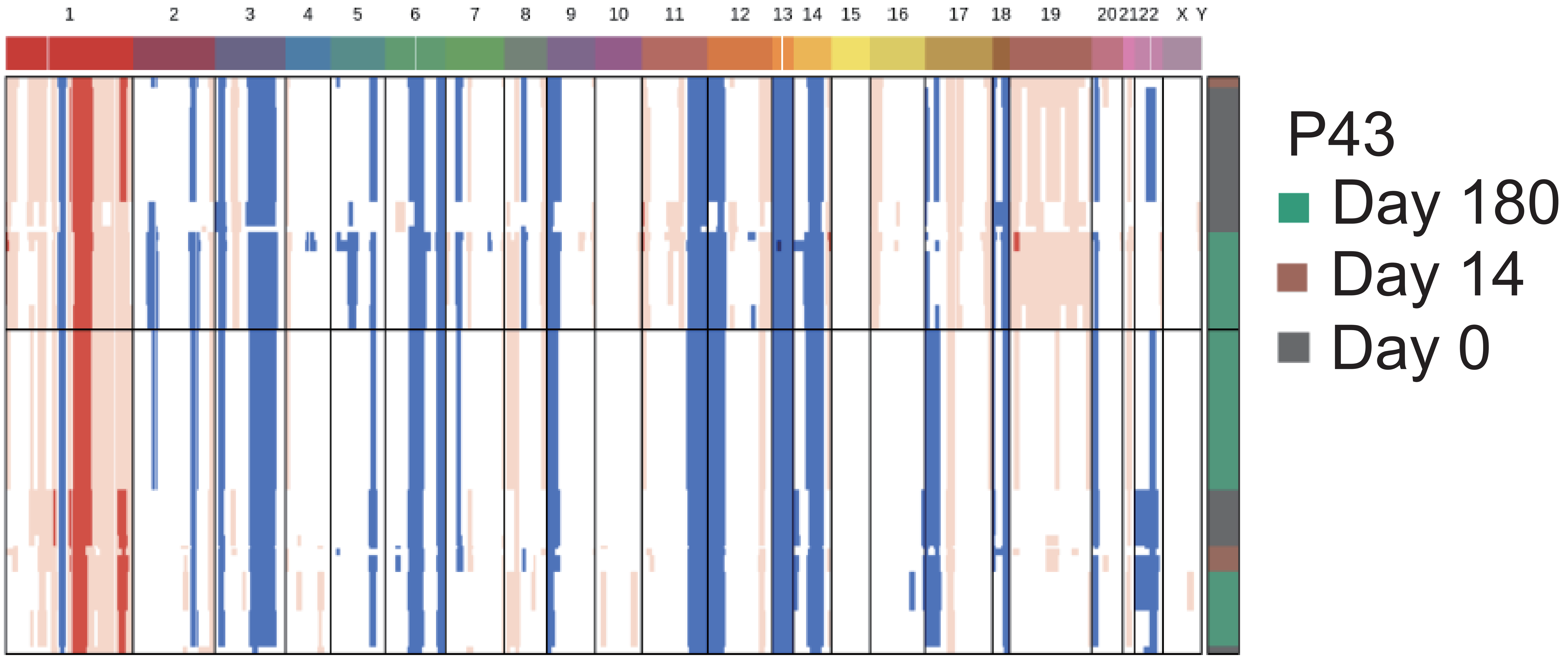

InferCNV
final

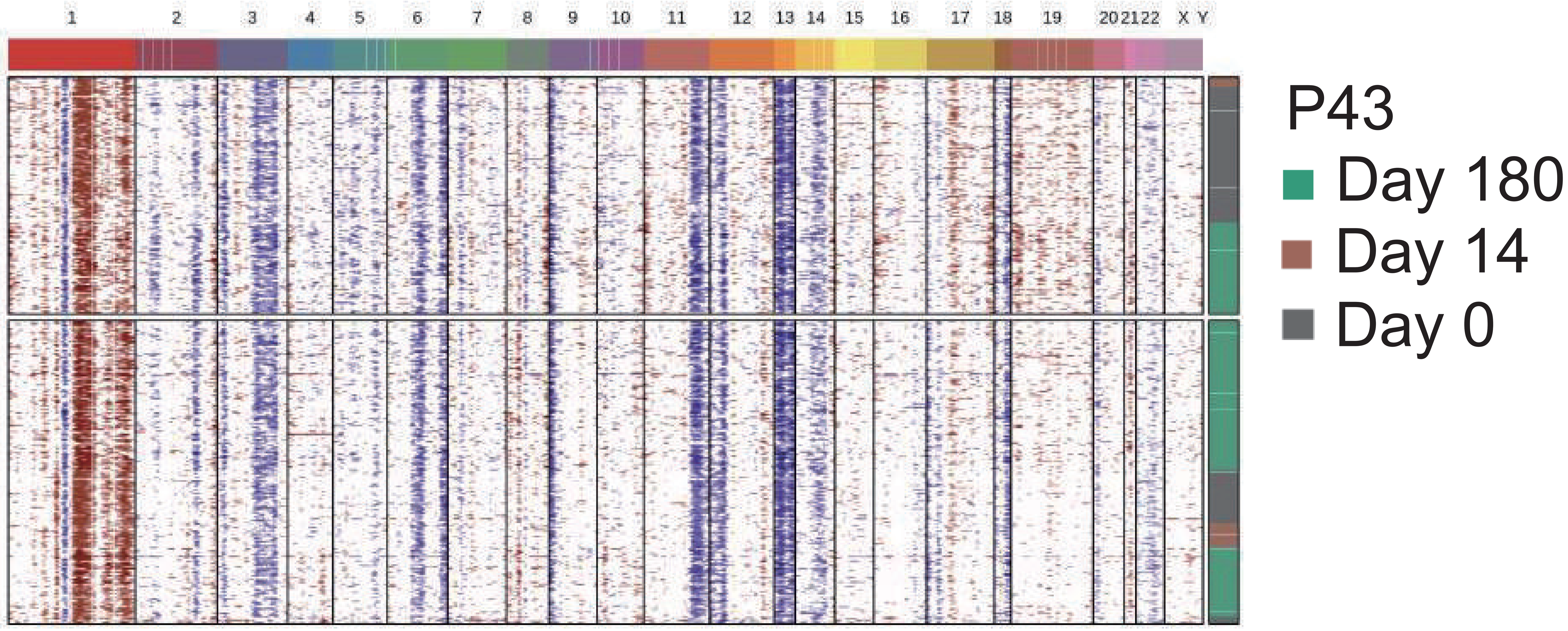

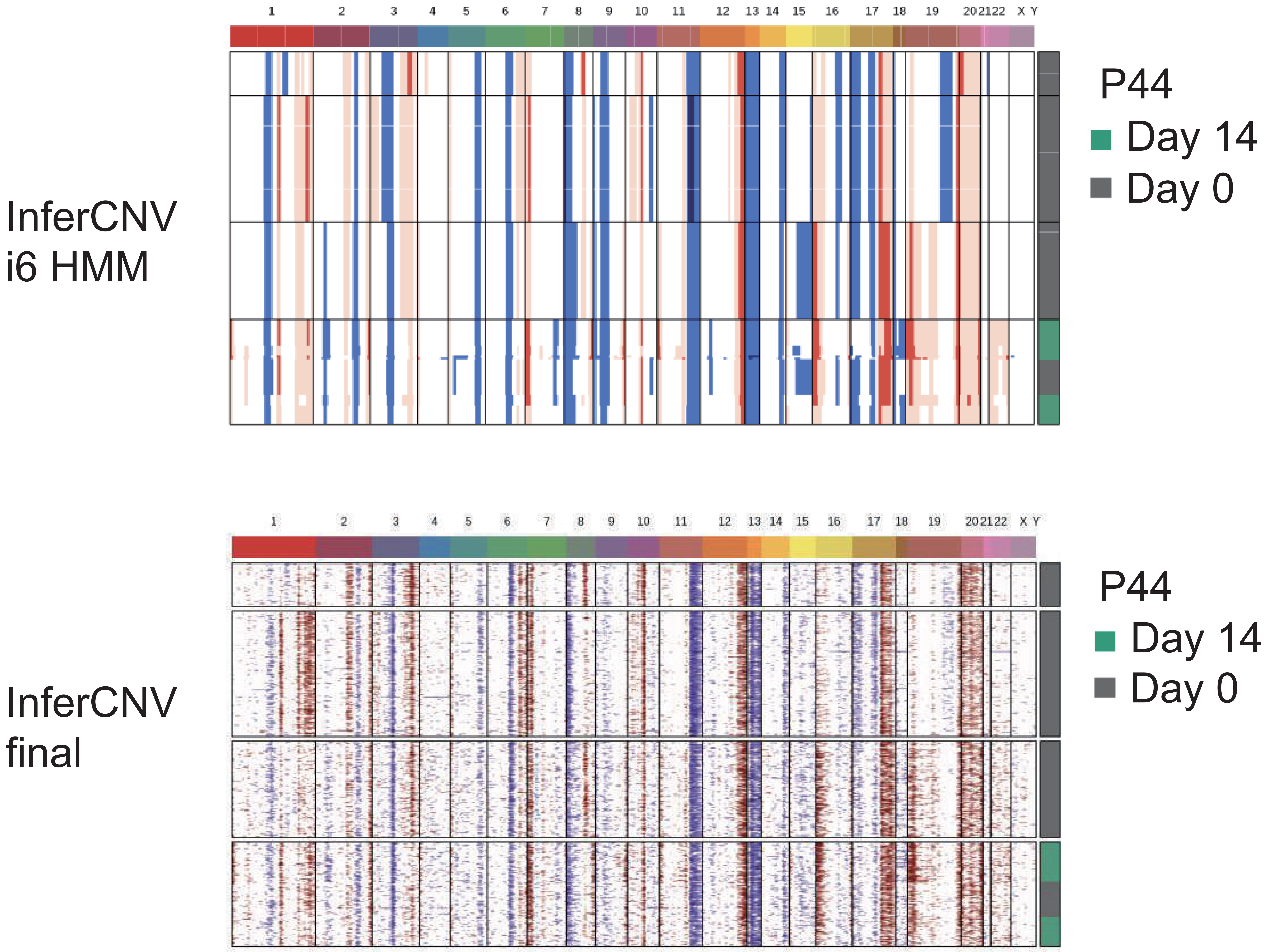
InferCNV
i6 HMM
1 2 3 4 5 6 7 8 9 10 11 12 13 14 15 16 17 18 19 20 21 22 X Y
P44
Day 14
Day 0
InferCNV
final
1 2 3 4 5 6 7 8 9 10 11 12 13 14 15 16 17 18 19 20 21 22 X Y
P44
Day 14
Day 0

InferCNV
i6 HMM

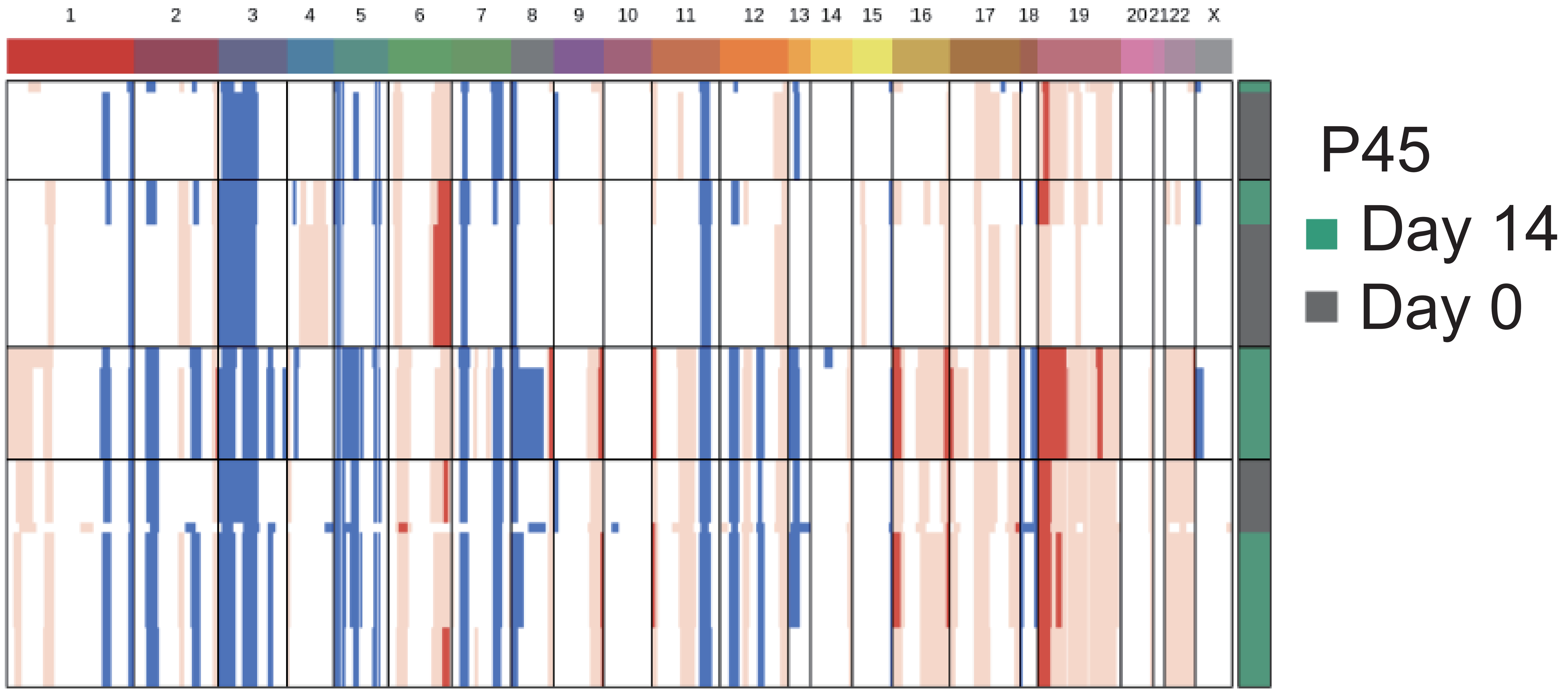

InferCNV
final

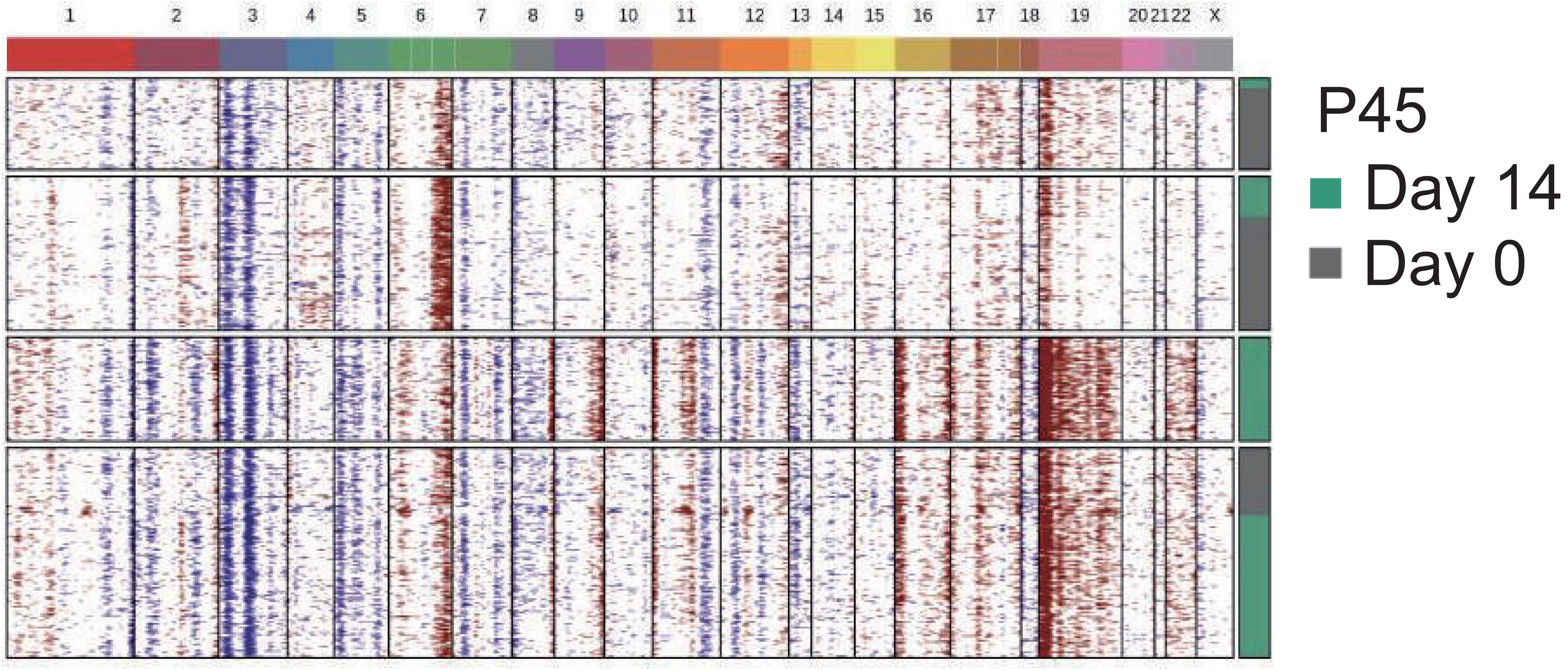

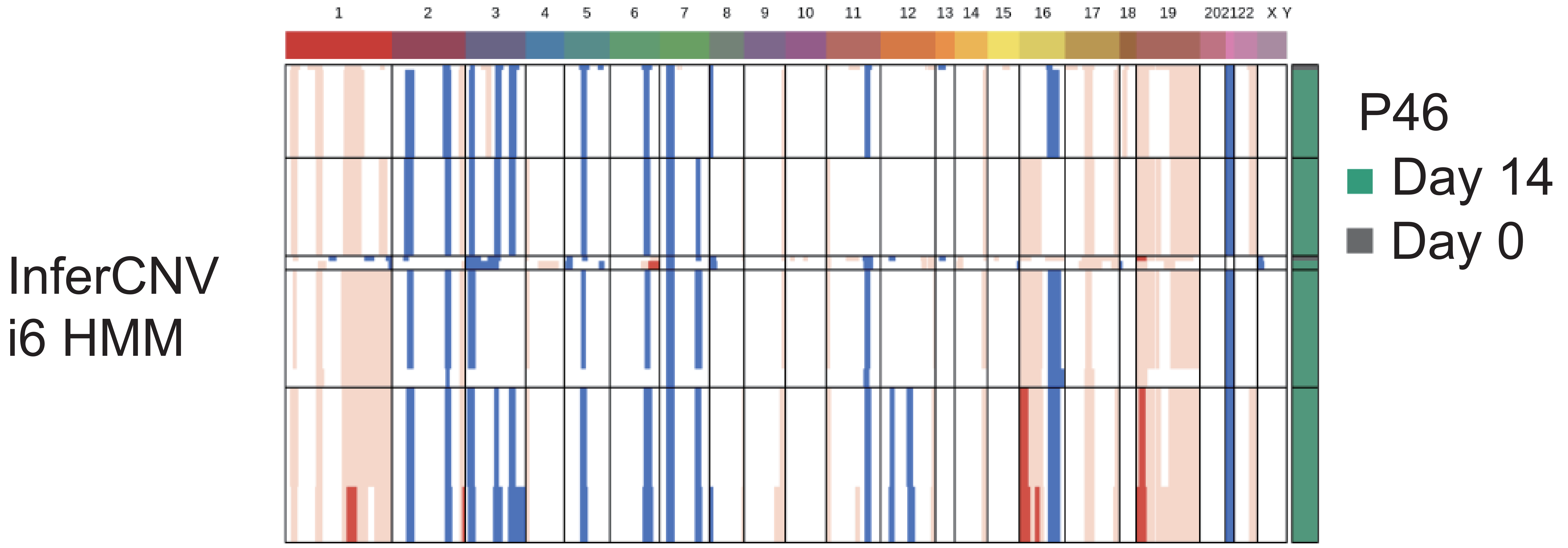
InferCNV
i6 HMM
1 2 3 4 5 6 7 8 9 10 11 12 13 14 15 16 17 18 19 20 21 22 X Y
P46
Day 14
Day 0

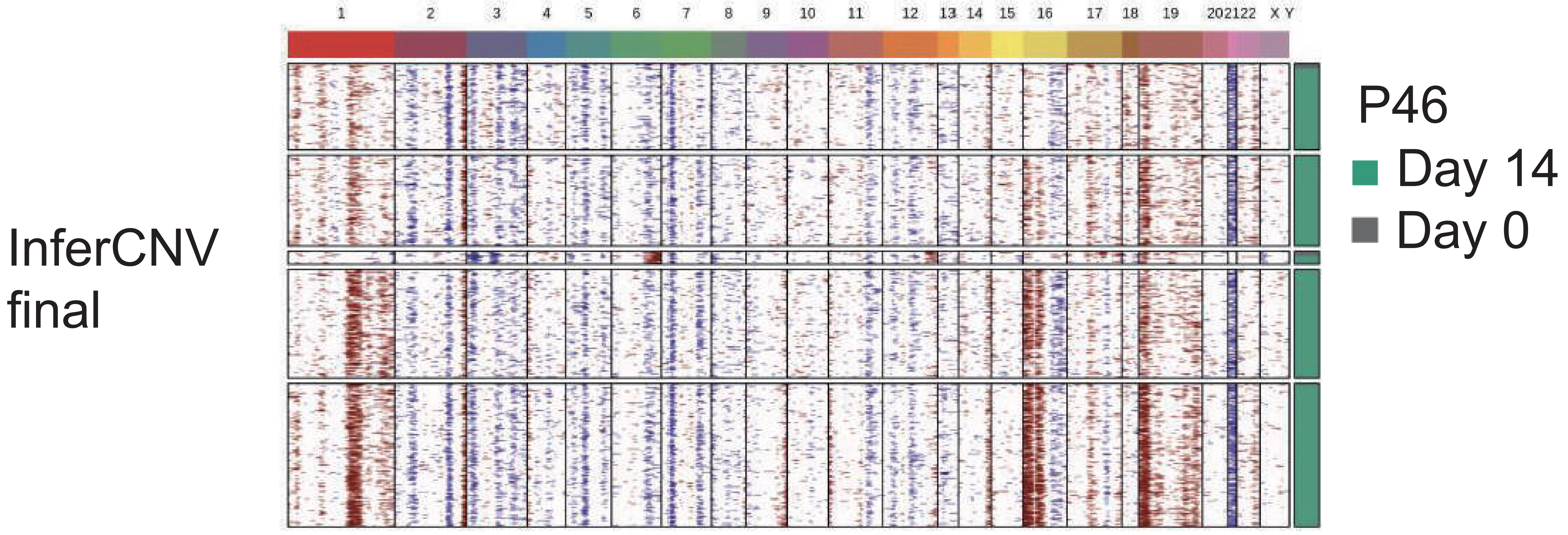
InferCNV
final
1 2 3 4 5 6 7 8 9 10 11 12 13 14 15 16 17 18 19 20 21 22 X Y
P46
Day 14
Day 0
